# MARLOWE: Taxonomic Characterization of Unknown Samples for Forensics Using *De Novo* Peptide Identification

**DOI:** 10.1101/2024.09.30.615220

**Authors:** Sarah C. Jenson, Fanny Chu, Gelio Alves, Aleksey Y. Ogurtsov, Anthony S. Barente, Dustin L. Crockett, Natalie C. Lamar, Eric D. Merkley, Yi-Kuo Yu, Kristin H. Jarman

## Abstract

We present a computational tool, MARLOWE, for source organism characterization of unknown, forensic biological samples. The intent of MARLOWE is to address a gap in applying proteomics data analysis to forensic applications. MARLOWE produces a list of potential source organisms given confident peptide tags derived from *de novo* peptide sequencing and a statistical approach to assign peptides to organisms in a probabilistic manner, based on a broad sequence database. In this way, the algorithm assumes no *a priori* knowledge of potential sources, and the probabilistic way peptides are taxonomically assigned and then scored enables results to be unbiased (within the constraints of the sequence database). In a proof-of-concept study, we examined MARLOWE’s performance on two datasets, the Biodiversity dataset and the *Bacillus cereus* superspecies dataset. Not only did MARLOWE demonstrate successful characterization to true contributors in single source and binary mixtures in the Biodiversity dataset, but also provided sufficient specificity to distinguish species within a bacterial superspecies group. We also compared MARLOWE’s results to those of MiCId, a leading microbial identification/characterization tool based on proteomics database search. Comparison of the two tools using 225 mass spectrometry data files yielded comparable performance, with slightly higher accuracy and specificity for MiCId. At the species level, MARLOWE achieved a specificity of 91.4% at 5% FDR. These results suggest that MARLOWE is suitable for candidate- or lead-generation identification of single-organism and binary samples that can generate forensic leads and aid in selecting appropriate follow-on analyses in a forensic context.

## Introduction

The last ten years has seen an increase in the use of proteomics methods for forensic applications ^1,2^, including sex estimation in human remains using sex-specific forms of amelogenin in tooth enamel ^3^, identification of individuals by genetically variant peptides in hair proteins ^4,5^ or touch samples ^6^, biofluid identification ^7,8^, and protein toxin detection ^9,10^. This paper focuses on identification (taxonomic characterization) of unknown forensic samples using bottom-up proteomics, which is closely related to the identification of microorganisms motivated by clinical needs. It is also related to metaproteomics (proteomics analysis of microbial communities in environmental or clinical samples), but forensics samples are not always complex communities; seized commodities, wildlife remains, or organisms cultivated for illegal purposes may contain only one or a few species.

Forensic samples present special challenges that rarely occur in fundamental science research. If the organism that produced the sample is unknown, the protein sequence database that should be used to identify peptides and proteins in the sample is also unknown, which is effectively the same case as the study of an organism with an unsequenced genome. Analysis of unknown samples may be qualitative, quantitative, or both. Qualitative analysis seeks to identify or at least provide some taxonomic indication of the organism(s) present; quantitative analysis seeks to characterize the abundance (biomass fraction) of the various organisms present. Both qualitative and quantitative analyses can be important in various applications. But, in many forensic contexts, the emphasis is on qualitative analysis, because laws in various jurisdictions make any quantity of certain organisms or toxins illegal. In this paper, we focus primarily on qualitative analysis, and refer to this process as taxonomic characterization. In addition to forensics, taxonomic characterization is also relevant to clinical, environmental, and archaeological/cultural applications.^11^

Several software tools or algorithms for taxonomic characterization of unknown samples using bottom-up proteomics and a database search have been described, but this general approach requires careful consideration of database composition and shared peptide assignment. Relevant software tools include ABOID ^12^, TCUP ^13^, MiCId ^14–18^, phyloproteomics ^19,20^, a statistical approach described by Jarman et al. ^21^, and two approaches specific for species identification from bone samples by Rüther et al. ^22^ and Yang et al. ^23^. All these protocols focus on analysis of untargeted, data-dependent, bottom-up proteomics data using database search with either a broad, multispecies database (either as a single database, or as multiple parallel searches with individual organism databases). The database may focus on particular organisms (e.g., pathogens) depending on the application. After the database search, peptide-spectrum matches (PSMs) or peptide sequences are assigned to organisms, and potential organisms are scored based on that mapping. However, the variations on this general framework all face two problems: database creation, and how to account for peptides that are shared between organisms.

*Database Creation.* There are known issues with searching a very large sequence database.^24^ As the number of organisms’ proteomes included in the database grows, the search space increases, leading to a higher number of false positive PSMs based purely on statistics. To control for false positive PSMs via the false discovery rate, score thresholds are increased, which leads to decreased sensitivity. (It is worth mentioning that MiCId ^25^ uses a different statistical approach meant to address this issue.) Conversely, if the database is too small, the correct organism may not be present, leading to a failure to identify it. Worse, a narrow database can lead to an incorrect or unrealistically specific identification, such as apparently identifying a strain when only species is feasible. ^26,27^ All of the general tools mentioned here use a broad or even comprehensive database initially, such as all available microbial genomes (e.g., ABOID,^12^) or the NCBI non-redundant database. Phylopeptidomics starts with a broad database, then conducts a second-round search using only organisms detected in the first round. ^19^ The SPIN method ^22^, designed for identifying mammals via bone fragment proteomics, constructs a multispecies database of a small number of proteins found in mammalian bones. Clearly, there is no standardized approach for database creation; bespoke databases are created to suit each application and tool, but their composition needs to be carefully considered to avoid the issues with false positives and false negatives described above.

*Shared peptides*. The second issue centers on assignment of shared peptides, that is, peptides that appear in the protein sequences of multiple organisms. Peptides that are unique to a single taxon in the database necessarily indicate that taxon, or one that is not present in the database. The simplest approach is to use only unique peptides, but these may not exist, or may not be expressed or detected; this approach can severely limit the ability to identify source organisms. On the other hand, many more peptides are shared among organisms than are unique to a single organism. Can shared peptides instead augment the presence of unique peptides and help identify taxa, given that peptides that appear in multiple taxa tend to occur in related taxa?

Several algorithms and tools exploit information from shared peptides to inform taxa, in different ways. TCUP ^13^ and UniPept ^28^ use the lowest common ancestor algorithm, which assigns peptides to the most basal node of the taxonomic tree to which they are unique. After matching PSMs to a MySQL database, ABOID uses a combination of clustering and principal components analysis to determine the most likely organism. ^12^. MiCId ^14^ also clusters organisms that share peptides, and weights the calculated p-value for each peptide based on an inverse function of the number of clusters in which that peptide appears. Recent efforts to augment organism identification functionality in MiCId have focused on statistical and algorithm improvements ^15,16^, usability ^17^, and extensibility to different types of proteomics data ^18^.

Phylopeptidomics, using an iterative database creation and search approach that increases in specificity per iteration, leverages a mapping of mass spectra to taxa, called taxon-spectrum matches (TSMs), which is analogous to peptide-spectrum matches (PSMs). This approach has been used to identify pathogens in archeological bone samples ^29^ and to characterize the gut microbiome of COVID-19 patients ^19^. This same research group has also expanded phylopeptidomics to estimate the biomass of each species ^20^ by plotting TSMs for a hypothesized species as a function of the phylogenetic distance from a larger group of species and fitting a multiexponential decay function to the resulting curve. This is a unique way to derive useful taxonomic information from shared peptides and the amplitude parameters from this fit reflect the known biomass ratios. Finally, Jarman et al. ^21^ have created an approach that introduced strong peptides, that is, peptides that are not uniquely assigned to a single taxon, but that are strongly biased to appear in only a few taxa. Phylogenetic organism clusters produced using the shared and strong peptides concept approximates “supergenus” or “superspecies” groups, as an alternative to classical taxonomy (e.g., genus, species). Again, as with database creation, there is no standardized approach to leveraging shared peptide assignments to identify organisms, but all recognize the value of shared peptides to inform taxonomy from proteomics data.

In this paper, we introduce an approach that provides unique answers to these two questions. First, we attempt to decrease the reliance on sequence databases by deploying *de novo* peptide identification ^30–32^. Because *de novo* methods often lead to only partially correct peptides, we use only confident sequence regions (tags) to match to potential organisms. This approach separates at least the peptide identification step from the limitations of existing databases, although matching sequence tags to organisms still requires a database of peptide sequences. Several researchers in metaproteomics have explored the use of *de novo* peptide identifications in various ways ^33–35^; the current paper attempts to use *de novo* sequencing for unknown identification in a forensic context. We combine *de novo* sequencing with (1) an adaptation of the strong peptide concept and (2) a novel statistical procedure for assigning peptides to organisms that accounts for shared peptides. Here, our algorithm estimates the number of peptides detected in a sample that could belong to an organism or its nearest neighbors based on proteomic similarity even if that particular organism were absent from our database.

Briefly, our tool, which we call MARLOWE (after the fictional private investigator created by author Raymond Chandler, in honor of the intended investigative use of the tool), uses *de novo* peptide identification, tag extraction and filtering by peptide strength, assignment of tags to taxonomic sources, adjustment of tag counts for tags shared between organisms, and final scoring of source organisms using non-negative least squares regression. Rather than relying solely on unique peptides, this procedure appropriately leverages the information contained in shared peptides by using the peptide strength concept referenced above.

In the proof-of-concept examination of MARLOWE’s performance presented here, we find that MARLOWE correctly characterizes most contributors of microbial samples in single-source and binary mixtures at the species level. Further, it is also able to correctly characterize microbial species within the closely related bacterial group *Bacillus cereus* to the species level, demonstrating its specificity. Comparing MARLOWE’s results with those of MiCId on the same datasets with the same database, we find that both tools successfully point to the correct taxonomic sources. MiCId is slightly more accurate, but MARLOWE achieved >90% specificity at the species level. This finding is in line with the reduced accuracy of sequences derived from *de novo* peptide identification. These results demonstrate that MARLOWE is capable of preliminary, qualitative organism identification/taxonomic characterization of unknown protein-containing samples. As such, it fills an important gap in forensic proteomics.

## Methods

### The MARLOWE Algorithm

MARLOWE (Figure 1) uses the following steps: (1) *de novo* peptide identification, (2) tag extraction and peptide strength filtering, (3) tag matching to a broad sequence database, (4) tag assignment to organisms, including a correction for the number of tags expected due to shared peptides if the organism is not present, and (5) taxonomic group scoring. Each of these steps is described below, along with a detailed explanation of the concept of peptide strength.

**Figure 1.**
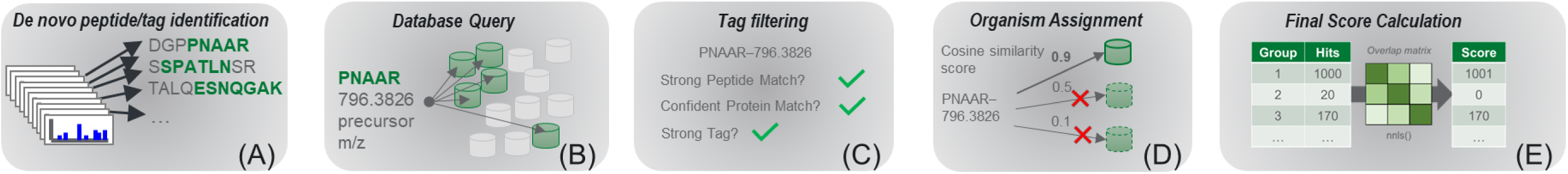
Workflow representation for the MARLOWE algorithm. (A) *De novo* peptide identification (in this paper we used Novor) produces partially correct sequences. High-confidence regions are extracted as tags. (B) Tag sequence and precursor mass information are used to query a broad sequence database stored as tryptic peptides linked to protein and organism accessions. (C) Tags are filtered for peptide strength, match to proteins, and tag strength. Tags that pass all of these filters proceed to the next step. (D) For each filtered tag, two vectors are constructed; first, a Boolean vector describing the presence or absence of that tag in each organism in the database; second, the overlap coefficient of an organism’s tryptic peptide set with every other organism in the database. This second vector is one row of the overlap matrix. The cosine similarity between the presence/absence vector for the current tag and each row of the overlap matrix is calculated, and summed across rows (i.e., organisms), and the highest value determines the organism to which the tag is assigned. (E) After assigning all tags to organisms, the count of tags per organisms becomes a raw score. This score is corrected for relatedness between organisms by means of a non-negative least squares function.

*De novo peptide identification.* With minor output file formatting, MARLOWE can accept input from PEAKS, ^36^ Novor,^37^ or Casanovo. ^38^ The only requirement is that the *de novo* tool provide (in addition to the usual PSM data) a local or per-amino acid residue confidence score that reflects the likelihood that an individual residue is correct. The listed tools all provide this local confidence score. In this paper, we used Novor.

*Tag extraction.* The accuracy of *de novo* PSMs is known to be lower than database search PSMs. However, *de novo* PSMs often contain correct subsequences ^39^ called tags, which have been used in various peptide identification schemes for a long time ^40,41^. In MARLOWE, we define a tag as a stretch of at least 5 consecutive amino acid residues in which each residue has a local confidence score of 80 (out of 100) or greater. Only tags from peptides where the overall PSM had an average local confidence score of 50 or greater were used; this ensured that only high-quality spectra were considered.

*Peptide strength filtering.* To optimize the taxonomic specificity available from *de novo* sequence tags, we calculate peptide strength for each and then filter them to retain only tags that match to peptides (strong tags). To understand the concept of strong peptides and their role in MARLOWE, it is important to keep the following points in mind:

1. Peptide strength is a way to avoid the limitations of “unique” peptides. Both genomics ^42^ and proteomics studies ^43,44^ have demonstrated that, as databases grow, sequences used as biomarkers that were once unique lose that uniqueness.
2. Peptide strength is dependent on the database being used, but far less dependent than peptide uniqueness. Jarman et al. showed that when new, related organisms are added to a sequence database, the number of unique peptides declines precipitously, but the peptide strength remains nearly constant. Thus, while peptide strength will change as databases are updated, we do not expect it to change dramatically, and peptide strength calculations from older databases will remain taxonomically informative for a long time.
3. Peptide strength is a way to leverage the information content of non-uniformly shared peptides. Strong peptides are shared peptides, but they are shared in a way that is biased across the taxonomic tree at a given level of taxonomy. Whereas unique peptides are specific to (for instance) a particular species, strong peptides are selective with respect to a group of related species. An accumulation of strong peptides can thus provide a very strong indication of the presence of one or more members of that group. This is very similar to the insight of Armengaud and coworkers ^20^ that shared peptides are informative in taxonomic characterization, and to the use of weighting factors assigned to shared peptides by Alves et al. in MiCId ^14^ (although MARLOWE uses a filtering rather than a weighting approach).

The peptide strength (ps) of a peptide sequence ρ is defined by Jarman et al. ^21^ as:

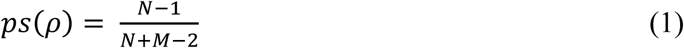

where N is the total number of taxa (at a given level) in the database, and M is the number of taxa in the database that contain peptide ρ. This equation is derived using the definitions of likelihood ratio and weight of evidence (statistical concepts often used in forensics), and expressions for the probability that peptide ρ occurs in organisms other than the one currently in question. Interested readers should consult the derivation given in the supplementary material of Jarman et al. ^21^ Peptide strength varies from ½ (for the case where N = M, meaning that peptide ρ is found in every taxon in the entire database) to unity (for the case where ρ occurs in only a single taxon, i.e., M = 1). Let Mc represent the critical number of taxa in which ρ appears. By setting a threshold value of 0.95 for the peptide strength and assuming N>>1, it follows from Eqn (1) that Mc = 0.053N, meaning that strong peptides occur in no more than 5.3% of the taxa at a given level in the database (5.3% is roughly the ratio 0.05/0.95). Importantly, as more organisms are sequenced and the database grows, as long as ρ appears in fewer than 5.3% of all taxa, it will remain a strong peptide.

This definition of peptide strength worked well for taxonomic characterization using database searching, but we found that using it with a broad sequence database resulted in nearly every peptide being classified as a strong peptide. To achieve more stringent filtering, we further defined a *pairwise* peptide strength as follows. (This description counts species in genera, but an analogous definition can be used for genera within a family or any other similar combination of adjacent taxonomic levels.) For each genus that contains ρ, the number of species in that genus that contain ρ is determined (analogous to M in Equation 1) and divided by the total number of species in that genus (analogous to N). We call this ratio a hit fraction. The genera with the largest and second-largest hit fractions are then identified. If the ratio of the largest and second-largest hit fractions is greater than 1/0.053, then the peptide is defined as strong. This procedure is equivalent to stating that a peptide is strong if it exists in 95% or more of the species in the genus where it occurs most often, and 5% or fewer of the species in the genus where it occurs the second most often. The pairwise peptide strength concept can be applied to tags as well.

*Tag matching to a broad sequence database.* To interpret *de novo* peptide identifications as evidence for the presence of particular taxa, *de novo* tags must be matched to a broad sequence database. In this proof-of-concept study, we have employed the KEGG Genomes database (release 91.0, https://www.genome.jp/kegg/genome/, downloaded July 1, 2019). All protein sequences (viral genomes were removed because the small size of viral proteomes would impact taxonomic clustering—see below) from this source were *in silico*- digested with trypsin and stored in a MySQL database. Tags extracted from Novor *de novo* sequencing output were used to query this database and retrieve peptide, protein, organism, and taxonomy tree (lineage) information. The query involves matching the overall mass of the *de novo* peptide—which includes the confident and unconfident residues—and the confident tag sequence. For *de novo* peptides with post translational modifications, the overall mass was adjusted to correspond to the mass of the peptide without modifications before it was used to query the database. The database was optimized to speed execution of these queries.

*Tag assignment to taxonomic groups.* To avoid the confusion created by the non-phylogenetic nature of formal taxonomy, we group all species in the KEGG database into phylogenetic taxonomic groups based on shared tryptic peptides, which we refer to as taxonomic groups. To accomplish this grouping, we first predict all tryptic peptides for each organism, then create a symmetric matrix (the overlap matrix) consisting of the Jaccard coefficients (overlap coefficients) for the sets of tryptic peptides of each pair of organisms. Taxonomic groups are then formed by clustering using the leader algorithm and Euclidean distance based on the overlap coefficients, with a radius of 0.5.

Tags extracted from *de novo* peptide identification results are first matched to the sequence database to identify those that could belong to genus-level strong peptides. The tag list is then further filtered for tags that match to proteins with at least two tags. Next, the list of tags is filtered for strong tags (not peptides) with respect to the clustered taxonomic groups. This final list of tags is the input for the assignment to taxonomic groups/organisms.

Outside of the peptide strength calculation and after filtering, we assign peptide tags to organisms in the following manner. Specifically, let *Ni* be the actual (true) number of peptides contributed by organism *i* in an unknown sample. The expected number of observed counts for organism *i*, *ei*, can be approximated as

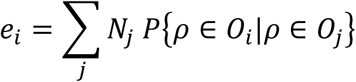

where 𝑃𝑃𝜌𝜌 ∈ 𝑂𝑂_𝑖𝑖_|𝜌𝜌 ∈ 𝑂𝑂_𝑗𝑗_ is the probability a randomly selected peptide 𝜌𝜌 belongs to organism *Oi*, given it also belongs to organism *Oj*. In matrix notation, the expected number of counts for all organisms <u>𝐸𝐸</u> can be written as

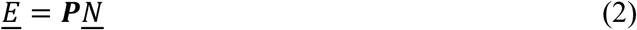

where <u>𝑁𝑁</u> is an *M* x *1* vector containing the actual number of peptides contributed by organism *i*=1,2,…*M*, and 𝑷𝑷 is an *M*x*M* matrix where entry *i,j* is 𝑃𝑃𝜌𝜌 ∈ 𝑂𝑂_𝑖𝑖_|𝜌𝜌 ∈ 𝑂𝑂_𝑗𝑗_, calculated as the number of peptides shared by organisms *i* and *j*, divided by the number of peptides belonging to organism *j*. By substituting <u>𝐸𝐸</u> with the observed number of counts <u>𝐸𝐸</u>, we estimate the actual number of peptides contributed by each organism (<u>𝑁𝑁</u>) by inverting Eqn (2),

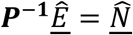

We note that this procedure, particularly the overlap matrix and the correction for the proteome similarity of closely related organisms, resembles the procedure of Penzlin et al. in the PIPASIC software.^45^

*Taxonomic group scoring*. The above scoring process adjusts the peptide counts associated with each source to reduce confusion caused by phylogenetic similarity between organisms (i.e., co-identifications). After tags are assigned to taxonomic groups, these tabulated assignments and an organism-level overlap matrix are used to assign scores via the non-negative least squares algorithm. In this step, the count of matching tags is corrected to the final taxonomic score based on the overlap coefficients. For any two organisms *i* and *j*, with *i* present in the sample and *j* absent, the expected number of hits for organism *j* is the product of the number of hits for *i* and the overlap coefficient. If fewer hits are found, the score is reduced. If more hits are found, the score is increased. MARLOWE reports both the corrected hit counts (taxonomic) scores for each detected taxonomic group, and, as a secondary score, the raw hit counts per species. To compare taxonomic scores across analyses of different data files, the taxonomic scores can be further normalized to a [0, 1] scale and/or transformed to a z-score. By applying these statistical concepts to tryptic peptide tags from the KEGG database, Release 91.0 (July 1, 2019), MARLOWE narrows down the protein sources in a sample, enabling a more focused follow-on confirmatory analysis.

### Publicly available raw mass spectrometry data for initial evaluation of MARLOWE

For initial evaluation of MARLOWE, raw spectrometry data files from the project PXD003669 were downloaded from ProteomeXchange and can be accessed via the ProteomeXchange online repository^46,47^ (http://proteomecentral.proteomexchange.org). Data from the PNNL Biodiversity Library^48^ can be accessed from ProteomeXchange via the MassIVE repository with identifiers PXD001860 and MSV000079053, although we accessed this data using an internal PNNL system. These datasets were converted to MGF format using ThermoRawFileParser for analysis with MARLOWE.

### MARLOWE implementation and execution time

MARLOWE is implemented in R (version 4.2.2) via three R packages: MakeSearchSim, CandidateSearchDatabase, and OrgIDPipeline. MakeSearchSim contains the functionality necessary to run Novor, the *de novo* peptide sequencing tool, and parse its output.

CandidateSearchDatabase contains all functionality relating to the creation and use of the MySQL database that houses a processed version of the KEGG database (Release 91.0 accessed July 2019). OrgIDPipeline contains the functions related to implementation of the statistical algorithms and methods necessary to (1) calculate the final taxonomic scores from the database query results and (2) generate the final list of potential source organisms.

An important consideration in algorithm usage is execution time. Here, we describe time estimates for use of MARLOWE itself, which is distinct from the time needed for *de novo* peptide sequencing, also described. This is because various *de novo* peptide sequencing tools exist and as such, there was no need for us to develop a bespoke tool specifically to integrate into MARLOWE for the same purpose. Further, MARLOWE provides user flexibility in its ability to utilize input from a few different *de novo* sequencing tools. For a mass spectrometry dataset, consisting of a collection of raw data files, MARLOWE requires on average, 4 minutes per data file, from ingesting *de novo* peptide sequence lists to producing a list of potential source organisms. Execution time for each individual third-party *de novo* sequencing tool will vary. In our hands, we have observed an average of 6 minutes per raw data file for Novor, and an execution time of up to 30 minutes per raw data file using the commercial tool PEAKS Online.

### Comparison with MiCId with publicly available mass spectrometry data

Data files from PXD001860 (PNNL Biodiversity Library),^49^ PXD003669,^44^ PXD014522,^50^ and PXD023033^51^ (see Supporting Information Table S1 for description of data files and references) were analyzed with both MARLOWE and MiCId.^14,15,17^ Any files from samples where the ground truth organism was not clearly and readily identified from the publication were discarded, as were files with sparse or otherwise low-quality data (see Supporting Information Table S2 for list of datafiles included for analysis). Data files were analyzed with MARLOWE as described above. The same files, in the original Thermo .raw format, were also analyzed with MiCId. Note that the precursor and fragment mass error tolerances applied were 15 ppm and 30 ppm, respectively for MiCId, and 15 ppm and 0.02 Da, respectively, for MARLOWE. The search database for MiCId was identical to the KEGG Genomes database used in MARLOWE (Supporting Information Table S3).

To compare the performance of the two software tools, data were summarized as receiver-operator characteristic (ROC) curves, which plot the cumulative positive detections against the false discovery rate across all examined datasets. For MARLOWE, we considered the organism with the greatest number of tag hits within each taxonomic group as a positive detection, and ranked these results by the z-score transformation of normalized taxonomic score (where true positive detections tend to be associated with larger scores) to build the ROC curves. For MiCId, we considered the organism with the smallest E-value according to the described algorithm as a positive result. Positive results were considered true if they matched the known, ground-truth organism at the indicated taxonomic level.

## Results & Discussion

### Proof-of-Concept: Analysis of data from the PNNL Biodiversity Library

We demonstrate the application of MARLOWE in a diverse set of 66 pure bacterial cultures and 25 simulated binary mixtures each of varying ratios (i.e., tandem mass spectra from 20, 30, or 40 proteins belonging to secondary source appended to primary source). Each simulated dataset was generated using a different random sampling of 20, 30, or 40 contributor proteins. Results are listed in Table 1 below.

**Table 1.**
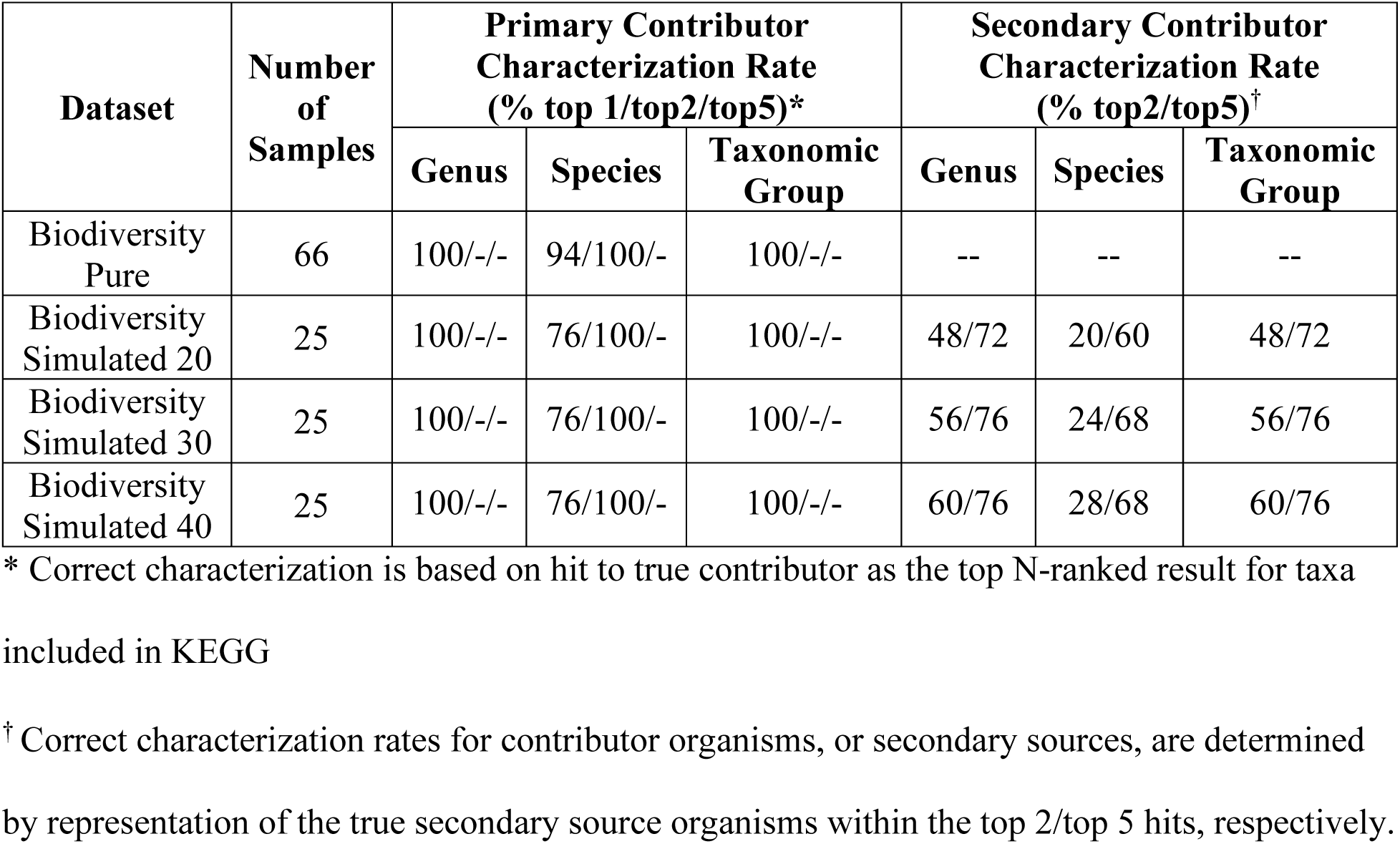
Summary of MARLOWE’s performance for single-source (pure) and simulated binary mixtures.

| Dataset | Number of Samples | Primary Contributor Characterization Rate (% top 1/top2/top5)* |  |  | Secondary Contributor Characterization Rate (% top2/top5)<sup>†</sup> |  |  |
| --- | --- | --- | --- | --- | --- | --- | --- |
|  |  | Genus | Species | Taxonomic Group | Genus | Species | Taxonomic Group |
| Biodiversity Pure | 66 | 100/-/- | 94/100/- | 100/-/- | -- | -- | -- |
| Biodiversity Simulated 20 | 25 | 100/-/- | 76/100/- | 100/-/- | 48/72 | 20/60 | 48/72 |
| Biodiversity Simulated 30 | 25 | 100/-/- | 76/100/- | 100/-/- | 56/76 | 24/68 | 56/76 |
| Biodiversity Simulated 40 | 25 | 100/-/- | 76/100/- | 100/-/- | 60/76 | 28/68 | 60/76 |

\* Correct characterization is based on hit to true contributor as the top N-ranked result for taxa included in KEGG
<sup>†</sup> Correct characterization rates for contributor organisms, or secondary sources, are determined by representation of the true secondary source organisms within the top 2/top 5 hits, respectively.

For pure cultures, we achieved 100% correct characterization specific to the species level; correct characterization is defined as MARLOWE returning the correct taxonomy in a ranked organism list (top organism, top two organisms, or top five organisms as show in Table 1). Of these correct characterizations, 100% was achieved with the correct genus and the correct taxonomic group as the top scoring organism, and 94% achieved with the correct species as the top scoring organism. 100% of the correct species were represented in the top 2 lists of highest scoring organisms.

In simulated mixtures, the primary source was 100% correctly identified at the genus and taxonomic group levels as the top-ranked hit for the various mixture ratios. At the species level, 100% correct characterization of the primary organism at the species level occurred when considering the top 2 hits. For the secondary sources, correctly identified secondary sources within the top 5 hits range between 72% and 76% at the genus and taxonomic group levels, and between 60% and 68% at the species level, across the various mixture ratios. Both the primary and contributor organisms in mixtures can be correctly characterized at a maximum rate of 76%. Characterization of mixtures is more successful when there is a larger contribution from secondary sources and when considering at taxonomic group or genus level compared to species-level. These results demonstrate initial implementation of the above statistical concepts in MARLOWE for successful organism source characterization.

### Characterization Specificity: Analysis of data from B. cereus group samples

We then examined MARLOWE’s performance—in particular, specificity—on a closely related bacterial group (Table 2). Characterization specificity was assessed using a dataset comprising seven species within the *Bacillus cereus* group that was described in Pfrunder et al. (2016).^44^ Strains include *B. cereus*, *B. cytotoxicus*, *B. mycoides*, *B. pseudomycoides*, *B. thuringiensis*, *B. anthracis* (not analyzed), and *B. toyonensis*, which all share great genomic similarity.^52^ These strains span a range of pathogen toxicity and health safety concerns,^53^ and as such, the ability to distinguish and correctly classify unknown samples to the different species in the group is of forensic, biosecurity, and health interest. Pfrunder and coworkers also examined *B. weihenstephanensis*, a strain within the *B. cereus* group that has been taxonomically reassigned to fall under *B. mycoides* species.^54^ In this analysis, the true contributor of *B. weihenstephanensis* samples at the species level is therefore assigned as *B. mycoides*. Characterization at the species level was chosen as a sufficiently specific taxonomic level since strains of the same species usually exhibit high degrees of genetic similarity. Further, actual strains of true contributors may not be included in the KEGG Genome database that forms the basis for MARLOWE, such as in the case of *B. weihenstephanensis*.

**Table 2.** Characterization summary of MARLOWE for *B. cereus* superspecies group with known true contributors.

| Dataset | Number of Samples | Characterization Rate (% top 1/top2/top5)* |  |  |  | Dataset Reference |
| --- | --- | --- | --- | --- | --- | --- |
|  |  | Family | Genus | Species | Taxonomic Group |  |
| *B. cereus* | 42 | 90/93/95 | 90/93/95 | 88/90/93 | 88/90/93 | Pfrunder et al. (2016)<sup>44</sup> |

\* Correct characterization is based on hit to true contributor as the top N-ranked result for taxa included in KEGG

To examine MARLOWE’s ability to specifically characterize the true contributor, characterization rate to the true contributor and taxonomic scores to the true contributor compared to all other taxa were quantified. We find that most samples yielded correct characterization to the true contributor *B. cereus* group member with high taxonomic scores, though characterization to *B. cereus* posed a challenge. Comparison of ranked organism lists among these samples showed high proteome diversity among the *B. cereus* group members, which are organized into different and distinct taxonomic groups. MARLOWE displays high characterization specificity for most samples containing *B. cereus* species, as lists of potential sources often unambiguously point to one *B. cereus* group member with a high taxonomic score, indicating great confidence in the characterization.

The true contributor at the species level was correctly characterized as the top-ranked organism in 88% of samples (Figure 2). Specifically, all biological and technical replicates of samples from *B. cytotoxicus*, *B. mycoides*, *B. weihenstephanensis*, *B. pseudomycoides*, and *B. thuringiensis* were correctly characterized to their respective true contributor as the organism with the highest taxonomic score and greatest number of tag-strong peptide matches. Note that samples containing *B. weihenstephanensis*, a strain reassigned as a subspecies of *B. mycoides*, were correctly characterized as *B. mycoides*. Interestingly, these species with correct characterization to the top-ranked organism had substantially high taxonomic scores (on average, 0.68 ± 0.24 (s.d.) normalized score) compared to all other ranked taxa. As discussed in the previous section, high taxonomic scores indicate greater confidence of the ranked organisms as true contributors, and the taxonomic score distributions for these samples suggest a single contributor sample rather than a mixture.

**Figure 2.**
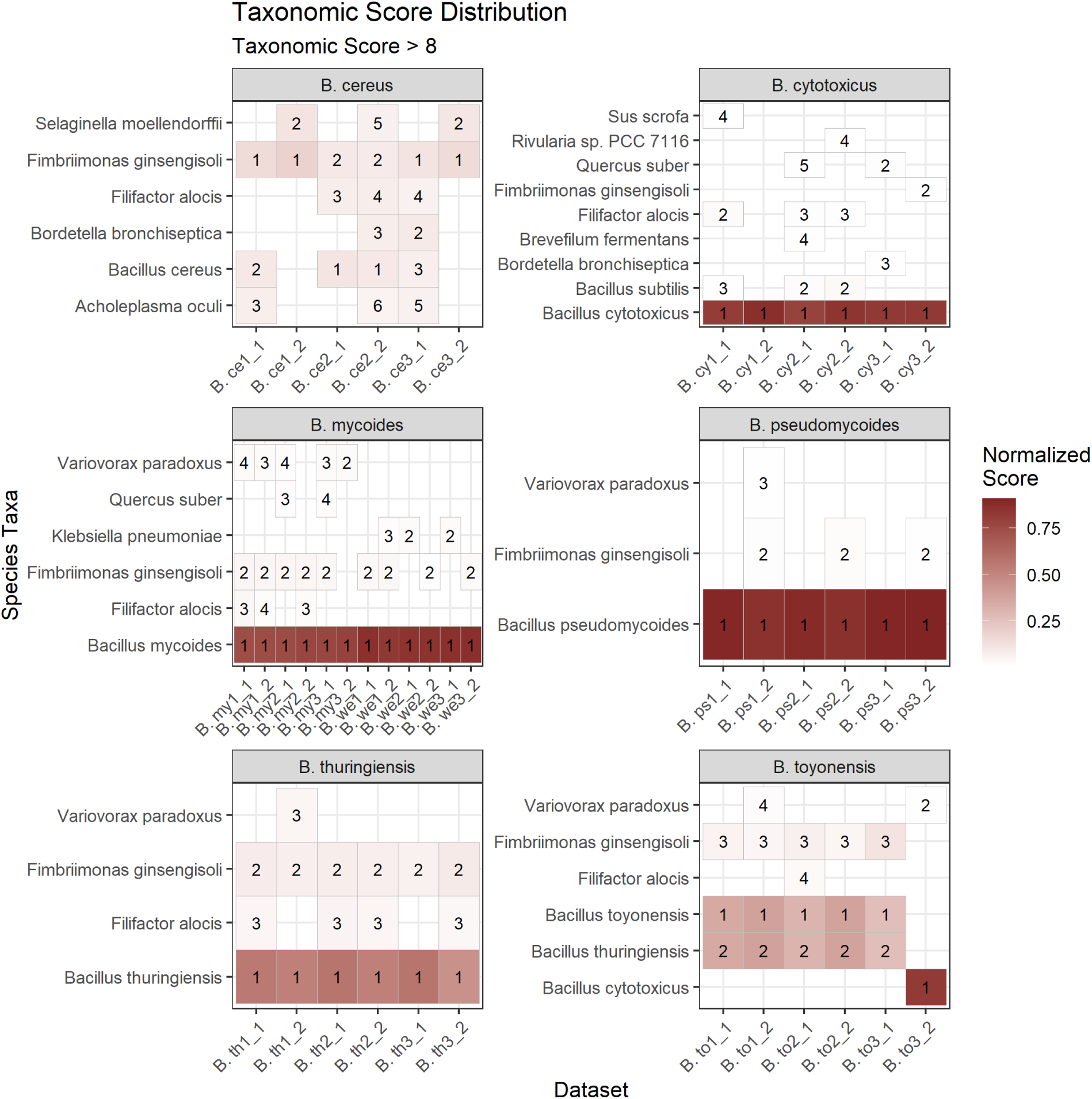
Heatmap of organism score ranking of all taxa with raw taxonomic scores > 8 for each sample within the *B. cereus* group, further grouped by the true contributor at the species level. Heatmap colors are based on normalized taxonomic score, normalized to the total taxonomic score per sample. Values in each cell represent the rank of the organism, assigned using taxonomic score and tag matches to strong peptides. MARLOWE achieved 88% correct characterization of the true contributor as the top-ranked organism in this dataset at the species level. For most samples, taxonomic scores of the true contributor are substantially higher than for all other taxa, indicating confident characterization.

For groups of samples where the true contributor was not the top-ranked organism, among the *B. toyonensis* samples, the characterization rate was 83%, as one of six replicates was characterized as *B. cytotoxicus* with a high taxonomic score (normalized score = 0.82). The high taxonomic score to *B. cytotoxicus* returned for one of the six replicates of *B. toyonensis* suggests an error in sample preparation rather than mischaracterization by MARLOWE; the distribution of taxonomic scores for this sample is also vastly different from the other *B. toyonensis* samples. A substantially greater number of peptide-spectrum matches to *B. cytotoxicus*-specific proteins (3935) compared to *B. toyonensis*-specific proteins (30) from conventional database searching confirms the sample preparation error for that replicate.

Characterization of *B. cereus* samples posed a challenge for MARLOWE, as only two of six replicates ranked the true contributor as the top organism, with *B. cereus* correctly characterized within the top three organisms in four of the six replicates. The results for *B. cereus* samples are also distinct in that the taxonomic scores for this set of samples are generally low and with a minimum taxonomic score of 6.3 (on average, 0.09 ± 0.02 (s.d.) normalized score and 11.4 ± 4.1 (s.d.) taxonomic score). The low taxonomic scores for *B. cereus* samples are due to the structure of the database and on taxonomic classifications. There are many strains of *B. cereus* in MARLOWE’s database (i.e., 14 strains of *B. cereus* split among 5 taxonomic groups). The presence of so many similar peptides in different species results in fewer strong peptides assigned to each specific strain of *B. cereus*, and thus similarly lower taxonomic scores overall. This illustrates the subtle dependence of MARLOWE results on database contents and the similarity of organisms across taxa. Possibly, this score could be improved by using groups constructed by phylogenetic clustering rather than taxonomy assignments. Despite not being the top-ranked organism, *B. cereus* is still returned as a potential organism by MARLOWE within the top three organisms in at least four of six replicates across *B. cereus* samples.

MARLOWE’s characterization specificity to members of the *B. cereus* group was examined by comparing taxonomic scores of the true contributor to all other taxa, as well as noting the different species that were ranked. Most notably, except for *B. toyonensis* samples, all ranked taxa other than the true contributor are not members of the *B. cereus* group, and in fact most do not belong to the *Bacillus* genus. Hits to other members of the *B. cereus* group yielded extremely low taxonomic scores (< 1.8 taxonomic score, n = 43 hits across 27 samples), which were filtered out, as the low magnitude of these scores represents spurious tag-peptide matches to peptides not distinctly strong to a single taxonomic group and indicates low confidence of the organism as a contributor in the sample. This suggests that proteomes of each species, particularly the strong peptide distributions, within the *B. cereus* group are sufficiently diverse from other *B. cereus* members and are also distinct from the proteomes of organisms in the *Bacillus* genus. As for the *B. toyonensis* samples, both *B. toyonensis* and *B. thuringiensis* were returned from MARLOWE for the samples that were correctly characterized. Further analysis of their taxonomic scores revealed that the two species belong to the same taxonomic group, and as such, have identical taxonomic scores. *B. toyonensis* was ranked higher than *B. thuringiensis* owing to a greater number of tag-strong peptide matches (on average, 75 ± 45 (s.d.) matches for *B. toyonensis* compared to 64 ± 39 (s.d.) matches for *B. thuringiensis*). In fact, *B. toyonensis* and *B. thuringiensis* MC28 reside in the same taxonomic group, but a different strain of *B. thuringiensis*, *B. thuringiensis* serovar chinensis CT-43, belongs to a different taxonomic group. The latter strain is the top-ranked organism for *B. thuringiensis* samples. Comparison of their genome sequences via NCBI BLAST showed 99% genetic similarity between *B. toyonensis* and *B. thuringiensis* MC28, but 97% similarity between the two strains of *B. thuringiensis*. This observation indicates not only the ability for MARLOWE to correctly characterize contributors to the species level, but also that similarity of organism proteomes may diverge from their assigned taxonomy and MARLOWE captures these differences.

### Comparison of MARLOWE and MiCId results

Comparable performance between MARLOWE and MiCId was observed, though MiCId demonstrated slightly higher accuracy and specificity (Figure 3). Using the same taxonomic database, analysis of 225 MS/MS data files from four different datasets (Biodiversity, PXD003669, PXD014522, and PXD023033) showed MARLOWE’s ability to identify the correct source organism 63% of the time within the top-ranked taxonomic group, and the correct organism was present in any taxonomic group 74% of the time. We note that the vast majority of false negatives derived from the low concentrations (i.e., ≤ 1 × 10^4^ cells/mL) of organisms in datasets from PXD023033, with similar results using MiCId. Through ROC curve analysis for the Biodiversity and PXD003669 datasets, we demonstrate 91.4% specificity for MARLOWE’s characterization at the species level, at a false discovery rate of 5% (Figure 3A). MiCId’s performance was slightly higher at 96.7% specificity at the species level. MARLOWE’s performance remained the same at the genus level, and MiCId’s performance increased to 97.8%, also at 5% FDR (Figure 3B).

**Figure 3.**
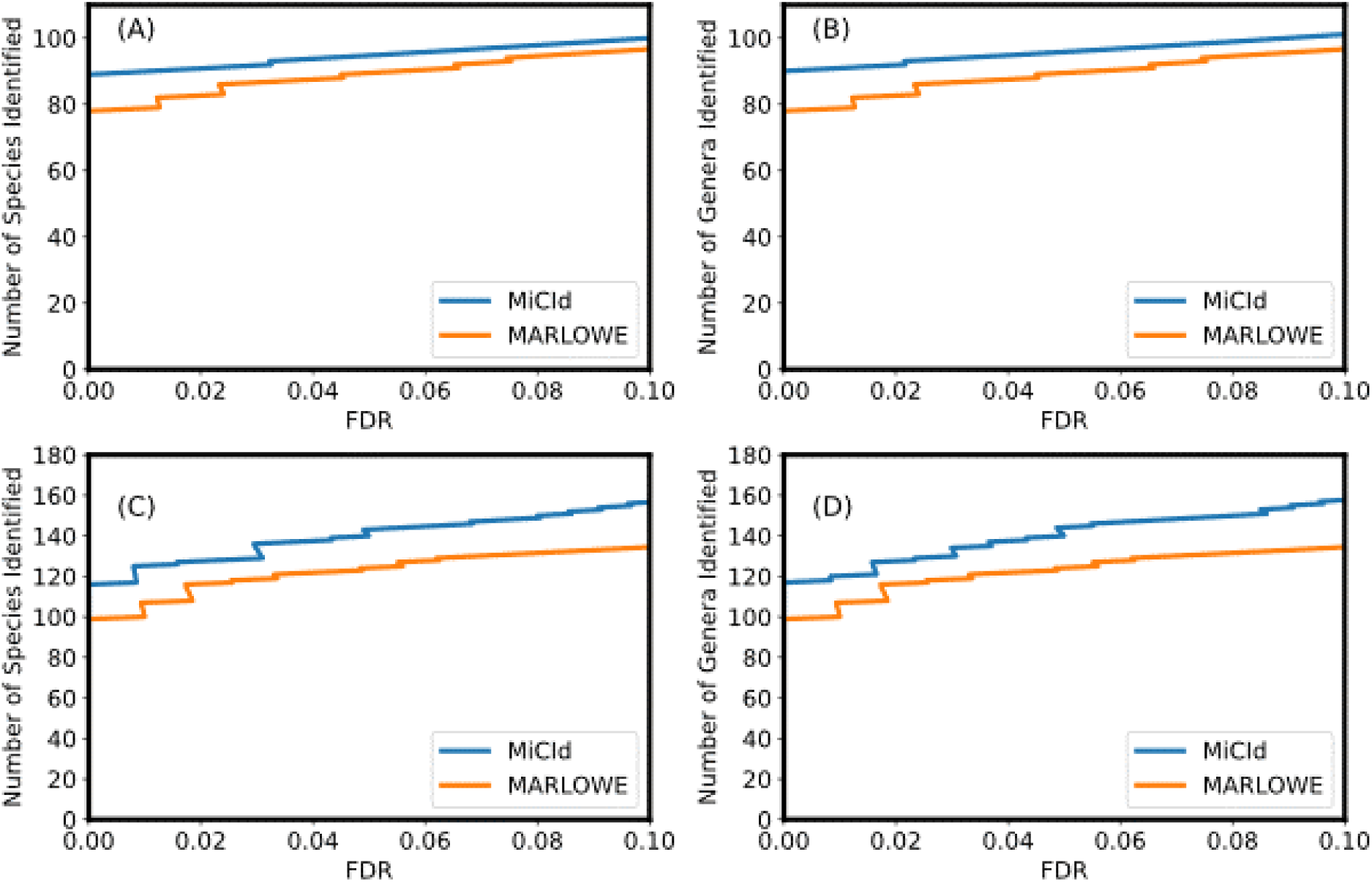
Comparison of MARLOWE and MiCId’s taxonomic characterization performance, demonstrated by receiver operator characteristic curve analysis at the (A) species and (B) genus levels for the Biodiversity and PXD003669 datasets, and at the (C) species and (D) genus levels for the Biodiversity, PXD003669, PXD014522, and PXD023033 datasets.

Comparison results suggest that MARLOWE’s approach to taxonomic characterization comes very close to MiCId’s performance, despite its intended use in generating investigate leads of potential source organisms. Given that MARLOWE’s approach relies on *de novo* sequenced peptides and knowing that *de novo* sequencing is less accurate than database searching for peptide identification, it is not surprising that there is a still a gap between the accuracies and specificities of the two tools. However, MARLOWE shows real promise and highlights the advantage of using *de novo* peptides for untargeted source characterization of unknown biological datasets.

### Conclusion: MARLOWE use and follow-up actions

The data presented show that MARLOWE performs almost as well as MiCId when using an identical organism database. However, MARLOWE does have certain limitations. In this paper, we examined datasets from samples that contain only one or two organisms. In a recent publication (Chu et al. 2025, *in press*), we have shown that MARLOWE performs well on samples with up to four organisms. Future studies should investigate the application of MARLOWE to more complex samples. Since MARLOWE relies on *de novo* peptide identification, data with high resolution for both precursor and fragment ion spectra is ideal.

However, Muth and coworkers^39^ have shown that *de novo* analysis of both high-resolution and low-resolution MS/MS spectra can produce accurate short sequence tags, so it is possible that MARLOWE could function with low-resolution MS/MS, but we have not tested this application. In this study, we used Novor^37^, but in principle MARLOWE can use *de novo* PSMs from any tool that supplies overall PSM scores *and* local (per residue) confidence scores, including Novor, PEAKS, and the newer deep-learning tool Casanovo.^38,55^ (Local scores are needed for defining tags in the MARLOWE workflow.) Further, newer versions of PEAKS can apply *de novo* analysis to data-independent data acquisition data, and export those results in the same format as for data-dependent data. Therefore, MARLOWE should in principle be able to conduct taxonomic characterization using DIA data as input, and we are investigating that avenue as well. MARLOWE also accommodates any post-translational modification that the underlying *de novo* tool accommodates.

Although MARLOWE uses a “database-free” *de novo* approach for microorganism identification/taxonomic characterization, the size, contents, and quality of the database used by MARLOWE does influence the outcome of organism identifications. The impact of database parameters deserves detailed investigation, but is outside the scope of the current study.

The intent of MARLOWE is to provide investigative leads on potential sources given an unknown bioforensic sample, without presumption of potential sources either during reassembly of peptide sequences from mass spectrometry data or during assignment of peptide tags to organisms and using a statistically robust method to do so. For this forensics-focused application, MARLOWE’s performance at the family, genus, and species levels is suitable; subspecies- or strain-level identification is generally not necessary. Results from the proof-of-concept analysis using the Biodiversity dataset as well as the *B. cereus* group dataset demonstrate MARLOWE’s ability to correctly characterize major contributors in single species and binary species, with a high degree of specificity. MARLOWE’s strengths are in narrowing down to potential source organisms, and as such, would be most suitable at the beginning of a bioinformatics pipeline or investigation. We expect and encourage follow-up actions on the results provided by MARLOWE to include confirmatory analyses, such as via database searching or applying bespoke data analysis pipelines for other qualitative and quantitative analyses. We believe that MARLOWE can be broadly applicable for several applications in mass spectrometry-based protein analysis, and its modular nature is amenable to being co-opted into other pipelines for bespoke bioinformatics analyses.

## Acknowledgments

The authors acknowledge the Environmental and Molecular Sciences Laboratory for liquid chromatography-tandem mass spectrometry data acquisition of samples. Development of MARLOWE’s workflow was supported by Department of Homeland Security, Science and Technology Directorate - Homeland Security Advanced Research Projects Agency - Chemical and Biological Division under Contract #HSHQPM16X00216. This research was supported in part by the Intramural Research Program of the National Library of Medicine at the NIH.

## Data Availability

Publicly available raw mass spectrometry data were acquired from ProteomeXchange and can be accessed via the online repository^46,47^ (http://proteomecentral.proteomexchange.org) under accessions PXD001860, PXD003669, PXD014522, and PXD023033.

## Author contributions

SCJ: algorithm design and development, data analysis

FC: data analysis, manuscript writing

GA: data analysis, manuscript writing

AYO: data analysis, manuscript writing

ASB: algorithm development and testing

DLC: algorithm design

NCH: algorithm development

EDM: algorithm conception and design, manuscript writing

YKY: manuscript review and editing

KHJ: algorithm conception, design, and development

