## Supplementary Information for "MARLOWE: Taxonomic Characterization of Unknown Samples for Forensics Using *De Novo* Peptide Identification"

**‡Current affiliation:**
Kristin H. Jarman
Karius Inc.
975 Island Dr., Suite 101
Redwood City, CA, 94065

**Table S1. ProteomeXchange Datasets Used in this Study**

| ProteomeXchange Identifier | Reference | Organisms |
|---|---|---|
| PXD001860 | Payne, S. H. et al. The Pacific Northwest National Laboratory library of bacterial and archaeal proteomic biodiversity. *Scientific Data* 2, 150041, doi:10.1038/sdata.2015.41 (2015). | 109 bacterial organisms |
| PXD003669 | Pfrunder, S. et al. Bacillus cereus Group-Type Strain-Specific Diagnostic Peptides. *J. Proteome Res.* 15, 3098-3107, doi:10.1021/acs.jproteome.6b00216 (2016). | *Bacillus cereus* group members |
| PXD014522 | Karlsson, R. et al. Discovery of Species-unique Peptide Biomarkers of Bacterial Pathogens by Tandem Mass Spectrometry-based Proteotyping. *Mol. Cell. Proteomics* 19, 518-528, doi:https://doi.org/10.1074/mcp.RA119.001667 (2020). | *Haemophilus influenzae, Moraxella catarrhalis, Staphylococcus aureus, Streptococcus pneumoniae* (strain ATCC BAA-255 / R6) |
| PXD023033 | Kondori, N. et al. Mass Spectrometry Proteotyping-Based Detection and Identification of Staphylococcus aureus, Escherichia coli, and Candida albicans in Blood. *Frontiers in Cellular and Infection Microbiology* 11, doi:10.3389/fcimb.2021.634215 (2021). | *Candida albicans, Escherichia coli, Staphylococcus aureus* |

Table S2. Datafiles from ProteomeXchange Datasets Used in this Study

| ProteomeXchange Identifier | Filename | Species Name | Strain Names | Cysteine Carbamylated? | Used in ROC Analysis? |
|---|---|---|---|---|---|
| PXD014522 | Hi_QEHF_171201_21.raw | Haemophilus influenzae | NA | No | Yes |
| PXD014522 | Hi_QE_170112_37.raw | Haemophilus influenzae | NA | No | Yes |
| PXD014522 | Hi_QE_170113_37.raw | Haemophilus influenzae | NA | No | Yes |
| PXD014522 | Hi_QE_170113_38.raw | Haemophilus influenzae | NA | No | Yes |
| PXD014522 | Hi_QE_170524_80.raw | Haemophilus influenzae | NA | No | Yes |
| PXD014522 | Hi_QE_171019_07.raw | Haemophilus influenzae | NA | No | Yes |
| PXD014522 | Hi_QE_171019_34.raw | Haemophilus influenzae | NA | No | Yes |
| PXD014522 | Hi_QE_171019_57.raw | Haemophilus influenzae | NA | No | Yes |
| PXD014522 | Hi_QE_171110_44.raw | Haemophilus influenzae | NA | No | Yes |
| PXD014522 | Hi_QE_171110_65.raw | Haemophilus influenzae | NA | No | Yes |
| PXD014522 | Hi_QE_171110_71.raw | Haemophilus influenzae | NA | No | Yes |
| PXD014522 | Hi_QE_171110_77.raw | Haemophilus influenzae | NA | No | Yes |
| PXD014522 | Hi_QE_180126_33.raw | Haemophilus influenzae | NA | No | Yes |
| PXD014522 | Mc_QEHF_171201_18.raw | Moraxella catarrhalis | NA | No | Yes |
| PXD014522 | Mc_QEHF_171201_24.raw | Moraxella catarrhalis | NA | No | Yes |
| PXD014522 | Mc_QE_170110_14.raw | Moraxella catarrhalis | NA | No | No |
| PXD014522 | Mc_QE_170110_20.raw | Moraxella catarrhalis | NA | No | No |
| PXD014522 | Mc_QE_171019_11.raw | Moraxella catarrhalis | NA | No | Yes |
| PXD014522 | Mc_QE_171019_61.raw | Moraxella catarrhalis | NA | No | Yes |
| PXD014522 | Mc_QE_171019_85.raw | Moraxella catarrhalis | NA | No | Yes |
| PXD014522 | Mc_QE_171019_86.raw | Moraxella catarrhalis | NA | No | Yes |
| PXD014522 | Sa_QEHF_171201_69.raw | Staphylococcus aureus | NA | No | Yes |
| PXD014522 | Sa_QEHF_171201_81.raw | Staphylococcus aureus | NA | No | No |
| PXD014522 | Sa_QEHF_171201_84.raw | Staphylococcus aureus | NA | No | No |
| PXD014522 | Sa_QEHF_181205_23.raw | Staphylococcus aureus | NA | No | Yes |
| PXD014522 | Sa_QEHF_181205_27.raw | Staphylococcus aureus | NA | No | Yes |
| PXD014522 | Sa_QEHF_181205_29.raw | Staphylococcus aureus | NA | No | Yes |

| ProteomeXchange Identifier | Filename | Species Name | Strain Names | Cysteine Carbamylated? | Used in ROC Analysis? |
|---|---|---|---|---|---|
| PXD014522 | Sa_QEHF_190218_71.raw | Staphylococcus aureus | NA | No | Yes |
| PXD014522 | Sa_QE_170412_11.raw | Staphylococcus aureus | NA | No | Yes |
| PXD014522 | Sa_QE_180126_21.raw | Staphylococcus aureus | NA | No | Yes |
| PXD014522 | Sp_QEHF_171201_27.raw | Streptococcus pneumoniae | NA | No | Yes |
| PXD014522 | Sp_QEHF_181205_49.raw | Streptococcus pneumoniae | NA | No | Yes |
| PXD014522 | Sp_QEHF_190218_83.raw | Streptococcus pneumoniae | NA | No | Yes |
| PXD014522 | Sp_QE_170110_08.raw | Streptococcus pneumoniae | NA | No | Yes |
| PXD014522 | Sp_QE_170110_20.raw | Streptococcus pneumoniae | NA | No | No |
| PXD014522 | Sp_QE_170112_12.raw | Streptococcus pneumoniae | NA | No | Yes |
| PXD014522 | Sp_QE_170112_13.raw | Streptococcus pneumoniae | NA | No | Yes |
| PXD014522 | Sp_QE_170112_21.raw | Streptococcus pneumoniae | NA | No | Yes |
| PXD014522 | Sp_QE_170112_28.raw | Streptococcus pneumoniae | NA | No | No |
| PXD014522 | Sp_QE_171019_23.raw | Streptococcus pneumoniae | NA | No | Yes |
| PXD014522 | Sp_QE_171019_26.raw | Streptococcus pneumoniae | NA | No | Yes |
| PXD014522 | Sp_QE_171019_69.raw | Streptococcus pneumoniae | NA | No | Yes |
| PXD014522 | Sp_QE_171019_70.raw | Streptococcus pneumoniae | NA | No | Yes |
| PXD014522 | Sp_QE_171110_38.raw | Streptococcus pneumoniae | NA | No | Yes |
| PXD023033 | QEHF_190520_37.raw | Staphylococcus aureus | S. aureus (CCUG 41582) | No | Yes |
| PXD023033 | QEHF_190520_38.raw | Staphylococcus aureus | S. aureus (CCUG 41582) | No | Yes |
| PXD023033 | QEHF_190614_42.raw | Staphylococcus aureus | S. aureus (CCUG 41582) | No | Yes |
| PXD023033 | QEHF_190614_43.raw | Staphylococcus aureus | S. aureus (CCUG 41582) | No | Yes |
| PXD023033 | QEHF_190614_47.raw | Staphylococcus aureus | S. aureus (CCUG 41582) | No | Yes |
| PXD023033 | QEHF_190614_48.raw | Staphylococcus aureus | S. aureus (CCUG 41582) | No | Yes |
| PXD023033 | QEHF_190614_49.raw | Staphylococcus aureus | S. aureus (CCUG 41582) | No | Yes |
| PXD023033 | QEHF_190520_41.raw | Staphylococcus aureus | S. aureus (CCUG 41582) | No | Yes |
| PXD023033 | QEHF_190614_54.raw | Staphylococcus aureus | S. aureus (CCUG 41582) | No | Yes |
| PXD023033 | QEHF_190614_55.raw | Staphylococcus aureus | S. aureus (CCUG 41582) | No | Yes |
| PXD023033 | QEHF_190523_08.raw | Escherichia coli | E. coli (CCUG 49263) | No | Yes |

| ProteomeXchange Identifier | Filename | Species Name | Strain Names | Cysteine Carbamylated? | Used in ROC Analysis? |
|---|---|---|---|---|---|
| PXD023033 | QEHF_190614_62.raw | Escherichia coli | E. coli (CCUG 49263) | No | Yes |
| PXD023033 | QEHF_190614_63.raw | Escherichia coli | E. coli (CCUG 49263) | No | Yes |
| PXD023033 | QEHF_190614_64.raw | Escherichia coli | E. coli (CCUG 49263) | No | Yes |
| PXD023033 | QEHF_190614_68.raw | Escherichia coli | E. coli (CCUG 49263) | No | Yes |
| PXD023033 | QEHF_190614_69.raw | Escherichia coli | E. coli (CCUG 49263) | No | Yes |
| PXD023033 | QEHF_190614_70.raw | Escherichia coli | E. coli (CCUG 49263) | No | Yes |
| PXD023033 | QEHF_190614_74.raw | Escherichia coli | E. coli (CCUG 49263) | No | Yes |
| PXD023033 | QEHF_190614_75.raw | Escherichia coli | E. coli (CCUG 49263) | No | Yes |
| PXD023033 | QEHF_190614_76.raw | Escherichia coli | E. coli (CCUG 49263) | No | Yes |
| PXD023033 | QEHF_190703_25.raw | Candida albicans | C. albicans (CCUG 32723) | No | Yes |
| PXD023033 | QEHF_190703_32.raw | Candida albicans | C. albicans (CCUG 32723) | No | Yes |
| PXD023033 | QEHF_190703_34.raw | Candida albicans | C. albicans (CCUG 32723) | No | Yes |
| PXD023033 | QEHF_190624_14.raw | Candida albicans | C. albicans (CCUG 32723) | No | Yes |
| PXD023033 | QEHF_190703_09.raw | Candida albicans | C. albicans (CCUG 32723) | No | Yes |
| PXD023033 | QEHF_190703_17.raw | Candida albicans | C. albicans (CCUG 32723) | No | Yes |
| PXD023033 | QEHF_190703_26.raw | Candida albicans | C. albicans (CCUG 32723) | No | Yes |
| PXD023033 | QEHF_190703_10.raw | Candida albicans | C. albicans (CCUG 32723) | No | Yes |
| PXD023033 | QEHF_190703_18.raw | Candida albicans | C. albicans (CCUG 32723) | No | Yes |
| PXD023033 | QEHF_190703_27.raw | Candida albicans | C. albicans (CCUG 32723) | No | Yes |
| PXD023033 | QEHF_190703_11.raw | Candida albicans | C. albicans (CCUG 32723) | No | Yes |
| PXD023033 | QEHF_190703_19.raw | Candida albicans | C. albicans (CCUG 32723) | No | Yes |
| PXD023033 | QEHF_190703_28.raw | Candida albicans | C. albicans (CCUG 32723) | No | Yes |
| PXD023033 | QEHF_190703_12.raw | Candida albicans | C. albicans (CCUG 32723) | No | Yes |
| PXD023033 | QEHF_190703_20.raw | Candida albicans | C. albicans (CCUG 32723) | No | Yes |
| PXD023033 | QEHF_190703_29.raw | Candida albicans | C. albicans (CCUG 32723) | No | Yes |
| PXD023033 | QEHF_190816_03.raw | Escherichia coli | E. coli (CCUG 49263) | No | Yes |
| PXD023033 | QEHF_190816_04.raw | Escherichia coli | E. coli (CCUG 49263) | No | Yes |
| PXD023033 | QEHF_190816_05.raw | Escherichia coli | E. coli (CCUG 49263) | No | Yes |

| ProteomeXchange Identifier | Filename | Species Name | Strain Names | Cysteine Carbamylated? | Used in ROC Analysis? |
|---|---|---|---|---|---|
| PXD023033 | QEHF_190816_06.raw | Escherichia coli | E. coli (CCUG 49263) | No | Yes |
| PXD023033 | QEHF_190816_07.raw | Escherichia coli | E. coli (CCUG 49263) | No | Yes |
| PXD023033 | QEHF_190816_08.raw | Escherichia coli | E. coli (CCUG 49263) | No | Yes |
| PXD023033 | QEHF_190816_10.raw | Escherichia coli | E. coli (CCUG 49263) | No | Yes |
| PXD023033 | QEHF_181106_06.raw | Escherichia coli | E. coli (CCUG 49263) | No | Yes |
| PXD023033 | QEHF_181106_20.raw | Escherichia coli | E. coli (CCUG 49263) | No | Yes |
| PXD023033 | QEHF_181106_07.raw | Escherichia coli | E. coli (CCUG 49263) | No | Yes |
| PXD023033 | QEHF_181106_28.raw | Escherichia coli | E. coli (CCUG 49263) | No | Yes |
| PXD023033 | QEHF_181106_08.raw | Escherichia coli | E. coli (CCUG 49263) | No | Yes |
| PXD023033 | QEHF_181106_21.raw | Escherichia coli | E. coli (CCUG 49263) | No | Yes |
| PXD023033 | QEHF_181106_09.raw | Escherichia coli | E. coli (CCUG 49263) | No | Yes |
| PXD023033 | QEHF_181106_22.raw | Escherichia coli | E. coli (CCUG 49263) | No | Yes |
| PXD023033 | QEHF_181106_10.raw | Escherichia coli | E. coli (CCUG 49263) | No | Yes |
| PXD023033 | QEHF_181106_23.raw | Escherichia coli | E. coli (CCUG 49263) | No | Yes |
| PXD023033 | QEHF_181106_11.raw | Escherichia coli | E. coli (CCUG 49263) | No | Yes |
| PXD023033 | QEHF_181106_24.raw | Escherichia coli | E. coli (CCUG 49263) | No | Yes |
| PXD023033 | QEHF_181106_12.raw | Escherichia coli | E. coli (CCUG 49263) | No | Yes |
| PXD023033 | QEHF_181106_25.raw | Escherichia coli | E. coli (CCUG 49263) | No | Yes |
| PXD023033 | QEHF_181106_13.raw | Escherichia coli | E. coli (CCUG 49263) | No | Yes |
| PXD023033 | QEHF_181106_26.raw | Escherichia coli | E. coli (CCUG 49263) | No | Yes |
| PXD023033 | Lumos_201104_13.raw | Staphylococcus aureus | S. aureus (CCUG 41582) | No | Yes |
| PXD023033 | Lumos_201104_14.raw | Staphylococcus aureus | S. aureus (CCUG 41582) | No | Yes |
| PXD023033 | Lumos_201104_15.raw | Staphylococcus aureus | S. aureus (CCUG 41582) | No | Yes |
| PXD023033 | Lumos_201104_16.raw | Staphylococcus aureus | S. aureus (CCUG 41582) | No | Yes |
| PXD023033 | Lumos_201104_17.raw | Staphylococcus aureus | S. aureus (CCUG 41582) | No | Yes |
| PXD023033 | Lumos_201104_18.raw | Staphylococcus aureus | S. aureus (CCUG 41582) | No | Yes |
| PXD023033 | Lumos_201104_19.raw | Staphylococcus aureus | S. aureus (CCUG 41582) | No | Yes |
| PXD023033 | Lumos_201104_20.raw | Staphylococcus aureus | S. aureus (CCUG 41582) | No | Yes |

| **ProteomeXchange Identifier** | **Filename** | **Species Name** | **Strain Names** | **Cysteine Carbamylated?** | **Used in ROC Analysis?** |
|---|---|---|---|---|---|
| PXD023033 | Lumos_201104_21.raw | Staphylococcus aureus | S. aureus (CCUG 41582) | No | Yes |
| PXD023033 | Lumos_201104_22.raw | Staphylococcus aureus | S. aureus (CCUG 41582) | No | Yes |
| PXD023033 | Lumos_201104_23.raw | Staphylococcus aureus | S. aureus (CCUG 41582) | No | Yes |
| PXD023033 | Lumos_201104_24.raw | Staphylococcus aureus | S. aureus (CCUG 41582) | No | Yes |
| PXD023033 | Lumos_201104_25.raw | Staphylococcus aureus | S. aureus (CCUG 41582) | No | Yes |
| PXD023033 | Lumos_201104_26.raw | Staphylococcus aureus | S. aureus (CCUG 41582) | No | Yes |
| PXD023033 | Lumos_201104_33.raw | Staphylococcus aureus | S. aureus (CCUG 41582) | No | Yes |
| PXD023033 | Lumos_201104_34.raw | Staphylococcus aureus | S. aureus (CCUG 41582) | No | Yes |
| PXD023033 | Lumos_201104_35.raw | Staphylococcus aureus | S. aureus (CCUG 41582) | No | Yes |
| PXD023033 | Lumos_201104_36.raw | Staphylococcus aureus | S. aureus (CCUG 41582) | No | Yes |
| PXD023033 | Lumos_201104_37.raw | Staphylococcus aureus | S. aureus (CCUG 41582) | No | Yes |
| PXD023033 | Lumos_201104_38.raw | Staphylococcus aureus | S. aureus (CCUG 41582) | No | Yes |
| PXD023033 | Lumos_201104_39.raw | Staphylococcus aureus | S. aureus (CCUG 41582) | No | Yes |
| PXD023033 | Lumos_201104_40.raw | Staphylococcus aureus | S. aureus (CCUG 41582) | No | Yes |
| PXD023033 | Lumos_201104_41.raw | Staphylococcus aureus | S. aureus (CCUG 41582) | No | Yes |
| PXD023033 | Lumos_201104_42.raw | Staphylococcus aureus | S. aureus (CCUG 41582) | No | Yes |
| PXD023033 | Lumos_201104_43.raw | Staphylococcus aureus | S. aureus (CCUG 41582) | No | Yes |
| PXD023033 | Lumos_201104_44.raw | Staphylococcus aureus | S. aureus (CCUG 41582) | No | Yes |
| PXD023033 | Lumos_201104_45.raw | Staphylococcus aureus | S. aureus (CCUG 41582) | No | Yes |
| PXD023033 | Lumos_201104_46.raw | Staphylococcus aureus | S. aureus (CCUG 41582) | No | Yes |
| PXD023033 | QEHF_181122_19.raw | Candida albicans | C. albicans (CCUG 32723) | No | Yes |
| PXD023033 | QEHF_181122_20.raw | Candida albicans | C. albicans (CCUG 32723) | No | Yes |
| PXD023033 | QEHF_181122_21.raw | Candida albicans | C. albicans (CCUG 32723) | No | Yes |
| PXD023033 | QEHF_181122_22.raw | Candida albicans | C. albicans (CCUG 32723) | No | Yes |
| PXD023033 | QEHF_181122_23.raw | Candida albicans | C. albicans (CCUG 32723) | No | Yes |
| PXD023033 | QEHF_181122_24.raw | Candida albicans | C. albicans (CCUG 32723) | No | No |
| PXD023033 | QEHF_181122_25.raw | Candida albicans | C. albicans (CCUG 32723) | No | Yes |
| PXD023033 | QEHF_181122_26.raw | Candida albicans | C. albicans (CCUG 32723) | No | Yes |

| ProteomeXchange Identifier | Filename | Species Name | Strain Names | Cysteine Carbamylated? | Used in ROC Analysis? |
|---|---|---|---|---|---|
| PXD003669 | 20150219sp_6843_B12_a1_1.raw | Bacillus mycoides | Bacillus cereus (DSM 31) | Yes | Yes |
| PXD003669 | 20150219sp_6843_B12_a1_2.raw | Bacillus mycoides | Bacillus cereus (DSM 31) | Yes | Yes |
| PXD003669 | 20150219sp_6843_B12_a2_1.raw | Bacillus mycoides | Bacillus cereus (DSM 31) | Yes | Yes |
| PXD003669 | 20150219sp_6843_B12_a2_2.raw | Bacillus mycoides | Bacillus cereus (DSM 31) | Yes | Yes |
| PXD003669 | 20150219sp_6843_B12_a3_1.raw | Bacillus mycoides | Bacillus cereus (DSM 31) | Yes | Yes |
| PXD003669 | 20150219sp_6843_B12_a3_2.raw | Bacillus mycoides | Bacillus cereus (DSM 31) | Yes | Yes |
| PXD003669 | 20150224sp_6836_B18_a1_1.raw | Bacillus cereus | Bacillus cereus (DSM 31) | Yes | Yes |
| PXD003669 | 20150224sp_6836_B18_a1_2.raw | Bacillus cereus | Bacillus cereus (DSM 31) | Yes | Yes |
| PXD003669 | 20150224sp_6836_B18_a2_1.raw | Bacillus cereus | Bacillus cereus (DSM 31) | Yes | Yes |
| PXD003669 | 20150224sp_6836_B18_a2_2.raw | Bacillus cereus | Bacillus cereus (DSM 31) | Yes | Yes |
| PXD003669 | 20150224sp_6836_B18_a3_1.raw | Bacillus cereus | Bacillus cereus (DSM 31) | Yes | Yes |
| PXD003669 | 20150224sp_6836_B18_a3_2.raw | Bacillus cereus | Bacillus cereus (DSM 31) | Yes | Yes |
| PXD003669 | 20150224sp_6841_B17_a1_1.raw | Bacillus thuringiensis | Bacillus toyonensis (CECT 876) | Yes | Yes |
| PXD003669 | 20150224sp_6841_B17_a1_2.raw | Bacillus thuringiensis | Bacillus toyonensis (CECT 876) | Yes | Yes |
| PXD003669 | 20150224sp_6841_B17_a2_1.raw | Bacillus thuringiensis | Bacillus toyonensis (CECT 876) | Yes | Yes |
| PXD003669 | 20150224sp_6841_B17_a2_2.raw | Bacillus thuringiensis | Bacillus toyonensis (CECT 876) | Yes | Yes |
| PXD003669 | 20150224sp_6841_B17_a3_1.raw | Bacillus thuringiensis | Bacillus toyonensis (CECT 876) | Yes | Yes |
| PXD003669 | 20150224sp_6841_B17_a3_2.raw | Bacillus thuringiensis | Bacillus toyonensis (CECT 876) | Yes | Yes |
| PXD003669 | 20150306sp_7007_B9_a1_1.raw | Bacillus mycoides;Bacillus weihenstephanensis | B. pseudomycoides (DSM 12442) | Yes | Yes |
| PXD003669 | 20150306sp_7007_B9_a1_2.raw | Bacillus mycoides;Bacillus weihenstephanensis | B. pseudomycoides (DSM 12442) | Yes | Yes |
| PXD003669 | 20150306sp_7007_B9_a2_1.raw | Bacillus mycoides;Bacillus weihenstephanensis | B. pseudomycoides (DSM 12442) | Yes | Yes |
| PXD003669 | 20150306sp_7007_B9_a2_2.raw | Bacillus mycoides;Bacillus weihenstephanensis | B. pseudomycoides (DSM 12442) | Yes | Yes |

| ProteomeXchange Identifier | Filename | Species Name | Strain Names | Cysteine Carbamylated? | Used in ROC Analysis? |
|---|---|---|---|---|---|
| PXD003669 | 20150306sp_7007_B9_a3_1.raw | Bacillus mycoides;Bacillus weihenstephanensis | B. pseudomycoides (DSM 12442) | Yes | Yes |
| PXD003669 | 20150306sp_7007_B9_a3_2.raw | Bacillus mycoides;Bacillus weihenstephanensis | B. pseudomycoides (DSM 12442) | Yes | Yes |
| PXD003669 | 20150309sp_7008_B13_a1_1.raw | Bacillus pseudomycoides | Bacillus toyonensis (CECT 876) | Yes | Yes |
| PXD003669 | 20150309sp_7008_B13_a1_2.raw | Bacillus pseudomycoides | Bacillus toyonensis (CECT 876) | Yes | Yes |
| PXD003669 | 20150309sp_7008_B13_a2_1.raw | Bacillus pseudomycoides | Bacillus toyonensis (CECT 876) | Yes | Yes |
| PXD003669 | 20150309sp_7008_B13_a2_2.raw | Bacillus pseudomycoides | Bacillus toyonensis (CECT 876) | Yes | Yes |
| PXD003669 | 20150309sp_7008_B13_a3_1.raw | Bacillus pseudomycoides | Bacillus toyonensis (CECT 876) | Yes | Yes |
| PXD003669 | 20150309sp_7008_B13_a3_2.raw | Bacillus pseudomycoides | Bacillus toyonensis (CECT 876) | Yes | Yes |
| PXD003669 | 20150417sp_7165_B20_a1_1.raw | Bacillus cytotoxicus | Bacillus cereus (DSM 31) | Yes | Yes |
| PXD003669 | 20150417sp_7165_B20_a1_2.raw | Bacillus cytotoxicus | Bacillus cereus (DSM 31) | Yes | Yes |
| PXD003669 | 20150417sp_7165_B20_a2_1.raw | Bacillus cytotoxicus | Bacillus cereus (DSM 31) | Yes | Yes |
| PXD003669 | 20150417sp_7165_B20_a2_2.raw | Bacillus cytotoxicus | Bacillus cereus (DSM 31) | Yes | Yes |
| PXD003669 | 20150417sp_7165_B20_a3_1.raw | Bacillus cytotoxicus | Bacillus cereus (DSM 31) | Yes | Yes |
| PXD003669 | 20150417sp_7165_B20_a3_2.raw | Bacillus cytotoxicus | Bacillus cereus (DSM 31) | Yes | Yes |
| PXD003669 | 20150507sp_7221_B21a1_1.raw | Bacillus toyonensis | B. weihenstephanensis (DSM 11821) | Yes | Yes |
| PXD003669 | 20150507sp_7221_B21a1_2.raw | Bacillus toyonensis | B. weihenstephanensis (DSM 11821) | Yes | Yes |
| PXD003669 | 20150507sp_7221_B21a2_1.raw | Bacillus toyonensis | B. weihenstephanensis (DSM 11821) | Yes | Yes |
| PXD003669 | 20150507sp_7221_B21a2_2.raw | Bacillus toyonensis | B. weihenstephanensis (DSM 11821) | Yes | Yes |
| PXD003669 | 20150507sp_7221_B21a3_1.raw | Bacillus toyonensis | B. weihenstephanensis (DSM 11821) | Yes | Yes |
| PXD003669 | 20150507sp_7221_B21a3_2.raw | Bacillus toyonensis | B. weihenstephanensis (DSM 11821) | Yes | No |

| ProteomeXchange Identifier | Filename | Species Name | Strain Names | Cysteine Carbamylated? | Used in ROC Analysis? |
|---|---|---|---|---|---|
| PXD001860 | Biodiversity_A_cryptum_FeTSB_anaerobic_2_01Jun16_Pippin_16-03-39.raw | Acidiphilium cryptum | Acidiphilium cryptum JF-5 | No | No |
| PXD001860 | Biodiversity_A_muciniphila_test_27Feb17_Pippin_16-11-03.raw | Akkermansia muciniphila | Akkermansia muciniphila ATCC BAA-835 | No | Yes |
| PXD001860 | Biodiversity_A_tumefaciens_R2A_aerobic_2_23Nov16_Pippin_16-09-11.raw | Agrobacterium tumefaciens | Agrobacterium tumefaciens | No | No |
| PXD001860 | Biodiversity_A_tumefaciens_R2A_aerobic_3_23Nov16_Pippin_16-09-11.raw | Agrobacterium tumefaciens | Agrobacterium tumefaciens | No | No |
| PXD001860 | Biodiversity_A_faecalis_LB_aerobic_02_26Feb16_Arwen_16-01-01.raw | Alcaligenes faecalis | Alcaligenes faecalis | No | No |
| PXD001860 | Biodiversity_A_faecalis_LB_aerobic_01_26Feb16_Arwen_16-01-01.raw | Alcaligenes faecalis | Alcaligenes faecalis | No | No |
| PXD001860 | Biodiversity_B_cereus_PN_L_CL_3_09Oct16_Pippin_16-05-06.raw | Bacillus cereus | Bacillus cereus ATCC 14579 | No | Yes |
| PXD001860 | Biodiversity_B_cereus_PN_Z_CL_3_rr_11Oct16_Pippin_16-05-06.raw | Bacillus cereus | Bacillus cereus ATCC 14579 | No | Yes |
| PXD001860 | Biodiversity_Bacillus_subtilis_LB_02_27Dec15_Arwen_15-07-13.raw | Bacillus subtilis | Bacillus subtilis subsp. subtilis str. 168 | No | Yes |
| PXD001860 | Biodiversity_B_subtilis_NCIB3610_plates_1_03May16_Samwise_16-03-32.raw | Bacillus subtilis | Bacillus subtilis subsp. subtilis NCIB 3610 = ATCC 6051 | No | Yes |
| PXD001860 | Biodiversity_B_fragilis_Carb_01_28Oct15_Arwen_15-07-13.raw | Bacteroides fragilis | Bacteroides fragilis 638R | No | No |
| PXD001860 | Biodiversity_B_fragilis_CMgluc_anaerobic_03_01Feb16_Arwen_15-07-13.raw | Bacteroides fragilis | Bacteroides fragilis 638R | No | Yes |
| PXD001860 | Biodiversity_B_thet_CMgluc_anaerobic_03_01Feb16_Arwen_15-07-13.raw | Bacteroides thetaiotaomicron | Bacteroides thetaiotaomicron VPI-5482 | No | No |

| ProteomeXchange Identifier | Filename | Species Name | Strain Names | Cysteine Carbamylated? | Used in ROC Analysis? |
|---|---|---|---|---|---|
| PXD001860 | Biodiversity_B_bifidum_CMcarb_anaerobic_01_26Feb16_Arwen_16-01-01.raw | Bifidobacterium bifidum | Bifidobacterium bifidum ATCC 29521 = JCM 1255 = DSM 20456 | No | Yes |
| PXD001860 | Biodiversity_B_bifidum_CMcarb_anaerobic_03_26Feb16_Arwen_16-01-01.raw | Bifidobacterium bifidum | Bifidobacterium bifidum ATCC 29521 = JCM 1255 = DSM 20456 | No | Yes |
| PXD001860 | Biodiversity_B_infantis_CMcarb_anaerobic_03_26Feb16_Arwen_16-01-01.raw | Bifidobacterium longum | Bifidobacterium longum subsp. infantis ATCC 15697 = JCM 1222 = DSM 20088 | No | Yes |
| PXD001860 | Biodiversity_Cellulomonas_gilvus_GS2_02_27Dec15_Arwen_15-07-13.raw | Cellulomonas gilvus | Cellulomonas gilvus ATCC 13127 | No | Yes |
| PXD001860 | Biodiversity_Cellulomonas_gilvus_GS2_03_27Dec15_Arwen_15-07-13.raw | Cellulomonas gilvus | Cellulomonas gilvus ATCC 13127 | No | Yes |
| PXD001860 | Biodiversity_C_Baltica_T240_R1_C_27Jan16_Arwen_15-07-13.raw | Cellulophaga baltica | Cellulophaga baltica 18 | No | Yes |
| PXD001860 | Biodiversity_C_Baltica_T240_R3_Inf_27Jan16_Arwen_15-07-13.raw | Cellulophaga baltica | Cellulophaga baltica 18 | No | Yes |
| PXD001860 | Biodiversity_C_indologenes_LIB_aerobic_01_03May16_Samwise_16-03-32.raw | Chryseobacterium indologenes | Chryseobacterium indologenes | No | No |
| PXD001860 | Biodiversity_C_indologenes_LIB_aerobic_03_03May16_Samwise_16-03-32.raw | Chryseobacterium indologenes | Chryseobacterium indologenes | No | No |
| PXD001860 | Biodiversity_C_freundii_LIB_01_28Oct15_Arwen_15-07-13.raw | Citrobacter freundii | Citrobacter freundii | No | No |
| PXD001860 | Biodiversity_Cibrobacter_freundii_LB_aerobic_03_01Feb16_Arwen_15-07-13.raw | Citrobacter freundii | Citrobacter freundii | No | No |
| PXD001860 | Biodiversity_C_ljungdahlii_Fructose_anaerobic_1_04Oct16_Pippin_16-05-06.raw | Clostridium ljungdahlii | Clostridium ljungdahlii DSM 13528 | No | Yes |
| PXD001860 | Biodiversity_C_ljungdahlii_Fructose_anaerobic_2_04Oct16_Pippin_16-05-06.raw | Clostridium ljungdahlii | Clostridium ljungdahlii DSM 13528 | No | Yes |

| ProteomeXchange Identifier | Filename | Species Name | Strain Names | Cysteine Carbamylated? | Used in ROC Analysis? |
|---|---|---|---|---|---|
| PXD001860 | Biodiversity_C_necator_R2A_aerobic_2_23Nov16_Pippin_16-09-11.raw | Cupriavidus necator | Cupriavidus necator N-1 | No | Yes |
| PXD001860 | Biodiversity_C_necator_R2A_aerobic_1_23Nov16_Pippin_16-09-11.raw | Cupriavidus necator | Cupriavidus necator N-1 | No | Yes |
| PXD001860 | Biodiversity_HL69_HLA_aerobic_2_05Oct16_Pippin_16-05-06.raw | Cyanobacterium sp. HL-69 | Cyanobacterium sp. HL-69 | No | Yes |
| PXD001860 | Biodiversity_HL69_HLA_aerobic_1_05Oct16_Pippin_16-05-06.raw | Cyanobacterium sp. HL-69 | Cyanobacterium sp. HL-69 | No | Yes |
| PXD001860 | Biodiversity_D_acidovorans_TGY_aerobic_02_29Apr16_Samwise_16-03-32_renamed.raw | Delftia acidovorans | Delftia acidovorans SPH-1 | No | Yes |
| PXD001860 | Biodiversity_D_acidovorans_TGY_aerobic_03_29Apr16_Samwise_16-03-32_renamed.raw | Delftia acidovorans | Delftia acidovorans SPH-1 | No | Yes |
| PXD001860 | Biodiversity_Ecoli_PmrA_3d_TB7_4c_QEP_15Aug18_Wally_18-07-04.raw | Escherichia coli | Escherichia coli K-12 | No | Yes |
| PXD001860 | Biodiversity_Ecoli_PmrA_24h_TB7gluc_1b_QEP_15Aug18_Wally_18-07-04.raw | Escherichia coli | Escherichia coli K-12 | No | Yes |
| PXD001860 | Biodiversity_F_prausnitzii_Carb_01_28Oct15_Arwen_15-07-13.raw | Faecalibacterium prausnitzii | Faecalibacterium prausnitzii | No | Yes |
| PXD001860 | Biodiversity_F_prausnitzii_LIB_01_28Oct15_Arwen_15-07-13.raw | Faecalibacterium prausnitzii | Faecalibacterium prausnitzii | No | No |
| PXD001860 | Biodiversity_F_succinogenes_MDM_03_27Dec15_Arwen_15-07-13.raw | Fibrobacter succinogenes | Fibrobacter succinogenes subsp. succinogenes S85 | No | Yes |
| PXD001860 | Biodiversity_F_succinogenes_MDM_02_27Dec15_Arwen_15-07-13.raw | Fibrobacter succinogenes | Fibrobacter succinogenes subsp. succinogenes S85 | No | Yes |

| ProteomeXchange Identifier | Filename | Species Name | Strain Names | Cysteine Carbamylated? | Used in ROC Analysis? |
|---|---|---|---|---|---|
| PXD001860 | Biodiversity_F_novicida_TSB_aerobic_03_01Feb16_Arwen_15-07-13.raw | Francisella tularensis | Francisella tularensis subsp. novicida U112 | No | Yes |
| PXD001860 | Biodiversity_F_novicida_TSB_aerobic_01_01Feb16_Arwen_15-07-13.raw | Francisella tularensis | Francisella tularensis subsp. novicida U112 | No | Yes |
| PXD001860 | Biodiversity_Lactobacillus_casei_MRS_02_27Dec15_Arwen_15-07-13.raw | Lactobacillus casei | Lactobacillus casei | No | Yes |
| PXD001860 | Biodiversity_Lactobacillus_casei_MRS_03_27Dec15_Arwen_15-07-13.raw | Lactobacillus casei | Lactobacillus casei | No | Yes |
| PXD001860 | Biodiversity_L_monocytogenes_BHI_aerobic_01_27Feb17_Pippin_16-11-03.raw | Listeria monocytogenes | Listeria monocytogenes 10403S | No | Yes |
| PXD001860 | Biodiversity_L_monocytogenes_BHI_aerobic_03_27Feb17_Pippin_16-11-03.raw | Listeria monocytogenes | Listeria monocytogenes 10403S | No | Yes |
| PXD001860 | Biodiversity_M_luteus_LIB_aerobic_03_26Feb16_Arwen_16-01-01.raw | Micrococcus luteus | Micrococcus luteus | No | Yes |
| PXD001860 | Biodiversity_M_luteus_LIB_aerobic_01_26Feb16_Arwen_16-01-01.raw | Micrococcus luteus | Micrococcus luteus | No | Yes |
| PXD001860 | Biodiversity_M_smegmatis_BHI_aerobic_2_05Oct16_Pippin_16-05-06.raw | Mycolicibacterium smegmatis | Mycolicibacterium smegmatis | No | Yes |
| PXD001860 | Biodiversity_M_smegmatis_BHI_aerobic_1_05Oct16_Pippin_16-05-06.raw | Mycolicibacterium smegmatis | Mycolicibacterium smegmatis | No | Yes |
| PXD001860 | Biodiversity_M_xanthus_DZ2_48h_plates_2_13Jun16_Pippin_16-03-39.raw | Myxococcus xanthus | Myxococcus xanthus DZ2 | No | Yes |
| PXD001860 | Biodiversity_M_xanthus_DZ2_24h_plates_2_13Jun16_Pippin_16-03-39.raw | Myxococcus xanthus | Myxococcus xanthus DZ2 | No | Yes |

| ProteomeXchange Identifier | Filename | Species Name | Strain Names | Cysteine Carbamylated? | Used in ROC Analysis? |
|---|---|---|---|---|---|
| PXD001860 | Biodiversity_P_polymyxa_TBS_aerobic_2_17July16_Samwise_16-04-10.raw | Paenibacillus polymyxa | Paenibacillus polymyxa ATCC 842 | No | Yes |
| PXD001860 | Biodiversity_P_polymyxa_TBS_aerobic_3_17July16_Samwise_16-04-10.raw | Paenibacillus polymyxa | Paenibacillus polymyxa ATCC 842 | No | Yes |
| PXD001860 | Biodiversity_P_denitrificans_LIB_aerobic_02_29Apr16_Samwise_16-03-32_renamed.raw | Paracoccus denitrificans | Paracoccus denitrificans | No | Yes |
| PXD001860 | Biodiversity_P_denitrificans_LIB_aerobic_03_29Apr16_Samwise_16-03-32_renamed.raw | Paracoccus denitrificans | Paracoccus denitrificans | No | Yes |
| PXD001860 | Biodiversity_P_ruminicola_MDM_anaerobic_2_09Jun16_Pippin_16-03-39.raw | Prevotella ruminicola | Prevotella ruminicola 23 | No | No |
| PXD001860 | Biodiversity_P_ruminicola_MDM_anaerobic_1_09Jun16_Pippin_16-03-39.raw | Prevotella ruminicola | Prevotella ruminicola 23 | No | Yes |
| PXD001860 | Biodiversity_R_jostii_R2A_aerobic_1_23Nov16_Pippin_16-09-11.raw | Rhodococcus jostii | Rhodococcus jostii RHA1 | No | Yes |
| PXD001860 | Biodiversity_R_jostii_R2A_aerobic_2_23Nov16_Pippin_16-09-11.raw | Rhodococcus jostii | Rhodococcus jostii RHA1 | No | Yes |
| PXD001860 | Biodiversity_R_palustris_PMnitro_anaerobic_2_01Jun16_Pippin_16-03-39.raw | Rhodopseudomonas palustris | Rhodopseudomonas palustris | No | Yes |
| PXD001860 | Biodiversity_R_palustris_PMnonnitro_anaerobic_3_01Jun16_Pippin_16-03-39.raw | Rhodopseudomonas palustris | Rhodopseudomonas palustris | No | Yes |
| PXD001860 | Biodiversity_S_aurantiaca_CYE_aerobic_3_17July16_Samwise_16-04-10.raw | Stigmatella aurantiaca | Stigmatella aurantiaca DW4/3-1 | No | Yes |
| PXD001860 | Biodiversity_S_aurantiaca_CYE_aerobic_2_17July16_Samwise_16-04-10.raw | Stigmatella aurantiaca | Stigmatella aurantiaca DW4/3-1 | No | Yes |

| ProteomeXchange Identifier | Filename | Species Name | Strain Names | Cysteine Carbamylated? | Used in ROC Analysis? |
|---|---|---|---|---|---|
| PXD001860 | Biodiversity_S_agalactiae_LIB_aerobic_03_26Feb16_Arwen_16-01-01.raw | Streptococcus agalactiae | Streptococcus agalactiae | No | Yes |
| PXD001860 | Biodiversity_S_agalactiae_LIB_aerobic_01_26Feb16_Arwen_16-01-01.raw | Streptococcus agalactiae | Streptococcus agalactiae | No | Yes |
| PXD001860 | Biodiversity_S_elongatus_BG11_aerobic_3_14July16_Pippin_16-05-01.raw | Synechococcus elongatus | Synechococcus elongatus PCC 7942 = FACHB-805 | No | Yes |
| PXD001860 | Biodiversity_S_elongatus_BG11NaCl_aerobic_3_05Oct16_Pippin_16-05-06.raw | Synechococcus elongatus | Synechococcus elongatus PCC 7942 = FACHB-805 | No | Yes |

Table S3. Species Included in the MARLOWE Database Used in this Study. MiCId used an identical database.

| NCBI Taxon Identifier | Organism Name |
|---|---|
| 535289 | [Acidovorax] ebreus TPSY |
| 1909395 | [Actinomadura] parvosata subsp. kistnae |
| 1771959 | [Arthrobacter] sp. ATCC 21022 |
| 281309 | [Bacillus thuringiensis] serovar konkukian str. 97-27 |
| 439292 | [Bacillus] selenitireducens MLS10 |
| 1232385 | [Brevibacterium] flavum ZL-1 |
| 29343 | [Clostridium] cellulosi |
| 717608 | [Clostridium] cf. saccharolyticum K10 |
| 610130 | [Clostridium] saccharolyticum WM1 |
| 1334193 | [Enterobacter] lignolyticus |
| 701347 | [Enterobacter] lignolyticus SCF1 |
| 515620 | [Eubacterium] eligens ATCC 27750 |
| 657319 | [Eubacterium] siraeum 70/3 |
| 717961 | [Eubacterium] siraeum V10Sc8a |
| 233412 | [Haemophilus] ducreyi 35000HP |
| 111781 | [Leptolyngbya] sp. PCC 7376 |
| 221988 | [Mannheimia] succiniciproducens MBEL55E |
| 1520670 | [Mycobacterium] stephanolepidis |
| 142651 | [Mycoplasma] phocae |
| 657313 | [Ruminococcus] torques L2-14 |
| 1817405 | Abyssicoccus albus |
| 1257118 | Acanthamoeba castellanii str. Neff |
| 133434 | Acanthaster planci |
| 329726 | Acaryochloris marina MBIC11017 |
| 720554 | Acetivibrio clariflavus DSM 19732 |
| 1677857 | Acetivibrio saccincola |
| 203119 | Acetivibrio thermocellus ATCC 27405 |
| 637887 | Acetivibrio thermocellus DSM 1313 |
| 499177 | Acetoanaerobium sticklandii DSM 519 |
| 435 | Acetobacter aceti |
| 481146 | Acetobacter ascendens |
| 1633874 | Acetobacter oryzifermentans |
| 1266844 | Acetobacter pasteurianus 386B |
| 634452 | Acetobacter pasteurianus IFO 3283-01 |
| 634458 | Acetobacter pasteurianus IFO 3283-01-42C |
| 634453 | Acetobacter pasteurianus IFO 3283-03 |
| 634454 | Acetobacter pasteurianus IFO 3283-07 |
| 634459 | Acetobacter pasteurianus IFO 3283-12 |

| NCBI Taxon Identifier | Organism Name |
|---|---|
| 634455 | Acetobacter pasteurianus IFO 3283-22 |
| 634456 | Acetobacter pasteurianus IFO 3283-26 |
| 634457 | Acetobacter pasteurianus IFO 3283-32 |
| 1076596 | Acetobacter persici |
| 65959 | Acetobacter pomorum |
| 446692 | Acetobacter senegalensis |
| 2259883 | Acetobacter sp. JWB |
| 104102 | Acetobacter tropicalis |
| 931626 | Acetobacterium woodii DSM 1030 |
| 574087 | Acetohalobium arabaticum DSM 5501 |
| 891968 | Acetomicrobium mobile DSM 13181 |
| 441768 | Acholeplasma laidlawii PG-8A |
| 35623 | Acholeplasma oculi |
| 32002 | Achromobacter denitrificans |
| 217204 | Achromobacter insolitus |
| 1758194 | Achromobacter sp. AONIH1 |
| 2282475 | Achromobacter sp. B7 |
| 217203 | Achromobacter spanius |
| 85698 | Achromobacter xylosoxidans |
| 762376 | Achromobacter xylosoxidans A8 |
| 1216976 | Achromobacter xylosoxidans NBRC 15126 = ATCC 27061 |
| 1167634 | Achromobacter xylosoxidans NH44784-1996 |
| 591001 | Acidaminococcus fermentans DSM 20731 |
| 568816 | Acidaminococcus intestini RyC-MR95 |
| 41673 | Acidianus brierleyi |
| 933801 | Acidianus hospitalis W1 |
| 282676 | Acidianus manzaensis |
| 619593 | Acidianus sulfidivorans JP7 |
| 1281578 | Acidiferrobacter sp. SPIII_3 |
| 160660 | Acidihalobacter prosperus |
| 666510 | Acidilobus saccharovorans 345-15 |
| 1577685 | Acidilobus sp. 7A |
| 2507161 | Acidilutibacter cellobiosedens |
| 525909 | Acidimicrobium ferrooxidans DSM 10331 |
| 349163 | Acidiphilium cryptum JF-5 |
| 926570 | Acidiphilium multivorum AIU301 |
| 1748 | Acidipropionibacterium acidipropionici |
| 1171373 | Acidipropionibacterium acidipropionici ATCC 4875 |
| 1749 | Acidipropionibacterium jensenii |
| 2057246 | Acidipropionibacterium virtanenii |
| 2211140 | Acidisarcina polymorpha |

| NCBI Taxon Identifier | Organism Name |
|---|---|
| 637389 | Acidithiobacillus caldus ATCC 51756 |
| 990288 | Acidithiobacillus caldus SM-1 |
| 1232575 | Acidithiobacillus ferridurans |
| 743299 | Acidithiobacillus ferrivorans SS3 |
| 243159 | Acidithiobacillus ferrooxidans ATCC 23270 |
| 380394 | Acidithiobacillus ferrooxidans ATCC 53993 |
| 240015 | Acidobacterium capsulatum ATCC 51196 |
| 351607 | Acidothermus cellulolyticus 11B |
| 553814 | Acidovorax carolinensis |
| 553814 | Acidovorax carolinensis |
| 553814 | Acidovorax carolinensis |
| 553814 | Acidovorax carolinensis |
| 2478662 | Acidovorax sp. 1608163 |
| 232721 | Acidovorax sp. JS42 |
| 358220 | Acidovorax sp. KKS102 |
| 1842533 | Acidovorax sp. RAC01 |
| 439481 | Aciduliprofundum boonei T469 |
| 673860 | Aciduliprofundum sp. MAR08-339 |
| 470 | Acinetobacter baumannii |
| 470 | Acinetobacter baumannii |
| 470 | Acinetobacter baumannii |
| 470 | Acinetobacter baumannii |
| 696749 | Acinetobacter baumannii 1656-2 |
| 480119 | Acinetobacter baumannii AB0057 |
| 557600 | Acinetobacter baumannii AB307-0294 |
| 405416 | Acinetobacter baumannii ACICU |
| 400667 | Acinetobacter baumannii ATCC 17978 |
| 509173 | Acinetobacter baumannii AYE |
| 1096995 | Acinetobacter baumannii BJAB07104 |
| 1096996 | Acinetobacter baumannii BJAB0715 |
| 1096997 | Acinetobacter baumannii BJAB0868 |
| 945556 | Acinetobacter baumannii D1279779 |
| 1455315 | Acinetobacter baumannii LAC-4 |
| 889738 | Acinetobacter baumannii MDR-TJ |
| 497978 | Acinetobacter baumannii MDR-ZJ06 |
| 509170 | Acinetobacter baumannii SDF |
| 980514 | Acinetobacter baumannii TCDC-AB0715 |
| 1100841 | Acinetobacter baumannii TYTH-1 |
| 1400867 | Acinetobacter baumannii ZW85-1 |
| 62977 | Acinetobacter baylyi ADP1 |
| 471 | Acinetobacter calcoaceticus |

| NCBI Taxon Identifier | Organism Name |
|---|---|
| 1871111 | Acinetobacter defluvii |
| 1324350 | Acinetobacter equi |
| 29430 | Acinetobacter haemolyticus |
| 756892 | Acinetobacter indicus |
| 1242245 | Acinetobacter johnsonii XBB1 |
| 40215 | Acinetobacter junii |
| 1785128 | Acinetobacter lactucae |
| 1789224 | Acinetobacter larvae |
| 106654 | Acinetobacter nosocomialis |
| 436717 | Acinetobacter oleivorans DR1 |
| 871585 | Acinetobacter pittii PHEA-2 |
| 40216 | Acinetobacter radioresistens |
| 108981 | Acinetobacter schindleri |
| 487316 | Acinetobacter soli |
| 1809055 | Acinetobacter sp. DUT-2 |
| 1407071 | Acinetobacter sp. TGL-Y2 |
| 1646498 | Acinetobacter sp. TTH0-4 |
| 1879050 | Acinetobacter wuhouensis |
| 32536 | Acinonyx jubatus |
| 103372 | Acromyrmex echinatior |
| 70779 | Acropora digitifera |
| 1609966 | Actibacterium sp. EMB200-NS6 |
| 1612552 | Actinoalloteichus fjordicus |
| 1470176 | Actinoalloteichus hoggarensis |
| 340345 | Actinoalloteichus hymeniacidonis |
| 2072503 | Actinoalloteichus sp. AHMU CJ021 |
| 1612551 | Actinoalloteichus sp. GBA129-24 |
| 202947 | Actinobacillus equuli subsp. equuli |
| 434271 | Actinobacillus pleuropneumoniae serovar 3 str. JL03 |
| 416269 | Actinobacillus pleuropneumoniae serovar 5b str. L20 |
| 537457 | Actinobacillus pleuropneumoniae serovar 7 str. AP76 |
| 189834 | Actinobacillus porcitonsillarum |
| 339671 | Actinobacillus succinogenes 130Z |
| 743972 | Actinobacillus suis ATCC 33415 |
| 696748 | Actinobacillus suis H91-0380 |
| 1650658 | Actinobacteria bacterium IMCC26256 |
| 2495645 | Actinobaculum sp. 313 |
| 1960083 | Actinomyces gaoshouyii |
| 1655 | Actinomyces naeslundii |
| 544580 | Actinomyces oris |
| 111015 | Actinomyces radicidentis |

| NCBI Taxon Identifier | Organism Name |
|---|---|
| 1851395 | Actinomyces sp. Chiba101 |
| 712122 | Actinomyces sp. oral taxon 414 |
| 2081702 | Actinomyces sp. oral taxon 897 |
| 512565 | Actinoplanes missouriensis 431 |
| 649831 | Actinoplanes sp. N902-109 |
| 2033844 | Actinoplanes sp. SE50 |
| 134676 | Actinoplanes sp. SE50/110 |
| 414996 | Actinopolyspora erythraea |
| 446462 | Actinosynnema mirum DSM 43827 |
| 42197 | Actinosynnema pretiosum |
| 59505 | Actinotignum schaalii |
| 7029 | Acyrthosiphon pisum |
| 1384484 | Adlercreutzia equolifaciens DSM 19450 |
| 1036672 | Advenella kashmirensis WT001 |
| 1247726 | Advenella mimigardefordensis DPN7 |
| 7159 | Aedes aegypti |
| 169297 | Aegilops tauschii subsp. tauschii |
| 2494375 | Aequorivita sp. H23M31 |
| 746697 | Aequorivita sublithincola DSM 14238 |
| 33936 | Aeribacillus pallidus |
| 87541 | Aerococcus christensenii |
| 119206 | Aerococcus sanguinicola |
| 2976812 | Aerococcus sp. Group 1 |
| 1376 | Aerococcus urinae |
| 51665 | Aerococcus urinaeequi |
| 128944 | Aerococcus urinaehominis |
| 1377 | Aerococcus viridans |
| 2079793 | Aeromicrobium chenweiae |
| 2041 | Aeromicrobium erythreum |
| 2107713 | Aeromicrobium sp. A1-2 |
| 648 | Aeromonas caviae |
| 196024 | Aeromonas dhakensis |
| 644 | Aeromonas hydrophila |
| 644 | Aeromonas hydrophila |
| 644 | Aeromonas hydrophila |
| 1419584 | Aeromonas hydrophila J-1 |
| 1288394 | Aeromonas hydrophila ML09-119 |
| 1418107 | Aeromonas hydrophila pc104A |
| 1321367 | Aeromonas hydrophila subsp. hydrophila AL09-71 |
| 380703 | Aeromonas hydrophila subsp. hydrophila ATCC 7966 |
| 1448139 | Aeromonas hydrophila YL17 |

| NCBI Taxon Identifier | Organism Name |
|---|---|
| 1208104 | Aeromonas media WS |
| 948519 | Aeromonas rivipollensis |
| 645 | Aeromonas salmonicida |
| 382245 | Aeromonas salmonicida subsp. salmonicida A449 |
| 652 | Aeromonas schubertii |
| 1636608 | Aeromonas sp. ASNIH3 |
| 1758179 | Aeromonas sp. ASNIH5 |
| 2033033 | Aeromonas sp. CU5 |
| 654 | Aeromonas veronii |
| 998088 | Aeromonas veronii B565 |
| 1198449 | Aeropyrum camini SY1 = JCM 12091 |
| 272557 | Aeropyrum pernix K1 |
| 1031710 | Afipia carboxidovorans OM4 |
| 504832 | Afipia carboxidovorans OM5 |
| 504832 | Afipia carboxidovorans OM5 |
| 936046 | Agaricus bisporus var. bisporus H97 |
| 597362 | Agaricus bisporus var. burnettii JB137-S8 |
| 515619 | Agathobacter rectalis ATCC 33656 |
| 657318 | Agathobacter rectalis DSM 17629 |
| 657317 | Agathobacter rectalis M104/1 |
| 714 | Aggregatibacter actinomycetemcomitans |
| 754507 | Aggregatibacter actinomycetemcomitans ANH9381 |
| 668336 | Aggregatibacter actinomycetemcomitans D11S-1 |
| 694569 | Aggregatibacter actinomycetemcomitans D7S-1 |
| 272556 | Aggregatibacter actinomycetemcomitans HK1651 |
| 1407647 | Aggregatibacter actinomycetemcomitans NUM4039 |
| 732 | Aggregatibacter aphrophilus |
| 634176 | Aggregatibacter aphrophilus NJ8700 |
| 176299 | Agrobacterium fabrum str. C58 |
| 1842536 | Agrobacterium sp. RAC06 |
| 358 | Agrobacterium tumefaciens |
| 358 | Agrobacterium tumefaciens |
| 358 | Agrobacterium tumefaciens |
| 453304 | Agromyces aureus |
| 2080742 | Agromyces badenianii |
| 2509455 | Agromyces protaetiae |
| 2021234 | Ahniella affigens |
| 9646 | Ailuropoda melanoleuca |
| 1679444 | Akkermansia glycaniphila |
| 349741 | Akkermansia muciniphila ATCC BAA-835 |
| 323284 | Alcaligenes aquatilis |

| NCBI Taxon Identifier | Organism Name |
|---|---|
| 511 | Alcaligenes faecalis |
| 511 | Alcaligenes faecalis |
| 393595 | Alcanivorax borkumensis SK2 |
| 1113728 | Alcanivorax sp. NBRC 101098 |
| 1727163 | Algoriphagus sanaruensis |
| 1036172 | Aliarcobacter butzleri 7h1h |
| 944546 | Aliarcobacter butzleri ED-1 |
| 367737 | Aliarcobacter butzleri RM4018 |
| 28200 | Aliarcobacter skirrowii |
| 596153 | Alicycliphilus denitrificans BC |
| 596154 | Alicycliphilus denitrificans K601 |
| 521098 | Alicyclobacillus acidocaldarius subsp. acidocaldarius DSM 446 |
| 1048834 | Alicyclobacillus acidocaldarius subsp. acidocaldarius Tc-4-1 |
| 312309 | Aliivibrio fischeri ES114 |
| 388396 | Aliivibrio fischeri MJ11 |
| 316275 | Aliivibrio salmonicida LFI1238 |
| 80852 | Aliivibrio wodanis |
| 679935 | Alistipes finegoldii DSM 17242 |
| 717959 | Alistipes shahii WAL 8301 |
| 79881 | Alkalicoccobacillus gibsonii |
| 398511 | Alkalihalophilus pseudofirmus OF4 |
| 187272 | Alkalilimnicola ehrlichii MLHE-1 |
| 293826 | Alkaliphilus metalliredigens QYMF |
| 350688 | Alkaliphilus oremlandii OhILAs |
| 889453 | Alkalitalea saponilacus |
| 8496 | Alligator mississippiensis |
| 38654 | Alligator sinensis |
| 1653480 | Alloactinosynnema sp. L-07 |
| 930169 | Alloalcanivorax dieselolei B5 |
| 1094342 | Alloalcanivorax xenomutans |
| 572477 | Allochromatium vinosum DSM 180 |
| 1173027 | Allocoleopsis franciscana PCC 7113 |
| 594679 | Allofrancisella guangzhouensis |
| 526227 | Allomeiothermus silvanus DSM 9946 |
| 886377 | Allomuricauda ruestringensis DSM 13258 |
| 311402 | Allorhizobium ampelinum S4 |
| 311180 | Alloyangia pacifica |
| 859653 | alpha proteobacterium HIMB5 |
| 744985 | alpha proteobacterium HIMB59 |
| 1318466 | Alteracholeplasma palmae J233 |
| 361183 | Altererythrobacter epoxidivorans |

| NCBI Taxon Identifier | Organism Name |
|---|---|
| 2060312 | Altererythrobacter sp. B11 |
| 5599 | Alternaria alternata |
| 589873 | Alteromonas australica |
| 589873 | Alteromonas australica |
| 529120 | Alteromonas macleodii ATCC 27126 |
| 1004787 | Alteromonas macleodii str. 'Balearic Sea AD45' |
| 1004785 | Alteromonas macleodii str. 'Black Sea 11' |
| 1004788 | Alteromonas macleodii str. 'English Channel 673' |
| 1300253 | Alteromonas mediterranea 615 |
| 1774373 | Alteromonas mediterranea DE |
| 1004786 | Alteromonas mediterranea DE1 |
| 1300254 | Alteromonas mediterranea MED64 |
| 1300255 | Alteromonas mediterranea U4 |
| 1300256 | Alteromonas mediterranea U7 |
| 1300257 | Alteromonas mediterranea U8 |
| 1300259 | Alteromonas mediterranea UM4b |
| 1300258 | Alteromonas mediterranea UM7 |
| 715451 | Alteromonas naphthalenivorans |
| 1777491 | Alteromonas sp. Mac1 |
| 2267264 | Alteromonas sp. RKMC-009 |
| 233316 | Alteromonas stellipolaris |
| 233316 | Alteromonas stellipolaris |
| 1160720 | Alteromonas stellipolaris LMG 21856 |
| 13333 | Amborella trichopoda |
| 2507160 | Aminipila luticellarii |
| 83263 | Aminobacter aminovorans |
| 374606 | Aminobacter sp. MSH1 |
| 572547 | Aminobacterium colombiense DSM 12261 |
| 429009 | Ammonifex degensii KC4 |
| 1246995 | Amorphoplanes friuliensis DSM 7358 |
| 698758 | Amphibacillus xylanus NBRC 15112 |
| 400682 | Amphimedon queenslandica |
| 1804986 | Amycolatopsis albispora |
| 208439 | Amycolatopsis japonica |
| 1221524 | Amycolatopsis mediterranei RB |
| 713604 | Amycolatopsis mediterranei S699 |
| 713604 | Amycolatopsis mediterranei S699 |
| 749927 | Amycolatopsis mediterranei U32 |
| 1068978 | Amycolatopsis methanolica 239 |
| 1156913 | Amycolatopsis orientalis HCCB10007 |
| 1896961 | Amycolatopsis sp. AA4 |

| NCBI Taxon Identifier | Organism Name |
|---|---|
| 1911175 | Amycolatopsis sp. BJA-103 |
| 1423721 | Amylolactobacillus amylophilus DSM 20533 = JCM 1125 |
| 272123 | Anabaena cylindrica PCC 7122 |
| 46234 | Anabaena sp. 90 |
| 1647413 | Anabaena sp. WA102 |
| 39488 | Anaerobutyricum hallii |
| 525919 | Anaerococcus prevotii DSM 20548 |
| 926569 | Anaerolinea thermophila UNI-1 |
| 1889813 | Anaerolineaceae bacterium oral taxon 439 |
| 455488 | Anaeromyxobacter dehalogenans 2CP-1 |
| 290397 | Anaeromyxobacter dehalogenans 2CP-C |
| 404589 | Anaeromyxobacter sp. Fw109-5 |
| 447217 | Anaeromyxobacter sp. K |
| 649756 | Anaerostipes hadrus |
| 991789 | Anaerotignum propionicum DSM 1682 |
| 574556 | Anaplasma centrale str. Israel |
| 1412841 | Anaplasma marginale str. Dawn |
| 320483 | Anaplasma marginale str. Florida |
| 1412840 | Anaplasma marginale str. Gypsy Plains |
| 234826 | Anaplasma marginale str. St. Maries |
| 1248439 | Anaplasma ovis str. Haibei |
| 1184254 | Anaplasma phagocytophilum str. Dog2 |
| 212042 | Anaplasma phagocytophilum str. HZ |
| 1184253 | Anaplasma phagocytophilum str. HZ2 |
| 1173064 | Anaplasma phagocytophilum str. JM |
| 8839 | Anas platyrhynchos |
| 639283 | Ancylobacter novellus DSM 506 |
| 1500254 | Aneurinibacillus soli |
| 1450761 | Aneurinibacillus sp. XH2 |
| 28377 | Anolis carolinensis |
| 180454 | Anopheles gambiae str. PEST |
| 294699 | Anoxybacillus amylolyticus |
| 491915 | Anoxybacillus flavithermus WK1 |
| 198467 | Anoxybacillus gonensis |
| 1490052 | Anoxybacillus sp. B2M1 |
| 1490057 | Anoxybacillus sp. B7M1 |
| 1323375 | Anoxybacter fermentans |
| 8845 | Anser cygnoides |
| 74033 | Antarctobacter heliothermus |
| 80765 | Aphis gossypii |
| 148814 | Apilactobacillus kunkeei |

| NCBI Taxon Identifier | Organism Name |
|---|---|
| 7460 | Apis mellifera |
| 202946 | Apteryx mantelli mantelli |
| 1296669 | Aquabacterium olei |
| 1938604 | Aquaspirillum sp. LM1 |
| 2067065 | Aquella oligotrophica |
| 1670800 | Aquibium oceanicum |
| 224324 | Aquifex aeolicus VF5 |
| 1714848 | Aquimarina sp. AD1 |
| 1714849 | Aquimarina sp. AD10 |
| 1714860 | Aquimarina sp. BL5 |
| 2283318 | Aquirhabdus parva |
| 2516557 | Aquirufa nivalisilvae |
| 332411 | Aquitalea magnusonii |
| 1590041 | Aquitalea sp. USM4 |
| 81972 | Arabidopsis lyrata subsp. lyrata |
| 3702 | Arabidopsis thaliana |
| 2341117 | Arachidicoccus soli |
| 1850526 | Arachidicoccus sp. BS20 |
| 130453 | Arachis duranensis |
| 130454 | Arachis ipaensis |
| 767029 | Arachnia propionica F0230a |
| 644284 | Arcanobacterium haemolyticum DSM 20595 |
| 224325 | Archaeoglobus fulgidus DSM 4304 |
| 1344584 | Archaeoglobus fulgidus DSM 8774 |
| 572546 | Archaeoglobus profundus DSM 5631 |
| 387631 | Archaeoglobus sulfaticallidus PM70-1 |
| 693661 | Archaeoglobus veneficus SNP6 |
| 1579370 | archaeon GW2011_AR10 |
| 1579378 | archaeon GW2011_AR20 |
| 48 | Archangium gephyra |
| 572480 | Arcobacter nitrofigilis DSM 7299 |
| 944547 | Arcobacter sp. L |
| 1784714 | Arcticibacterium luteifluviistationis |
| 616991 | Arenibacter algicola |
| 76114 | Aromatoleum aromaticum EbN1 |
| 1658671 | Arsenicicoccus sp. oral taxon 190 |
| 235559 | Arsenophonus endosymbiont of Aleurodicus dispersus |
| 656366 | Arthrobacter alpinus |
| 656366 | Arthrobacter alpinus |
| 37928 | Arthrobacter crystallopoietes |
| 1704044 | Arthrobacter sp. ERGS1:01 |

| NCBI Taxon Identifier | Organism Name |
|---|---|
| 290399 | Arthrobacter sp. FB24 |
| 1588023 | Arthrobacter sp. Hiyo8 |
| 1690248 | Arthrobacter sp. LS16 |
| 1494608 | Arthrobacter sp. PAMC 25486 |
| 2079227 | Arthrobacter sp. PGP41 |
| 1357915 | Arthrobacter sp. QXT-31 |
| 1118963 | Arthrobacter sp. Rue61a |
| 1849032 | Arthrobacter sp. U41 |
| 1652545 | Arthrobacter sp. YC-RL1 |
| 2020486 | Arthrobacter sp. YN |
| 1806905 | Arthrobacter sp. ZXY-2 |
| 696747 | Arthrospira platensis NIES-39 |
| 1231624 | Asaia bogorensis NBRC 16594 |
| 4686 | Asparagus officinalis |
| 344612 | Aspergillus clavatus NRRL 1 |
| 331117 | Aspergillus fischeri NRRL 181 |
| 332952 | Aspergillus flavus NRRL3357 |
| 330879 | Aspergillus fumigatus Af293 |
| 227321 | Aspergillus nidulans FGSC A4 |
| 425011 | Aspergillus niger CBS 513.88 |
| 510516 | Aspergillus oryzae RIB40 |
| 322098 | Aster yellows witches'-broom phytoplasma AYWB |
| 573065 | Asticcacaulis excentricus CB 48 |
| 7994 | Astyanax mexicanus |
| 194338 | Athene cunicularia |
| 565 | Atlantibacter hermannii |
| 12957 | Atta cephalotes |
| 1648404 | Aurantiacibacter atlanticus |
| 502682 | Aurantiacibacter gangjinensis |
| 708131 | Aurantimicrobium minutum |
| 1987356 | Aurantimicrobium photophilum |
| 2079575 | Aurantimicrobium sp. MWH-Uga1 |
| 1349819 | Aureimonas sp. AU20 |
| 44056 | Aureococcus anophagefferens |
| 717982 | Auricularia subglabra TFB-10046 SS5 |
| 2170745 | Auritidibacter sp. NML130574 |
| 3075 | Auxenochlorella protothecoides |
| 728 | Avibacterium paragallinarum |
| 418699 | Azoarcus olearius |
| 418699 | Azoarcus olearius |
| 198107 | Azoarcus sp. CIB |

| NCBI Taxon Identifier | Organism Name |
|---|---|
| 356837 | Azoarcus sp. DN11 |
| 748247 | Azoarcus sp. KH32C |
| 438753 | Azorhizobium caulinodans ORS 571 |
| 640081 | Azospira oryzae PS |
| 1064539 | Azospirillum baldaniorum |
| 192 | Azospirillum brasilense |
| 192 | Azospirillum brasilense |
| 1226968 | Azospirillum humicireducens |
| 862719 | Azospirillum lipoferum 4B |
| 682998 | Azospirillum ramasamyi |
| 137722 | Azospirillum sp. B510 |
| 664962 | Azospirillum sp. TSH58 |
| 528244 | Azospirillum thiophilum |
| 1328314 | Azotobacter chroococcum NCIMB 8003 |
| 1283330 | Azotobacter vinelandii CA |
| 1283331 | Azotobacter vinelandii CA6 |
| 322710 | Azotobacter vinelandii DJ |
| 484906 | Babesia bovis T2Bo |
| 1133968 | Babesia microti strain RI |
| 293387 | Bacillus altitudinis |
| 1412898 | Bacillus amyloliquefaciens CC178 |
| 692420 | Bacillus amyloliquefaciens DSM 7 = ATCC 23350 |
| 1091041 | Bacillus amyloliquefaciens IT-45 |
| 1415165 | Bacillus amyloliquefaciens LFB112 |
| 1001582 | Bacillus amyloliquefaciens LL3 |
| 999891 | Bacillus amyloliquefaciens TA208 |
| 1034836 | Bacillus amyloliquefaciens XH7 |
| 1392 | Bacillus anthracis |
| 592021 | Bacillus anthracis str. A0248 |
| 743835 | Bacillus anthracis str. A16 |
| 673518 | Bacillus anthracis str. A16R |
| 198094 | Bacillus anthracis str. Ames |
| 261594 | Bacillus anthracis str. 'Ames Ancestor' |
| 568206 | Bacillus anthracis str. CDC 684 |
| 768494 | Bacillus anthracis str. H9401 |
| 260799 | Bacillus anthracis str. Sterne |
| 1392837 | Bacillus anthracis str. SVA11 |
| 261591 | Bacillus anthracis str. Vollum |
| 720555 | Bacillus atrophaeus 1942 |
| 1330043 | Bacillus bombysepticus str. Wang |
| 1396 | Bacillus cereus |

| NCBI Taxon Identifier | Organism Name |
|---|---|
| 572264 | Bacillus cereus 03BB102 |
| 405534 | Bacillus cereus AH187 |
| 405535 | Bacillus cereus AH820 |
| 222523 | Bacillus cereus ATCC 10987 |
| 226900 | Bacillus cereus ATCC 14579 |
| 405532 | Bacillus cereus B4264 |
| 637380 | Bacillus cereus biovar anthracis str. CI |
| 288681 | Bacillus cereus E33L |
| 347495 | Bacillus cereus F837/76 |
| 1217984 | Bacillus cereus FRI-35 |
| 405531 | Bacillus cereus G9842 |
| 334406 | Bacillus cereus NC7401 |
| 361100 | Bacillus cereus Q1 |
| 315749 | Bacillus cytotoxicus NVH 391-98 |
| 1664069 | Bacillus glycinifermentans |
| 1367477 | Bacillus infantis NRRL B-14911 |
| 279010 | Bacillus licheniformis DSM 13 = ATCC 14580 |
| 279010 | Bacillus licheniformis DSM 13 = ATCC 14580 |
| 796606 | Bacillus methanolicus MGA3 |
| 1405 | Bacillus mycoides |
| 1405 | Bacillus mycoides |
| 315730 | Bacillus mycoides KBAB4 |
| 766760 | Bacillus paralicheniformis ATCC 9945a |
| 64104 | Bacillus pseudomycoides |
| 1408 | Bacillus pumilus |
| 1408 | Bacillus pumilus |
| 315750 | Bacillus pumilus SAFR-032 |
| 561879 | Bacillus safensis |
| 1479 | Bacillus smithii |
| 666686 | Bacillus sp. 1NLA3E |
| 1570330 | Bacillus sp. BH072 |
| 1806506 | Bacillus sp. BS34A |
| 1127744 | Bacillus sp. JS |
| 1628753 | Bacillus sp. LM 4-2 |
| 98228 | Bacillus sp. OxB-1 |
| 1446792 | Bacillus sp. Pc3 |
| 1774743 | Bacillus sp. SDLI1 |
| 756828 | Bacillus sp. WP8 |
| 1565991 | Bacillus sp. X1(2014) |
| 1574141 | Bacillus sp. YP1 |
| 96241 | Bacillus spizizenii |

| NCBI Taxon Identifier | Organism Name |
|---|---|
| 655816 | Bacillus spizizenii str. W23 |
| 1052585 | Bacillus spizizenii TU-B-10 |
| 936156 | Bacillus subtilis BSn5 |
| 1415167 | Bacillus subtilis PY79 |
| 1220533 | Bacillus subtilis QB928 |
| 645657 | Bacillus subtilis subsp. natto BEST195 |
| 1147161 | Bacillus subtilis subsp. subtilis 6051-HGW |
| 224308 | Bacillus subtilis subsp. subtilis str. 168 |
| 1221328 | Bacillus subtilis subsp. subtilis str. AG1839 |
| 1302650 | Bacillus subtilis subsp. subtilis str. BAB-1 |
| 1192196 | Bacillus subtilis subsp. subtilis str. BSP1 |
| 1232554 | Bacillus subtilis subsp. subtilis str. JH642 substr. AG174 |
| 1404258 | Bacillus subtilis subsp. subtilis str. OH 131.1 |
| 1052588 | Bacillus subtilis subsp. subtilis str. RO-NN-1 |
| 1233100 | Bacillus subtilis XF-1 |
| 1428 | Bacillus thuringiensis |
| 1428 | Bacillus thuringiensis |
| 714359 | Bacillus thuringiensis BMB171 |
| 527021 | Bacillus thuringiensis Bt407 |
| 1218175 | Bacillus thuringiensis HD-771 |
| 1217737 | Bacillus thuringiensis HD-789 |
| 1195464 | Bacillus thuringiensis MC28 |
| 541229 | Bacillus thuringiensis serovar chinensis CT-43 |
| 930170 | Bacillus thuringiensis serovar finitimus YBT-020 |
| 1261129 | Bacillus thuringiensis serovar kurstaki str. HD-1 |
| 1279365 | Bacillus thuringiensis serovar kurstaki str. HD73 |
| 570416 | Bacillus thuringiensis serovar kurstaki str. YBT-1520 |
| 1286404 | Bacillus thuringiensis serovar thuringiensis str. IS5056 |
| 412694 | Bacillus thuringiensis str. Al Hakam |
| 529122 | Bacillus thuringiensis YBT-1518 |
| 1415784 | Bacillus toyonensis BCT-7112 |
| 72361 | Bacillus vallismortis |
| 492670 | Bacillus velezensis |
| 1225788 | Bacillus velezensis AS43.3 |
| 1114958 | Bacillus velezensis CAU B946 |
| 326423 | Bacillus velezensis FZB42 |
| 1385727 | Bacillus velezensis NAU-B3 |
| 1423138 | Bacillus velezensis SQR9 |
| 1449088 | Bacillus velezensis TrigoCor1448 |
| 1338518 | Bacillus velezensis UCMB5033 |
| 1150475 | Bacillus velezensis UCMB5036 |

| NCBI Taxon Identifier | Organism Name |
|---|---|
| 1150476 | Bacillus velezensis UCMB5113 |
| 1155777 | Bacillus velezensis YAU B9601-Y2 |
| 1155777 | Bacillus velezensis YAU B9601-Y2 |
| 1547283 | Bacillus weihaiensis |
| 2494319 | Bacillus wiedmannii bv. thuringiensis |
| 1178537 | Bacillus xiamenensis |
| 1249553 | Bacterioplanes sanyensis |
| 960 | Bacteriovorax stolpii |
| 1898108 | bacterium AB1 |
| 1400053 | Bacteroidales bacterium CF |
| 47678 | Bacteroides caccae |
| 1796613 | Bacteroides caecimuris |
| 246787 | Bacteroides cellulosilyticus |
| 817 | Bacteroides fragilis |
| 862962 | Bacteroides fragilis 638R |
| 272559 | Bacteroides fragilis NCTC 9343 |
| 295405 | Bacteroides fragilis YCH46 |
| 693979 | Bacteroides helcogenes P 36-108 |
| 28113 | Bacteroides heparinolyticus |
| 28116 | Bacteroides ovatus |
| 818 | Bacteroides thetaiotaomicron |
| 226186 | Bacteroides thetaiotaomicron VPI-5482 |
| 657309 | Bacteroides xylanisolvens XB1A |
| 28119 | Bacteroides zoogleoformans |
| 1690483 | Bacteroidetes bacterium UKL13-3 |
| 310752 | Balaenoptera acutorostrata scammoni |
| 880074 | Barnesiella viscericola DSM 18177 |
| 1318743 | Bartonella ancashensis |
| 1686310 | Bartonella apis |
| 1094489 | Bartonella australis AUST/NH1 |
| 360095 | Bartonella bacilliformis KC583 |
| 696125 | Bartonella clarridgeiae 73 |
| 634504 | Bartonella grahamii as4aup |
| 38323 | Bartonella henselae |
| 38323 | Bartonella henselae |
| 283166 | Bartonella henselae str. Houston-1 |
| 1225179 | Bartonella quintana RM-11 |
| 283165 | Bartonella quintana str. Toulouse |
| 515256 | Bartonella sp. 1-1C |
| 1933910 | Bartonella sp. A1379B |
| 1933906 | Bartonella sp. JB15 |

| NCBI Taxon Identifier | Organism Name |
|---|---|
| 1933907 | Bartonella sp. JB63 |
| 1933912 | Bartonella sp. Raccoon60 |
| 1933904 | Bartonella sp. WD16.2 |
| 85701 | Bartonella tribocorum |
| 382640 | Bartonella tribocorum CIP 105476 |
| 1094497 | Bartonella vinsonii subsp. berkhoffii str. Winnie |
| 1072685 | Basilea psittacipulmonis DSM 24701 |
| 41875 | Bathycoccus prasinos |
| 717646 | Baudoinia panamericana UAMH 10762 |
| 374463 | Baumannia cicadellinicola str. Hc (Homalodisca coagulata) |
| 959 | Bdellovibrio bacteriovorus |
| 264462 | Bdellovibrio bacteriovorus HD100 |
| 1069642 | Bdellovibrio bacteriovorus str. Tiberius |
| 765869 | Bdellovibrio bacteriovorus W |
| 1916293 | Bdellovibrio sp. qaytius |
| 288004 | Beggiatoa leptomitoformis |
| 395963 | Beijerinckia indica subsp. indica ATCC 9039 |
| 866536 | Belliella baltica DSM 15883 |
| 1618337 | Berkelbacteria bacterium GW2011_GWE1_39_12 |
| 880071 | Bernardetia litoralis DSM 6794 |
| 1531429 | Berryella intestinalis |
| 543913 | beta proteobacterium CB |
| 3555 | Beta vulgaris subsp. vulgaris |
| 1904640 | Betaproteobacteria bacterium GR16-43 |
| 1690485 | Betaproteobacteria bacterium UKL13-2 |
| 471853 | Beutenbergia cavernae DSM 12333 |
| 1263829 | Bibersteinia trehalosi USDA-ARS-USMARC-188 |
| 1263831 | Bibersteinia trehalosi USDA-ARS-USMARC-189 |
| 1263832 | Bibersteinia trehalosi USDA-ARS-USMARC-190 |
| 1171377 | Bibersteinia trehalosi USDA-ARS-USMARC-192 |
| 1341694 | Bifidobacterium [indicum] DSM 20214 = LMG 11587 |
| 1437605 | Bifidobacterium actinocoloniiforme DSM 22766 |
| 1680 | Bifidobacterium adolescentis |
| 1680 | Bifidobacterium adolescentis |
| 367928 | Bifidobacterium adolescentis ATCC 15703 |
| 518635 | Bifidobacterium angulatum DSM 20098 = JCM 7096 |
| 28025 | Bifidobacterium animalis |
| 703613 | Bifidobacterium animalis subsp. animalis ATCC 25527 |
| 442563 | Bifidobacterium animalis subsp. lactis AD011 |
| 1167629 | Bifidobacterium animalis subsp. lactis ATCC 27673 |
| 1168290 | Bifidobacterium animalis subsp. lactis B420 |

| NCBI Taxon Identifier | Organism Name |
|---|---|
| 552531 | Bifidobacterium animalis subsp. lactis BB-12 |
| 742729 | Bifidobacterium animalis subsp. lactis Bi-07 |
| 580050 | Bifidobacterium animalis subsp. lactis Bl-04 |
| 1281781 | Bifidobacterium animalis subsp. lactis Bl12 |
| 1075106 | Bifidobacterium animalis subsp. lactis BLC1 |
| 1042403 | Bifidobacterium animalis subsp. lactis CNCM I-2494 |
| 555970 | Bifidobacterium animalis subsp. lactis DSM 10140 |
| 573236 | Bifidobacterium animalis subsp. lactis V9 |
| 1147128 | Bifidobacterium asteroides PRL2011 |
| 484020 | Bifidobacterium bifidum BGN4 |
| 702459 | Bifidobacterium bifidum PRL2010 |
| 883062 | Bifidobacterium bifidum S17 |
| 1385938 | Bifidobacterium breve 12L |
| 1385942 | Bifidobacterium breve 689b |
| 866777 | Bifidobacterium breve ACS-071-V-Sch8b |
| 518634 | Bifidobacterium breve DSM 20213 = JCM 1192 |
| 1385939 | Bifidobacterium breve JCM 7017 |
| 1385940 | Bifidobacterium breve JCM 7019 |
| 1385941 | Bifidobacterium breve NCFB 2258 |
| 936351 | Bifidobacterium breve S27 |
| 326426 | Bifidobacterium breve UCC2003 |
| 566552 | Bifidobacterium catenulatum DSM 16992 = JCM 1194 = LMG 11043 |
| 1447716 | Bifidobacterium catenulatum PV20-2 |
| 1150460 | Bifidobacterium catenulatum subsp. kashiwanohense JCM 15439 = DSM 21854 |
| 35760 | Bifidobacterium choerinum |
| 1687 | Bifidobacterium coryneforme |
| 401473 | Bifidobacterium dentium Bd1 |
| 1150423 | Bifidobacterium dentium JCM 1195 = DSM 20436 |
| 216816 | Bifidobacterium longum |
| 205913 | Bifidobacterium longum DJO10A |
| 206672 | Bifidobacterium longum NCC2705 |
| 565040 | Bifidobacterium longum subsp. infantis 157F |
| 391904 | Bifidobacterium longum subsp. infantis ATCC 15697 = JCM 1222 = DSM 20088 |
| 391904 | Bifidobacterium longum subsp. infantis ATCC 15697 = JCM 1222 = DSM 20088 |
| 890402 | Bifidobacterium longum subsp. longum BBMN68 |
| 722911 | Bifidobacterium longum subsp. longum F8 |
| 1300227 | Bifidobacterium longum subsp. longum GT15 |
| 565042 | Bifidobacterium longum subsp. longum JCM 1217 |
| 759350 | Bifidobacterium longum subsp. longum JDM301 |
| 1035817 | Bifidobacterium longum subsp. longum KACC 91563 |
| 547043 | Bifidobacterium pseudocatenulatum DSM 20438 = JCM 1200 = LMG 10505 |

| NCBI Taxon Identifier | Organism Name |
|---|---|
| 1447715 | Bifidobacterium pseudolongum PV8-2 |
| 1150461 | Bifidobacterium scardovii JCM 12489 = DSM 13734 |
| 1254439 | Bifidobacterium thermophilum RBL67 |
| 930090 | Bipolaris oryzae ATCC 44560 |
| 665912 | Bipolaris sorokiniana ND90Pr |
| 930089 | Bipolaris zeicola 26-R-13 |
| 2233851 | Blastochloris tepida |
| 1079 | Blastochloris viridis |
| 1146883 | Blastococcus saxobsidens DD2 |
| 1550728 | Blastomonas fulva |
| 1842535 | Blastomonas sp. RAC04 |
| 1229512 | Blattabacterium cuenoti BPAA |
| 1186051 | Blattabacterium sp. (Blaberus giganteus) |
| 1072467 | Blattabacterium sp. (Blatta orientalis) str. Tarazona |
| 331104 | Blattabacterium sp. (Blattella germanica) str. Bge |
| 298656 | Blattabacterium sp. (Cryptocercus kyebangensis) |
| 1075399 | Blattabacterium sp. (Cryptocercus punctulatus) str. Cpu |
| 1074889 | Blattabacterium sp. (Mastotermes darwiniensis) str. MADAR |
| 1316444 | Blattabacterium sp. (Nauphoeta cinerea) |
| 600809 | Blattabacterium sp. (Periplaneta americana) str. BPLAN |
| 1912897 | Blautia argi |
| 537007 | Blautia hansenii DSM 20583 |
| 657314 | Blautia obeum A2-162 |
| 33035 | Blautia producta |
| 1796616 | Blautia pseudococcoides |
| 1505597 | Blochmannia endosymbiont of Camponotus (Colobopsis) obliquus |
| 1505596 | Blochmannia endosymbiont of Polyrhachis (Hedomyrma) turneri |
| 150288 | Boleophthalmus pectinirostris |
| 132113 | Bombus impatiens |
| 30195 | Bombus terrestris |
| 7091 | Bombyx mori |
| 360910 | Bordetella avium 197N |
| 463025 | Bordetella bronchialis |
| 518 | Bordetella bronchiseptica |
| 568707 | Bordetella bronchiseptica 253 |
| 1208658 | Bordetella bronchiseptica MO149 |
| 257310 | Bordetella bronchiseptica RB50 |
| 463014 | Bordetella flabilis |
| 463040 | Bordetella genomosp. 13 |
| 103855 | Bordetella hinzii |
| 1172205 | Bordetella holmesii 44057 |

| NCBI Taxon Identifier | Organism Name |
|---|---|
| 1247649 | Bordetella holmesii ATCC 51541 |
| 257311 | Bordetella parapertussis 12822 |
| 1208660 | Bordetella parapertussis Bpp5 |
| 1403053 | Bordetella pertussis 137 |
| 568706 | Bordetella pertussis 18323 |
| 743277 | Bordetella pertussis B1917 |
| 1017264 | Bordetella pertussis CS |
| 257313 | Bordetella pertussis Tohama I |
| 340100 | Bordetella petrii DSM 12804 |
| 1331258 | Bordetella pseudohinzii |
| 1697043 | Bordetella sp. H567 |
| 1977852 | Bordetella sp. J329 |
| 123899 | Bordetella trematum |
| 1365188 | Borrelia anserina Es |
| 1155096 | Borrelia crocidurae str. Achema |
| 412419 | Borrelia duttonii Ly |
| 1329908 | Borrelia hermsii CC1 |
| 314723 | Borrelia hermsii DAH |
| 47466 | Borrelia miyamotoi |
| 1302858 | Borrelia miyamotoi LB-2001 |
| 1435045 | Borrelia parkeri HR1 |
| 412418 | Borrelia recurrentis A1 |
| 1104446 | Borrelia turcica IST7 |
| 314724 | Borrelia turicatae 91E135 |
| 1239934 | Borreliella afzelii HLJ01 |
| 411555 | Borreliella afzelii K78 |
| 390236 | Borreliella afzelii PKo |
| 390236 | Borreliella afzelii PKo |
| 1398490 | Borreliella afzelii Tom3107 |
| 290434 | Borreliella bavariensis PBi |
| 521010 | Borreliella bissettiae DN127 |
| 224326 | Borreliella burgdorferi B31 |
| 1328311 | Borreliella burgdorferi CA382 |
| 521008 | Borreliella burgdorferi JD1 |
| 521007 | Borreliella burgdorferi N40 |
| 445985 | Borreliella burgdorferi ZS7 |
| 1245910 | Borreliella chilensis |
| 29519 | Borreliella garinii |
| 1081646 | Borreliella garinii BgVir |
| 1234596 | Borreliella garinii NMJW1 |
| 1421551 | Borreliella garinii SZ |

| NCBI Taxon Identifier | Organism Name |
|---|---|
| 1674146 | Borreliella mayonii |
| 1398491 | Borreliella valaisiana Tom4006 |
| 9915 | Bos indicus |
| 72004 | Bos mutus |
| 9913 | Bos taurus |
| 1792307 | Bosea sp. PAMC 26642 |
| 1842539 | Bosea sp. RAC05 |
| 1867715 | Bosea sp. Tri-49 |
| 1526658 | Bosea vaviloviae |
| 332648 | Botrytis cinerea B05.10 |
| 1912795 | Boudabousia tangfeifanii |
| 2017485 | Brachybacterium avium |
| 446465 | Brachybacterium faecium DSM 4810 |
| 1331682 | Brachybacterium ginsengisoli |
| 556288 | Brachybacterium saurashtrense |
| 1903186 | Brachybacterium sp. P6-10-X1 |
| 2017484 | Brachybacterium vulturis |
| 15368 | Brachypodium distachyon |
| 1287055 | Brachyspira hampsonii |
| 1266923 | Brachyspira hyodysenteriae ATCC 27164 |
| 565034 | Brachyspira hyodysenteriae WA1 |
| 1045858 | Brachyspira intermedia PWS/A |
| 526224 | Brachyspira murdochii DSM 12563 |
| 759914 | Brachyspira pilosicoli 95/1000 |
| 1133568 | Brachyspira pilosicoli B2904 |
| 1042417 | Brachyspira pilosicoli P43/6/78 |
| 1161918 | Brachyspira pilosicoli WesB |
| 1548548 | Bradymonas sediminis |
| 1404768 | Bradyrhizobium amphicarpaeae |
| 1404864 | Bradyrhizobium cosmicum |
| 224911 | Bradyrhizobium diazoefficiens USDA 110 |
| 1274631 | Bradyrhizobium icense |
| 375 | Bradyrhizobium japonicum |
| 1037409 | Bradyrhizobium japonicum USDA 6 |
| 1245469 | Bradyrhizobium oligotrophicum S58 |
| 931866 | Bradyrhizobium ottawaense |
| 376 | Bradyrhizobium sp. |
| 288000 | Bradyrhizobium sp. BTAi1 |
| 1223566 | Bradyrhizobium sp. CCGE-LA001 |
| 114615 | Bradyrhizobium sp. ORS 278 |
| 115808 | Bradyrhizobium sp. ORS 285 |

| NCBI Taxon Identifier | Organism Name |
|---|---|
| 2057741 | Bradyrhizobium sp. SK17 |
| 7739 | Branchiostoma floridae |
| 3708 | Brassica napus |
| 109376 | Brassica oleracea var. oleracea |
| 3711 | Brassica rapa |
| 1109412 | Brenneria goodwinii |
| 2304600 | Breoghania sp. L-A4 |
| 1986204 | Brevefilum fermentans |
| 51101 | Brevibacillus agri |
| 358681 | Brevibacillus brevis NBRC 100599 |
| 54913 | Brevibacillus formosus |
| 1042163 | Brevibacillus laterosporus LMG 15441 |
| 273384 | Brevibacterium aurantiacum |
| 1703 | Brevibacterium linens |
| 1280413 | Breviolum minutum Mf 1.05b.01 |
| 293 | Brevundimonas diminuta |
| 588932 | Brevundimonas naejangsanensis |
| 1532555 | Brevundimonas sp. DS20 |
| 1827469 | Brevundimonas sp. GW460-12-10-14-LB2 |
| 1938605 | Brevundimonas sp. LM2 |
| 633149 | Brevundimonas subvibrioides ATCC 15264 |
| 41276 | Brevundimonas vesicularis |
| 2756 | Brochothrix thermosphacta |
| 235 | Brucella abortus |
| 235 | Brucella abortus |
| 235 | Brucella abortus |
| 235 | Brucella abortus |
| 359391 | Brucella abortus 2308 |
| 1104320 | Brucella abortus A13334 |
| 262698 | Brucella abortus bv. 1 str. 9-941 |
| 520450 | Brucella abortus bv. 2 str. 86/8/59 |
| 520454 | Brucella abortus bv. 6 str. 870 |
| 520455 | Brucella abortus bv. 9 str. C68 |
| 430066 | Brucella abortus S19 |
| 529 | Brucella anthropi |
| 439375 | Brucella anthropi ATCC 49188 |
| 36855 | Brucella canis |
| 36855 | Brucella canis |
| 483179 | Brucella canis ATCC 23365 |
| 1104321 | Brucella canis HSK A52141 |
| 1408887 | Brucella canis str. Oliveri |

| NCBI Taxon Identifier | Organism Name |
|---|---|
| 1423891 | Brucella ceti TE10759-12 |
| 1407053 | Brucella ceti TE28753-12 |
| 546272 | Brucella melitensis ATCC 23457 |
| 224914 | Brucella melitensis bv. 1 str. 16M |
| 224914 | Brucella melitensis bv. 1 str. 16M |
| 520466 | Brucella melitensis bv. 3 str. Ether |
| 941967 | Brucella melitensis M28 |
| 703352 | Brucella melitensis M5-90 |
| 1029825 | Brucella melitensis NI |
| 568815 | Brucella microti CCM 4915 |
| 444178 | Brucella ovis ATCC 25840 |
| 120576 | Brucella pinnipedialis |
| 520461 | Brucella pinnipedialis B2/94 |
| 419475 | Brucella pseudogrignonensis |
| 1891098 | Brucella sp. 2002734562 |
| 29461 | Brucella suis |
| 29461 | Brucella suis |
| 204722 | Brucella suis 1330 |
| 204722 | Brucella suis 1330 |
| 470137 | Brucella suis ATCC 23445 |
| 1004954 | Brucella suis bv. 1 str. S2 |
| 645170 | Brucella suis bv. 2 |
| 645170 | Brucella suis bv. 2 |
| 645170 | Brucella suis bv. 2 |
| 645170 | Brucella suis bv. 2 |
| 520487 | Brucella suis bv. 3 str. 686 |
| 1112912 | Brucella suis VBI22 |
| 981386 | Brucella vulpis |
| 6279 | Brugia malayi |
| 1265350 | Buchnera aphidicola (Aphis glycines) |
| 261317 | Buchnera aphidicola (Cinara tujafilina) |
| 372461 | Buchnera aphidicola BCc |
| 563178 | Buchnera aphidicola str. 5A (Acyrthosiphon pisum) |
| 1005090 | Buchnera aphidicola str. Ak (Acyrthosiphon kondoi) |
| 107806 | Buchnera aphidicola str. APS (Acyrthosiphon pisum) |
| 224915 | Buchnera aphidicola str. Bp (Baizongia pistaciae) |
| 1009859 | Buchnera aphidicola str. F009 (Myzus persicae) |
| 1009858 | Buchnera aphidicola str. G002 (Myzus persicae) |
| 713600 | Buchnera aphidicola str. JF98 (Acyrthosiphon pisum) |
| 713601 | Buchnera aphidicola str. JF99 (Acyrthosiphon pisum) |
| 713603 | Buchnera aphidicola str. LL01 (Acyrthosiphon pisum) |

| NCBI Taxon Identifier | Organism Name |
|---|---|
| 198804 | Buchnera aphidicola str. Sg (Schizaphis graminum) |
| 713602 | Buchnera aphidicola str. TLW03 (Acyrthosiphon pisum) |
| 561501 | Buchnera aphidicola str. Tuc7 (Acyrthosiphon pisum) |
| 1005057 | Buchnera aphidicola str. Ua (Uroleucon ambrosiae) |
| 1009856 | Buchnera aphidicola str. USDA (Myzus persicae) |
| 1009857 | Buchnera aphidicola str. W106 (Myzus persicae) |
| 339670 | Burkholderia ambifaria AMMD |
| 398577 | Burkholderia ambifaria MC40-6 |
| 95486 | Burkholderia cenocepacia |
| 95486 | Burkholderia cenocepacia |
| 1055524 | Burkholderia cenocepacia H111 |
| 331272 | Burkholderia cenocepacia HI2424 |
| 216591 | Burkholderia cenocepacia J2315 |
| 292 | Burkholderia cepacia |
| 983594 | Burkholderia cepacia ATCC 25416 |
| 1009846 | Burkholderia cepacia GG4 |
| 488447 | Burkholderia contaminans |
| 488732 | Burkholderia diffusa |
| 350701 | Burkholderia dolosa AU0158 |
| 28095 | Burkholderia gladioli |
| 999541 | Burkholderia gladioli BSR3 |
| 626418 | Burkholderia glumae BGR1 |
| 1176492 | Burkholderia glumae LMG 2196 = ATCC 33617 |
| 430531 | Burkholderia humptydooensis |
| 482957 | Burkholderia lata |
| 488446 | Burkholderia latens |
| 13373 | Burkholderia mallei |
| 13373 | Burkholderia mallei |
| 13373 | Burkholderia mallei |
| 13373 | Burkholderia mallei |
| 13373 | Burkholderia mallei |
| 13373 | Burkholderia mallei |
| 243160 | Burkholderia mallei ATCC 23344 |
| 412022 | Burkholderia mallei NCTC 10229 |
| 320389 | Burkholderia mallei NCTC 10247 |
| 320389 | Burkholderia mallei NCTC 10247 |
| 320388 | Burkholderia mallei SAVP1 |
| 488729 | Burkholderia metallica |
| 87883 | Burkholderia multivorans |
| 395019 | Burkholderia multivorans ATCC 17616 |
| 395019 | Burkholderia multivorans ATCC 17616 |

| NCBI Taxon Identifier | Organism Name |
|---|---|
| 985079 | Burkholderia multivorans ATCC BAA-247 |
| 342113 | Burkholderia oklahomensis |
| 441162 | Burkholderia oklahomensis C6786 |
| 331271 | Burkholderia orbicola AU 1054 |
| 406425 | Burkholderia orbicola MC0-3 |
| 41899 | Burkholderia plantarii |
| 41899 | Burkholderia plantarii |
| 28450 | Burkholderia pseudomallei |
| 884204 | Burkholderia pseudomallei 1026b |
| 357348 | Burkholderia pseudomallei 1106a |
| 320372 | Burkholderia pseudomallei 1710b |
| 320373 | Burkholderia pseudomallei 668 |
| 1439855 | Burkholderia pseudomallei A79A |
| 1229785 | Burkholderia pseudomallei BPC006 |
| 1306418 | Burkholderia pseudomallei HBPUB10134a |
| 272560 | Burkholderia pseudomallei K96243 |
| 1249471 | Burkholderia pseudomallei MSHR146 |
| 1335307 | Burkholderia pseudomallei MSHR305 |
| 536230 | Burkholderia pseudomallei MSHR346 |
| 1249474 | Burkholderia pseudomallei MSHR511 |
| 1249659 | Burkholderia pseudomallei MSHR520 |
| 1249477 | Burkholderia pseudomallei NAU35A-3 |
| 1241583 | Burkholderia pseudomallei NCTC 13179 |
| 1207504 | Burkholderia pseudomultivorans |
| 60550 | Burkholderia pyrrocinia |
| 488731 | Burkholderia seminalis |
| 1468409 | Burkholderia sp. 2002721687 |
| 640510 | Burkholderia sp. CCGE1001 |
| 640512 | Burkholderia sp. CCGE1003 |
| 1678678 | Burkholderia sp. HB1 |
| 416344 | Burkholderia sp. KJ006 |
| 1795043 | Burkholderia sp. PAMC 26561 |
| 1795874 | Burkholderia sp. PAMC 28687 |
| 758796 | Burkholderia sp. RPE67 |
| 1097668 | Burkholderia sp. YI23 |
| 95485 | Burkholderia stabilis |
| 1503054 | Burkholderia stagnalis |
| 1503055 | Burkholderia territorii |
| 57975 | Burkholderia thailandensis |
| 1249660 | Burkholderia thailandensis 2002721643 |
| 1241582 | Burkholderia thailandensis 2002721723 |

| NCBI Taxon Identifier | Organism Name |
|---|---|
| 1249665 | Burkholderia thailandensis E254 |
| 271848 | Burkholderia thailandensis E264 |
| 1249662 | Burkholderia thailandensis E444 |
| 1249663 | Burkholderia thailandensis H0587 |
| 1249661 | Burkholderia thailandensis MSMB121 |
| 1250339 | Burkholderia thailandensis MSMB59 |
| 1249667 | Burkholderia thailandensis USAMRU Malaysia #20 |
| 1249668 | Burkholderia ubonensis MSMB22 |
| 269482 | Burkholderia vietnamiensis G4 |
| 1449978 | Burkholderia vietnamiensis LMG 10929 |
| 1469502 | Burkholderiales bacterium GJ-E10 |
| 1834205 | Burkholderiales bacterium YL45 |
| 2479367 | Buttiauxella sp. 3AFRM03 |
| 245012 | butyrate-producing bacterium SM4/1 |
| 245014 | butyrate-producing bacterium SS3/4 |
| 2093856 | Butyricimonas faecalis |
| 657324 | Butyrivibrio fibrisolvens 16/4 |
| 185008 | Butyrivibrio hungatei |
| 515622 | Butyrivibrio proteoclasticus B316 |
| 758793 | Caballeronia insecticola |
| 1219035 | Caenibius tardaugens NBRC 16725 |
| 6238 | Caenorhabditis briggsae |
| 6239 | Caenorhabditis elegans |
| 3821 | Cajanus cajan |
| 273068 | Caldanaerobacter subterraneus subsp. tengcongensis MB4 |
| 632516 | Caldicellulosiruptor acetigenus 6A |
| 632335 | Caldicellulosiruptor acetigenus I77R1B |
| 521460 | Caldicellulosiruptor bescii DSM 6725 |
| 632292 | Caldicellulosiruptor hydrothermalis 108 |
| 632348 | Caldicellulosiruptor kronotskyensis 2002 |
| 608506 | Caldicellulosiruptor obsidiansis OB47 |
| 632518 | Caldicellulosiruptor owensensis OL |
| 351627 | Caldicellulosiruptor saccharolyticus DSM 8903 |
| 926550 | Caldilinea aerophila DSM 14535 = NBRC 104270 |
| 1653476 | Caldimicrobium thiodismutans |
| 413882 | Caldimonas brevitalea |
| 511051 | Caldisericum exile AZM16c01 |
| 1056495 | Caldisphaera lagunensis DSM 15908 |
| 768670 | Calditerrivibrio nitroreducens DSM 19672 |
| 880073 | Caldithrix abyssi DSM 13497 |
| 397948 | Caldivirga maquilingensis IC-167 |

| NCBI Taxon Identifier | Organism Name |
|---|---|
| 9483 | Callithrix jacchus |
| 7868 | Callorhinchus milii |
| 1337936 | Calothrix sp. 336/3 |
| 1170562 | Calothrix sp. PCC 6303 |
| 99598 | Calothrix sp. PCC 7507 |
| 90675 | Camelina sativa |
| 9838 | Camelus dromedarius |
| 419612 | Camelus ferus |
| 104421 | Camponotus floridanus |
| 522484 | Campylobacter avium LMG 24591 |
| 195 | Campylobacter coli |
| 195 | Campylobacter coli |
| 1358410 | Campylobacter coli 15-537360 |
| 1367491 | Campylobacter coli 76339 |
| 1273173 | Campylobacter coli CVM N29710 |
| 1183378 | Campylobacter coli RM1875 |
| 1183379 | Campylobacter coli RM4661 |
| 1183380 | Campylobacter coli RM5611 |
| 199 | Campylobacter concisus |
| 360104 | Campylobacter concisus 13826 |
| 1121267 | Campylobacter cuniculorum DSM 23162 = LMG 24588 |
| 360105 | Campylobacter curvus 525.92 |
| 1240980 | Campylobacter fetus subsp. fetus 04/554 |
| 360106 | Campylobacter fetus subsp. fetus 82-40 |
| 1507806 | Campylobacter fetus subsp. testudinum |
| 1244528 | Campylobacter fetus subsp. testudinum 03-427 |
| 1093099 | Campylobacter fetus subsp. venerealis 97/608 |
| 1273266 | Campylobacter fetus subsp. venerealis cfvi03/293 |
| 688354 | Campylobacter fetus subsp. venerealis str. 84-112 |
| 824 | Campylobacter gracilis |
| 28898 | Campylobacter helveticus |
| 1813019 | Campylobacter hepaticus |
| 360107 | Campylobacter hominis ATCC BAA-381 |
| 1031746 | Campylobacter hyointestinalis subsp. hyointestinalis LMG 9260 |
| 1244531 | Campylobacter iguaniorum |
| 1031564 | Campylobacter insulaenigrae NCTC 12927 |
| 1338035 | Campylobacter jejuni 32488 |
| 1347340 | Campylobacter jejuni 4031 |
| 195099 | Campylobacter jejuni RM1221 |
| 360109 | Campylobacter jejuni subsp. doylei 269.97 |
| 32022 | Campylobacter jejuni subsp. jejuni |

| NCBI Taxon Identifier | Organism Name |
|---|---|
| 32022 | Campylobacter jejuni subsp. jejuni |
| 32022 | Campylobacter jejuni subsp. jejuni |
| 32022 | Campylobacter jejuni subsp. jejuni |
| 32022 | Campylobacter jejuni subsp. jejuni |
| 1357994 | Campylobacter jejuni subsp. jejuni 00-2425 |
| 1380767 | Campylobacter jejuni subsp. jejuni 00-2426 |
| 1380768 | Campylobacter jejuni subsp. jejuni 00-2538 |
| 1383068 | Campylobacter jejuni subsp. jejuni 00-2544 |
| 407148 | Campylobacter jejuni subsp. jejuni 81116 |
| 354242 | Campylobacter jejuni subsp. jejuni 81-176 |
| 567106 | Campylobacter jejuni subsp. jejuni IA3902 |
| 757425 | Campylobacter jejuni subsp. jejuni ICDCCJ07001 |
| 645464 | Campylobacter jejuni subsp. jejuni M1 |
| 192222 | Campylobacter jejuni subsp. jejuni NCTC 11168 = ATCC 700819 |
| 1211776 | Campylobacter jejuni subsp. jejuni NCTC 11168-BN148 |
| 1201032 | Campylobacter jejuni subsp. jejuni PT14 |
| 1295010 | Campylobacter jejuni subsp. jejuni R14 |
| 718271 | Campylobacter jejuni subsp. jejuni S3 |
| 1031753 | Campylobacter lanienae NCTC 13004 |
| 1388750 | Campylobacter lari CCUG 22395 |
| 1388749 | Campylobacter lari NCTC 11845 |
| 1453989 | Campylobacter lari RM16701 |
| 1453988 | Campylobacter lari RM16712 |
| 306263 | Campylobacter lari RM2100 |
| 1031536 | Campylobacter lari subsp. concheus LMG 11760 |
| 1388753 | Campylobacter peloridis LMG 23910 |
| 1874362 | Campylobacter pinnipediorum subsp. caledonicus |
| 1500960 | Campylobacter sp. RM16704 |
| 1031920 | Campylobacter sputorum bv. faecalis CCUG 20703 |
| 1388751 | Campylobacter subantarcticus LMG 24374 |
| 1388752 | Campylobacter subantarcticus LMG 24377 |
| 1032069 | Campylobacter ureolyticus RIGS 9880 |
| 1660076 | Campylobacter vicugnae |
| 580033 | Campylobacter volucris LMG 24379 |
| 237561 | Candida albicans SC5314 |
| 573826 | Candida dubliniensis CD36 |
| 1136231 | Candida orthopsilosis Co 90-125 |
| 294747 | Candida tropicalis MYA-3404 |
| 1620412 | candidate division Kazan bacterium GW2011_GWA1_50_15 |
| 1643353 | candidate division SR1 bacterium Aalborg_AAW-1 |
| 1394709 | candidate division SR1 bacterium RAAC1_SR1_1 |

| NCBI Taxon Identifier | Organism Name |
|---|---|
| 1619077 | candidate division TM6 bacterium GW2011_GWF2_28_16 |
| 1394710 | candidate division WWE3 bacterium RAAC2_WWE3_1 |
| 522306 | Candidatus Accumulibacter regalis |
| 452471 | Candidatus Amoebophilus asiaticus 5a2 |
| 634113 | Candidatus Arsenophonus lipoptenae |
| 1029718 | Candidatus Arthromitus sp. SFB-mouse-Japan |
| 1508644 | Candidatus Arthromitus sp. SFB-mouse-NL |
| 1041809 | Candidatus Arthromitus sp. SFB-mouse-Yit |
| 1041504 | Candidatus Arthromitus sp. SFB-rat-Yit |
| 1453429 | Candidatus Atelocyanobacterium thalassa isolate ALOHA |
| 511995 | Candidatus Azobacteroides pseudotrichonymphae genomovar. CFP2 |
| 673862 | Candidatus Babela massiliensis |
| 1700835 | Candidatus Bathyarchaeota archaeon BA1 |
| 1700836 | Candidatus Bathyarchaeota archaeon BA2 |
| 1618372 | Candidatus Beckwithbacteria bacterium GW2011_GWC1_49_16 |
| 2026885 | Candidatus Bipolaricaulis anaerobius |
| 2501609 | Candidatus Bipolaricaulis sibiricus |
| 1240471 | Candidatus Blochmanniella chromaiodes str. 640 |
| 203907 | Candidatus Blochmanniella floridana |
| 291272 | Candidatus Blochmanniella pennsylvanica str. BPEN |
| 859654 | Candidatus Blochmanniella vafra str. BVAF |
| 311458 | Candidatus Caldarchaeum subterraneum |
| 1618633 | Candidatus Campbellbacteria bacterium GW2011_OD1_34_28 |
| 247481 | Candidatus Cardinium hertigii |
| 1202536 | Candidatus Carsonella ruddii CE isolate Thao2000 |
| 1202537 | Candidatus Carsonella ruddii CS isolate Thao2000 |
| 667013 | Candidatus Carsonella ruddii DC |
| 1202538 | Candidatus Carsonella ruddii HC isolate Thao2000 |
| 1202539 | Candidatus Carsonella ruddii HT isolate Thao2000 |
| 1202540 | Candidatus Carsonella ruddii PC isolate NHV |
| 387662 | Candidatus Carsonella ruddii PV |
| 459349 | Candidatus Cloacimonas acidaminovorans str. Evry |
| 2054173 | Candidatus Coxiella mudrowiae |
| 477974 | Candidatus Desulforudis audaxviator MP104C |
| 1725232 | Candidatus Desulfovibrio trichonymphae |
| 1778262 | Candidatus Doolittlea endobia |
| 1193729 | Candidatus Endolissoclinum faulkneri L2 |
| 1401328 | Candidatus Endolissoclinum faulkneri L5 |
| 1408204 | Candidatus Endomicrobiellum trichonymphae |
| 1408204 | Candidatus Endomicrobiellum trichonymphae |
| 1608628 | Candidatus Filomicrobium marinum |

| NCBI Taxon Identifier | Organism Name |
|---|---|
| 1608628 | Candidatus Filomicrobium marinum |
| 1798018 | Candidatus Fluviicola riflensis |
| 1802984 | Candidatus Fokinia solitaria |
| 653937 | Candidatus Francisella endociliophora |
| 1070130 | Candidatus Gullanella endobia |
| 572265 | Candidatus Hamiltonella defensa 5AT (Acyrthosiphon pisum) |
| 1427984 | Candidatus Hepatoplasma crinochetorum Av |
| 1778263 | Candidatus Hoaglandella endobia |
| 573658 | Candidatus Hodgkinia cicadicola |
| 573658 | Candidatus Hodgkinia cicadicola |
| 573658 | Candidatus Hodgkinia cicadicola |
| 573234 | Candidatus Hodgkinia cicadicola Dsem |
| 476281 | Candidatus Ishikawaella capsulata Mpkobe |
| 1541959 | Candidatus Izimaplasma bacterium HR1 |
| 1495769 | Candidatus Johnevansia muelleri |
| 336810 | Candidatus Karelsulcia muelleri |
| 336810 | Candidatus Karelsulcia muelleri |
| 336810 | Candidatus Karelsulcia muelleri |
| 706194 | Candidatus Karelsulcia muelleri CARI |
| 641892 | Candidatus Karelsulcia muelleri DMIN |
| 444179 | Candidatus Karelsulcia muelleri GWSS |
| 1189303 | Candidatus Karelsulcia muelleri PSPU |
| 595499 | Candidatus Karelsulcia muelleri SMDSEM |
| 1343076 | Candidatus Karelsulcia muelleri str. Sulcia-ALF |
| 1208923 | Candidatus Kinetoplastibacterium blastocrithidii (ex Strigomonas culicis) |
| 1208922 | Candidatus Kinetoplastibacterium blastocrithidii TCC012E |
| 1267577 | Candidatus Kinetoplastibacterium crithidii (ex Angomonas deanei ATCC 30255) |
| 1208918 | Candidatus Kinetoplastibacterium crithidii TCC036E |
| 1208919 | Candidatus Kinetoplastibacterium desouzaii TCC079E |
| 1208921 | Candidatus Kinetoplastibacterium galatii TCC219 |
| 1208920 | Candidatus Kinetoplastibacterium oncopeltii TCC290E |
| 1576550 | Candidatus Kinetoplastibacterium sorsogonicusi |
| 374847 | Candidatus Korarchaeum cryptofilum OPF8 |
| 204669 | Candidatus Koribacter versatilis Ellin345 |
| 174633 | Candidatus Kuenenia stuttgartiensis |
| 2005262 | Candidatus Legionella polyplacis |
| 1277257 | Candidatus Liberibacter africanus PTSAPSY |
| 1261131 | Candidatus Liberibacter americanus str. Sao Paulo |
| 1174529 | Candidatus Liberibacter asiaticus str. gxpsy |
| 931202 | Candidatus Liberibacter asiaticus str. Ishi-1 |
| 537021 | Candidatus Liberibacter asiaticus str. psy62 |

| NCBI Taxon Identifier | Organism Name |
|---|---|
| 658172 | Candidatus Liberibacter solanacearum CLso-ZC1 |
| 1538547 | Candidatus Lokiarchaeum sp. GC14_75 |
| 1318617 | Candidatus Malacoplasma girerdii |
| 1920749 | Candidatus Mancarchaeum acidiphilum |
| 1899017 | Candidatus Melainabacteria bacterium MEL.A1 |
| 1295009 | Candidatus Methanomassiliicoccus intestinalis Issoire-Mx1 |
| 1236689 | Candidatus Methanomethylophilus alvi Mx1201 |
| 1577791 | Candidatus Methanoplasma termitum |
| 671143 | Candidatus Methylomirabilis oxygeniifera |
| 1581557 | Candidatus Methylopumilus planktonicus |
| 1581680 | Candidatus Methylopumilus turicensis |
| 696127 | Candidatus Midichloria mitochondrii IricVA |
| 1778264 | Candidatus Mikella endobia |
| 903503 | Candidatus Moranella endobia PCIT |
| 1234603 | Candidatus Moranella endobia PCVAL |
| 1212765 | Candidatus Mycoplasma haematolamae str. Purdue |
| 1116213 | Candidatus Mycoplasma haematominutum 'Birmingham 1' |
| 1884916 | Candidatus Nanopelagicus abundans |
| 1884915 | Candidatus Nanopelagicus hibericus |
| 1884634 | Candidatus Nanopelagicus limnes |
| 1577684 | Candidatus Nanopusillus acidilobi |
| 2093824 | Candidatus Nanosynbacter lyticus |
| 1343077 | Candidatus Nasuia deltocephalinicola str. NAS-ALF |
| 2058097 | Candidatus Nitrosocaldus cavascurensis |
| 1353260 | Candidatus Nitrosocosmicus oleophilus |
| 1630141 | Candidatus Nitrosoglobus terrae |
| 1898749 | Candidatus Nitrosomarinus catalina |
| 1410606 | Candidatus Nitrosopelagicus brevis |
| 1229908 | Candidatus Nitrosopumilus koreensis AR1 |
| 1229909 | Candidatus Nitrosopumilus sediminis |
| 1459636 | Candidatus Nitrososphaera evergladensis SR1 |
| 1237085 | Candidatus Nitrososphaera gargensis Ga9.2 |
| 1078905 | Candidatus Nitrosotalea devanaterra |
| 1603555 | Candidatus Nitrosotenuis cloacae |
| 1715989 | Candidatus Nitrospira inopinata |
| 1414854 | Candidatus Nucleicultrix amoebiphila FS5 |
| 186490 | Candidatus Palibaumannia cicadellinicola |
| 186490 | Candidatus Palibaumannia cicadellinicola |
| 1235990 | Candidatus Pantoea carbekii |
| 1235990 | Candidatus Pantoea carbekii |
| 91604 | Candidatus Paracaedibacter acanthamoebae |

| NCBI Taxon Identifier | Organism Name |
|---|---|
| 244581 | Candidatus Paracaedimonas acanthamoebae |
| 1002672 | Candidatus Pelagibacter sp. IMCC9063 |
| 335992 | Candidatus Pelagibacter ubique HTCC1062 |
| 1735162 | Candidatus Peribacter riflensis |
| 2115978 | Candidatus Phycorickettsia trachydisci |
| 59748 | Candidatus Phytoplasma australiense |
| 37692 | Candidatus Phytoplasma mali |
| 69896 | Candidatus Phytoplasma solani |
| 135727 | Candidatus Phytoplasma ziziphi |
| 1884914 | Candidatus Planktophila dulcis |
| 1884913 | Candidatus Planktophila lacus |
| 573600 | Candidatus Planktophila limnetica |
| 1884904 | Candidatus Planktophila sulfonica |
| 1884907 | Candidatus Planktophila vernalis |
| 1884905 | Candidatus Planktophila versatilis |
| 91844 | Candidatus Portiera aleyrodidarum |
| 91844 | Candidatus Portiera aleyrodidarum |
| 91844 | Candidatus Portiera aleyrodidarum |
| 1206109 | Candidatus Portiera aleyrodidarum BT-B-HRs |
| 1206109 | Candidatus Portiera aleyrodidarum BT-B-HRs |
| 1239881 | Candidatus Portiera aleyrodidarum BT-QVLC |
| 1239881 | Candidatus Portiera aleyrodidarum BT-QVLC |
| 1163752 | Candidatus Portiera aleyrodidarum MED (Bemisia tabaci) |
| 1297582 | Candidatus Portiera aleyrodidarum TV |
| 669502 | Candidatus Profftella armatura |
| 1806508 | Candidatus Promineifilum breve |
| 264201 | Candidatus Protochlamydia amoebophila UWE25 |
| 389348 | Candidatus Protochlamydia naegleriophila |
| 2716812 | Candidatus Protofrankia datiscae |
| 1302376 | Candidatus Pseudomonas adelgestsugas |
| 1125411 | Candidatus Pseudothioglobus singularis PS1 |
| 488538 | Candidatus Puniceispirillum marinum IMCC1322 |
| 472834 | Candidatus Purcelliella pentastirinorum |
| 535712 | Candidatus Rhodoluna planktonica |
| 676208 | Candidatus Rickettsiella viridis |
| 515618 | Candidatus Riesia pediculicola USDA |
| 1719125 | Candidatus Riesia sp. GBBU |
| 413404 | Candidatus Ruthia magnifica str. Cm (Calyptogena magnifica) |
| 1619070 | Candidatus Saccharibacteria bacterium GW2011_GWC2_44_17 |
| 1394711 | Candidatus Saccharibacteria bacterium RAAC3_TM7_1 |
| 1332188 | Candidatus Saccharimonas aalborgensis |

| NCBI Taxon Identifier | Organism Name |
|---|---|
| 2342 | Candidatus Sodalis pierantonius str. SOPE |
| 234267 | Candidatus Solibacter usitatus Ellin6076 |
| 1249480 | Candidatus Sulfuricurvum sp. RIFRC-1 |
| 946483 | Candidatus Symbiobacter mobilis CR |
| 1410383 | Candidatus Tachikawaea gelatinosa |
| 1748243 | Candidatus Tenderia electrophaga |
| 1166950 | Candidatus Thiodictyon syntrophicum |
| 1705394 | Candidatus Thioglobus autotrophicus |
| 2508687 | Candidatus Thioglobus sp. NP1 |
| 1902579 | Candidatus Tokpelaia hoelldoblerii |
| 1266371 | Candidatus Tremblaya phenacola PAVE |
| 891398 | Candidatus Tremblaya princeps PCIT |
| 1053648 | Candidatus Tremblaya princeps PCVAL |
| 1133592 | Candidatus Uzinura diaspidicola str. ASNER |
| 1930593 | Candidatus Velamenicoccus archaeovorus |
| 412965 | Candidatus Vesicomyosocius okutanii |
| 1759059 | Candidatus Viadribacter manganicus |
| 881286 | Candidatus Vidania fulgoroideae |
| 1415657 | Candidatus Walczuchella monophlebidarum |
| 1618600 | Candidatus Woesebacteria bacterium GW2011_GWF1_31_35 |
| 1619007 | Candidatus Wolfebacteria bacterium GW2011_GWB1_47_1 |
| 1704307 | Candidatus Xiphinematobacter sp. Idaho Grape |
| 871271 | Candidatus Zinderia insecticola CARI |
| 498019 | Candidozyma auris |
| 9615 | Canis lupus familiaris |
| 860228 | Capnocytophaga canimorsus Cc5 |
| 28189 | Capnocytophaga cynodegmi |
| 1017 | Capnocytophaga gingivalis |
| 45243 | Capnocytophaga haemolytica |
| 327575 | Capnocytophaga leadbetteri |
| 521097 | Capnocytophaga ochracea DSM 7271 |
| 209053 | Capnocytophaga sp. ChDC OS43 |
| 1945657 | Capnocytophaga sp. H2931 |
| 1945658 | Capnocytophaga sp. H4358 |
| 1705617 | Capnocytophaga sp. oral taxon 323 |
| 1019 | Capnocytophaga sputigena |
| 1848904 | Capnocytophaga stomatis |
| 9925 | Capra hircus |
| 2507162 | Caproiciproducens sp. NJN-50 |
| 81985 | Capsella rubella |
| 4072 | Capsicum annuum |

| NCBI Taxon Identifier | Organism Name |
|---|---|
| 178899 | Carboxydocella thermautotrophica |
| 246194 | Carboxydothermus hydrogenoformans Z-2901 |
| 1231626 | Cardinium endosymbiont cEper1 of Encarsia pergandiella |
| 650378 | Cardinium endosymbiont of Sogatella furcifera |
| 2718 | Cardiobacterium hominis |
| 3649 | Carica papaya |
| 2748 | Carnobacterium divergens |
| 1266845 | Carnobacterium inhibens subsp. gilichinskyi |
| 1234679 | Carnobacterium maltaromaticum LMA28 |
| 208596 | Carnobacterium sp. 17-4 |
| 1564681 | Carnobacterium sp. CP1 |
| 1437824 | Castellaniella defragrans 65Phen |
| 51338 | Castor canadensis |
| 479433 | Catenulispora acidiphila DSM 44928 |
| 1679497 | Caulobacter flavus |
| 69395 | Caulobacter henricii |
| 69666 | Caulobacter mirabilis |
| 509190 | Caulobacter segnis ATCC 21756 |
| 366602 | Caulobacter sp. K31 |
| 190650 | Caulobacter vibrioides CB15 |
| 565050 | Caulobacter vibrioides NA1000 |
| 261658 | Cavenderia fasciculata |
| 158823 | Cedecea lapagei |
| 158822 | Cedecea neteri |
| 158822 | Cedecea neteri |
| 158822 | Cedecea neteri |
| 1758178 | Celeribacter ethanolicus |
| 1208324 | Celeribacter indicus |
| 1397108 | Celeribacter marinus |
| 590998 | Cellulomonas fimi ATCC 484 |
| 446466 | Cellulomonas flavigena DSM 20109 |
| 593907 | Cellulomonas gilvus ATCC 13127 |
| 2003551 | Cellulomonas sp. PSBB021 |
| 688270 | Cellulophaga algicola DSM 14237 |
| 1348584 | Cellulophaga baltica 18 |
| 1348585 | Cellulophaga baltica NN016038 |
| 979 | Cellulophaga lytica |
| 867900 | Cellulophaga lytica DSM 7489 |
| 642492 | Cellulosilyticum lentocellum DSM 5427 |
| 1710 | Cellulosimicrobium cellulans |
| 1980001 | Cellulosimicrobium sp. TH-20 |

| NCBI Taxon Identifier | Organism Name |
|---|---|
| 498211 | Cellvibrio japonicus Ueda107 |
| 1987723 | Cellvibrio sp. PSBB006 |
| 1945512 | Cellvibrio sp. PSBB023 |
| 414004 | Cenarchaeum symbiosum A |
| 272943 | Cereibacter sphaeroides 2.4.1 |
| 349102 | Cereibacter sphaeroides ATCC 17025 |
| 349101 | Cereibacter sphaeroides ATCC 17029 |
| 557760 | Cereibacter sphaeroides KD131 |
| 1173020 | Chamaesiphon minutus PCC 6605 |
| 1441930 | Chania multitudinisentens RB-25 |
| 266779 | Chelativorans sp. BNC1 |
| 444444 | Chelatococcus daeguensis |
| 1702325 | Chelatococcus sp. CO-6 |
| 8469 | Chelonia mydas |
| 63459 | Chenopodium quinoa |
| 2029983 | Chitinophaga caeni |
| 485918 | Chitinophaga pinensis DSM 2588 |
| 83555 | Chlamydia abortus |
| 218497 | Chlamydia abortus S26/3 |
| 1229831 | Chlamydia avium 10DC88 |
| 227941 | Chlamydia caviae GPIC |
| 264202 | Chlamydia felis Fe/C-56 |
| 1143323 | Chlamydia gallinacea 08-1274/3 |
| 83560 | Chlamydia muridarum |
| 83560 | Chlamydia muridarum |
| 83560 | Chlamydia muridarum |
| 243161 | Chlamydia muridarum str. Nigg |
| 1460373 | Chlamydia muridarum str. Nigg 2 MCR |
| 1434773 | Chlamydia muridarum str. Nigg CM972 |
| 1434770 | Chlamydia muridarum str. Nigg3 CMUT3-5 |
| 331635 | Chlamydia pecorum E58 |
| 1234369 | Chlamydia pecorum P787 |
| 1234367 | Chlamydia pecorum PV3056/3 |
| 1234368 | Chlamydia pecorum W73 |
| 115711 | Chlamydia pneumoniae AR39 |
| 115713 | Chlamydia pneumoniae CWL029 |
| 138677 | Chlamydia pneumoniae J138 |
| 406984 | Chlamydia pneumoniae LPCoLN |
| 182082 | Chlamydia pneumoniae TW-183 |
| 1967783 | Chlamydia poikilotherma |
| 1112252 | Chlamydia psittaci 01DC11 |

| NCBI Taxon Identifier | Organism Name |
|---|---|
| 1221877 | Chlamydia psittaci 01DC12 |
| 1112254 | Chlamydia psittaci 02DC15 |
| 1112267 | Chlamydia psittaci 08DC60 |
| 331636 | Chlamydia psittaci 6BC |
| 331636 | Chlamydia psittaci 6BC |
| 1218176 | Chlamydia psittaci 84/55 |
| 1112250 | Chlamydia psittaci C19/98 |
| 1050219 | Chlamydia psittaci CP3 |
| 1218353 | Chlamydia psittaci GR9 |
| 1218357 | Chlamydia psittaci M56 |
| 500464 | Chlamydia psittaci Mat116 |
| 1218354 | Chlamydia psittaci MN |
| 1050221 | Chlamydia psittaci NJ1 |
| 929557 | Chlamydia psittaci RD1 |
| 1218355 | Chlamydia psittaci VS225 |
| 1218358 | Chlamydia psittaci WC |
| 1218356 | Chlamydia psittaci WS/RT/E30 |
| 1071753 | Chlamydia trachomatis A/363 |
| 1071754 | Chlamydia trachomatis A/5291 |
| 1071755 | Chlamydia trachomatis A/7249 |
| 315277 | Chlamydia trachomatis A/HAR-13 |
| 580047 | Chlamydia trachomatis A2497 |
| 580047 | Chlamydia trachomatis A2497 |
| 580049 | Chlamydia trachomatis B/Jali20/OT |
| 672161 | Chlamydia trachomatis B/TZ1A828/OT |
| 1431547 | Chlamydia trachomatis C/TW-3 |
| 1071756 | Chlamydia trachomatis D/SotonD1 |
| 1071757 | Chlamydia trachomatis D/SotonD5 |
| 1071758 | Chlamydia trachomatis D/SotonD6 |
| 272561 | Chlamydia trachomatis D/UW-3/CX |
| 759363 | Chlamydia trachomatis D-EC |
| 759364 | Chlamydia trachomatis D-LC |
| 707183 | Chlamydia trachomatis E/11023 |
| 707184 | Chlamydia trachomatis E/150 |
| 596777 | Chlamydia trachomatis E/Bour |
| 1100832 | Chlamydia trachomatis E/C599 |
| 1071760 | Chlamydia trachomatis E/SotonE4 |
| 1071761 | Chlamydia trachomatis E/SotonE8 |
| 1071759 | Chlamydia trachomatis E/SW3 |
| 1340853 | Chlamydia trachomatis F/11-96 |
| 1071764 | Chlamydia trachomatis F/SotonF3 |

| NCBI Taxon Identifier | Organism Name |
|---|---|
| 1071762 | Chlamydia trachomatis F/SW4 |
| 1071763 | Chlamydia trachomatis F/SW5 |
| 1100833 | Chlamydia trachomatis F/SWFPminus |
| 707187 | Chlamydia trachomatis G/11074 |
| 707186 | Chlamydia trachomatis G/11222 |
| 718219 | Chlamydia trachomatis G/9301 |
| 707185 | Chlamydia trachomatis G/9768 |
| 1071765 | Chlamydia trachomatis G/SotonG1 |
| 1071766 | Chlamydia trachomatis Ia/SotonIa1 |
| 1071767 | Chlamydia trachomatis Ia/SotonIa3 |
| 1260222 | Chlamydia trachomatis IU824 |
| 1260223 | Chlamydia trachomatis IU888 |
| 907272 | Chlamydia trachomatis J/6276tet1 |
| 1071768 | Chlamydia trachomatis K/SotonK1 |
| 1071770 | Chlamydia trachomatis L1/115 |
| 1075087 | Chlamydia trachomatis L1/1322/p2 |
| 1071771 | Chlamydia trachomatis L1/224 |
| 1071769 | Chlamydia trachomatis L1/440/LN |
| 1071772 | Chlamydia trachomatis L2/25667R |
| 471472 | Chlamydia trachomatis L2/434/Bu |
| 1262673 | Chlamydia trachomatis L2/434/Bu(f) |
| 1263406 | Chlamydia trachomatis L2/434/Bu(i) |
| 1071777 | Chlamydia trachomatis L2b/795 |
| 1071773 | Chlamydia trachomatis L2b/8200/07 |
| 1071778 | Chlamydia trachomatis L2b/Ams1 |
| 1071779 | Chlamydia trachomatis L2b/Ams2 |
| 1071780 | Chlamydia trachomatis L2b/Ams3 |
| 1071781 | Chlamydia trachomatis L2b/Ams4 |
| 1071782 | Chlamydia trachomatis L2b/Ams5 |
| 1075085 | Chlamydia trachomatis L2b/Canada1 |
| 1075086 | Chlamydia trachomatis L2b/Canada2 |
| 1071776 | Chlamydia trachomatis L2b/CV204 |
| 1071775 | Chlamydia trachomatis L2b/LST |
| 471473 | Chlamydia trachomatis L2b/UCH-1/proctitis |
| 1071774 | Chlamydia trachomatis L2b/UCH-2 |
| 887712 | Chlamydia trachomatis L2c |
| 1071783 | Chlamydia trachomatis L3/404/LN |
| 907269 | Chlamydia trachomatis RC-F(s)/342 |
| 907265 | Chlamydia trachomatis RC-F(s)/852 |
| 907263 | Chlamydia trachomatis RC-F/69 |
| 907270 | Chlamydia trachomatis RC-J(s)/122 |

| NCBI Taxon Identifier | Organism Name |
|---|---|
| 907266 | Chlamydia trachomatis RC-J/943 |
| 907267 | Chlamydia trachomatis RC-J/953 |
| 907271 | Chlamydia trachomatis RC-J/966 |
| 1007871 | Chlamydia trachomatis RC-J/971 |
| 907268 | Chlamydia trachomatis RC-L2(s)/3 |
| 907264 | Chlamydia trachomatis RC-L2(s)/46 |
| 1007870 | Chlamydia trachomatis RC-L2/55 |
| 634464 | Chlamydia trachomatis Sweden2 |
| 3055 | Chlamydomonas reinhardtii |
| 981222 | Chloracidobacterium thermophilum B |
| 554065 | Chlorella variabilis |
| 274537 | Chlorobaculum limnaeum |
| 517417 | Chlorobaculum parvum NCIB 8327 |
| 194439 | Chlorobaculum tepidum TLS |
| 340177 | Chlorobium chlorochromatii CaD3 |
| 290315 | Chlorobium limicola DSM 245 |
| 331678 | Chlorobium phaeobacteroides BS1 |
| 290317 | Chlorobium phaeobacteroides DSM 266 |
| 290318 | Chlorobium phaeovibrioides DSM 265 |
| 60711 | Chlorocebus sabaeus |
| 326427 | Chloroflexus aggregans DSM 9485 |
| 324602 | Chloroflexus aurantiacus J-10-fl |
| 480224 | Chloroflexus aurantiacus Y-400-fl |
| 517418 | Chloroherpeton thalassium ATCC 35110 |
| 2005460 | Chondrocystis sp. NIES-4102 |
| 52 | Chondromyces crocatus |
| 2769 | Chondrus crispus |
| 626937 | Christensenella minuta |
| 1229726 | Christiangramia flava JLT2011 |
| 411154 | Christiangramia forsetii KT0803 |
| 2126553 | Christiangramia fulva |
| 1913577 | Christiangramia salexigens |
| 2202141 | Chromobacterium phragmitis |
| 2202141 | Chromobacterium phragmitis |
| 1778675 | Chromobacterium rhizoryzae |
| 2059672 | Chromobacterium sp. ATCC 53434 |
| 1108595 | Chromobacterium vaccinii |
| 243365 | Chromobacterium violaceum ATCC 12472 |
| 290398 | Chromohalobacter israelensis DSM 3043 |
| 251229 | Chroococcidiopsis thermalis PCC 7203 |
| 8478 | Chrysemys picta bellii |

| NCBI Taxon Identifier | Organism Name |
|---|---|
| 651561 | Chryseobacterium arthrosphaerae |
| 1324352 | Chryseobacterium gallinarum |
| 1685010 | Chryseobacterium glaciei |
| 253 | Chryseobacterium indologenes |
| 558152 | Chryseobacterium piperi |
| 2478663 | Chryseobacterium sp. 3008163 |
| 2039166 | Chryseobacterium sp. 6424 |
| 1721091 | Chryseobacterium sp. IHB B 17019 |
| 878220 | Chryseobacterium sp. StRB126 |
| 2015076 | Chryseobacterium sp. T16E-39 |
| 536441 | Chryseobacterium taklimakanense |
| 2321403 | Chryseolinea soli |
| 1303518 | Chthonomonas calidirosea T49 |
| 3827 | Cicer arietinum |
| 79782 | Cimex lectularius |
| 7719 | Ciona intestinalis |
| 404380 | Citrifermentans bemidjiense Bem |
| 35703 | Citrobacter amalonaticus |
| 1261127 | Citrobacter amalonaticus Y19 |
| 57706 | Citrobacter braakii |
| 67824 | Citrobacter farmeri |
| 1333848 | Citrobacter freundii CFNIH1 |
| 290338 | Citrobacter koseri ATCC BAA-895 |
| 1563222 | Citrobacter pasteurii |
| 637910 | Citrobacter rodentium ICC168 |
| 1920110 | Citrobacter sp. CFNIH10 |
| 1703250 | Citrobacter sp. CRE-46 |
| 1702170 | Citrobacter sp. FDAARGOS_156 |
| 67827 | Citrobacter werkmanii |
| 133448 | Citrobacter youngae |
| 1634516 | Citromicrobium sp. JL477 |
| 2711 | Citrus sinensis |
| 85681 | Citrus x clementina |
| 1874630 | Clavibacter capsici |
| 33014 | Clavibacter michiganensis subsp. insidiosus |
| 443906 | Clavibacter michiganensis subsp. michiganensis NCPPB 382 |
| 1097677 | Clavibacter nebraskensis NCPPB 2581 |
| 31964 | Clavibacter sepedonicus |
| 36911 | Clavispora lusitaniae |
| 306902 | Clavispora lusitaniae ATCC 42720 |
| 1197717 | Cloacibacillus porcorum |

| NCBI Taxon Identifier | Organism Name |
|---|---|
| 237258 | Cloacibacterium normanense |
| 1496 | Clostridioides difficile |
| 272563 | Clostridioides difficile 630 |
| 272563 | Clostridioides difficile 630 |
| 645462 | Clostridioides difficile CD196 |
| 645463 | Clostridioides difficile R20291 |
| 84022 | Clostridium aceticum |
| 272562 | Clostridium acetobutylicum ATCC 824 |
| 991791 | Clostridium acetobutylicum DSM 1731 |
| 863638 | Clostridium acetobutylicum EA 2018 |
| 29341 | Clostridium argentinense |
| 1341692 | Clostridium autoethanogenum DSM 10061 |
| 1415775 | Clostridium baratii str. Sullivan |
| 1520 | Clostridium beijerinckii |
| 864803 | Clostridium beijerinckii ATCC 35702 |
| 290402 | Clostridium beijerinckii NCIMB 8052 |
| 1216932 | Clostridium bornimense |
| 441770 | Clostridium botulinum A str. ATCC 19397 |
| 413999 | Clostridium botulinum A str. ATCC 3502 |
| 441771 | Clostridium botulinum A str. Hall |
| 536232 | Clostridium botulinum A2 str. Kyoto |
| 498214 | Clostridium botulinum A3 str. Loch Maree |
| 935198 | Clostridium botulinum B str. Eklund 17B (NRP) |
| 498213 | Clostridium botulinum B1 str. Okra |
| 515621 | Clostridium botulinum Ba4 str. 657 |
| 929506 | Clostridium botulinum BKT015925 |
| 508767 | Clostridium botulinum E3 str. Alaska E43 |
| 758678 | Clostridium botulinum F str. 230613 |
| 441772 | Clostridium botulinum F str. Langeland |
| 941968 | Clostridium botulinum H04402 065 |
| 1492 | Clostridium butyricum |
| 536227 | Clostridium carboxidivorans P7 |
| 573061 | Clostridium cellulovorans 743B |
| 46867 | Clostridium chauvoei |
| 332101 | Clostridium drakei |
| 1552 | Clostridium estertheticum subsp. estertheticum |
| 1497 | Clostridium formicaceticum |
| 182773 | Clostridium isatidis |
| 431943 | Clostridium kluyveri DSM 555 |
| 583346 | Clostridium kluyveri NBRC 12016 |
| 748727 | Clostridium ljungdahlii DSM 13528 |

| NCBI Taxon Identifier | Organism Name |
|---|---|
| 386415 | Clostridium novyi NT |
| 86416 | Clostridium pasteurianum BC1 |
| 1262449 | Clostridium pasteurianum DSM 525 = ATCC 6013 |
| 1262449 | Clostridium pasteurianum DSM 525 = ATCC 6013 |
| 195103 | Clostridium perfringens ATCC 13124 |
| 289380 | Clostridium perfringens SM101 |
| 195102 | Clostridium perfringens str. 13 |
| 1345695 | Clostridium saccharobutylicum DSM 13864 |
| 931276 | Clostridium saccharoperbutylacetonicum N1-4(HMT) |
| 1548 | Clostridium scatologenes |
| 755731 | Clostridium sp. BNL1100 |
| 1042156 | Clostridium sp. SY8519 |
| 1509 | Clostridium sporogenes |
| 394958 | Clostridium taeniosporum |
| 1231072 | Clostridium tetani 12124569 |
| 212717 | Clostridium tetani E88 |
| 84032 | Clostridium thermosuccinogenes |
| 1519 | Clostridium tyrobutyricum |
| 1619308 | Cnuibacter physcomitrellae |
| 28258 | Cobetia marina |
| 246410 | Coccidioides immitis RS |
| 222929 | Coccidioides posadasii C735 delta SOWgp |
| 574566 | Coccomyxa subellipsoidea C-169 |
| 1967665 | Cognaticolwellia beringensis |
| 2674991 | Cohnella candidum |
| 1445577 | Colletotrichum fioriniae PJ7 |
| 279058 | Collimonas arenae |
| 1005048 | Collimonas fungivorans Ter331 |
| 279113 | Collimonas pratensis |
| 74426 | Collinsella aerofaciens |
| 8932 | Columba livia |
| 167879 | Colwellia psychrerythraea 34H |
| 2161872 | Colwellia sp. Arc7-D |
| 58049 | Colwellia sp. MT41 |
| 1816218 | Colwellia sp. PAMC 20917 |
| 1816219 | Colwellia sp. PAMC 21821 |
| 225992 | Comamonas kerstersii |
| 1082851 | Comamonas serinivorans |
| 1392005 | Comamonas testosteroni TK102 |
| 363952 | Comamonas thiooxydans |
| 2070537 | Commensalibacter melissae |

| NCBI Taxon Identifier | Organism Name |
|---|---|
| 1423720 | Companilactobacillus alimentarius DSM 20249 |
| 1847728 | Companilactobacillus allii |
| 392416 | Companilactobacillus crustorum |
| 1007676 | Companilactobacillus ginsenosidimutans |
| 1074467 | Companilactobacillus heilongjiangensis |
| 469383 | Conexibacter woesei DSM 14684 |
| 741705 | Coniophora puteana RWD-64-598 SS2 |
| 240176 | Coprinopsis cinerea okayama7#130 |
| 717962 | Coprococcus catus GD/7 |
| 751585 | Coprococcus sp. ART55/1 |
| 309798 | Coprothermobacter proteolyticus DSM 5265 |
| 583355 | Coraliomargarita akajimensis DSM 45221 |
| 1144275 | Corallococcus coralloides DSM 2259 |
| 1189310 | Corallococcus macrosporus DSM 14697 |
| 983644 | Cordyceps militaris CM01 |
| 700015 | Coriobacterium glomerans PW2 |
| 932674 | Corvus cornix cornix |
| 649754 | Corynebacterium ammoniagenes DSM 20306 |
| 1431546 | Corynebacterium aquilae DSM 44791 |
| 1348662 | Corynebacterium argentoratense DSM 44202 |
| 191610 | Corynebacterium atypicum |
| 548476 | Corynebacterium aurimucosum ATCC 700975 |
| 1121353 | Corynebacterium callunae DSM 20147 |
| 161896 | Corynebacterium camporealensis |
| 1285583 | Corynebacterium casei LMG S-19264 |
| 1652495 | Corynebacterium crudilactis |
| 931089 | Corynebacterium deserti GIMN1.010 |
| 1717 | Corynebacterium diphtheriae |
| 698966 | Corynebacterium diphtheriae 241 |
| 698962 | Corynebacterium diphtheriae 31A |
| 698973 | Corynebacterium diphtheriae BH8 |
| 698963 | Corynebacterium diphtheriae C7 (beta) |
| 698965 | Corynebacterium diphtheriae CDCE 8392 |
| 698967 | Corynebacterium diphtheriae HC01 |
| 698968 | Corynebacterium diphtheriae HC02 |
| 698969 | Corynebacterium diphtheriae HC03 |
| 698970 | Corynebacterium diphtheriae HC04 |
| 698972 | Corynebacterium diphtheriae INCA 402 |
| 257309 | Corynebacterium diphtheriae NCTC 13129 |
| 698964 | Corynebacterium diphtheriae PW8 |
| 698971 | Corynebacterium diphtheriae VA01 |

| NCBI Taxon Identifier | Organism Name |
|---|---|
| 558173 | Corynebacterium doosanense CAU 212 = DSM 45436 |
| 196164 | Corynebacterium efficiens YS-314 |
| 1050174 | Corynebacterium epidermidicanis |
| 1451189 | Corynebacterium falsenii DSM 44353 |
| 28028 | Corynebacterium flavescens |
| 1437875 | Corynebacterium frankenforstense DSM 45800 |
| 187491 | Corynebacterium glaucum |
| 1718 | Corynebacterium glutamicum |
| 1718 | Corynebacterium glutamicum |
| 1718 | Corynebacterium glutamicum |
| 196627 | Corynebacterium glutamicum ATCC 13032 |
| 196627 | Corynebacterium glutamicum ATCC 13032 |
| 1204414 | Corynebacterium glutamicum K051 |
| 1310161 | Corynebacterium glutamicum MB001 |
| 340322 | Corynebacterium glutamicum R |
| 1232381 | Corynebacterium glutamicum SCgG1 |
| 1232383 | Corynebacterium glutamicum SCgG2 |
| 1404245 | Corynebacterium glyciniphilum AJ 3170 |
| 1121362 | Corynebacterium halotolerans YIM 70093 = DSM 44683 |
| 1223515 | Corynebacterium humireducens NBRC 106098 = DSM 45392 |
| 156978 | Corynebacterium imitans |
| 306537 | Corynebacterium jeikeium K411 |
| 645127 | Corynebacterium kroppenstedtii DSM 44385 |
| 35755 | Corynebacterium kutscheri |
| 1408189 | Corynebacterium lactis RW2-5 |
| 1224162 | Corynebacterium marinum DSM 44953 |
| 1224163 | Corynebacterium maris DSM 45190 |
| 38301 | Corynebacterium minutissimum |
| 571915 | Corynebacterium mustelae |
| 161895 | Corynebacterium phocae |
| 1719 | Corynebacterium pseudotuberculosis |
| 1719 | Corynebacterium pseudotuberculosis |
| 1719 | Corynebacterium pseudotuberculosis |
| 1087454 | Corynebacterium pseudotuberculosis 1/06-A |
| 679896 | Corynebacterium pseudotuberculosis 1002 |
| 1168865 | Corynebacterium pseudotuberculosis 258 |
| 1089446 | Corynebacterium pseudotuberculosis 267 |
| 1087452 | Corynebacterium pseudotuberculosis 3/99-5 |
| 1087451 | Corynebacterium pseudotuberculosis 31 |
| 1074485 | Corynebacterium pseudotuberculosis 316 |
| 1087453 | Corynebacterium pseudotuberculosis 42/02-A |

| NCBI Taxon Identifier | Organism Name |
|---|---|
| 681645 | Corynebacterium pseudotuberculosis C231 |
| 935697 | Corynebacterium pseudotuberculosis CIP 52.97 |
| 1161911 | Corynebacterium pseudotuberculosis Cp162 |
| 765874 | Corynebacterium pseudotuberculosis FRC41 |
| 889513 | Corynebacterium pseudotuberculosis I19 |
| 1117942 | Corynebacterium pseudotuberculosis P54B96 |
| 935298 | Corynebacterium pseudotuberculosis PAT10 |
| 1006579 | Corynebacterium ramonii FRC0011 |
| 662755 | Corynebacterium resistens DSM 45100 |
| 146827 | Corynebacterium simulans |
| 161899 | Corynebacterium singulare |
| 1487956 | Corynebacterium sp. ATCC 6931 |
| 1437874 | Corynebacterium sphenisci DSM 44792 |
| 1705 | Corynebacterium stationis |
| 43770 | Corynebacterium striatum |
| 1200352 | Corynebacterium terpenotabidum Y-11 |
| 136857 | Corynebacterium testudinoris |
| 65058 | Corynebacterium ulcerans |
| 65058 | Corynebacterium ulcerans |
| 65058 | Corynebacterium ulcerans |
| 65058 | Corynebacterium ulcerans |
| 996634 | Corynebacterium ulcerans 0102 |
| 945711 | Corynebacterium ulcerans 809 |
| 945712 | Corynebacterium ulcerans BR-AD22 |
| 504474 | Corynebacterium urealyticum DSM 7109 |
| 1267754 | Corynebacterium urealyticum DSM 7111 |
| 401472 | Corynebacterium ureicelerivorans |
| 1072256 | Corynebacterium uterequi |
| 858619 | Corynebacterium variabile DSM 44702 |
| 1224164 | Corynebacterium vitaeruminis DSM 20294 |
| 93934 | Coturnix japonica |
| 434923 | Coxiella burnetii CbuG_Q212 |
| 434924 | Coxiella burnetii CbuK_Q154 |
| 434922 | Coxiella burnetii Dugway 5J108-111 |
| 360115 | Coxiella burnetii RSA 331 |
| 227377 | Coxiella burnetii RSA 493 |
| 325775 | Coxiella endosymbiont of Amblyomma americanum |
| 10029 | Cricetulus griseus |
| 1173022 | Crinalium epipsammum PCC 9333 |
| 216432 | Croceibacter atlanticus HTCC2559 |
| 1267766 | Croceibacterium atlanticum |

| NCBI Taxon Identifier | Organism Name |
|---|---|
| 450378 | Croceicoccus marinus |
| 1348774 | Croceicoccus naphthovorans |
| 43989 | Crocosphaera subtropica ATCC 51142 |
| 165597 | Crocosphaera watsonii WH 8501 |
| 1073999 | Cronobacter condimenti 1330 |
| 1159554 | Cronobacter dublinensis subsp. dublinensis LMG 23823 |
| 413503 | Cronobacter malonaticus |
| 1159491 | Cronobacter malonaticus LMG 23826 |
| 1159613 | Cronobacter muytjensii ATCC 51329 |
| 28141 | Cronobacter sakazakii |
| 290339 | Cronobacter sakazakii ATCC BAA-894 |
| 1138308 | Cronobacter sakazakii ES15 |
| 956149 | Cronobacter sakazakii SP291 |
| 693216 | Cronobacter turicensis z3032 |
| 1074000 | Cronobacter universalis NCTC 9529 |
| 670052 | Cryobacterium arcticum |
| 1978566 | Cryobacterium sp. LW097 |
| 469378 | Cryptobacterium curtum DSM 15641 |
| 367775 | Cryptococcus gattii WM276 |
| 283643 | Cryptococcus neoformans var. neoformans B-3501A |
| 214684 | Cryptococcus neoformans var. neoformans JEC21 |
| 353151 | Cryptosporidium hominis TU502 |
| 353152 | Cryptosporidium parvum Iowa II |
| 3656 | Cucumis melo |
| 3659 | Cucumis sativus |
| 3661 | Cucurbita maxima |
| 3662 | Cucurbita moschata |
| 3664 | Cucurbita pepo subsp. pepo |
| 7176 | Culex quinquefasciatus |
| 1673428 | Cuniculiplasma divulgatum |
| 68895 | Cupriavidus basilensis |
| 1267562 | Cupriavidus gilardii CR3 |
| 367825 | Cupriavidus malaysiensis |
| 266264 | Cupriavidus metallidurans CH34 |
| 106590 | Cupriavidus necator |
| 381666 | Cupriavidus necator H16 |
| 1042878 | Cupriavidus necator N-1 |
| 82633 | Cupriavidus pauculus |
| 264198 | Cupriavidus pinatubonensis JMP134 |
| 876364 | Cupriavidus sp. USMAA2-4 |
| 1389192 | Cupriavidus sp. USMAHM13 |

| NCBI Taxon Identifier | Organism Name |
|---|---|
| 977880 | Cupriavidus taiwanensis LMG 19424 |
| 1905847 | Curtobacterium sp. BH-2-1-1 |
| 1561023 | Curtobacterium sp. MR_MD2014 |
| 1747 | Cutibacterium acnes |
| 909952 | Cutibacterium acnes 266 |
| 1031709 | Cutibacterium acnes 6609 |
| 1234380 | Cutibacterium acnes C1 |
| 1276648 | Cutibacterium acnes hdn-1 |
| 1134454 | Cutibacterium acnes HL096PA1 |
| 267747 | Cutibacterium acnes KPA171202 |
| 553199 | Cutibacterium acnes SK137 |
| 1091045 | Cutibacterium acnes subsp. defendens ATCC 11828 |
| 1114967 | Cutibacterium acnes TypeIA2 P.acn17 |
| 1114969 | Cutibacterium acnes TypeIA2 P.acn31 |
| 1114966 | Cutibacterium acnes TypeIA2 P.acn33 |
| 1170318 | Cutibacterium avidum 44067 |
| 33011 | Cutibacterium granulosum |
| 2559073 | Cutibacterium modestum |
| 280699 | Cyanidioschyzon merolae strain 10D |
| 156563 | Cyanistes caeruleus |
| 755178 | Cyanobacterium aponinum PCC 10605 |
| 1228987 | cyanobacterium endosymbiont of Epithemia turgida isolate EtSB Lake Yunoko |
| 1763363 | cyanobacterium endosymbiont of Rhopalodia gibberula |
| 2054282 | Cyanobacterium sp. HL-69 |
| 292563 | Cyanobacterium stanieri PCC 7202 |
| 292564 | Cyanobium gracile PCC 6307 |
| 1851505 | Cyanobium sp. NIES-981 |
| 395961 | Cyanothece sp. PCC 7425 |
| 320787 | Cyclobacterium amurskyense |
| 880070 | Cyclobacterium marinum DSM 745 |
| 385025 | Cycloclasticus sp. P1 |
| 728003 | Cycloclasticus sp. PY97N |
| 1198232 | Cycloclasticus zancles 78-ME |
| 1457365 | Cyclonatronum proteinivorum |
| 56107 | Cylindrospermum stagnale PCC 7417 |
| 59895 | Cynara cardunculus var. scolymus |
| 244447 | Cynoglossus semilaevis |
| 7962 | Cyprinus carpio |
| 43 | Cystobacter fuscus |
| 859143 | Cytobacillus kochii |
| 1196031 | Cytobacillus oceanisediminis 2691 |

| NCBI Taxon Identifier | Organism Name |
|---|---|
| 269798 | Cytophaga hutchinsonii ATCC 33406 |
| 13035 | Dactylococcopsis salina PCC 8305 |
| 278856 | Danaus plexippus plexippus |
| 7955 | Danio rerio |
| 6669 | Daphnia pulex |
| 79200 | Daucus carota subsp. sativus |
| 284592 | Debaryomyces hansenii CBS767 |
| 159087 | Dechloromonas aromatica RCB |
| 2231055 | Dechloromonas sp. HYN0024 |
| 639282 | Deferribacter desulfuricans SSM1 |
| 1006576 | Defluviitoga tunisiensis |
| 871738 | Dehalobacter restrictus DSM 9455 |
| 1131462 | Dehalobacter sp. CF |
| 1147129 | Dehalobacter sp. DCA |
| 243164 | Dehalococcoides mccartyi 195 |
| 216389 | Dehalococcoides mccartyi BAV1 |
| 1193806 | Dehalococcoides mccartyi BTF08 |
| 255470 | Dehalococcoides mccartyi CBDB1 |
| 1432059 | Dehalococcoides mccartyi CG1 |
| 1432060 | Dehalococcoides mccartyi CG4 |
| 1432061 | Dehalococcoides mccartyi CG5 |
| 1193807 | Dehalococcoides mccartyi DCMB5 |
| 633145 | Dehalococcoides mccartyi GT |
| 1388758 | Dehalococcoides mccartyi GY50 |
| 311424 | Dehalococcoides mccartyi VS |
| 1522671 | Dehalococcoides sp. UCH007 |
| 1839801 | Dehalogenimonas formicexedens |
| 552811 | Dehalogenimonas lykanthroporepellens BL-DC-9 |
| 943347 | Dehalogenimonas sp. WBC-2 |
| 1768108 | Deinococcus actinosclerus |
| 546414 | Deinococcus deserti VCD115 |
| 317577 | Deinococcus ficus |
| 319795 | Deinococcus geothermalis DSM 11300 |
| 745776 | Deinococcus gobiensis I-0 |
| 2202254 | Deinococcus irradiatisoli |
| 709986 | Deinococcus maricopensis DSM 21211 |
| 937777 | Deinococcus peraridilitoris DSM 19664 |
| 693977 | Deinococcus proteolyticus MRP |
| 1182568 | Deinococcus puniceus |
| 243230 | Deinococcus radiodurans R1 = ATCC 13939 = DSM 20539 |
| 57497 | Deinococcus radiopugnans |

| NCBI Taxon Identifier | Organism Name |
|---|---|
| 1309411 | Deinococcus soli (ex Cha et al. 2016) |
| 980427 | Deinococcus wulumuqiensis |
| 398578 | Delftia acidovorans SPH-1 |
| 742013 | Delftia sp. Cs1-4 |
| 1920191 | Delftia sp. HK171 |
| 180282 | Delftia tsuruhatensis |
| 906689 | Dendrobium catenatum |
| 77166 | Dendroctonus ponderosae |
| 79604 | Denitrobacterium detoxificans |
| 522772 | Denitrovibrio acetiphilus DSM 12809 |
| 1630135 | Dermabacter vaginalis |
| 1274 | Dermacoccus nishinomiyaensis |
| 6956 | Dermatophagoides pteronyssinus |
| 1863 | Dermatophilus congolensis |
| 644282 | Desulfarculus baarsii DSM 2075 |
| 439235 | Desulfatibacillum alkenivorans AK-01 |
| 756499 | Desulfitobacterium dehalogenans ATCC 51507 |
| 871963 | Desulfitobacterium dichloroeliminans LMG P-21439 |
| 272564 | Desulfitobacterium hafniense DCB-2 |
| 138119 | Desulfitobacterium hafniense Y51 |
| 871968 | Desulfitobacterium metallireducens DSM 15288 |
| 880072 | Desulfobacca acetoxidans DSM 11109 |
| 651182 | Desulfobacula toluolica Tol2 |
| 1986146 | Desulfobulbus oralis |
| 577650 | Desulfobulbus propionicus DSM 2032 |
| 1167006 | Desulfocapsa sulfexigens DSM 10523 |
| 897 | Desulfococcus multivorans |
| 690850 | Desulfocurvibacter africanus subsp. africanus str. Walvis Bay |
| 485916 | Desulfofarcimen acetoxidans DSM 771 |
| 760568 | Desulfofundulus kuznetsovii DSM 6115 |
| 485915 | Desulfohalobium retbaense DSM 5692 |
| 525897 | Desulfomicrobium baculatum DSM 4028 |
| 888061 | Desulfomicrobium orale DSM 12838 |
| 706587 | Desulfomonile tiedjei DSM 6799 |
| 1833852 | Desulforamulus ferrireducens |
| 349161 | Desulforamulus reducens MI-1 |
| 696281 | Desulforamulus ruminis DSM 2154 |
| 177437 | Desulforapulum autotrophicum HRM2 |
| 767817 | Desulfoscipio gibsoniae DSM 7213 |
| 646529 | Desulfosporosinus acidiphilus SJ4 |
| 768704 | Desulfosporosinus meridiei DSM 13257 |

| NCBI Taxon Identifier | Organism Name |
|---|---|
| 768706 | Desulfosporosinus orientis DSM 765 |
| 96561 | Desulfosudis oleivorans Hxd3 |
| 177439 | Desulfotalea psychrophila LSv54 |
| 868595 | Desulfotomaculum nigrificans CO-1-SRB |
| 525146 | Desulfovibrio desulfuricans ATCC 27774 |
| 44742 | Desulfovibrio fairfieldensis |
| 241368 | Desulfovibrio ferrophilus |
| 901 | Desulfovibrio piger |
| 631220 | Desulfovibrio sp. G11 |
| 694431 | Desulfurella acetivorans A63 |
| 653733 | Desulfurispirillum indicum S5 |
| 589865 | Desulfurivibrio alkaliphilus AHT 2 |
| 868864 | Desulfurobacterium thermolithotrophum DSM 11699 |
| 490899 | Desulfurococcus amylolyticus 1221n |
| 768672 | Desulfurococcus amylolyticus DSM 16532 |
| 765177 | Desulfurococcus mucosus DSM 2162 |
| 1603606 | Desulfuromonas soudanensis |
| 1823759 | Desulfuromonas sp. DDH964 |
| 1643450 | Devosia sp. H5989 |
| 2083786 | Devosia sp. I507 |
| 39950 | Dialister pneumosintes |
| 164759 | Diaphorobacter nitroreducens |
| 246195 | Dichelobacter nodosus VCS1703A |
| 732165 | Dichomitus squalens LYAD-421 SS1 |
| 561229 | Dickeya chrysanthemi Ech1591 |
| 198628 | Dickeya dadantii 3937 |
| 1306405 | Dickeya dianthicola RNS04.9 |
| 1778540 | Dickeya fangzhongdai |
| 590409 | Dickeya parazeae Ech586 |
| 1089444 | Dickeya solani |
| 1225786 | Dickeya solani IPO 2222 |
| 1427366 | Dickeya zeae EC1 |
| 309799 | Dictyoglomus thermophilum H-6-12 |
| 515635 | Dictyoglomus turgidum DSM 6724 |
| 352472 | Dictyostelium discoideum AX4 |
| 5786 | Dictyostelium purpureum |
| 546160 | Dietzia lutea |
| 139021 | Dietzia psychralcaliphila |
| 2052657 | Dietzia sp. JS16-p6b |
| 712270 | Dietzia sp. oral taxon 368 |
| 499555 | Dietzia timorensis |

| NCBI Taxon Identifier | Organism Name |
|---|---|
| 609295 | Dinoponera quadriceps |
| 398580 | Dinoroseobacter shibae DFL 12 = DSM 16493 |
| 143948 | Diuraphis noxia |
| 1300342 | Dokdonella koreensis DS-123 |
| 1300343 | Dokdonia donghaensis DSW-1 |
| 983548 | Dokdonia sp. 4H-3-7-5 |
| 2173169 | Dokdonia sp. Dokd-P16 |
| 313590 | Dokdonia sp. MED134 |
| 1168034 | Draconibacterium orientale |
| 1072389 | Drepanopeziza brunnea f. sp. 'multigermtubi' MB_m1 |
| 7217 | Drosophila ananassae |
| 7220 | Drosophila erecta |
| 7222 | Drosophila grimshawi |
| 7227 | Drosophila melanogaster |
| 7230 | Drosophila mojavensis |
| 7234 | Drosophila persimilis |
| 46245 | Drosophila pseudoobscura pseudoobscura |
| 7238 | Drosophila sechellia |
| 7240 | Drosophila simulans |
| 7244 | Drosophila virilis |
| 7260 | Drosophila willistoni |
| 7245 | Drosophila yakuba |
| 1619313 | Duffyella gerundensis |
| 66656 | Durio zibethinus |
| 471854 | Dyadobacter fermentans DSM 18053 |
| 1217721 | Dyella japonica A8 |
| 1379159 | Dyella jiangningensis |
| 2501295 | Dyella sp. M7H15-1 |
| 445710 | Dyella thiooxydans |
| 1795355 | Echinicola strongylocentroti |
| 926556 | Echinicola vietnamensis DSM 17526 |
| 6210 | Echinococcus granulosus |
| 1442136 | Ectothiorhodospira sp. BSL-9 |
| 667120 | Edwardsiella anguillarum ET080813 |
| 93378 | Edwardsiella hoshinae |
| 634503 | Edwardsiella ictaluri 93-146 |
| 1288122 | Edwardsiella piscicida C07-087 |
| 498217 | Edwardsiella piscicida EIB202 |
| 1578828 | Edwardsiella sp. EA181011 |
| 1650654 | Edwardsiella sp. LADL05-105 |
| 718251 | Edwardsiella tarda FL6-60 |

| NCBI Taxon Identifier | Organism Name |
|---|---|
| 479437 | Eggerthella lenta DSM 2243 |
| 502558 | Eggerthella sp. YY7918 |
| 1670831 | Egibacter rhizosphaerae |
| 188379 | Egretta garzetta |
| 269484 | Ehrlichia canis str. Jake |
| 205920 | Ehrlichia chaffeensis str. Arkansas |
| 1249651 | Ehrlichia chaffeensis str. Heartland |
| 1249652 | Ehrlichia chaffeensis str. Jax |
| 1249653 | Ehrlichia chaffeensis str. Liberty |
| 1249654 | Ehrlichia chaffeensis str. Osceola |
| 1249655 | Ehrlichia chaffeensis str. Saint Vincent |
| 1249656 | Ehrlichia chaffeensis str. Wakulla |
| 1249650 | Ehrlichia chaffeensis str. West Paces |
| 391036 | Ehrlichia japonica |
| 1423892 | Ehrlichia muris AS145 |
| 302409 | Ehrlichia ruminantium str. Gardel |
| 254945 | Ehrlichia ruminantium str. Welgevonden |
| 254945 | Ehrlichia ruminantium str. Welgevonden |
| 539 | Eikenella corrodens |
| 51953 | Elaeis guineensis |
| 8005 | Electrophorus electricus |
| 1117645 | Elizabethkingia anophelis |
| 1352940 | Elizabethkingia anophelis FMS-007 |
| 1338011 | Elizabethkingia anophelis NUHP1 |
| 1756149 | Elizabethkingia bruuniana |
| 238 | Elizabethkingia meningoseptica |
| 172045 | Elizabethkingia miricola |
| 1756150 | Elizabethkingia ursingii |
| 445932 | Elusimicrobium minutum Pei191 |
| 280463 | Emiliania huxleyi CCMP1516 |
| 929562 | Emticicia oligotrophica DSM 17448 |
| 284813 | Encephalitozoon cuniculi GB-M1 |
| 907965 | Encephalitozoon hellem ATCC 50504 |
| 876142 | Encephalitozoon intestinalis ATCC 50506 |
| 1178016 | Encephalitozoon romaleae SJ-2008 |
| 1408281 | Endomicrobium proavitum |
| 86106 | endosymbiont of Acanthamoeba sp. UWC8 |
| 1303921 | endosymbiont of Bathymodiolus septemdierum str. Myojin knoll |
| 1248727 | endosymbiont of unidentified scaly snail isolate Monju |
| 106592 | Ensifer adhaerens |
| 1416753 | Ensifer adhaerens OV14 |

| NCBI Taxon Identifier | Organism Name |
|---|---|
| 370354 | Entamoeba dispar SAW760 |
| 294381 | Entamoeba histolytica HM-1:IMSS |
| 370355 | Entamoeba invadens IP1 |
| 1421338 | Enterobacter asburiae L1 |
| 69218 | Enterobacter cancerogenus |
| 550 | Enterobacter cloacae |
| 550 | Enterobacter cloacae |
| 550 | Enterobacter cloacae |
| 1333850 | Enterobacter cloacae ECNIH2 |
| 716541 | Enterobacter cloacae subsp. cloacae ATCC 13047 |
| 1211025 | Enterobacter cloacae subsp. cloacae ENHKU01 |
| 718254 | Enterobacter cloacae subsp. cloacae NCTC 9394 |
| 1104326 | Enterobacter cloacae subsp. dissolvens SDM |
| 158836 | Enterobacter hormaechei |
| 1333851 | Enterobacter hormaechei subsp. hoffmannii ECNIH3 |
| 1333849 | Enterobacter hormaechei subsp. hoffmannii ECR091 |
| 301105 | Enterobacter hormaechei subsp. hormaechei |
| 299766 | Enterobacter hormaechei subsp. steigerwaltii |
| 1296536 | Enterobacter hormaechei subsp. xiangfangensis |
| 1296536 | Enterobacter hormaechei subsp. xiangfangensis |
| 208224 | Enterobacter kobei |
| 299767 | Enterobacter ludwigii |
| 299767 | Enterobacter ludwigii |
| 1812935 | Enterobacter roggenkampii |
| 1812935 | Enterobacter roggenkampii |
| 885040 | Enterobacter soli |
| 399742 | Enterobacter sp. 638 |
| 1560339 | Enterobacter sp. E20 |
| 1692238 | Enterobacter sp. FY-07 |
| 1166130 | Enterobacter sp. R4-368 |
| 1920109 | Enterobacteriaceae bacterium ENNIH2 |
| 2052938 | Enterobacteriaceae bacterium S05 |
| 693444 | Enterobacteriaceae bacterium strain FGI 57 |
| 208479 | Enterocloster bolteae |
| 565655 | Enterococcus casseliflavus EC20 |
| 53345 | Enterococcus durans |
| 1351 | Enterococcus faecalis |
| 936153 | Enterococcus faecalis 62 |
| 1201292 | Enterococcus faecalis ATCC 29212 |
| 1206105 | Enterococcus faecalis D32 |
| 1287066 | Enterococcus faecalis DENG1 |

| NCBI Taxon Identifier | Organism Name |
|---|---|
| 474186 | Enterococcus faecalis OG1RF |
| 1261557 | Enterococcus faecalis str. Symbioflor 1 |
| 226185 | Enterococcus faecalis V583 |
| 1158600 | Enterococcus faecium ATCC 8459 = NRRL B-2354 |
| 1155766 | Enterococcus faecium Aus0004 |
| 1305849 | Enterococcus faecium Aus0085 |
| 333849 | Enterococcus faecium DO |
| 1344042 | Enterococcus faecium T110 |
| 1353 | Enterococcus gallinarum |
| 160453 | Enterococcus gilvus |
| 768486 | Enterococcus hirae ATCC 9790 |
| 1300150 | Enterococcus mundtii QU 25 |
| 332949 | Enterococcus silesiacus |
| 417368 | Enterococcus thailandicus |
| 74700 | Entomoplasma freundtii |
| 2268024 | Ephemeroptericola cinctiostellae |
| 9793 | Equus asinus |
| 9796 | Equus caballus |
| 9798 | Equus przewalskii |
| 931890 | Eremothecium cymbalariae DBVPG#7215 |
| 284811 | Eremothecium gossypii ATCC 10895 |
| 716540 | Erwinia amylovora ATCC 49946 |
| 665029 | Erwinia amylovora CFBP1430 |
| 634500 | Erwinia billingiae Eb661 |
| 55211 | Erwinia persicina |
| 644651 | Erwinia pyrifoliae DSM 12163 |
| 634499 | Erwinia pyrifoliae Ep1/96 |
| 215689 | Erwinia sp. Ejp617 |
| 465817 | Erwinia tasmaniensis Et1/99 |
| 1514105 | Erysipelothrix larvae |
| 2485784 | Erysipelothrix piscisicarius |
| 650150 | Erysipelothrix rhusiopathiae str. Fujisawa |
| 1313290 | Erysipelothrix rhusiopathiae SY1027 |
| 1834207 | Erysipelotrichaceae bacterium I46 |
| 2182384 | Erythrobacter aureus |
| 39960 | Erythrobacter litoralis |
| 314225 | Erythrobacter litoralis HTCC2594 |
| 1112 | Erythrobacter neustonensis |
| 2011159 | Erythrobacter sp. KY5 |
| 1440052 | Escherichia albertii KF1 |
| 216592 | Escherichia coli 042 |

| NCBI Taxon Identifier | Organism Name |
|---|---|
| 362663 | Escherichia coli 536 |
| 585055 | Escherichia coli 55989 |
| 655817 | Escherichia coli ABU 83972 |
| 405955 | Escherichia coli APEC O1 |
| 1274814 | Escherichia coli APEC O78 |
| 481805 | Escherichia coli ATCC 8739 |
| 413997 | Escherichia coli B str. REL606 |
| 469008 | Escherichia coli BL21(DE3) |
| 469008 | Escherichia coli BL21(DE3) |
| 866768 | Escherichia coli 'BL21-Gold(DE3)pLysS AG' |
| 595496 | Escherichia coli BW2952 |
| 199310 | Escherichia coli CFT073 |
| 536056 | Escherichia coli DH1 |
| 536056 | Escherichia coli DH1 |
| 585397 | Escherichia coli ED1a |
| 316401 | Escherichia coli ETEC H10407 |
| 331112 | Escherichia coli HS |
| 585034 | Escherichia coli IAI1 |
| 585057 | Escherichia coli IAI39 |
| 714962 | Escherichia coli IHE3034 |
| 1355100 | Escherichia coli JJ1886 |
| 595495 | Escherichia coli KO11FL |
| 595495 | Escherichia coli KO11FL |
| 591946 | Escherichia coli LF82 |
| 1335916 | Escherichia coli LY180 |
| 1033813 | Escherichia coli NA114 |
| 585395 | Escherichia coli O103:H2 str. 12009 |
| 1134782 | Escherichia coli O104:H4 str. 2009EL-2050 |
| 1133853 | Escherichia coli O104:H4 str. 2009EL-2071 |
| 1133852 | Escherichia coli O104:H4 str. 2011C-3493 |
| 585396 | Escherichia coli O111:H- str. 11128 |
| 574521 | Escherichia coli O127:H6 str. E2348/69 |
| 331111 | Escherichia coli O139:H28 str. E24377A |
| 1248902 | Escherichia coli O145:H28 str. RM13514 |
| 1248915 | Escherichia coli O145:H28 str. RM13516 |
| 444450 | Escherichia coli O157:H7 str. EC4115 |
| 155864 | Escherichia coli O157:H7 str. EDL933 |
| 386585 | Escherichia coli O157:H7 str. Sakai |
| 544404 | Escherichia coli O157:H7 str. TW14359 |
| 941322 | Escherichia coli O25b:H4-ST131 |
| 573235 | Escherichia coli O26:H11 str. 11368 |

| NCBI Taxon Identifier | Organism Name |
|---|---|
| 701177 | Escherichia coli O55:H7 str. CB9615 |
| 1048689 | Escherichia coli O55:H7 str. RM12579 |
| 1072459 | Escherichia coli O7:K1 str. CE10 |
| 685038 | Escherichia coli O83:H1 str. NRG 857C |
| 910348 | Escherichia coli P12b |
| 1382700 | Escherichia coli PMV-1 |
| 585035 | Escherichia coli S88 |
| 409438 | Escherichia coli SE11 |
| 431946 | Escherichia coli SE15 |
| 439855 | Escherichia coli SMS-3-5 |
| 885275 | Escherichia coli str. 'clone D i14' |
| 885276 | Escherichia coli str. 'clone D i2' |
| 316385 | Escherichia coli str. K-12 substr. DH10B |
| 1110693 | Escherichia coli str. K-12 substr. MDS42 |
| 511145 | Escherichia coli str. K-12 substr. MG1655 |
| 316407 | Escherichia coli str. K-12 substr. W3110 |
| 869729 | Escherichia coli UM146 |
| 585056 | Escherichia coli UMN026 |
| 696406 | Escherichia coli UMNK88 |
| 364106 | Escherichia coli UTI89 |
| 566546 | Escherichia coli W |
| 566546 | Escherichia coli W |
| 741093 | Escherichia coli Xuzhou21 |
| 585054 | Escherichia fergusonii ATCC 35469 |
| 1499973 | Escherichia marmotae |
| 8010 | Esox lucius |
| 663278 | Ethanoligenens harbinense YUAN-3 |
| 903814 | Eubacterium limosum KIST612 |
| 888727 | Eubacterium sulci ATCC 35585 |
| 71139 | Eucalyptus grandis |
| 72664 | Eutrema salsugineum |
| 1287681 | Eutypa lata UCREL1 |
| 1761453 | Euzebyella marina |
| 649639 | Evansella cellulosilytica DSM 2522 |
| 2652724 | Exaiptasia diaphana |
| 1087448 | Exiguobacterium antarcticum B7 |
| 262543 | Exiguobacterium sibiricum 255-15 |
| 360911 | Exiguobacterium sp. AT1b |
| 1399115 | Exiguobacterium sp. MH3 |
| 1849031 | Exiguobacterium sp. U13-1 |
| 718252 | Faecalibacterium prausnitzii L2-6 |

| NCBI Taxon Identifier | Organism Name |
|---|---|
| 657322 | Faecalibacterium prausnitzii SL3/3 |
| 1702221 | Faecalibaculum rodentium |
| 717960 | Faecalitalea cylindroides T2-87 |
| 345164 | Falco cherrug |
| 8954 | Falco peregrinus |
| 1579316 | Falsihalocynthiibacter arcticus |
| 236753 | Fastidiosipila sanguinis |
| 9685 | Felis catus |
| 1562970 | Fermentimonas caenicola |
| 550540 | Ferrimonas balearica DSM 9799 |
| 1188319 | Ferriphaselus amnicola |
| 589924 | Ferroglobus placidus DSM 10642 |
| 333146 | Ferroplasma acidarmanus Fer1 |
| 74969 | Ferroplasma acidiphilum |
| 1163730 | Fervidicoccus fontis Kam940 |
| 2423 | Fervidobacterium islandicum |
| 381764 | Fervidobacterium nodosum Rt17-B1 |
| 771875 | Fervidobacterium pennivorans DSM 9078 |
| 1166018 | Fibrella aestuarina BUZ 2 |
| 1834519 | Fibrella sp. ES10-3-2-2 |
| 59374 | Fibrobacter succinogenes subsp. succinogenes S85 |
| 59374 | Fibrobacter succinogenes subsp. succinogenes S85 |
| 59894 | Ficedula albicollis |
| 255247 | Fictibacillus arsenicus |
| 1221500 | Fictibacillus phosphorivorans |
| 546269 | Filifactor alocis ATCC 35896 |
| 477680 | Filimonas lacunae |
| 661478 | Fimbriimonas ginsengisoli Gsoil 348 |
| 334413 | Finegoldia magna ATCC 29328 |
| 1752063 | Fischerella sp. NIES-3754 |
| 516051 | Flagellimonas lutaonensis |
| 1383885 | Flagellimonas maritima |
| 2494373 | Flammeovirga pectinis |
| 1191459 | Flammeovirga sp. MY04 |
| 1257021 | Flammeovirgaceae bacterium 311 |
| 2496867 | Flaviflexus ciconiae |
| 1282737 | Flaviflexus salsibiostraticola |
| 1492898 | Flavisolibacter tropicus |
| 1803846 | Flavivirga eckloniae |
| 531844 | Flavobacteriaceae bacterium 3519-10 |
| 1150389 | Flavobacteriaceae bacterium UJ101 |

| NCBI Taxon Identifier | Organism Name |
|---|---|
| 459526 | Flavobacterium anhuiense |
| 1784713 | Flavobacterium arcticum |
| 1034807 | Flavobacterium branchiophilum FL-15 |
| 1041826 | Flavobacterium columnare ATCC 49512 |
| 1306519 | Flavobacterium commune |
| 1355330 | Flavobacterium faecale |
| 1492737 | Flavobacterium gilvum |
| 1094466 | Flavobacterium indicum GPTSA100-9 = DSM 17447 |
| 376686 | Flavobacterium johnsoniae UW101 |
| 1678728 | Flavobacterium kingsejongi |
| 96345 | Flavobacterium psychrophilum |
| 96345 | Flavobacterium psychrophilum |
| 96345 | Flavobacterium psychrophilum |
| 96345 | Flavobacterium psychrophilum |
| 96345 | Flavobacterium psychrophilum |
| 96345 | Flavobacterium psychrophilum |
| 1452725 | Flavobacterium psychrophilum FPG101 |
| 1452724 | Flavobacterium psychrophilum FPG3 |
| 402612 | Flavobacterium psychrophilum JIP02/86 |
| 292800 | Flavonifractor plautii |
| 717231 | Flexistipes sinusarabici DSM 4947 |
| 755732 | Fluviicola taffensis DSM 16823 |
| 158441 | Folsomia candida |
| 694068 | Fomitiporia mediterranea MF3/22 |
| 1336794 | Formosa sp. Hel1_33_131 |
| 1336795 | Formosa sp. Hel3_A1_48 |
| 101020 | Fragaria vesca subsp. vesca |
| 635003 | Fragilariopsis cylindrus CCMP1102 |
| 984129 | Francisella cf. novicida Fx1 |
| 1542390 | Francisella frigiditurris |
| 549298 | Francisella halioticida |
| 622488 | Francisella hispaniensis |
| 1088883 | Francisella hispaniensis FSC454 |
| 1356861 | Francisella orientalis LADL 07-285A |
| 1163389 | Francisella orientalis str. Toba 04 |
| 1086726 | Francisella persica ATCC VR-331 |
| 28110 | Francisella philomiragia |
| 28110 | Francisella philomiragia |
| 28110 | Francisella philomiragia |
| 28110 | Francisella philomiragia |
| 28110 | Francisella philomiragia |

| NCBI Taxon Identifier | Organism Name |
|---|---|
| 539329 | Francisella philomiragia subsp. philomiragia ATCC 25015 |
| 484022 | Francisella philomiragia subsp. philomiragia ATCC 25017 |
| 573569 | Francisella salina |
| 475375 | Francisella sp. MA067296 |
| 119857 | Francisella tularensis subsp. holarctica |
| 119857 | Francisella tularensis subsp. holarctica |
| 119857 | Francisella tularensis subsp. holarctica |
| 1232394 | Francisella tularensis subsp. holarctica F92 |
| 351581 | Francisella tularensis subsp. holarctica FSC200 |
| 458234 | Francisella tularensis subsp. holarctica FTNF002-00 |
| 376619 | Francisella tularensis subsp. holarctica LVS |
| 393011 | Francisella tularensis subsp. holarctica OSU18 |
| 1432652 | Francisella tularensis subsp. holarctica PHIT-FT049 |
| 441952 | Francisella tularensis subsp. mediasiatica FSC147 |
| 264 | Francisella tularensis subsp. novicida |
| 1452728 | Francisella tularensis subsp. novicida F6168 |
| 401614 | Francisella tularensis subsp. novicida U112 |
| 401614 | Francisella tularensis subsp. novicida U112 |
| 393115 | Francisella tularensis subsp. tularensis FSC198 |
| 510831 | Francisella tularensis subsp. tularensis NE061598 |
| 177416 | Francisella tularensis subsp. tularensis SCHU S4 |
| 1341656 | Francisella tularensis subsp. tularensis str. SCHU S4 substr. NR-28534 |
| 1001534 | Francisella tularensis subsp. tularensis TI0902 |
| 1001542 | Francisella tularensis subsp. tularensis TIGB03 |
| 418136 | Francisella tularensis subsp. tularensis WY96-3418 |
| 573570 | Francisella uliginis |
| 326424 | Frankia alni ACN14a |
| 106370 | Frankia casuarinae |
| 767434 | Frateuria aurantia DSM 6220 |
| 651822 | Fretibacterium fastidiosum |
| 1335048 | Frigidibacter mobilis |
| 1267021 | Frischella perrara |
| 1795630 | Frondihabitans sp. PAMC 28766 |
| 53444 | Fructilactobacillus lindneri |
| 714313 | Fructilactobacillus sanfranciscensis TMW 1.1304 |
| 1891926 | Fuerstiella marisgermanici |
| 229533 | Fusarium graminearum PH-1 |
| 426428 | Fusarium oxysporum f. sp. lycopersici 4287 |
| 1028729 | Fusarium pseudograminearum CS3096 |
| 660122 | Fusarium vanettenii 77-13-4 |
| 334819 | Fusarium verticillioides 7600 |

| NCBI Taxon Identifier | Organism Name |
|---|---|
| 1075 | Fuscovulum blasticum |
| 469607 | Fusobacterium animalis 4_8 |
| 457405 | Fusobacterium animalis 7_1 |
| 469615 | Fusobacterium gonidiaformans ATCC 25563 |
| 1307442 | Fusobacterium hwasookii ChDC F174 |
| 469616 | Fusobacterium mortiferum ATCC 9817 |
| 190304 | Fusobacterium nucleatum subsp. nucleatum ATCC 25586 |
| 860 | Fusobacterium periodonticum |
| 861 | Fusobacterium ulcerans |
| 856 | Fusobacterium varium |
| 469602 | Fusobacterium vincentii 3_1_27 |
| 469604 | Fusobacterium vincentii 3_1_36A2 |
| 130081 | Galdieria sulphuraria |
| 1005058 | Gallibacterium anatis UMN179 |
| 395494 | Gallionella capsiferriformans ES-2 |
| 9031 | Gallus gallus |
| 83406 | gamma proteobacterium HdN1 |
| 553190 | Gardnerella vaginalis 409-05 |
| 525284 | Gardnerella vaginalis ATCC 14019 |
| 1009464 | Gardnerella vaginalis HMP9231 |
| 1173025 | Geitlerinema sp. PCC 7407 |
| 146911 | Gekko japonicus |
| 29391 | Gemella morbillorum |
| 2040624 | Gemella sp. ND 6198 |
| 1785995 | Gemella sp. oral taxon 928 |
| 1615909 | Geminocystis sp. NIES-3708 |
| 1617448 | Geminocystis sp. NIES-3709 |
| 114 | Gemmata obscuriglobus |
| 1630693 | Gemmata sp. SH-PL17 |
| 379066 | Gemmatimonas aurantiaca T-27 |
| 1379270 | Gemmatimonas phototrophica |
| 861299 | Gemmatirosa kalamazoonensis |
| 2169400 | Gemmobacter aquarius |
| 483547 | Geoalkalibacter subterraneus |
| 1921421 | Geobacillus genomosp. 3 |
| 235909 | Geobacillus kaustophilus HTA426 |
| 169283 | Geobacillus lituanicus |
| 1629723 | Geobacillus sp. 12AMOR1 |
| 691437 | Geobacillus sp. C56-T3 |
| 1233873 | Geobacillus sp. GHH01 |
| 1813182 | Geobacillus sp. JS12 |

| NCBI Taxon Identifier | Organism Name |
|---|---|
| 1519377 | Geobacillus sp. LC300 |
| 471223 | Geobacillus sp. WCH70 |
| 581103 | Geobacillus sp. Y4.1MC1 |
| 550542 | Geobacillus sp. Y412MC52 |
| 544556 | Geobacillus sp. Y412MC61 |
| 272567 | Geobacillus stearothermophilus 10 |
| 129338 | Geobacillus subterraneus |
| 33938 | Geobacillus thermocatenulatus |
| 420246 | Geobacillus thermodenitrificans NG80-2 |
| 33941 | Geobacillus thermoleovorans |
| 1111068 | Geobacillus thermoleovorans CCB_US3_UF5 |
| 1340425 | Geobacter anodireducens |
| 269799 | Geobacter metallireducens GS-15 |
| 345632 | Geobacter pickeringii |
| 443143 | Geobacter sp. M18 |
| 443144 | Geobacter sp. M21 |
| 663917 | Geobacter sulfurreducens KN400 |
| 243231 | Geobacter sulfurreducens PCA |
| 526225 | Geodermatophilus obscurus DSM 43160 |
| 565033 | Geoglobus acetivorans |
| 113653 | Geoglobus ahangari |
| 2341112 | Georhizobium profundi |
| 48883 | Geospiza fortis |
| 1424294 | Geosporobacter ferrireducens |
| 316067 | Geotalea daltonii FRC-32 |
| 351605 | Geotalea uraniireducens Rf4 |
| 139438 | Geovibrio thiophilus |
| 184922 | Giardia lamblia ATCC 50803 |
| 929813 | Gibbsiella quercinecans |
| 1196095 | Gilliamella apicola |
| 1085623 | Glaciecola nitratireducens FR1064 |
| 983545 | Glaciecola sp. 4H-3-7+YE-5 |
| 738 | Glaesserella parasuis |
| 738 | Glaesserella parasuis |
| 557723 | Glaesserella parasuis SH0165 |
| 1322346 | Glaesserella parasuis ZJ0906 |
| 2030797 | Glaesserella sp. 15-184 |
| 1116229 | Glarea lozoyensis ATCC 20868 |
| 1183438 | Gloeobacter kilaueensis JS1 |
| 251221 | Gloeobacter violaceus PCC 7421 |
| 1173026 | Gloeocapsa sp. PCC 7428 |

| NCBI Taxon Identifier | Organism Name |
| --- | --- |
| 670483 | Gloeophyllum trabeum ATCC 11539 |
| 65393 | Gloeothece citriformis PCC 7424 |
| 497965 | Gloeothece verrucosa PCC 7822 |
| 272568 | Gluconacetobacter diazotrophicus PA1 5 |
| 272568 | Gluconacetobacter diazotrophicus PA1 5 |
| 318683 | Gluconobacter albidus |
| 290633 | Gluconobacter oxydans 621H |
| 1288313 | Gluconobacter oxydans DSM 3504 |
| 1224746 | Gluconobacter oxydans H24 |
| 861360 | Glutamicibacter arilaitensis Re117 |
| 1933880 | Glutamicibacter halophytocola |
| 3847 | Glycine max |
| 1434191 | Glycocaulis alkaliphilus |
| 84096 | Gordonia alkanivorans |
| 526226 | Gordonia bronchialis DSM 43247 |
| 2420509 | Gordonia insulae |
| 1004901 | Gordonia iterans |
| 1136941 | Gordonia phthalatica |
| 1112204 | Gordonia polyisoprenivorans VH2 |
| 36822 | Gordonia rubripertincta |
| 337191 | Gordonia sp. KTR9 |
| 2059875 | Gordonia sp. YC-JH1 |
| 2055 | Gordonia terrae |
| 657308 | Gordonibacter pamelaeae 7-10-1-b |
| 9595 | Gorilla gorilla gorilla |
| 29729 | Gossypium arboreum |
| 3635 | Gossypium hirsutum |
| 29730 | Gossypium raimondii |
| 1128398 | Gottschalkia acidurici 9a |
| 744872 | Gracilinema caldarium DSM 7334 |
| 391165 | Granulibacter bethesdensis CGDNIH1 |
| 1051974 | Granulibacter bethesdensis CGDNIH2 |
| 1052004 | Granulibacter bethesdensis CGDNIH3 |
| 1052095 | Granulibacter bethesdensis CGDNIH4 |
| 682795 | Granulicella mallensis MP5ACTX8 |
| 1198114 | Granulicella tundricola MP5ACTX9 |
| 1192854 | Granulosicoccus antarcticus IMCC3135 |
| 673 | Grimontia hollisae |
| 2419771 | Gryllotalpicola protaetiae |
| 905079 | Guillardia theta CCMP2712 |
| 1445510 | Gynuella sunshinyii YC6258 |

| NCBI Taxon Identifier | Organism Name |
|---|---|
| 195105 | Haematobacter massiliensis |
| 1549855 | Haematospirillum jordaniae |
| 727 | Haemophilus influenzae |
| 727 | Haemophilus influenzae |
| 727 | Haemophilus influenzae |
| 727 | Haemophilus influenzae |
| 862964 | Haemophilus influenzae 10810 |
| 281310 | Haemophilus influenzae 86-028NP |
| 1295140 | Haemophilus influenzae CGSHiCZ412602 |
| 866630 | Haemophilus influenzae F3031 |
| 935897 | Haemophilus influenzae F3047 |
| 1334187 | Haemophilus influenzae KR494 |
| 374930 | Haemophilus influenzae PittEE |
| 374931 | Haemophilus influenzae PittGG |
| 262727 | Haemophilus influenzae R2846 |
| 262728 | Haemophilus influenzae R2866 |
| 71421 | Haemophilus influenzae Rd KW20 |
| 862965 | Haemophilus parainfluenzae T3T1 |
| 712310 | Haemophilus sp. oral taxon 036 |
| 1453496 | Hafnia alvei FB1 |
| 546367 | Hafnia paralvei |
| 349521 | Hahella chejuensis KCTC 2396 |
| 1628392 | Hahella sp. KA22 |
| 199441 | Halalkalibacter krulwichiae |
| 272558 | Halalkalibacterium halodurans C-125 |
| 795797 | Halalkalicoccus jeotgali B3 |
| 1604004 | Halanaeroarchaeum sulfurireducens |
| 1604004 | Halanaeroarchaeum sulfurireducens |
| 656519 | Halanaerobium hydrogeniformans |
| 572479 | Halanaerobium praevalens DSM 2228 |
| 502025 | Haliangium ochraceum DSM 14365 |
| 930805 | Halioglobus japonicus |
| 760192 | Haliscomenobacter hydrossis DSM 1100 |
| 634497 | Haloarcula hispanica ATCC 33960 |
| 1417673 | Haloarcula hispanica N601 |
| 272569 | Haloarcula marismortui ATCC 43049 |
| 1592728 | Haloarcula sp. CBA1115 |
| 1932004 | Haloarcula taiwanensis |
| 866895 | Halobacillus halophilus DSM 2266 |
| 45668 | Halobacillus litoralis |
| 402384 | Halobacillus mangrovi |

| NCBI Taxon Identifier | Organism Name |
|---|---|
| 862908 | Halobacteriovorax marinus SJ |
| 2109558 | Halobacteriovorax sp. BALOs_7 |
| 1407499 | Halobacterium hubeiense |
| 64091 | Halobacterium salinarum NRC-1 |
| 478009 | Halobacterium salinarum R1 |
| 751944 | Halobacterium sp. DL1 |
| 748449 | Halobacteroides halobius DSM 5150 |
| 358396 | Halobiforma lacisalsi AJ5 |
| 2382161 | Halocella sp. SP3-1 |
| 1679096 | Halococcoides cellulosivorans |
| 1873524 | Halodesulfurarchaeum formicicum |
| 1873524 | Halodesulfurarchaeum formicicum |
| 35746 | Haloferax gibbonsii |
| 523841 | Haloferax mediterranei ATCC 33500 |
| 309800 | Haloferax volcanii DS2 |
| 469382 | Halogeometricum borinquense DSM 11551 |
| 1073996 | Halohasta litchfieldiae |
| 485914 | Halomicrobium mukohataei DSM 12286 |
| 1641165 | Halomicronema hongdechloris C2206 |
| 1897729 | Halomonas aestuarii |
| 475662 | Halomonas beimenensis |
| 213554 | Halomonas campaniensis |
| 507626 | Halomonas chromatireducens |
| 768066 | Halomonas elongata DSM 2581 |
| 1178482 | Halomonas huangheensis |
| 115561 | Halomonas hydrothermalis |
| 1883416 | Halomonas sp. 1513 |
| 1118153 | Halomonas sp. GFAJ-1 |
| 1504981 | Halomonas sp. KO116 |
| 1610576 | Halomonas sp. R57-5 |
| 2136172 | Halomonas sp. SF2003 |
| 756883 | halophilic archaeon DL31 |
| 797210 | Halopiger xanaduensis SH-6 |
| 660522 | Haloplanus aerogenes |
| 1547898 | Haloplanus rubicundus |
| 1547898 | Haloplanus rubicundus |
| 768065 | Haloquadratum walsbyi C23 |
| 362976 | Haloquadratum walsbyi DSM 16790 |
| 1033806 | Halorhabdus tiamatea SARL4B |
| 519442 | Halorhabdus utahensis DSM 12940 |
| 1354791 | Halorhodospira halochloris str. A |

| NCBI Taxon Identifier | Organism Name |
|---|---|
| 349124 | Halorhodospira halophila SL1 |
| 1932360 | Halorientalis sp. IM1011 |
| 416348 | Halorubrum lacusprofundi ATCC 49239 |
| 2497325 | Halorubrum sp. BOL3-1 |
| 634157 | Halorubrum sp. PV6 |
| 797299 | Halostagnicola larsenii XH-48 |
| 376489 | Halotalea alkalilenta |
| 543526 | Haloterrigena turkmenica DSM 5511 |
| 65093 | Halothece sp. PCC 7418 |
| 373903 | Halothermothrix orenii H 168 |
| 1860122 | Halothiobacillus diazotrophicus |
| 555778 | Halothiobacillus neapolitanus c2 |
| 797302 | Halovivax ruber XH-70 |
| 610380 | Harpegnathos saltator |
| 1482074 | Hartmannibacter diazotrophicus |
| 4232 | Helianthus annuus |
| 382638 | Helicobacter acinonychis str. Sheeba |
| 135569 | Helicobacter apodemus |
| 37372 | Helicobacter bilis |
| 1002804 | Helicobacter bizzozeronii CIII-1 |
| 182217 | Helicobacter cetorum MIT 00-7128 |
| 1163745 | Helicobacter cetorum MIT 99-5656 |
| 537971 | Helicobacter cinaedi CCUG 18818 = ATCC BAA-847 |
| 1172562 | Helicobacter cinaedi PAGU611 |
| 936155 | Helicobacter felis ATCC 49179 |
| 1216962 | Helicobacter heilmannii ASB1.4 |
| 235279 | Helicobacter hepaticus ATCC 51449 |
| 679897 | Helicobacter mustelae 12198 |
| 985081 | Helicobacter pylori 2017 |
| 985080 | Helicobacter pylori 2018 |
| 85962 | Helicobacter pylori 26695 |
| 85962 | Helicobacter pylori 26695 |
| 585535 | Helicobacter pylori 35A |
| 290847 | Helicobacter pylori 51 |
| 684950 | Helicobacter pylori 52 |
| 585538 | Helicobacter pylori 83 |
| 869727 | Helicobacter pylori 908 |
| 1055531 | Helicobacter pylori Aklavik117 |
| 1055532 | Helicobacter pylori Aklavik86 |
| 592205 | Helicobacter pylori B38 |
| 693745 | Helicobacter pylori B8 |

| NCBI Taxon Identifier | Organism Name |
|---|---|
| 1407462 | Helicobacter pylori BM012A |
| 1407463 | Helicobacter pylori BM012S |
| 765964 | Helicobacter pylori Cuz20 |
| 1055527 | Helicobacter pylori ELS37 |
| 866344 | Helicobacter pylori F16 |
| 866345 | Helicobacter pylori F30 |
| 102608 | Helicobacter pylori F32 |
| 866346 | Helicobacter pylori F57 |
| 563041 | Helicobacter pylori G27 |
| 907240 | Helicobacter pylori Gambia94/24 |
| 357544 | Helicobacter pylori HPAG1 |
| 1163743 | Helicobacter pylori HUP-B14 |
| 907238 | Helicobacter pylori India7 |
| 85963 | Helicobacter pylori J99 |
| 907237 | Helicobacter pylori Lithuania75 |
| 1248725 | Helicobacter pylori OK113 |
| 1248726 | Helicobacter pylori OK310 |
| 1382920 | Helicobacter pylori oki102 |
| 1382921 | Helicobacter pylori oki112 |
| 1382922 | Helicobacter pylori oki128 |
| 1382923 | Helicobacter pylori oki154 |
| 1382924 | Helicobacter pylori oki422 |
| 1382925 | Helicobacter pylori oki673 |
| 1382926 | Helicobacter pylori oki828 |
| 1382927 | Helicobacter pylori oki898 |
| 570508 | Helicobacter pylori P12 |
| 1163742 | Helicobacter pylori PeCan18 |
| 765963 | Helicobacter pylori PeCan4 |
| 1055528 | Helicobacter pylori Puno120 |
| 1055529 | Helicobacter pylori Puno135 |
| 1234365 | Helicobacter pylori Rif1 |
| 1234600 | Helicobacter pylori Rif2 |
| 794851 | Helicobacter pylori Sat464 |
| 1163740 | Helicobacter pylori Shi112 |
| 1163741 | Helicobacter pylori Shi169 |
| 1163739 | Helicobacter pylori Shi417 |
| 512562 | Helicobacter pylori Shi470 |
| 765962 | Helicobacter pylori SJM180 |
| 1055530 | Helicobacter pylori SNT49 |
| 1352356 | Helicobacter pylori SouthAfrica20 |
| 907239 | Helicobacter pylori SouthAfrica7 |

| NCBI Taxon Identifier | Organism Name |
|---|---|
| 1311573 | Helicobacter pylori UM032 |
| 1321939 | Helicobacter pylori UM037 |
| 1321940 | Helicobacter pylori UM066 |
| 1321941 | Helicobacter pylori UM298 |
| 1321938 | Helicobacter pylori UM299 |
| 637913 | Helicobacter pylori v225d |
| 1127122 | Helicobacter pylori XZ274 |
| 76936 | Helicobacter typhlonius |
| 29058 | Helicoverpa armigera |
| 498761 | Heliomicrobium modesticaldum Ice1 |
| 6412 | Helobdella robusta |
| 1071380 | Henningerozyma blattae CBS 6284 |
| 1262470 | Herbaspirillum hiltneri N3 |
| 1078773 | Herbaspirillum rubrisubalbicans M1 |
| 964 | Herbaspirillum seropedicae |
| 757424 | Herbaspirillum seropedicae SmR1 |
| 2025949 | Herbaspirillum sp. meg3 |
| 1679721 | Herbinix luporum |
| 204773 | Herminiimonas arsenicoxydans |
| 316274 | Herpetosiphon aurantiacus DSM 785 |
| 747525 | Heterobasidion irregulare TC 32-1 |
| 10181 | Heterocephalus glaber |
| 3981 | Hevea brasiliensis |
| 941639 | Heyndrickxia coagulans 2-6 |
| 345219 | Heyndrickxia coagulans 36D1 |
| 1121088 | Heyndrickxia coagulans DSM 1 = ATCC 7050 |
| 760142 | Hippea maritima DSM 10411 |
| 109280 | Hippocampus comes |
| 186990 | Hipposideros armiger |
| 582402 | Hirschia baltica ATCC 49814 |
| 205914 | Histophilus somni 129PT |
| 228400 | Histophilus somni 2336 |
| 2059318 | Histoplasma mississippiense (nom. inval.) |
| 1620421 | Hoeflea sp. IMCC20628 |
| 9606 | Homo sapiens |
| 76123 | Hoylesella enoeca |
| 443218 | Hoyosella subflava DQS3-9A1 |
| 2282656 | Humibacter sp. BT305 |
| 1834198 | Hungateiclostridiaceae bacterium KB18 |
| 6087 | Hydra vulgaris |
| 2231112 | Hydrogenimonas sp. |

| NCBI Taxon Identifier | Organism Name |
|---|---|
| 608538 | Hydrogenobacter thermophilus TK-6 |
| 608538 | Hydrogenobacter thermophilus TK-6 |
| 547144 | Hydrogenobaculum sp. HO |
| 547146 | Hydrogenobaculum sp. SN |
| 380749 | Hydrogenobaculum sp. Y04AAS1 |
| 1763535 | Hydrogenophaga crassostreae |
| 795665 | Hydrogenophaga sp. PBC |
| 1842537 | Hydrogenophaga sp. RAC07 |
| 297 | Hydrogenophilus thermoluteolus |
| 317025 | Hydrogenovibrio crunogenus XCL-2 |
| 265883 | Hydrogenovibrio thermophilus |
| 1850093 | Hymenobacter nivis |
| 2319843 | Hymenobacter oligotrophus |
| 1484116 | Hymenobacter psoromatis |
| 1411621 | Hymenobacter sedentarius |
| 1356852 | Hymenobacter sp. APR13 |
| 1385663 | Hymenobacter sp. DG25A |
| 1385664 | Hymenobacter sp. DG25B |
| 1484118 | Hymenobacter sp. PAMC 26628 |
| 1227739 | Hymenobacter swuensis DY53 |
| 415426 | Hyperthermus butylicus DSM 5456 |
| 670307 | Hyphomicrobium denitrificans 1NES1 |
| 582899 | Hyphomicrobium denitrificans ATCC 51888 |
| 1029756 | Hyphomicrobium nitrativorans NL23 |
| 717785 | Hyphomicrobium sp. MC1 |
| 1690484 | Hyphomonadaceae bacterium UKL13-1 |
| 228405 | Hyphomonas neptunium ATCC 15444 |
| 242600 | Ichthyobacterium seriolicida |
| 7998 | Ictalurus punctatus |
| 1321370 | Idiomarina loihiensis GSL 199 |
| 283942 | Idiomarina loihiensis L2TR |
| 1096243 | Idiomarina piscisalsi |
| 2100422 | Idiomarina sp. OT37-5b |
| 2055892 | Idiomarina sp. X4 |
| 945713 | Ignavibacterium album JCM 16511 |
| 453591 | Ignicoccus hospitalis KIN4/I |
| 940295 | Ignicoccus islandicus DSM 13165 |
| 583356 | Ignisphaera aggregans DSM 17230 |
| 1313172 | Ilumatobacter coccineus YM16-304 |
| 572544 | Ilyobacter polytropus DSM 2926 |
| 1810504 | Immundisolibacter cernigliae |

| NCBI Taxon Identifier | Organism Name |
|---|---|
| 2487118 | Intestinibaculum porci |
| 1297617 | Intestinimonas butyriciproducens |
| 710696 | Intrasporangium calvum DSM 43043 |
| 2496266 | Iodobacter ciconiae |
| 35883 | Ipomoea nil |
| 391936 | Isoalcanivorax pacificus W11-5 |
| 1300344 | Isoptericola dokdonensis DS-3 |
| 743718 | Isoptericola variabilis 225 |
| 575540 | Isosphaera pallida ATCC 43644 |
| 6945 | Ixodes scapularis |
| 857417 | Janibacter indicus |
| 53458 | Janibacter limosus |
| 290400 | Jannaschia sp. CCS1 |
| 1349767 | Janthinobacterium agaricidamnosum NBRC 102515 = DSM 9628 |
| 1644131 | Janthinobacterium sp. 1_2014MBL_MicDiv |
| 2497863 | Janthinobacterium sp. 17J80-10 |
| 1236179 | Janthinobacterium sp. B9-8 |
| 1938606 | Janthinobacterium sp. LM6 |
| 375286 | Janthinobacterium sp. Marseille |
| 368607 | Janthinobacterium svalbardensis |
| 180498 | Jatropha curcas |
| 1906741 | Jeongeupia sp. USM3 |
| 2496265 | Jeotgalibaca ciconiae |
| 708126 | Jeotgalibaca dankookensis |
| 1903686 | Jeotgalibaca sp. PTS2502 |
| 1508404 | Jeotgalibacillus malaysiensis |
| 471856 | Jonesia denitrificans DSM 20603 |
| 51240 | Juglans regia |
| 914150 | Kangiella geojedonensis |
| 523791 | Kangiella koreensis DSM 16069 |
| 1561924 | Kangiella profundi |
| 1144748 | Kangiella sediminilitoris |
| 1071382 | Kazachstania africana CBS 2517 |
| 1917421 | Ketobacter alkanivorans |
| 92947 | Ketogulonicigenium robustum |
| 759362 | Ketogulonicigenium vulgare WSH-001 |
| 880591 | Ketogulonicigenium vulgare Y25 |
| 860235 | Kibdelosporangium phytohabitans |
| 266940 | Kineococcus radiotolerans SRS30216 = ATCC BAA-149 |
| 504 | Kingella kingae |
| 1307763 | Kiritimatiella glycovorans |

| NCBI Taxon Identifier | Organism Name |
|---|---|
| 68173 | Kitasatospora albolonga |
| 1894 | Kitasatospora aureofaciens |
| 452652 | Kitasatospora setae KM-6054 |
| 2018025 | Kitasatospora sp. MMS16-BH015 |
| 935296 | Klebsiella aerogenes EA1509E |
| 1028307 | Klebsiella aerogenes KCTC 2190 |
| 1134687 | Klebsiella michiganensis |
| 1134687 | Klebsiella michiganensis |
| 1191061 | Klebsiella michiganensis E718 |
| 1308980 | Klebsiella michiganensis HKOPL1 |
| 1006551 | Klebsiella michiganensis KCTC 1686 |
| 571 | Klebsiella oxytoca |
| 1333852 | Klebsiella oxytoca KONIH1 |
| 573 | Klebsiella pneumoniae |
| 573 | Klebsiella pneumoniae |
| 573 | Klebsiella pneumoniae |
| 573 | Klebsiella pneumoniae |
| 573 | Klebsiella pneumoniae |
| 1420012 | Klebsiella pneumoniae 30660/NJST258_1 |
| 1420013 | Klebsiella pneumoniae 30684/NJST258_2 |
| 507522 | Klebsiella pneumoniae 342 |
| 1244085 | Klebsiella pneumoniae CG43 |
| 1380908 | Klebsiella pneumoniae JM45 |
| 1049565 | Klebsiella pneumoniae KCTC 2242 |
| 72407 | Klebsiella pneumoniae subsp. pneumoniae |
| 72407 | Klebsiella pneumoniae subsp. pneumoniae |
| 72407 | Klebsiella pneumoniae subsp. pneumoniae |
| 72407 | Klebsiella pneumoniae subsp. pneumoniae |
| 72407 | Klebsiella pneumoniae subsp. pneumoniae |
| 1193292 | Klebsiella pneumoniae subsp. pneumoniae 1084 |
| 1125630 | Klebsiella pneumoniae subsp. pneumoniae HS11286 |
| 1094170 | Klebsiella pneumoniae subsp. pneumoniae KPNIH10 |
| 1225181 | Klebsiella pneumoniae subsp. pneumoniae KPNIH24 |
| 1328324 | Klebsiella pneumoniae subsp. pneumoniae KPNIH27 |
| 1328325 | Klebsiella pneumoniae subsp. pneumoniae KPR0928 |
| 272620 | Klebsiella pneumoniae subsp. pneumoniae MGH 78578 |
| 484021 | Klebsiella pneumoniae subsp. pneumoniae NTUH-K2044 |
| 861365 | Klebsiella pneumoniae subsp. rhinoscleromatis SB3432 |
| 1463165 | Klebsiella quasipneumoniae |
| 2026240 | Klebsiella quasivariicola |
| 1905288 | Klebsiella sp. LTGPAF-6F |

| NCBI Taxon Identifier | Organism Name |
|---|---|
| 244366 | Klebsiella variicola |
| 244366 | Klebsiella variicola |
| 244366 | Klebsiella variicola |
| 640131 | Klebsiella variicola At-22 |
| 73098 | Kluyvera georgiana |
| 284590 | Kluyveromyces lactis NRRL Y-1140 |
| 1003335 | Kluyveromyces marxianus DMKU3-1042 |
| 446860 | Kocuria flava |
| 1049583 | Kocuria indica |
| 71999 | Kocuria palustris |
| 378753 | Kocuria rhizophila DC2201 |
| 33995 | Komagataeibacter europaeus |
| 634177 | Komagataeibacter medellinensis NBRC 3288 |
| 265960 | Komagataeibacter nataicola |
| 265959 | Komagataeibacter saccharivorans |
| 1296990 | Komagataeibacter xylinus E25 |
| 644223 | Komagataella phaffii GS115 |
| 2282170 | Kordia sp. SMS9 |
| 208223 | Kosakonia cowanii |
| 497725 | Kosakonia oryzae |
| 283686 | Kosakonia radicincitans |
| 1235834 | Kosakonia sacchari SP1 |
| 2492396 | Kosakonia sp. CCTCC M2018092 |
| 521045 | Kosmotoga olearia TBF 19.5.1 |
| 1330330 | Kosmotoga pacifica |
| 153496 | Kozakia baliensis |
| 479435 | Kribbella flavida DSM 17836 |
| 37003 | Kryptolebias marmoratus |
| 1750719 | Kurthia sp. 11kri321 |
| 698828 | Kushneria konosiri |
| 157779 | Kushneria marisflavi |
| 1449976 | Kutzneria albida DSM 43870 |
| 2055160 | Kyrpidia spormannii |
| 562970 | Kyrpidia tusciae DSM 2912 |
| 478801 | Kytococcus sedentarius DSM 20547 |
| 1717717 | Labilibaculum antarcticum |
| 1391654 | Labilithrix luteola |
| 1674922 | Labrenzia sp. CP4 |
| 2021862 | Labrenzia sp. VG12 |
| 486041 | Laccaria bicolor S238N-H82 |
| 37482 | Laceyella sacchari |

| NCBI Taxon Identifier | Organism Name |
|---|---|
| 559295 | Lachancea thermotolerans CBS 6340 |
| 617123 | Lachnoanaerobaculum umeaense |
| 357809 | Lachnoclostridium phytofermentans ISDg |
| 1834196 | Lachnoclostridium sp. YL32 |
| 712991 | Lachnospiraceae bacterium oral taxon 500 |
| 1526571 | Lacimicrobium alkaliphilum |
| 983544 | Lacinutrix sp. 5H-3-7-4 |
| 2057808 | Lacinutrix sp. Bg11-31 |
| 1486034 | Lacinutrix venerupis |
| 1051650 | Lacticaseibacillus casei 12A |
| 998820 | Lacticaseibacillus casei BD-II |
| 543734 | Lacticaseibacillus casei BL23 |
| 999378 | Lacticaseibacillus casei LC2W |
| 1318635 | Lacticaseibacillus casei LOCK919 |
| 1215914 | Lacticaseibacillus casei W56 |
| 1597 | Lacticaseibacillus paracasei |
| 321967 | Lacticaseibacillus paracasei ATCC 334 |
| 1446494 | Lacticaseibacillus paracasei N1115 |
| 537973 | Lacticaseibacillus paracasei subsp. paracasei 8700:2 |
| 525337 | Lacticaseibacillus paracasei subsp. paracasei ATCC 25302 = DSM 5622 = JCM 8130 |
| 1088720 | Lacticaseibacillus rhamnosus ATCC 8530 |
| 568703 | Lacticaseibacillus rhamnosus GG |
| 568703 | Lacticaseibacillus rhamnosus GG |
| 568704 | Lacticaseibacillus rhamnosus Lc 705 |
| 1316933 | Lacticaseibacillus rhamnosus LOCK900 |
| 1318634 | Lacticaseibacillus rhamnosus LOCK908 |
| 60520 | Lactiplantibacillus paraplantarum |
| 1589 | Lactiplantibacillus pentosus |
| 1590 | Lactiplantibacillus plantarum |
| 1327988 | Lactiplantibacillus plantarum 16 |
| 644042 | Lactiplantibacillus plantarum JDM1 |
| 889932 | Lactiplantibacillus plantarum ST-III |
| 767468 | Lactiplantibacillus plantarum subsp. plantarum P-8 |
| 220668 | Lactiplantibacillus plantarum WCFS1 |
| 1284663 | Lactiplantibacillus plantarum ZJ316 |
| 1600 | Lactobacillus acetotolerans |
| 1579 | Lactobacillus acidophilus |
| 891391 | Lactobacillus acidophilus 30SC |
| 1314884 | Lactobacillus acidophilus La-14 |
| 272621 | Lactobacillus acidophilus NCFM |
| 83683 | Lactobacillus amylolyticus |

| NCBI Taxon Identifier | Organism Name |
|---|---|
| 695560 | Lactobacillus amylovorus GRL 1112 |
| 695562 | Lactobacillus amylovorus GRL1118 |
| 748671 | Lactobacillus crispatus ST1 |
| 353496 | Lactobacillus delbrueckii subsp. bulgaricus 2038 |
| 390333 | Lactobacillus delbrueckii subsp. bulgaricus ATCC 11842 = JCM 1002 |
| 321956 | Lactobacillus delbrueckii subsp. bulgaricus ATCC BAA-365 |
| 767455 | Lactobacillus delbrueckii subsp. bulgaricus ND02 |
| 52242 | Lactobacillus gallinarum |
| 324831 | Lactobacillus gasseri ATCC 33323 = JCM 1131 |
| 1587 | Lactobacillus helveticus |
| 326425 | Lactobacillus helveticus CNRZ32 |
| 405566 | Lactobacillus helveticus DPC 4571 |
| 767462 | Lactobacillus helveticus H10 |
| 767456 | Lactobacillus helveticus H9 |
| 880633 | Lactobacillus helveticus R0052 |
| 109790 | Lactobacillus jensenii |
| 909954 | Lactobacillus johnsonii DPC 6026 |
| 633699 | Lactobacillus johnsonii FI9785 |
| 1408186 | Lactobacillus johnsonii N6.2 |
| 257314 | Lactobacillus johnsonii NCC 533 |
| 1033837 | Lactobacillus kefiranofaciens ZW3 |
| 2107999 | Lactobacillus paragasseri |
| 1545702 | Lactobacillus sp. wkB8 |
| 2419773 | Lactococcus allomyrinae |
| 1104322 | Lactococcus cremoris subsp. cremoris A76 |
| 1295826 | Lactococcus cremoris subsp. cremoris KW2 |
| 416870 | Lactococcus cremoris subsp. cremoris MG1363 |
| 746361 | Lactococcus cremoris subsp. cremoris NZ9000 |
| 272622 | Lactococcus cremoris subsp. cremoris SK11 |
| 1111678 | Lactococcus cremoris subsp. cremoris UC509.9 |
| 420889 | Lactococcus garvieae ATCC 49156 |
| 420890 | Lactococcus garvieae Lg2 |
| 1358 | Lactococcus lactis |
| 929102 | Lactococcus lactis subsp. lactis CV56 |
| 272623 | Lactococcus lactis subsp. lactis Il1403 |
| 1046624 | Lactococcus lactis subsp. lactis IO-1 |
| 684738 | Lactococcus lactis subsp. lactis KF147 |
| 1399116 | Lactococcus lactis subsp. lactis KLDS 4.0325 |
| 1117941 | Lactococcus lactis subsp. lactis NCDO 2118 |
| 297352 | Lactococcus piscium MKFS47 |
| 1366 | Lactococcus raffinolactis |

| NCBI Taxon Identifier | Organism Name |
|---|---|
| 4236 | Lactuca sativa |
| 1838286 | Lacunisphaera limnophila |
| 521095 | Lancefieldella parvula DSM 20469 |
| 557598 | Laribacter hongkongensis HLHK9 |
| 215358 | Larimichthys crocea |
| 8187 | Lates calcarifer |
| 28038 | Latilactobacillus curvatus |
| 314315 | Latilactobacillus sakei subsp. sakei 23K |
| 7897 | Latimeria chalumnae |
| 47671 | Lautropia mirabilis |
| 1528099 | Lawsonella clevelandensis |
| 1234378 | Lawsonia intracellularis N343 |
| 363253 | Lawsonia intracellularis PHE/MN1-00 |
| 545695 | Leadbettera azotonutricia ZAS-9 |
| 649349 | Leadbetterella byssophila DSM 17132 |
| 83655 | Leclercia adecarboxylata |
| 1920114 | Leclercia sp. LSNIH1 |
| 1920116 | Leclercia sp. LSNIH3 |
| 1867846 | Legionella clemsonensis |
| 1212491 | Legionella fallonii LLAP-10 |
| 449 | Legionella hackeliae |
| 45067 | Legionella lansingensis |
| 661367 | Legionella longbeachae NSW150 |
| 451 | Legionella micdadei |
| 1268635 | Legionella oakridgensis ATCC 33761 = DSM 21215 |
| 423212 | Legionella pneumophila 2300/99 Alcoy |
| 400673 | Legionella pneumophila str. Corby |
| 297245 | Legionella pneumophila str. Lens |
| 297246 | Legionella pneumophila str. Paris |
| 91891 | Legionella pneumophila subsp. pneumophila |
| 91891 | Legionella pneumophila subsp. pneumophila |
| 933093 | Legionella pneumophila subsp. pneumophila ATCC 43290 |
| 1312904 | Legionella pneumophila subsp. pneumophila LPE509 |
| 272624 | Legionella pneumophila subsp. pneumophila str. Philadelphia 1 |
| 1199191 | Legionella pneumophila subsp. pneumophila str. Thunder Bay |
| 28087 | Legionella sainthelensi |
| 1389489 | Leifsonia xyli subsp. cynodontis DSM 46306 |
| 281090 | Leifsonia xyli subsp. xyli str. CTCB07 |
| 420245 | Leishmania braziliensis MHOM/BR/75/M2904 |
| 5661 | Leishmania donovani |
| 435258 | Leishmania infantum JPCM5 |

| NCBI Taxon Identifier | Organism Name |
|---|---|
| 347515 | Leishmania major strain Friedlin |
| 929439 | Leishmania mexicana MHOM/GT/2001/U1103 |
| 999552 | Leisingera methylohalidivorans DSM 14336 |
| 2508307 | Leisingera sp. NJS204 |
| 61646 | Lelliottia amnigena |
| 1907578 | Lelliottia jeotgali |
| 69220 | Lelliottia nimipressuralis |
| 2153385 | Lelliottia sp. WB101 |
| 158841 | Leminorella richardii |
| 1472767 | Lentibacillus amyloliquefaciens |
| 511437 | Lentilactobacillus buchneri NRRL B-30929 |
| 1071400 | Lentilactobacillus buchneri subsp. silagei CD034 |
| 1138822 | Lentilactobacillus curieae |
| 152331 | Lentilactobacillus parabuchneri |
| 1586287 | Lentzea guizhouensis |
| 411966 | Leptolyngbya boryana IAM M-101 |
| 1752064 | Leptolyngbya sp. NIES-3755 |
| 1080068 | Leptolyngbya sp. O-77 |
| 28452 | Leptospira alstonii |
| 355278 | Leptospira biflexa serovar Patoc strain 'Patoc 1 (Ames)' |
| 456481 | Leptospira biflexa serovar Patoc strain 'Patoc 1 (Paris)' |
| 355277 | Leptospira borgpetersenii serovar Hardjo-bovis str. JB197 |
| 355276 | Leptospira borgpetersenii serovar Hardjo-bovis str. L550 |
| 267671 | Leptospira interrogans serovar Copenhageni str. Fiocruz L1-130 |
| 189518 | Leptospira interrogans serovar Lai str. 56601 |
| 573825 | Leptospira interrogans serovar Lai str. IPAV |
| 1395589 | Leptospira interrogans serovar Linhai str. 56609 |
| 408139 | Leptospira kmetyi |
| 1137606 | Leptospira mayottensis |
| 758847 | Leptospira santarosai serovar Shermani str. LT 821 |
| 1048260 | Leptospirillum ferriphilum ML-04 |
| 1441628 | Leptospirillum ferriphilum YSK |
| 1162668 | Leptospirillum ferrooxidans C2-3 |
| 1660083 | Leptospirillum sp. Group II 'CF-1' |
| 395495 | Leptothrix cholodnii SP-6 |
| 523794 | Leptotrichia buccalis C-1013-b |
| 712357 | Leptotrichia sp. oral taxon 212 |
| 712368 | Leptotrichia sp. oral taxon 498 |
| 1785996 | Leptotrichia sp. oral taxon 847 |
| 1935379 | Leucobacter muris |
| 1784719 | Leucobacter triazinivorans |

| NCBI Taxon Identifier | Organism Name |
|---|---|
| 1229758 | Leuconostoc carnosum JB16 |
| 349519 | Leuconostoc citreum KM20 |
| 255248 | Leuconostoc garlicum |
| 762550 | Leuconostoc gasicomitatum LMG 18811 |
| 1229756 | Leuconostoc gelidum JB7 |
| 762051 | Leuconostoc kimchii IMSNU 11154 |
| 1246 | Leuconostoc lactis |
| 427140 | Leuconostoc mesenteroides KFRI-MG |
| 203120 | Leuconostoc mesenteroides subsp. mesenteroides ATCC 8293 |
| 1107880 | Leuconostoc mesenteroides subsp. mesenteroides J18 |
| 979982 | Leuconostoc sp. C2 |
| 1511761 | Leuconostoc suionicum |
| 387344 | Levilactobacillus brevis ATCC 367 |
| 1001583 | Levilactobacillus brevis KB290 |
| 637971 | Levilactobacillus koreensis |
| 267363 | Levilactobacillus zymae |
| 1215343 | Liberibacter crescens BT-1 |
| 89059 | Ligilactobacillus acidipiscis |
| 1601 | Ligilactobacillus agilis |
| 1069534 | Ligilactobacillus ruminis ATCC 27782 |
| 1624 | Ligilactobacillus salivarius |
| 712961 | Ligilactobacillus salivarius CECT 5713 |
| 362948 | Ligilactobacillus salivarius UCC118 |
| 1851148 | Limihaloglobus sulfuriphilus |
| 2172103 | Limnobaculum parvum |
| 1555112 | Limnochorda pilosa |
| 1678129 | Limnohabitans sp. 103DPR2 |
| 1678128 | Limnohabitans sp. 63ED37-2 |
| 712938 | Limosilactobacillus fermentum CECT 5716 |
| 767453 | Limosilactobacillus fermentum F-6 |
| 334390 | Limosilactobacillus fermentum IFO 3956 |
| 1130798 | Limosilactobacillus mucosae LM1 |
| 1632 | Limosilactobacillus oris |
| 1340495 | Limosilactobacillus reuteri I5007 |
| 491077 | Limosilactobacillus reuteri SD2112 |
| 557436 | Limosilactobacillus reuteri subsp. reuteri |
| 557433 | Limosilactobacillus reuteri subsp. reuteri JCM 1112 |
| 1358027 | Limosilactobacillus reuteri TD1 |
| 83485 | Linepithema humile |
| 7574 | Lingula anatina |
| 118797 | Lipotes vexillifer |

| NCBI Taxon Identifier | Organism Name |
|---|---|
| 272626 | Listeria innocua Clip11262 |
| 202751 | Listeria ivanovii subsp. ivanovii |
| 881621 | Listeria ivanovii subsp. ivanovii PAM 55 |
| 202752 | Listeria ivanovii subsp. londoniensis |
| 202752 | Listeria ivanovii subsp. londoniensis |
| 1457190 | Listeria ivanovii WSLC3009 |
| 1639 | Listeria monocytogenes |
| 1639 | Listeria monocytogenes |
| 1639 | Listeria monocytogenes |
| 1126011 | Listeria monocytogenes 07PF0776 |
| 653938 | Listeria monocytogenes 08-5578 |
| 637381 | Listeria monocytogenes 08-5923 |
| 393133 | Listeria monocytogenes 10403S |
| 1288295 | Listeria monocytogenes 6179 |
| 882095 | Listeria monocytogenes ATCC 19117 |
| 1334565 | Listeria monocytogenes EGD |
| 169963 | Listeria monocytogenes EGD-e |
| 393127 | Listeria monocytogenes Finland 1998 |
| 393126 | Listeria monocytogenes FSL R2-561 |
| 552536 | Listeria monocytogenes HCC23 |
| 393130 | Listeria monocytogenes J0161 |
| 930782 | Listeria monocytogenes J1-220 |
| 930781 | Listeria monocytogenes J1816 |
| 882094 | Listeria monocytogenes L312 |
| 563174 | Listeria monocytogenes L99 |
| 1234141 | Listeria monocytogenes La111 |
| 1030009 | Listeria monocytogenes M7 |
| 1234142 | Listeria monocytogenes N53-1 |
| 1437838 | Listeria monocytogenes R479a |
| 568819 | Listeria monocytogenes serotype 4b str. CLIP 80459 |
| 265669 | Listeria monocytogenes serotype 4b str. F2365 |
| 1230340 | Listeria monocytogenes serotype 4b str. LL195 |
| 863767 | Listeria monocytogenes serotype 7 str. SLCC2482 |
| 932920 | Listeria monocytogenes SLCC2372 |
| 882097 | Listeria monocytogenes SLCC2376 |
| 879088 | Listeria monocytogenes SLCC2378 |
| 882020 | Listeria monocytogenes SLCC2479 |
| 879089 | Listeria monocytogenes SLCC2540 |
| 932919 | Listeria monocytogenes SLCC2755 |
| 882096 | Listeria monocytogenes SLCC5850 |
| 879090 | Listeria monocytogenes SLCC7179 |

| NCBI Taxon Identifier | Organism Name |
|---|---|
| 1457187 | Listeria monocytogenes WSLC1001 |
| 1457188 | Listeria monocytogenes WSLC1042 |
| 683837 | Listeria seeligeri serovar 1/2b str. SLCC3954 |
| 1006155 | Listeria weihenstephanensis |
| 386043 | Listeria welshimeri serovar 6b str. SLCC5334 |
| 718192 | Litorilituus sediminis |
| 7209 | Loa loa |
| 379508 | Lodderomyces elongisporus NRRL YB-4239 |
| 375175 | Loigolactobacillus backii |
| 1423822 | Loigolactobacillus coryniformis subsp. torquens DSM 20004 = KCTC 3535 |
| 1082704 | Lonsdalea britannica |
| 225164 | Lottia gigantea |
| 34305 | Lotus japonicus |
| 9785 | Loxodonta africana |
| 3871 | Lupinus angustifolius |
| 1440763 | Luteibacter rhizovicinus DSM 16549 |
| 2006110 | Luteimonas chenhongjianii |
| 571913 | Luteipulveratus mongoliensis |
| 1855912 | Luteitalea pratensis |
| 1622118 | Lutibacter profundi |
| 28031 | Lysinibacillus fusiformis |
| 2169540 | Lysinibacillus sp. 2017 |
| 2081964 | Lysinibacillus sp. B2A1 |
| 2072025 | Lysinibacillus sp. YS11 |
| 444177 | Lysinibacillus sphaericus C3-41 |
| 1145276 | Lysinibacillus varians |
| 84531 | Lysobacter antibioticus |
| 84531 | Lysobacter antibioticus |
| 435897 | Lysobacter capsici |
| 69 | Lysobacter enzymogenes |
| 69 | Lysobacter enzymogenes |
| 262324 | Lysobacter gummosus |
| 1605891 | Lysobacter maris |
| 2698682 | Lysobacter oculi |
| 2290922 | Lysobacter sp. TY2-98 |
| 9541 | Macaca fascicularis |
| 9544 | Macaca mulatta |
| 1855823 | Macrococcoides canis |
| 458233 | Macrococcoides caseolyticum JCSC5402 |
| 1898474 | Macrococcus sp. IME1552 |
| 29159 | Magallana gigas |

| NCBI Taxon Identifier | Organism Name |
|---|---|
| 699246 | Mageeibacillus indolicus UPII9-5 |
| 156889 | Magnetococcus marinus MC-1 |
| 1288970 | Magnetospira sp. QH-2 |
| 431944 | Magnetospirillum gryphiswaldense MSR-1 |
| 1430440 | Magnetospirillum gryphiswaldense MSR-1 v2 |
| 1639348 | Magnetospirillum sp. ME-1 |
| 1663591 | Magnetospirillum sp. XM-1 |
| 697281 | Mahella australiensis 50-1 BON |
| 202462 | Maize bushy stunt phytoplasma |
| 197482 | Malaciobacter halophilus |
| 1032238 | Malaciobacter mytili LMG 24559 |
| 272633 | Malacoplasma penetrans HF-2 |
| 425265 | Malassezia globosa CBS 7966 |
| 3750 | Malus domestica |
| 1296 | Mammaliicoccus sciuri |
| 3983 | Manihot esculenta |
| 75985 | Mannheimia haemolytica |
| 75985 | Mannheimia haemolytica |
| 1261126 | Mannheimia haemolytica D153 |
| 1311759 | Mannheimia haemolytica D171 |
| 1311760 | Mannheimia haemolytica D174 |
| 1316932 | Mannheimia haemolytica M42548 |
| 1249531 | Mannheimia haemolytica USDA-ARS-USMARC-183 |
| 1222034 | Mannheimia haemolytica USDA-ARS-USMARC-184 |
| 1249526 | Mannheimia haemolytica USDA-ARS-USMARC-185 |
| 1366053 | Mannheimia haemolytica USMARC_2286 |
| 1432056 | Mannheimia sp. USDA-ARS-USMARC-1261 |
| 1433287 | Mannheimia varigena USDA-ARS-USMARC-1296 |
| 1434214 | Mannheimia varigena USDA-ARS-USMARC-1312 |
| 1434215 | Mannheimia varigena USDA-ARS-USMARC-1388 |
| 1178778 | Maribacter cobaltidurans |
| 1836467 | Maribacter hydrothermalis |
| 1644130 | Maribacter sp. 1_2014MBL_MicDiv |
| 313603 | Maribacter sp. HTCC2170 |
| 2496865 | Maribacter sp. MJ134 |
| 394221 | Maricaulis maris MCS10 |
| 765910 | Marichromatium purpuratum 984 |
| 1121451 | Maridesulfovibrio hydrothermalis AM13 = DSM 14728 |
| 526222 | Maridesulfovibrio salexigens DSM 2638 |
| 2027857 | Mariniflexile sp. TRM1-10 |
| 1911586 | Marinilactibacillus sp. 15R |

| NCBI Taxon Identifier | Organism Name |
|---|---|
| 869210 | Marinithermus hydrothermalis DSM 14884 |
| 443254 | Marinitoga piezophila KA3 |
| 1545835 | Marinitoga sp. 1137 |
| 225937 | Marinobacter adhaerens HP15 |
| 1163748 | Marinobacter nauticus ATCC 49840 |
| 351348 | Marinobacter nauticus VT8 |
| 330734 | Marinobacter psychrophilus |
| 1420917 | Marinobacter salarius |
| 1874317 | Marinobacter salinus |
| 1420916 | Marinobacter similis |
| 2304594 | Marinobacter sp. Arc7-DN-1 |
| 490759 | Marinobacter sp. BSs20148 |
| 1671721 | Marinobacter sp. CP1 |
| 1749259 | Marinobacter sp. LQ44 |
| 1821621 | Marinobacterium aestuarii |
| 717774 | Marinomonas mediterranea MMB-1 |
| 491952 | Marinomonas posidonica IVIA-Po-181 |
| 400668 | Marinomonas sp. MWYL1 |
| 988812 | Marinovum algicola DG 898 |
| 1921086 | Mariprofundus aestuarium |
| 1921087 | Mariprofundus ferrinatatus |
| 454601 | Maritalea myrionectae |
| 643867 | Marivirga tractuosa DSM 4126 |
| 1920883 | Marivivens sp. JLT3646 |
| 1486262 | Martelella endophytica |
| 1122214 | Martelella mediterranea DSM 17316 |
| 686597 | Martelella sp. AD-3 |
| 945844 | Massilia oculi |
| 1678028 | Massilia sp. NR 4-1 |
| 1707785 | Massilia sp. WG5 |
| 1593482 | Massilia sp. YMA4 |
| 2045208 | Massilia violaceinigra |
| 106582 | Maylandia zebra |
| 3880 | Medicago truncatula |
| 1121448 | Megalodesulfovibrio gigas DSM 1382 = ATCC 19364 |
| 657316 | Megamonas hypermegale ART12/1 |
| 1064535 | Megasphaera elsdenii DSM 20460 |
| 1675036 | Megasphaera hexanoica |
| 2144175 | Megasphaera stantonii |
| 504728 | Meiothermus ruber DSM 1279 |
| 504728 | Meiothermus ruber DSM 1279 |

| NCBI Taxon Identifier | Organism Name |
|---|---|
| 1339250 | Meiothermus taiwanensis WR-220 |
| 2109913 | Melaminivora suipulveris |
| 747676 | Melampsora larici-populina 98AG31 |
| 9103 | Meleagris gallopavo |
| 1191523 | Melioribacter roseus P3M-2 |
| 940190 | Melissococcus plutonius ATCC 35311 |
| 1090974 | Melissococcus plutonius DAT561 |
| 1294270 | Melittangium boletus DSM 14713 |
| 743966 | Mesomycoplasma bovoculi M165/69 |
| 572263 | Mesomycoplasma conjunctivae HRC/581 |
| 86660 | Mesomycoplasma dispar |
| 743971 | Mesomycoplasma flocculare ATCC 27399 |
| 907287 | Mesomycoplasma hyopneumoniae 168 |
| 1116211 | Mesomycoplasma hyopneumoniae 168-L |
| 295358 | Mesomycoplasma hyopneumoniae 232 |
| 754503 | Mesomycoplasma hyopneumoniae 7422 |
| 262722 | Mesomycoplasma hyopneumoniae 7448 |
| 262719 | Mesomycoplasma hyopneumoniae J |
| 634997 | Mesomycoplasma hyorhinis DBS 1050 |
| 1129369 | Mesomycoplasma hyorhinis GDL-1 |
| 872331 | Mesomycoplasma hyorhinis HUB-1 |
| 936139 | Mesomycoplasma hyorhinis MCLD |
| 1118964 | Mesomycoplasma hyorhinis SK76 |
| 216427 | Mesoplasma chauliocola |
| 324078 | Mesoplasma coleopterae |
| 2149 | Mesoplasma entomophilum |
| 265311 | Mesoplasma florum L1 |
| 1406864 | Mesoplasma florum W37 |
| 81460 | Mesoplasma lactucae ATCC 49193 |
| 81459 | Mesoplasma melaleucae |
| 225999 | Mesoplasma syrphidae |
| 219745 | Mesoplasma tabanidae |
| 1082933 | Mesorhizobium amorphae CCNWGS0123 |
| 754035 | Mesorhizobium australicum WSM2073 |
| 765698 | Mesorhizobium ciceri biovar biserrulae WSM1271 |
| 266835 | Mesorhizobium japonicum MAFF 303099 |
| 935546 | Mesorhizobium loti NZP2037 |
| 536019 | Mesorhizobium opportunistum WSM2075 |
| 278153 | Mesorhizobium sp. WSM1497 |
| 1236046 | Mesotoga infera |
| 660470 | Mesotoga prima MesG1.Ag.4.2 |

| NCBI Taxon Identifier | Organism Name |
|---|---|
| 1006006 | Metallosphaera cuprina Ar-4 |
| 1293036 | Metallosphaera hakonensis JCM 8857 = DSM 7519 |
| 399549 | Metallosphaera sedula DSM 5348 |
| 243272 | Metamycoplasma arthritidis 158L3-1 |
| 29554 | Metamycoplasma canadense |
| 347256 | Metamycoplasma hominis ATCC 23114 |
| 1267000 | Metamycoplasma hominis ATCC 27545 |
| 29559 | Metamycoplasma hyosynoviae |
| 655827 | Metarhizium acridum CQMa 102 |
| 655844 | Metarhizium robertsii ARSEF 23 |
| 118062 | Methanobacterium congolense |
| 2162 | Methanobacterium formicicum |
| 2162 | Methanobacterium formicicum |
| 877455 | Methanobacterium lacus |
| 868131 | Methanobacterium paludis |
| 2025351 | Methanobacterium sp. BAmetb5 |
| 2025350 | Methanobacterium sp. BRmetb2 |
| 1379702 | Methanobacterium sp. MB1 |
| 1911685 | Methanobacterium sp. MZ-A1 |
| 59277 | Methanobacterium subterraneum |
| 230361 | Methanobrevibacter millerae |
| 294671 | Methanobrevibacter olleyae |
| 634498 | Methanobrevibacter ruminantium M1 |
| 420247 | Methanobrevibacter smithii ATCC 35061 |
| 224719 | Methanobrevibacter sp. AbM4 |
| 1609968 | Methanobrevibacter sp. YE315 |
| 1301915 | Methanocaldococcus bathoardescens |
| 573064 | Methanocaldococcus fervens AG86 |
| 573063 | Methanocaldococcus infernus ME |
| 243232 | Methanocaldococcus jannaschii DSM 2661 |
| 644281 | Methanocaldococcus sp. FS406-22 |
| 579137 | Methanocaldococcus vulcanius M7 |
| 351160 | Methanocella arvoryzae MRE50 |
| 1041930 | Methanocella conradii HZ254 |
| 304371 | Methanocella paludicola SANAE |
| 259564 | Methanococcoides burtonii DSM 6242 |
| 1434104 | Methanococcoides methylutens MM1 |
| 419665 | Methanococcus aeolicus Nankai-3 |
| 402880 | Methanococcus maripaludis C5 |
| 444158 | Methanococcus maripaludis C6 |
| 426368 | Methanococcus maripaludis C7 |

| NCBI Taxon Identifier | Organism Name |
|---|---|
| 267377 | Methanococcus maripaludis S2 |
| 1053692 | Methanococcus maripaludis X1 |
| 406327 | Methanococcus vannielii SB |
| 456320 | Methanococcus voltae A3 |
| 410358 | Methanocorpusculum labreanum Z |
| 1201294 | Methanoculleus bourgensis MS2 |
| 368407 | Methanoculleus marisnigri JR1 |
| 86622 | Methanoculleus sp. MAB1 |
| 1495144 | methanogenic archaeon ISO4-H5 |
| 644295 | Methanohalobium evestigatum Z-7303 |
| 2177 | Methanohalophilus halophilus |
| 547558 | Methanohalophilus mahii DSM 5219 |
| 679926 | Methanolacinia petrolearia DSM 11571 |
| 1094980 | Methanolobus psychrophilus R15 |
| 867904 | Methanomethylovorans hollandica DSM 15978 |
| 190192 | Methanopyrus kandleri AV19 |
| 456442 | Methanoregula boonei 6A8 |
| 593750 | Methanoregula formicica SMSP |
| 679901 | Methanosalsum zhilinae DSM 4017 |
| 188937 | Methanosarcina acetivorans C2A |
| 1434106 | Methanosarcina barkeri 227 |
| 1434107 | Methanosarcina barkeri 3 |
| 1434108 | Methanosarcina barkeri MS |
| 269797 | Methanosarcina barkeri str. Fusaro |
| 1434109 | Methanosarcina barkeri str. Wiesmoor |
| 1715806 | Methanosarcina flavescens |
| 1434110 | Methanosarcina horonobensis HB-1 = JCM 15518 |
| 1434111 | Methanosarcina lacustris Z-7289 |
| 1434113 | Methanosarcina mazei C16 |
| 192952 | Methanosarcina mazei Go1 |
| 213585 | Methanosarcina mazei S-6 |
| 1236903 | Methanosarcina mazei Tuc01 |
| 1434118 | Methanosarcina siciliae C2J |
| 1434119 | Methanosarcina siciliae HI350 |
| 1434120 | Methanosarcina siciliae T4/M |
| 1434099 | Methanosarcina sp. Kolksee |
| 1434100 | Methanosarcina sp. MTP4 |
| 1434102 | Methanosarcina sp. WH1 |
| 1434103 | Methanosarcina sp. WWM596 |
| 1434121 | Methanosarcina thermophila CHTI-55 |
| 523844 | Methanosarcina thermophila TM-1 |

| NCBI Taxon Identifier | Organism Name |
|---|---|
| 1434123 | Methanosarcina vacuolata Z-761 |
| 1789762 | Methanosphaera sp. BMS |
| 339860 | Methanosphaera stadtmanae DSM 3091 |
| 521011 | Methanosphaerula palustris E1-9c |
| 323259 | Methanospirillum hungatei JF-1 |
| 79929 | Methanothermobacter marburgensis str. Marburg |
| 866790 | Methanothermobacter sp. CaT2 |
| 2017966 | Methanothermobacter sp. EMTCatA1 |
| 1898379 | Methanothermobacter sp. MT-2 |
| 187420 | Methanothermobacter thermautotrophicus str. Delta H |
| 145261 | Methanothermobacter wolfeii |
| 647113 | Methanothermococcus okinawensis IH1 |
| 523846 | Methanothermus fervidus DSM 2088 |
| 1110509 | Methanothrix harundinacea 6Ac |
| 990316 | Methanothrix soehngenii GP6 |
| 349307 | Methanothrix thermoacetophila PT |
| 880724 | Methanotorris igneus Kol 5 |
| 481448 | Methylacidiphilum infernorum V4 |
| 420662 | Methylibium petroleiphilum PM1 |
| 2082386 | Methylibium sp. Pch-M |
| 265072 | Methylobacillus flagellatus KT |
| 270351 | Methylobacterium aquaticum |
| 2051553 | Methylobacterium currus |
| 2202825 | Methylobacterium durans |
| 460265 | Methylobacterium nodulans ORS 2060 |
| 693986 | Methylobacterium oryzae CBMB20 |
| 418223 | Methylobacterium phyllosphaerae |
| 426355 | Methylobacterium radiotolerans JCM 2831 |
| 426117 | Methylobacterium sp. 4-46 |
| 925818 | Methylobacterium sp. AMS5 |
| 2067957 | Methylobacterium sp. DM1 |
| 739141 | Methylobacterium sp. XJLW |
| 1432792 | Methylocaldum marinum |
| 1384459 | Methyloceanibacter caenitepidi |
| 395965 | Methylocella silvestris BL2 |
| 243233 | Methylococcus capsulatus str. Bath |
| 655015 | Methylocystis bryophila |
| 173366 | Methylocystis rosea |
| 187303 | Methylocystis sp. SC2 |
| 1538553 | Methylomonas denitrificans |
| 702114 | Methylomonas koyamae |

| NCBI Taxon Identifier | Organism Name |
|---|---|
| 857087 | Methylomonas methanica MC09 |
| 1727196 | Methylomonas sp. DH-1 |
| 107637 | Methylomonas sp. LW13 |
| 1930071 | Methylomusa anaerophila |
| 754477 | Methylophaga frappieri |
| 754476 | Methylophaga nitratireducenticrescens |
| 1662285 | Methylophilus sp. TWE2 |
| 272630 | Methylorubrum extorquens AM1 |
| 440085 | Methylorubrum extorquens CM4 |
| 661410 | Methylorubrum extorquens DM4 |
| 419610 | Methylorubrum extorquens PA1 |
| 441620 | Methylorubrum populi BJ001 |
| 29429 | Methylorubrum zatmanii |
| 595536 | Methylosinus trichosporium OB3b |
| 583345 | Methylotenera mobilis JLW8 |
| 666681 | Methylotenera versatilis 301 |
| 1091494 | Methylotuvimicrobium alcaliphilum 20Z |
| 1842540 | Methyloversatilis sp. RAC08 |
| 569860 | Methylovirgula ligni |
| 582744 | Methylovorus glucosotrophus SIP3-4 |
| 887061 | Methylovorus sp. MP688 |
| 1704499 | Methylovulum psychrotolerans |
| 294746 | Meyerozyma guilliermondii ATCC 6260 |
| 856793 | Micavibrio aeruginosavorus ARL-13 |
| 349215 | Micavibrio aeruginosavorus EPB |
| 36805 | Microbacterium aurum |
| 84292 | Microbacterium chocolatum |
| 104336 | Microbacterium foliorum |
| 162426 | Microbacterium hominis |
| 273677 | Microbacterium oleivorans |
| 300019 | Microbacterium paludicola |
| 1916917 | Microbacterium sp. 1.5R |
| 1906742 | Microbacterium sp. BH-3-3-3 |
| 1696072 | Microbacterium sp. CGR1 |
| 1714373 | Microbacterium sp. No. 7 |
| 1795053 | Microbacterium sp. PAMC 28756 |
| 367477 | Microbacterium sp. XT11 |
| 979556 | Microbacterium testaceum StLB037 |
| 260552 | Microbulbifer agarilyticus |
| 1769779 | Microbulbifer aggregans |
| 359370 | Microbulbifer sp. A4B17 |

| NCBI Taxon Identifier | Organism Name |
|---|---|
| 252514 | Microbulbifer thermotolerans |
| 279828 | Microcella alkaliphila |
| 465515 | Micrococcus luteus NCTC 2665 |
| 449447 | Microcystis aeruginosa NIES-843 |
| 1638788 | Microcystis panniformis FACHB-1757 |
| 1967666 | Microcystis sp. MC19 |
| 1032480 | Microlunatus phosphovorus NM-1 |
| 296587 | Micromonas commoda |
| 564608 | Micromonas pusilla CCMP1545 |
| 644283 | Micromonospora aurantiaca ATCC 27029 |
| 263358 | Micromonospora maris AB-18-032 |
| 2201999 | Micromonospora sp. B006 |
| 648999 | Micromonospora sp. L5 |
| 479978 | Micromonospora tulbaghiae |
| 69319 | Microplitis demolitor |
| 75385 | Micropruina glycogenica |
| 412690 | Microterricola viridarii |
| 1882682 | Microvirga ossetica |
| 2082949 | Microvirga sp. 17 mud 1-3 |
| 888845 | Minicystis rosea |
| 1658665 | Mitsuaria sp. 7 |
| 665913 | Mixta calida |
| 665914 | Mixta gaviniae |
| 6573 | Mizuhopecten yessoensis |
| 548479 | Mobiluncus curtisii ATCC 43063 |
| 477641 | Modestobacter marinus |
| 114527 | Mogibacterium diversum |
| 149040 | Mollisia scopiformis |
| 3673 | Momordica charantia |
| 554373 | Moniliophthora perniciosa FA553 |
| 1381753 | Moniliophthora roreri MCA 2997 |
| 13616 | Monodelphis domestica |
| 307658 | Monomorium pharaonis |
| 43700 | Monopterus albus |
| 145388 | Monoraphidium neglectum |
| 431895 | Monosiga brevicollis MX1 |
| 264732 | Moorella thermoacetica ATCC 39073 |
| 1458985 | Moorena producens PAL-8-15-08-1 |
| 476 | Moraxella bovis |
| 386891 | Moraxella bovoculi |
| 480 | Moraxella catarrhalis |

| NCBI Taxon Identifier | Organism Name |
|---|---|
| 480 | Moraxella catarrhalis |
| 1236608 | Moraxella catarrhalis BBH18 |
| 34062 | Moraxella osloensis |
| 29433 | Moraxella ovis |
| 1124991 | Morganella morganii subsp. morganii KT |
| 80854 | Moritella viscosa |
| 69539 | Moritella yayanosii |
| 2305508 | Mucilaginibacter celer |
| 1550579 | Mucilaginibacter gotjawali |
| 1300914 | Mucilaginibacter sp. PAMC 26640 |
| 1234841 | Mucilaginibacter xinganensis |
| 1433126 | Mucinivorans hirudinis |
| 1796646 | Muribaculum intestinale |
| 2507538 | Muriicola soli |
| 10090 | Mus musculus |
| 214687 | Musa acuminata subsp. malaccensis |
| 7370 | Musca domestica |
| 579405 | Musicola paradisiaca Ech703 |
| 2079792 | Mycetocola zhujimingii |
| 882378 | Mycetohabitans rhizoxinica HKI 454 |
| 1553431 | Mycoavidus cysteinexigens |
| 243243 | Mycobacterium avium 104 |
| 1770 | Mycobacterium avium subsp. paratuberculosis |
| 1770 | Mycobacterium avium subsp. paratuberculosis |
| 262316 | Mycobacterium avium subsp. paratuberculosis K-10 |
| 1199187 | Mycobacterium avium subsp. paratuberculosis MAP4 |
| 1048245 | Mycobacterium canettii CIPT 140010059 |
| 1205676 | Mycobacterium canettii CIPT 140060008 |
| 1205675 | Mycobacterium canettii CIPT 140070008 |
| 1205674 | Mycobacterium canettii CIPT 140070010 |
| 1205677 | Mycobacterium canettii CIPT 140070017 |
| 482462 | Mycobacterium dioxanotrophicus |
| 1202450 | Mycobacterium haemophilum DSM 44634 |
| 487521 | Mycobacterium intracellulare ATCC 13950 |
| 1138382 | Mycobacterium intracellulare MOTT-02 |
| 222805 | Mycobacterium intracellulare subsp. chimaera |
| 1232724 | Mycobacterium intracellulare subsp. intracellulare MTCC 9506 |
| 1138871 | Mycobacterium intracellulare subsp. yongonense 05-1390 |
| 557599 | Mycobacterium kansasii ATCC 12478 |
| 561304 | Mycobacterium leprae Br4923 |
| 272631 | Mycobacterium leprae TN |

| NCBI Taxon Identifier | Organism Name |
|---|---|
| 64667 | Mycobacterium lepraemurium |
| 459424 | Mycobacterium liflandii 128FXT |
| 1131442 | Mycobacterium marinum E11 |
| 216594 | Mycobacterium marinum M |
| 701042 | Mycobacterium marseillense |
| 1138383 | Mycobacterium paraintracellulare |
| 722731 | Mycobacterium shigaense |
| 1545728 | Mycobacterium sp. EPa45 |
| 164757 | Mycobacterium sp. JLS |
| 212767 | Mycobacterium sp. JS623 |
| 189918 | Mycobacterium sp. KMS |
| 164756 | Mycobacterium sp. MCS |
| 1168287 | Mycobacterium sp. MOTT36Y |
| 1273687 | Mycobacterium sp. VKM Ac-1817D |
| 1138877 | Mycobacterium tuberculosis 7199-99 |
| 1010836 | Mycobacterium tuberculosis BT1 |
| 1010835 | Mycobacterium tuberculosis BT2 |
| 1310114 | Mycobacterium tuberculosis CAS/NITR204 |
| 443149 | Mycobacterium tuberculosis CCDC5079 |
| 443149 | Mycobacterium tuberculosis CCDC5079 |
| 443150 | Mycobacterium tuberculosis CCDC5180 |
| 83331 | Mycobacterium tuberculosis CDC1551 |
| 707235 | Mycobacterium tuberculosis CTRI-2 |
| 1306414 | Mycobacterium tuberculosis EAI5 |
| 1310115 | Mycobacterium tuberculosis EAI5/NITR206 |
| 336982 | Mycobacterium tuberculosis F11 |
| 419947 | Mycobacterium tuberculosis H37Ra |
| 83332 | Mycobacterium tuberculosis H37Rv |
| 83332 | Mycobacterium tuberculosis H37Rv |
| 1010834 | Mycobacterium tuberculosis HKBS1 |
| 478434 | Mycobacterium tuberculosis KZN 1435 |
| 478433 | Mycobacterium tuberculosis KZN 4207 |
| 478435 | Mycobacterium tuberculosis KZN 605 |
| 1091500 | Mycobacterium tuberculosis RGTB327 |
| 1091501 | Mycobacterium tuberculosis RGTB423 |
| 1306400 | Mycobacterium tuberculosis str. Beijing/NITR203 |
| 652616 | Mycobacterium tuberculosis str. Erdman = ATCC 35801 |
| 395095 | Mycobacterium tuberculosis str. Haarlem |
| 1304279 | Mycobacterium tuberculosis str. Haarlem/NITR202 |
| 1097669 | Mycobacterium tuberculosis UT205 |
| 572418 | Mycobacterium tuberculosis variant africanum GM041182 |

| NCBI Taxon Identifier | Organism Name |
|---|---|
| 233413 | Mycobacterium tuberculosis variant bovis AF2122/97 |
| 998092 | Mycobacterium tuberculosis variant bovis BCG str. ATCC 35743 |
| 1206780 | Mycobacterium tuberculosis variant bovis BCG str. Korea 1168P |
| 717522 | Mycobacterium tuberculosis variant bovis BCG str. Mexico |
| 410289 | Mycobacterium tuberculosis variant bovis BCG str. Pasteur 1173P2 |
| 561275 | Mycobacterium tuberculosis variant bovis BCG str. Tokyo 172 |
| 1806 | Mycobacterium tuberculosis variant microti |
| 362242 | Mycobacterium ulcerans Agy99 |
| 561007 | Mycobacteroides abscessus ATCC 19977 |
| 1303024 | Mycobacteroides abscessus subsp. bolletii 50594 |
| 1001714 | Mycobacteroides abscessus subsp. massiliense CCUG 48898 = JCM 15300 |
| 1198627 | Mycobacteroides abscessus subsp. massiliense str. GO 06 |
| 1460372 | Mycobacteroides chelonae CCUG 47445 |
| 83262 | Mycobacteroides immunogenum |
| 1578165 | Mycobacteroides saopaulense |
| 875328 | Mycolicibacter sinensis |
| 1788 | Mycolicibacter terrae |
| 710421 | Mycolicibacterium chubuense NBB4 |
| 1766 | Mycolicibacterium fortuitum |
| 350054 | Mycolicibacterium gilvum PYR-GCK |
| 278137 | Mycolicibacterium gilvum Spyr1 |
| 134601 | Mycolicibacterium goodii |
| 1122247 | Mycolicibacterium hassiacum DSM 44199 |
| 758802 | Mycolicibacterium litorale |
| 1795 | Mycolicibacterium neoaurum |
| 700508 | Mycolicibacterium neoaurum VKM Ac-1815D |
| 1771 | Mycolicibacterium phlei |
| 710685 | Mycolicibacterium rhodesiae NBB3 |
| 1772 | Mycolicibacterium smegmatis |
| 1772 | Mycolicibacterium smegmatis |
| 246196 | Mycolicibacterium smegmatis MC2 155 |
| 246196 | Mycolicibacterium smegmatis MC2 155 |
| 246196 | Mycolicibacterium smegmatis MC2 155 |
| 1797 | Mycolicibacterium thermoresistibile |
| 1354275 | Mycolicibacterium vaccae 95051 |
| 350058 | Mycolicibacterium vanbaalenii PYR-1 |
| 340047 | Mycoplasma capricolum subsp. capricolum ATCC 27343 |
| 40480 | Mycoplasma capricolum subsp. capripneumoniae |
| 40480 | Mycoplasma capricolum subsp. capripneumoniae |
| 40480 | Mycoplasma capricolum subsp. capripneumoniae |
| 1124992 | Mycoplasma capricolum subsp. capripneumoniae 87001 |

| NCBI Taxon Identifier | Organism Name |
|---|---|
| 512564 | Mycoplasma crocodyli MP145 |
| 1111676 | Mycoplasma haemocanis str. Illinois |
| 859194 | Mycoplasma haemofelis Ohio2 |
| 941640 | Mycoplasma haemofelis str. Langford 1 |
| 866629 | Mycoplasma leachii 99/014/6 |
| 880447 | Mycoplasma leachii PG50 |
| 267748 | Mycoplasma mobile 163K |
| 862259 | Mycoplasma mycoides subsp. capri LC str. 95010 |
| 2103 | Mycoplasma mycoides subsp. mycoides |
| 865867 | Mycoplasma mycoides subsp. mycoides SC str. Gladysdale |
| 272632 | Mycoplasma mycoides subsp. mycoides SC str. PG1 |
| 1415773 | Mycoplasma ovis str. Michigan |
| 1403316 | Mycoplasma parvum str. Indiana |
| 743965 | Mycoplasma putrefaciens KS1 |
| 1292033 | Mycoplasma putrefaciens Mput9231 |
| 1749074 | Mycoplasma sp. (ex Biomphalaria glabrata) |
| 708248 | Mycoplasma suis KI3806 |
| 768700 | Mycoplasma suis str. Illinois |
| 1197325 | Mycoplasma wenyonii str. Massachusetts |
| 743967 | Mycoplasma yeatsii GM274B |
| 1159203 | Mycoplasmoides gallisepticum CA06_2006.052-5-2P |
| 1159202 | Mycoplasmoides gallisepticum NC06_2006.080-5-2P |
| 1159204 | Mycoplasmoides gallisepticum NC08_2008.031-4-3P |
| 1159198 | Mycoplasmoides gallisepticum NC95_13295-2-2P |
| 1159199 | Mycoplasmoides gallisepticum NC96_1596-4-2P |
| 1159200 | Mycoplasmoides gallisepticum NY01_2001.047-5-1P |
| 1006581 | Mycoplasmoides gallisepticum S6 |
| 708616 | Mycoplasmoides gallisepticum str. F |
| 710128 | Mycoplasmoides gallisepticum str. R(high) |
| 710127 | Mycoplasmoides gallisepticum str. R(low) |
| 1159197 | Mycoplasmoides gallisepticum VA94_7994-1-7P |
| 1159201 | Mycoplasmoides gallisepticum WI01_2001.043-13-2P |
| 243273 | Mycoplasmoides genitalium G37 |
| 662947 | Mycoplasmoides genitalium M2288 |
| 663918 | Mycoplasmoides genitalium M2321 |
| 662946 | Mycoplasmoides genitalium M6282 |
| 662945 | Mycoplasmoides genitalium M6320 |
| 1112856 | Mycoplasmoides pneumoniae 309 |
| 722438 | Mycoplasmoides pneumoniae FH |
| 272634 | Mycoplasmoides pneumoniae M129 |
| 1238993 | Mycoplasmoides pneumoniae M129-B7 |

| NCBI Taxon Identifier | Organism Name |
|---|---|
| 2110 | Mycoplasmopsis agalactiae |
| 347257 | Mycoplasmopsis agalactiae PG2 |
| 2094 | Mycoplasmopsis arginini |
| 2112 | Mycoplasmopsis bovigenitalium |
| 1316930 | Mycoplasmopsis bovis CQ-W70 |
| 767465 | Mycoplasmopsis bovis HB0801 |
| 956483 | Mycoplasmopsis bovis Hubei-1 |
| 289397 | Mycoplasmopsis bovis PG45 |
| 2113 | Mycoplasmopsis californica |
| 1397850 | Mycoplasmopsis californica HAZ160_1 |
| 29555 | Mycoplasmopsis canis |
| 1117644 | Mycoplasmopsis canis PG 14 |
| 1246955 | Mycoplasmopsis cynos C142 |
| 637387 | Mycoplasmopsis fermentans JER |
| 943945 | Mycoplasmopsis fermentans M64 |
| 496833 | Mycoplasmopsis fermentans PG18 |
| 29556 | Mycoplasmopsis gallinacea |
| 48003 | Mycoplasmopsis pullorum |
| 272635 | Mycoplasmopsis pulmonis UAB CTIP |
| 262723 | Mycoplasmopsis synoviae 53 |
| 1267001 | Mycoplasmopsis synoviae ATCC 25204 |
| 109478 | Myotis brandtii |
| 225400 | Myotis davidii |
| 76832 | Myroides odoratimimus |
| 480520 | Myroides profundi |
| 1583100 | Myroides sp. A21 |
| 1458492 | Myroides sp. ZB35 |
| 483219 | Myxococcus fulvus HW-1 |
| 1297742 | Myxococcus hansupus |
| 1278073 | Myxococcus stipitatus DSM 14675 |
| 246197 | Myxococcus xanthus DK 1622 |
| 744533 | Naegleria gruberi strain NEG-M |
| 1902245 | Nakamurella antarctica |
| 479431 | Nakamurella multipartita DSM 44233 |
| 284593 | Nakaseomyces glabratus CBS 138 |
| 1093141 | Nannochloropsis gaditana CCMP526 |
| 1026970 | Nannospalax galili |
| 228908 | Nanoarchaeum equitans Kin4-M |
| 1737403 | Nanohaloarchaea archaeon SG9 |
| 125878 | Nanorana parkeri |
| 7425 | Nasonia vitripennis |

| NCBI Taxon Identifier | Organism Name |
|---|---|
| 457570 | Natranaerobius thermophilus JW/NM-WN-LF |
| 745377 | Natrarchaeobaculum aegyptiacum |
| 2044521 | Natrarchaeobaculum sulfurireducens |
| 2044521 | Natrarchaeobaculum sulfurireducens |
| 547559 | Natrialba magadii ATCC 43099 |
| 797303 | Natrinema pellirubrum DSM 15624 |
| 406552 | Natrinema sp. J7-2 |
| 797304 | Natronobacterium gregoryi SP2 |
| 694430 | Natronococcus occultus SP4 |
| 268739 | Natronomonas moolapensis 8.8.11 |
| 348780 | Natronomonas pharaonis DSM 2160 |
| 588898 | Natronorubrum daqingense |
| 27288 | Naumovozyma castellii |
| 1071378 | Naumovozyma dairenensis CBS 421 |
| 598659 | Nautilia profundicola AmH |
| 51031 | Necator americanus |
| 1853278 | Neisseria chenwenguii |
| 546263 | Neisseria elongata subsp. glycolytica ATCC 29315 |
| 242231 | Neisseria gonorrhoeae FA 1090 |
| 521006 | Neisseria gonorrhoeae NCCP11945 |
| 489653 | Neisseria lactamica 020-06 |
| 487 | Neisseria meningitidis |
| 374833 | Neisseria meningitidis 053442 |
| 604162 | Neisseria meningitidis 8013 |
| 662598 | Neisseria meningitidis alpha14 |
| 630588 | Neisseria meningitidis alpha710 |
| 272831 | Neisseria meningitidis FAM18 |
| 935599 | Neisseria meningitidis G2136 |
| 909420 | Neisseria meningitidis H44/76 |
| 935591 | Neisseria meningitidis M01-240149 |
| 935588 | Neisseria meningitidis M01-240355 |
| 935593 | Neisseria meningitidis M04-240196 |
| 122586 | Neisseria meningitidis MC58 |
| 935589 | Neisseria meningitidis NZ-05/33 |
| 942513 | Neisseria meningitidis WUE 2594 |
| 122587 | Neisseria meningitidis Z2491 |
| 488 | Neisseria mucosa |
| 490 | Neisseria sicca |
| 28091 | Neisseria weaveri |
| 4432 | Nelumbo nucifera |
| 45351 | Nematostella vectensis |

| NCBI Taxon Identifier | Organism Name |
|---|---|
| 320497 | Neoasaia chiangmaiensis |
| 1353976 | Neochlamydia sp. S13 |
| 1287680 | Neofusicoccum parvum UCRNP2 |
| 556325 | Neomicrococcus aestuarii |
| 1028801 | Neorhizobium galegae bv. officinalis bv. officinalis str. HAMBI 1141 |
| 1028800 | Neorhizobium galegae bv. orientalis str. HAMBI 540 |
| 1825976 | Neorhizobium sp. NCHU2750 |
| 2060726 | Neorhizobium sp. SOG26 |
| 1286528 | Neorickettsia helminthoeca str. Oregon |
| 434131 | Neorickettsia risticii str. Illinois |
| 222891 | Neorickettsia sennetsu str. Miyayama |
| 367110 | Neurospora crassa OR74A |
| 510951 | Neurospora tetrasperma FGSC 2508 |
| 1176587 | Niabella ginsenosidivorans |
| 929713 | Niabella soli DSM 19437 |
| 700598 | Niastella koreensis GR20-10 |
| 49451 | Nicotiana attenuata |
| 4096 | Nicotiana sylvestris |
| 4097 | Nicotiana tabacum |
| 4098 | Nicotiana tomentosiformis |
| 110193 | Nicrophorus vespilloides |
| 128390 | Nipponia nippon |
| 391774 | Nitratidesulfovibrio vulgaris DP4 |
| 573059 | Nitratidesulfovibrio vulgaris RCH1 |
| 882 | Nitratidesulfovibrio vulgaris str. Hildenborough |
| 883 | Nitratidesulfovibrio vulgaris str. 'Miyazaki F' |
| 749222 | Nitratifractor salsuginis DSM 16511 |
| 387092 | Nitratiruptor sp. SB155-2 |
| 323097 | Nitrobacter hamburgensis X14 |
| 323098 | Nitrobacter winogradskyi Nb-255 |
| 472759 | Nitrosococcus halophilus Nc 4 |
| 323261 | Nitrosococcus oceani ATCC 19707 |
| 105559 | Nitrosococcus watsonii C-113 |
| 44574 | Nitrosomonas communis |
| 228410 | Nitrosomonas europaea ATCC 19718 |
| 335283 | Nitrosomonas eutropha C91 |
| 153948 | Nitrosomonas sp. AL212 |
| 261292 | Nitrosomonas sp. Is79A3 |
| 44577 | Nitrosomonas ureae |
| 1580092 | Nitrosopumilus adriaticus |
| 436308 | Nitrosopumilus maritimus SCM1 |

| NCBI Taxon Identifier | Organism Name |
|---|---|
| 1582439 | Nitrosopumilus piranensis |
| 926571 | Nitrososphaera viennensis EN76 |
| 1288494 | Nitrosospira lacus |
| 323848 | Nitrosospira multiformis ATCC 25196 |
| 330214 | Nitrospira defluvii |
| 1325564 | Nitrospira japonica |
| 42253 | Nitrospira moscoviensis |
| 1441467 | Nitrospirillum viridazoti CBAmc |
| 1612173 | Niveispirillum cyanobacteriorum |
| 1133849 | Nocardia brasiliensis ATCC 700358 |
| 1127134 | Nocardia cyriacigeorgica GUH-2 |
| 37329 | Nocardia farcinica |
| 247156 | Nocardia farcinica IFM 10152 |
| 2213200 | Nocardia mangyaensis |
| 1415166 | Nocardia nova SH22a |
| 37332 | Nocardia seriolae |
| 455432 | Nocardia terpenica |
| 1300347 | Nocardioides dokdonensis FR1436 |
| 196162 | Nocardioides sp. JS614 |
| 1205910 | Nocardiopsis alba ATCC BAA-2165 |
| 446468 | Nocardiopsis dassonvillei subsp. dassonvillei DSM 43111 |
| 1235441 | Nocardiopsis gilva YIM 90087 |
| 1914872 | Nodularia spumigena UHCC 0039 |
| 61853 | Nomascus leucogenys |
| 592029 | Nonlabens dokdonensis DSW-6 |
| 2496866 | Nonlabens ponticola |
| 2058134 | Nonlabens sp. MB-3u-79 |
| 1476901 | Nonlabens sp. MIC269 |
| 323273 | Nonlabens tegetincola |
| 551115 | 'Nostoc azollae' 0708 |
| 2038116 | Nostoc flagelliforme CCNUN1 |
| 63737 | Nostoc punctiforme PCC 73102 |
| 1869241 | Nostoc sp. CENA543 |
| 1751286 | Nostoc sp. NIES-3756 |
| 317936 | Nostoc sp. PCC 7107 |
| 103690 | Nostoc sp. PCC 7120 = FACHB-418 |
| 28072 | Nostoc sp. PCC 7524 |
| 1940762 | Nostocales cyanobacterium HT-58-2 |
| 105023 | Nothobranchius furzeri |
| 8208 | Notothenia coriiceps |
| 1471761 | Novibacillus thermophilus |

| NCBI Taxon Identifier | Organism Name |
|---|---|
| 279238 | Novosphingobium aromaticivorans DSM 12444 |
| 1088721 | Novosphingobium pentaromativorans US6-1 |
| 158500 | Novosphingobium resinovorum |
| 1609758 | Novosphingobium sp. P6W |
| 702113 | Novosphingobium sp. PP1Y |
| 1016987 | Novosphingobium sp. THN1 |
| 82983 | Obesumbacterium proteus |
| 716816 | Oceanicoccus sagamiensis |
| 511062 | Oceanimonas sp. GK1 |
| 1903694 | Oceanisphaera avium |
| 1416627 | Oceanisphaera profunda |
| 670487 | Oceanithermus profundus DSM 14977 |
| 221109 | Oceanobacillus iheyensis HTE831 |
| 2052660 | Oceanobacillus zhaokaii |
| 271865 | Ochrobactrum quorumnocens |
| 391626 | Octadecabacter antarcticus 307 |
| 391616 | Octadecabacter arcticus 238 |
| 1458307 | Octadecabacter temperatus |
| 37653 | Octopus bimaculoides |
| 9708 | Odobenus rosmarus divergens |
| 709991 | Odoribacter splanchnicus DSM 20712 |
| 203123 | Oenococcus oeni PSU-1 |
| 2203724 | Oenococcus sicerae |
| 2259623 | Oenococcus sp. UCMA 16435 |
| 158386 | Olea europaea var. sylvestris |
| 207559 | Oleidesulfovibrio alaskensis G20 |
| 141451 | Oleiphilus messinensis |
| 698738 | Oleispira antarctica RB-8 |
| 90245 | Oligella urethralis |
| 639310 | Olleya aquimaris |
| 2058135 | Olleya sp. Bg11-27 |
| 712411 | Olsenella sp. oral taxon 807 |
| 633147 | Olsenella uli DSM 7084 |
| 74940 | Oncorhynchus tshawytscha |
| 262768 | Onion yellows phytoplasma OY-M |
| 2015173 | Ooceraea biroi |
| 6198 | Opisthorchis viverrini |
| 794903 | Opitutaceae bacterium TAV5 |
| 452637 | Opitutus terrae PB90-1 |
| 9733 | Orcinus orca |
| 357244 | Orientia tsutsugamushi str. Boryong |

| NCBI Taxon Identifier | Organism Name |
|---|---|
| 334380 | Orientia tsutsugamushi str. Ikeda |
| 2283195 | Ornithinimicrobium avium |
| 867902 | Ornithobacterium rhinotracheale DSM 15997 |
| 1401325 | Ornithobacterium rhinotracheale ORT-UMN 88 |
| 9258 | Ornithorhynchus anatinus |
| 1851544 | Orrella dioscoreae |
| 2163011 | Orrella marina |
| 9986 | Oryctolagus cuniculus |
| 4533 | Oryza brachyantha |
| 39947 | Oryza sativa Japonica Group |
| 39947 | Oryza sativa Japonica Group |
| 8090 | Oryzias latipes |
| 56110 | Oscillatoria acuminata PCC 6304 |
| 179408 | Oscillatoria nigro-viridis PCC 7112 |
| 693746 | Oscillibacter valericigenes Sjm18-20 |
| 436017 | Ostreococcus lucimarinus CCE9901 |
| 70448 | Ostreococcus tauri |
| 2109914 | Ottowia oryzae |
| 1658672 | Ottowia sp. oral taxon 894 |
| 9940 | Ovis aries |
| 926562 | Owenweeksia hongkongensis DSM 17368 |
| 847 | Oxalobacter formigenes |
| 1411902 | Pacificitalea manganoxidans |
| 643674 | Paenalcaligenes hominis |
| 290340 | Paenarthrobacter aurescens TC1 |
| 1126833 | Paenibacillus beijingensis |
| 160799 | Paenibacillus borealis |
| 1616788 | Paenibacillus bovis |
| 1763538 | Paenibacillus crassostreae |
| 414771 | Paenibacillus donghaensis |
| 44251 | Paenibacillus durus |
| 189425 | Paenibacillus graminis |
| 1870820 | Paenibacillus ihbetae |
| 172713 | Paenibacillus kribbensis |
| 697284 | Paenibacillus larvae subsp. larvae DSM 25430 |
| 1401 | Paenibacillus lautus |
| 1116391 | Paenibacillus mucilaginosus 3016 |
| 997761 | Paenibacillus mucilaginosus K02 |
| 1036673 | Paenibacillus mucilaginosus KNP414 |
| 162209 | Paenibacillus naphthalenovorans |
| 189426 | Paenibacillus odorifer |

| NCBI Taxon Identifier | Organism Name |
|---|---|
| 59893 | Paenibacillus peoriae |
| 1406 | Paenibacillus polymyxa |
| 1429244 | Paenibacillus polymyxa CR1 |
| 349520 | Paenibacillus polymyxa E681 |
| 1052684 | Paenibacillus polymyxa M1 |
| 886882 | Paenibacillus polymyxa SC2 |
| 1413214 | Paenibacillus polymyxa SQR-21 |
| 1073571 | Paenibacillus riograndensis SBR5 |
| 1268072 | Paenibacillus sabinae T27 |
| 1695218 | Paenibacillus sp. 32O-W |
| 1536774 | Paenibacillus sp. FSL H7-0357 |
| 1536775 | Paenibacillus sp. FSL H7-0737 |
| 1536769 | Paenibacillus sp. FSL P4-0081 |
| 1536770 | Paenibacillus sp. FSL R5-0345 |
| 1536771 | Paenibacillus sp. FSL R5-0912 |
| 1536772 | Paenibacillus sp. FSL R7-0273 |
| 1536773 | Paenibacillus sp. FSL R7-0331 |
| 867076 | Paenibacillus sp. IHB B 3084 |
| 1566358 | Paenibacillus sp. IHBB 10380 |
| 324057 | Paenibacillus sp. JDR-2 |
| 481743 | Paenibacillus sp. Y412MC10 |
| 169760 | Paenibacillus stellifer |
| 1178515 | Paenibacillus swuensis |
| 985665 | Paenibacillus terrae HPL-003 |
| 715225 | Paenibacillus vortex V453 |
| 528191 | Paenibacillus xylanexedens |
| 1462996 | Paenibacillus yonginensis |
| 2320858 | Paenisporosarcina cavernae |
| 1343739 | Palaeococcus pacificus DY20341 |
| 694427 | Paludibacter propionicigenes WB4 |
| 1387353 | Paludisphaera borealis |
| 9597 | Pan paniscus |
| 9598 | Pan troglodytes |
| 93218 | Pandoraea apista |
| 656179 | Pandoraea faecigallinarum |
| 93219 | Pandoraea norimbergensis |
| 573737 | Pandoraea oxalativorans |
| 93220 | Pandoraea pnomenusa |
| 93220 | Pandoraea pnomenusa |
| 93220 | Pandoraea pnomenusa |
| 1416914 | Pandoraea pnomenusa 3kgm |

| NCBI Taxon Identifier | Organism Name |
|---|---|
| 93221 | Pandoraea pulmonicola |
| 93222 | Pandoraea sputorum |
| 445709 | Pandoraea thiooxydans |
| 656178 | Pandoraea vervacti |
| 310915 | Pangasianodon hypophthalmus |
| 121719 | Pannonibacter phragmitetus |
| 74533 | Panthera tigris altaica |
| 59538 | Pantholops hodgsonii |
| 549 | Pantoea agglomerans |
| 1891675 | Pantoea alhagi |
| 932677 | Pantoea ananatis AJ13355 |
| 706191 | Pantoea ananatis LMG 20103 |
| 1123863 | Pantoea ananatis LMG 5342 |
| 1095774 | Pantoea ananatis PA13 |
| 1076550 | Pantoea rwandensis |
| 592316 | Pantoea sp. At-9b |
| 1484158 | Pantoea sp. PSNIH1 |
| 1484157 | Pantoea sp. PSNIH2 |
| 660596 | Pantoea stewartii subsp. stewartii DC283 |
| 470934 | Pantoea vagans |
| 712898 | Pantoea vagans C9-1 |
| 3469 | Papaver somniferum |
| 76193 | Papilio machaon |
| 435591 | Parabacteroides distasonis ATCC 8503 |
| 2025876 | Parabacteroides sp. CT06 |
| 2026199 | Paraburkholderia aromaticivorans |
| 2654982 | Paraburkholderia atlantica |
| 134536 | Paraburkholderia caledonica |
| 1323664 | Paraburkholderia caribensis MBA4 |
| 134537 | Paraburkholderia fungorum |
| 60548 | Paraburkholderia graminis |
| 169430 | Paraburkholderia hospita |
| 1229205 | Paraburkholderia phenoliruptrix BR3459a |
| 391038 | Paraburkholderia phymatum STM815 |
| 1417228 | Paraburkholderia phytofirmans OLGA172 |
| 398527 | Paraburkholderia phytofirmans PsJN |
| 1926494 | Paraburkholderia sp. SOS3 |
| 754502 | Paraburkholderia sprentiae WSM5005 |
| 311230 | Paraburkholderia terrae |
| 169427 | Paraburkholderia terricola |
| 266265 | Paraburkholderia xenovorans LB400 |

| NCBI Taxon Identifier | Organism Name |
|---|---|
| 266265 | Paraburkholderia xenovorans LB400 |
| 765952 | Parachlamydia acanthamoebae UV-7 |
| 61635 | Paracholeplasma brassicae |
| 643561 | Paracidovorax avenae ATCC 19860 |
| 397945 | Paracidovorax citrulli AAC00-1 |
| 1505 | Paraclostridium sordellii |
| 1904441 | Paracoccaceae bacterium |
| 502780 | Paracoccidioides brasiliensis Pb18 |
| 502779 | Paracoccidioides lutzii Pb01 |
| 1367847 | Paracoccus aminophilus JCM 7686 |
| 34004 | Paracoccus aminovorans |
| 1945662 | Paracoccus contaminans |
| 318586 | Paracoccus denitrificans PD1222 |
| 2065379 | Paracoccus jeotgali |
| 1499308 | Paracoccus mutanolyticus |
| 2259340 | Paracoccus suum |
| 1529068 | Paracoccus tegillarcae |
| 147645 | Paracoccus yeei |
| 1077935 | Paracoccus zhejiangensis |
| 2315862 | Paraflavitalea soli |
| 298653 | Parafrankia sp. EAN1pec |
| 1426 | Parageobacillus thermoglucosidasius |
| 634956 | Parageobacillus thermoglucosidasius C56-YS93 |
| 1129794 | Paraglaciecola psychrophila 170 |
| 3042615 | Paraglaciecola sp. T6c |
| 342108 | Paramagnetospirillum magneticum AMB-1 |
| 412030 | Paramecium tetraurelia strain d4-2 |
| 1676925 | Paramormyrops kingsleyae |
| 1755811 | Paraphotobacterium marinum |
| 1150469 | Pararhodospirillum photometricum DSM 122 |
| 864564 | Parascardovia denticolens DSM 10105 = JCM 12538 |
| 2483033 | Parasedimentitalea marina |
| 760011 | Parasphaerochaeta coccoides DSM 17374 |
| 266812 | Parasphingorhabdus flavimaris |
| 321614 | Parastagonospora nodorum SN15 |
| 84588 | Parasynechococcus marenigrum WH 8102 |
| 645517 | Paraurantiacibacter namhicola |
| 41977 | Parazoarcus communis |
| 2003188 | Parolsenella catena |
| 9157 | Parus major |
| 402881 | Parvibaculum lavamentivorans DS-1 |

| NCBI Taxon Identifier | Organism Name |
|---|---|
| 33033 | Parvimonas micra |
| 314260 | Parvularcula bermudensis HTCC2503 |
| 754 | Pasteurella dagmatis |
| 747 | Pasteurella multocida |
| 1075089 | Pasteurella multocida 36950 |
| 584721 | Pasteurella multocida subsp. multocida str. 3480 |
| 1132496 | Pasteurella multocida subsp. multocida str. HN06 |
| 272843 | Pasteurella multocida subsp. multocida str. Pm70 |
| 1768242 | Paucibacter sp. KCTC 42545 |
| 1291742 | Paucilactobacillus hokkaidonensis JCM 18461 |
| 29443 | Paucimonas lemoignei |
| 29471 | Pectobacterium atrosepticum |
| 29471 | Pectobacterium atrosepticum |
| 218491 | Pectobacterium atrosepticum SCRI1043 |
| 561230 | Pectobacterium carotovorum subsp. carotovorum PC1 |
| 1218933 | Pectobacterium carotovorum subsp. carotovorum PCC21 |
| 78398 | Pectobacterium odoriferum |
| 1905730 | Pectobacterium parmentieri |
| 1905730 | Pectobacterium parmentieri |
| 561231 | Pectobacterium parmentieri WPP163 |
| 2042057 | Pectobacterium polaris |
| 1175631 | Pectobacterium wasabiae CFBP 3304 |
| 121224 | Pediculus humanus corporis |
| 1254 | Pediococcus acidilactici |
| 701521 | Pediococcus claussenii ATCC BAA-344 |
| 51663 | Pediococcus damnosus |
| 114090 | Pediococcus inopinatus |
| 278197 | Pediococcus pentosaceus ATCC 25745 |
| 1408206 | Pediococcus pentosaceus SL4 |
| 188932 | Pedobacter cryoconitis |
| 363852 | Pedobacter ginsengisoli |
| 485917 | Pedobacter heparinus DSM 2366 |
| 1727164 | Pedobacter sp. PACM 27299 |
| 430522 | Pedobacter steynii |
| 543877 | Pelagerythrobacter marensis |
| 1082931 | Pelagibacterium halotolerans B2 |
| 338966 | Pelobacter propionicus DSM 2379 |
| 319225 | Pelodictyon luteolum DSM 273 |
| 324925 | Pelodictyon phaeoclathratiforme BU-1 |
| 13735 | Pelodiscus sinensis |
| 913107 | Pelolinea submarina |

| NCBI Taxon Identifier | Organism Name |
|---|---|
| 1192197 | Pelosinus fermentans JBW45 |
| 484770 | Pelosinus sp. UFO1 |
| 370438 | Pelotomaculum thermopropionicum SI |
| 1170230 | Penicillium digitatum Pd1 |
| 500485 | Penicillium rubens Wisconsin 54-1255 |
| 1286171 | Peptoclostridium acidaminophilum DSM 3953 |
| 54005 | Peptoniphilus harei |
| 1912856 | Peptoniphilus sp. ING2-D1G |
| 2081703 | Peptostreptococcaceae bacterium oral taxon 929 |
| 264697 | Peribacillus muralis |
| 1478 | Peribacillus simplex |
| 123214 | Persephonella marina EX-H1 |
| 1229662 | Pestalotiopsis fici W106-1 |
| 2126346 | Peterkaempfera bronchialis |
| 1642646 | Petrimonas mucosa |
| 1978337 | Petrimonas sp. IBARAKI |
| 2173034 | Petrocella atlantisensis |
| 403833 | Petrotoga mobilis SJ95 |
| 1286976 | Phaeoacremonium minimum UCRPA7 |
| 1423144 | Phaeobacter gallaeciensis DSM 26640 |
| 383629 | Phaeobacter inhibens 2.10 |
| 391619 | Phaeobacter inhibens DSM 17395 |
| 1580596 | Phaeobacter piscinae |
| 1844006 | Phaeobacter porticola |
| 681157 | Phaeobacter sp. LSS9 |
| 556484 | Phaeodactylum tricornutum CCAP 1055/1 |
| 78828 | Phalaenopsis equestris |
| 650164 | Phanerochaete carnosa HHB-10118-sp |
| 33025 | Phascolarctobacterium faecium |
| 38626 | Phascolarctos cinereus |
| 3885 | Phaseolus vulgaris |
| 2201350 | Phenylobacterium parvum |
| 450851 | Phenylobacterium zucineum HLK1 |
| 357276 | Phocaeicola dorei |
| 357276 | Phocaeicola dorei |
| 667015 | Phocaeicola salanitronis DSM 18170 |
| 435590 | Phocaeicola vulgatus ATCC 8482 |
| 42345 | Phoenix dactylifera |
| 85581 | Photobacterium damselae subsp. damselae |
| 658445 | Photobacterium gaetbulicola Gung47 |
| 298386 | Photobacterium profundum SS9 |

| NCBI Taxon Identifier | Organism Name |
|---|---|
| 291112 | Photorhabdus asymbiotica |
| 141679 | Photorhabdus laumondii subsp. laumondii |
| 243265 | Photorhabdus laumondii subsp. laumondii TTO1 |
| 230089 | Photorhabdus thracensis |
| 1868589 | Phreatobacter cathodiphilus |
| 1142394 | Phycisphaera mikurensis NBRC 102666 |
| 1867719 | Phyllobacterium zundukense |
| 3218 | Physcomitrium patens |
| 2487150 | Phytobacter sp. MRY16-398 |
| 1972431 | Phytobacter ursingii |
| 403677 | Phytophthora infestans T30-4 |
| 67593 | Phytophthora sojae |
| 32049 | Picosynechococcus sp. PCC 7002 |
| 374982 | Picosynechococcus sp. PCC 73109 |
| 1122961 | Picrophilus oshimae DSM 9789 |
| 64459 | Pieris rapae |
| 2488560 | Pigmentiphaga sp. H8 |
| 2045 | Pimelobacter simplex |
| 1632865 | Pirellula sp. SH-Sr6A |
| 530564 | Pirellula staleyi DSM 6068 |
| 946333 | Piscinibacter gummiphilus |
| 1227812 | Piscirickettsia salmonis LF-89 = ATCC VR-1361 |
| 1632864 | Planctomyces sp. SH-PL14 |
| 1636152 | Planctomyces sp. SH-PL62 |
| 521674 | Planctopirus limnophila DSM 3776 |
| 666509 | Planktomarina temperata RCA23 |
| 282423 | Planktothrix agardhii NIES-204 |
| 1185653 | Planococcus antarcticus DSM 14505 |
| 414778 | Planococcus donghaensis |
| 1598147 | Planococcus faecalis |
| 1215089 | Planococcus halocryophilus |
| 1374 | Planococcus kocurii |
| 192421 | Planococcus maritimus |
| 1038856 | Planococcus plakortidis |
| 200991 | Planococcus rifietoensis |
| 2058136 | Planococcus sp. MB-3u-03 |
| 1526927 | Planococcus sp. PAMC 21323 |
| 1302659 | Planococcus versutus |
| 2071627 | Plantactinospora sp. BB1 |
| 2108470 | Plantactinospora sp. BC1 |
| 2024580 | Plantactinospora sp. KBS50 |

| NCBI Taxon Identifier | Organism Name |
|---|---|
| 2480625 | Plantibacter sp. PA-3-X8 |
| 5823 | Plasmodium berghei ANKA |
| 31271 | Plasmodium chabaudi chabaudi |
| 1120755 | Plasmodium cynomolgi strain B |
| 36329 | Plasmodium falciparum 3D7 |
| 57267 | Plasmodium falciparum Dd2 |
| 137071 | Plasmodium falciparum HB3 |
| 5851 | Plasmodium knowlesi strain H |
| 126793 | Plasmodium vivax Sal-1 |
| 5861 | Plasmodium yoelii |
| 891974 | Plautia stali symbiont |
| 1885025 | Pleomorphomonas sp. SM30 |
| 703 | Plesiomonas shigelloides |
| 118163 | Pleurocapsa sp. PCC 7327 |
| 61647 | Pluralibacter gergoviae |
| 51655 | Plutella xylostella |
| 46731 | Pocillopora damicornis |
| 515849 | Podospora anserina S mat+ |
| 8081 | Poecilia reticulata |
| 103695 | Pogona vitticeps |
| 144034 | Pogonomyrmex barbatus |
| 996801 | Polaribacter reichenbachii |
| 2058137 | Polaribacter sp. ALD11 |
| 1529069 | Polaribacter sp. BM10 |
| 313598 | Polaribacter sp. MED152 |
| 1774273 | Polaribacter vadi |
| 365044 | Polaromonas naphthalenivorans CJ2 |
| 296591 | Polaromonas sp. JS666 |
| 2268087 | Polaromonas sp. SP1 |
| 91411 | Polistes canadensis |
| 2028345 | Pollutimonas thiosulfatoxidans |
| 991905 | Polymorphum gilvum SL003B-26A1 |
| 312153 | Polynucleobacter asymbioticus QLW-P1DMWA-1 |
| 1835254 | Polynucleobacter duraquae |
| 452638 | Polynucleobacter necessarius STIR1 |
| 2527775 | Polynucleobacter paneuropaeus |
| 400727 | Pomacea canaliculata |
| 9601 | Pongo abelii |
| 323450 | Pontibacter actiniarum |
| 400092 | Pontibacter korlensis |
| 1159327 | Pontimonas salivibrio |

| NCBI Taxon Identifier | Organism Name |
|---|---|
| 3694 | Populus trichocarpa |
| 2023229 | Porphyrobacter sp. HT-58-2 |
| 1896196 | Porphyrobacter sp. LM 6 |
| 879243 | Porphyromonas asaccharolytica DSM 20707 |
| 393921 | Porphyromonas crevioricanis |
| 431947 | Porphyromonas gingivalis ATCC 33277 |
| 1030843 | Porphyromonas gingivalis TDC60 |
| 242619 | Porphyromonas gingivalis W83 |
| 1850254 | Poseidonibacter parvus |
| 561896 | Postia placenta Mad-698-R |
| 82985 | Pragia fontium |
| 530584 | Prauserella marina |
| 685727 | Prescottella equi 103S |
| 908937 | Prevotella dentalis DSM 3688 |
| 767031 | Prevotella denticola F0289 |
| 1236517 | Prevotella fusca JCM 17724 |
| 246198 | Prevotella intermedia 17 |
| 1177574 | Prevotella jejuni |
| 553174 | Prevotella melaninogenica ATCC 25845 |
| 575614 | Prevotella sp. oral taxon 299 str. F0039 |
| 135735 | Priestia endophytica |
| 86664 | Priestia flexa |
| 592022 | Priestia megaterium DSM 319 |
| 1348623 | Priestia megaterium NBRC 15308 = ATCC 14581 |
| 545693 | Priestia megaterium QM B1551 |
| 1006007 | Priestia megaterium WSH-002 |
| 146891 | Prochlorococcus marinus str. AS9601 |
| 93059 | Prochlorococcus marinus str. MIT 9211 |
| 93060 | Prochlorococcus marinus str. MIT 9215 |
| 167546 | Prochlorococcus marinus str. MIT 9301 |
| 59922 | Prochlorococcus marinus str. MIT 9303 |
| 74546 | Prochlorococcus marinus str. MIT 9312 |
| 74547 | Prochlorococcus marinus str. MIT 9313 |
| 167542 | Prochlorococcus marinus str. MIT 9515 |
| 167555 | Prochlorococcus marinus str. NATL1A |
| 59920 | Prochlorococcus marinus str. NATL2A |
| 167539 | Prochlorococcus marinus subsp. marinus str. CCMP1375 |
| 59919 | Prochlorococcus marinus subsp. pastoris str. CCMP1986 |
| 1501268 | Prochlorococcus sp. MIT 0604 |
| 1501269 | Prochlorococcus sp. MIT 0801 |
| 104623 | Prodigiosinella confusarubida |

| NCBI Taxon Identifier | Organism Name |
|---|---|
| 556499 | Propionibacterium acidifaciens |
| 66712 | Propionibacterium freudenreichii subsp. freudenreichii |
| 754252 | Propionibacterium freudenreichii subsp. shermanii CIRM-BIA1 |
| 290512 | Prosthecochloris aestuarii DSM 271 |
| 1868325 | Prosthecochloris sp. CIB 2401 |
| 281093 | Prosthecochloris sp. GSB1 |
| 1974213 | Prosthecochloris sp. HL-130-GSB |
| 2419774 | Protaetiibacter intestinalis |
| 1642647 | Proteiniphilum saccharofermentans |
| 183417 | Proteus hauseri |
| 584 | Proteus mirabilis |
| 1266738 | Proteus mirabilis BB2000 |
| 529507 | Proteus mirabilis HI4320 |
| 585 | Proteus vulgaris |
| 103944 | Protobothrops mucrosquamatus |
| 126385 | Providencia alcalifaciens |
| 333962 | Providencia heimbachae |
| 2027290 | Providencia huaxiensis |
| 587 | Providencia rettgeri |
| 588 | Providencia stuartii |
| 588 | Providencia stuartii |
| 1157951 | Providencia stuartii MRSN 2154 |
| 42229 | Prunus avium |
| 102107 | Prunus mume |
| 3760 | Prunus persica |
| 1736674 | Pseudalgibacter alginicilyticus |
| 685565 | Pseudanabaena sp. ABRG5-3 |
| 82654 | Pseudanabaena sp. PCC 7367 |
| 452863 | Pseudarthrobacter chlorophenolicus A6 |
| 930171 | Pseudarthrobacter phenanthrenivorans Sphe3 |
| 121292 | Pseudarthrobacter sulfonivorans |
| 2067960 | Pseudazoarcus pumilus |
| 247523 | Pseudoalteromonas aliena |
| 1117313 | Pseudoalteromonas arctica A 37-1-2 |
| 621376 | Pseudoalteromonas donghaensis |
| 1314869 | Pseudoalteromonas espejiana DSM 9414 |
| 152297 | Pseudoalteromonas issachenkonii |
| 43657 | Pseudoalteromonas luteoviolacea |
| 28109 | Pseudoalteromonas nigrifaciens |
| 161398 | Pseudoalteromonas phenolica |
| 1348114 | Pseudoalteromonas piratica |

| NCBI Taxon Identifier | Organism Name |
|---|---|
| 43662 | Pseudoalteromonas piscicida |
| 43658 | Pseudoalteromonas rubra |
| 283699 | Pseudoalteromonas sp. Bsw20308 |
| 1514074 | Pseudoalteromonas sp. NC201 |
| 234831 | Pseudoalteromonas sp. SM9913 |
| 1117319 | Pseudoalteromonas spongiae UST010723-006 |
| 43659 | Pseudoalteromonas tetraodonis |
| 1315283 | Pseudoalteromonas translucida KMM 520 |
| 326442 | Pseudoalteromonas translucida TAC125 |
| 314281 | Pseudoalteromonas tunicata |
| 1184267 | Pseudobdellovibrio exovorus JSS |
| 383855 | Pseudocercospora fijiensis CIRAD86 |
| 643562 | Pseudodesulfovibrio aespoeensis Aspo-2 |
| 1716143 | Pseudodesulfovibrio indicus |
| 641491 | Pseudodesulfovibrio mercurii |
| 1322246 | Pseudodesulfovibrio piezophilus C1TLV30 |
| 57320 | Pseudodesulfovibrio profundus |
| 2072590 | Pseudoduganella armeniaca |
| 298654 | Pseudofrankia inefficax |
| 748280 | Pseudogulbenkiania sp. NH8B |
| 1249552 | Pseudohongiella spirulinae |
| 216595 | Pseudomonas [fluorescens] SBW25 |
| 287 | Pseudomonas aeruginosa |
| 1280938 | Pseudomonas aeruginosa B136-33 |
| 1352355 | Pseudomonas aeruginosa c7447m |
| 1093787 | Pseudomonas aeruginosa DK2 |
| 1408272 | Pseudomonas aeruginosa LES431 |
| 557722 | Pseudomonas aeruginosa LESB58 |
| 941193 | Pseudomonas aeruginosa M18 |
| 1415629 | Pseudomonas aeruginosa MTB-1 |
| 1089456 | Pseudomonas aeruginosa NCGM2.S1 |
| 1279007 | Pseudomonas aeruginosa PA1 |
| 1279008 | Pseudomonas aeruginosa PA1R |
| 1407059 | Pseudomonas aeruginosa PA38182 |
| 381754 | Pseudomonas aeruginosa PA7 |
| 208964 | Pseudomonas aeruginosa PAO1 |
| 1367494 | Pseudomonas aeruginosa PAO1-VE13 |
| 1367493 | Pseudomonas aeruginosa PAO1-VE2 |
| 1352354 | Pseudomonas aeruginosa PAO581 |
| 1340851 | Pseudomonas aeruginosa RP73 |
| 1427342 | Pseudomonas aeruginosa SCV20265 |

| NCBI Taxon Identifier | Organism Name |
|---|---|
| 208963 | Pseudomonas aeruginosa UCBPP-PA14 |
| 1448140 | Pseudomonas aeruginosa YL84 |
| 43263 | Pseudomonas alcaligenes |
| 237609 | Pseudomonas alkylphenolica |
| 129138 | Pseudomonas amygdali pv. morsprunorum |
| 219572 | Pseudomonas antarctica |
| 46257 | Pseudomonas avellanae |
| 47878 | Pseudomonas azotoformans |
| 930166 | Pseudomonas brassicacearum |
| 994484 | Pseudomonas brassicacearum subsp. brassicacearum NFM421 |
| 587753 | Pseudomonas chlororaphis |
| 587753 | Pseudomonas chlororaphis |
| 86192 | Pseudomonas chlororaphis subsp. aurantiaca |
| 1441629 | Pseudomonas cichorii JBC1 |
| 53408 | Pseudomonas citronellolis |
| 47879 | Pseudomonas corrugata |
| 157783 | Pseudomonas cremoricolorata |
| 384676 | Pseudomonas entomophila L48 |
| 294 | Pseudomonas fluorescens |
| 294 | Pseudomonas fluorescens |
| 294 | Pseudomonas fluorescens |
| 1037911 | Pseudomonas fluorescens A506 |
| 205922 | Pseudomonas fluorescens Pf0-1 |
| 1334632 | Pseudomonas fluorescens PICF7 |
| 296 | Pseudomonas fragi |
| 104087 | Pseudomonas frederiksbergensis |
| 743720 | Pseudomonas fulva 12-X |
| 1301098 | Pseudomonas knackmussii B13 |
| 198620 | Pseudomonas koreensis |
| 86185 | Pseudomonas lundensis |
| 1147786 | Pseudomonas mandelii JR-1 |
| 1001585 | Pseudomonas mendocina NK-01 |
| 399739 | Pseudomonas mendocina ymp |
| 1435044 | Pseudomonas monteilii SB3078 |
| 1435058 | Pseudomonas monteilii SB3101 |
| 1114970 | Pseudomonas ogarae |
| 1182590 | Pseudomonas oleovorans CECT 5344 |
| 76758 | Pseudomonas orientalis |
| 47885 | Pseudomonas oryzihabitans |
| 47885 | Pseudomonas oryzihabitans |
| 157782 | Pseudomonas parafulva |

| NCBI Taxon Identifier | Organism Name |
|---|---|
| 70775 | Pseudomonas plecoglossicida |
| 1282356 | Pseudomonas poae RE*1-1-14 |
| 1420599 | Pseudomonas protegens Cab57 |
| 1124983 | Pseudomonas protegens CHA0 |
| 220664 | Pseudomonas protegens Pf-5 |
| 303 | Pseudomonas putida |
| 931281 | Pseudomonas putida BIRD-1 |
| 1196325 | Pseudomonas putida DOT-T1E |
| 351746 | Pseudomonas putida F1 |
| 76869 | Pseudomonas putida GB-1 |
| 1331671 | Pseudomonas putida H8234 |
| 1215088 | Pseudomonas putida HB3267 |
| 160488 | Pseudomonas putida KT2440 |
| 1211579 | Pseudomonas putida NBRC 14164 |
| 231023 | Pseudomonas putida ND6 |
| 1042876 | Pseudomonas putida S16 |
| 390235 | Pseudomonas putida W619 |
| 1245471 | Pseudomonas resinovorans NBRC 106553 |
| 216142 | Pseudomonas rhizosphaerae |
| 264730 | Pseudomonas savastanoi pv. phaseolicola 1448A |
| 1853130 | Pseudomonas silesiensis |
| 1306993 | Pseudomonas soli |
| 1294143 | Pseudomonas sp. ATCC 13867 |
| 1649877 | Pseudomonas sp. CCOS 191 |
| 1611770 | Pseudomonas sp. MRSN 12121 |
| 1500686 | Pseudomonas sp. Os17 |
| 1028989 | Pseudomonas sp. StFLB209 |
| 1856685 | Pseudomonas sp. TCU-HL1 |
| 1415630 | Pseudomonas sp. TKP |
| 1207075 | Pseudomonas sp. UW4 |
| 69328 | Pseudomonas sp. VLB120 |
| 47883 | Pseudomonas synxantha |
| 1357279 | Pseudomonas syringae CC1557 |
| 205918 | Pseudomonas syringae pv. syringae B728a |
| 223283 | Pseudomonas syringae pv. tomato str. DC3000 |
| 200450 | Pseudomonas trivialis |
| 76761 | Pseudomonas veronii |
| 1788301 | Pseudomonas versuta |
| 515393 | Pseudomonas yamanorum |
| 219809 | Pseudomyrmex gracilis |
| 675635 | Pseudonocardia dioxanivorans CB1190 |

| NCBI Taxon Identifier | Organism Name |
|---|---|
| 445576 | Pseudonocardia sp. AL041005-10 |
| 1688404 | Pseudonocardia sp. EC080610-09 |
| 1096856 | Pseudonocardia sp. EC080619-01 |
| 1096868 | Pseudonocardia sp. EC080625-04 |
| 1641402 | Pseudonocardia sp. HH130629-09 |
| 1690815 | Pseudonocardia sp. HH130630-07 |
| 762903 | Pseudopedobacter saltans DSM 12145 |
| 181119 | Pseudopodoces humilis |
| 1125847 | Pseudorhizobium banfieldiae |
| 1235591 | Pseudorhodoplanes sinuspersici |
| 1402135 | Pseudosulfitobacter pseudonitzschiae |
| 1123384 | Pseudothermotoga hypogea DSM 11164 = NBRC 106472 |
| 416591 | Pseudothermotoga lettingae TMO |
| 688269 | Pseudothermotoga thermarum DSM 5069 |
| 911045 | Pseudovibrio sp. FO-BEG1 |
| 1045855 | Pseudoxanthomonas spadix BD-a59 |
| 314722 | Pseudoxanthomonas suwonensis |
| 743721 | Pseudoxanthomonas suwonensis 11-1 |
| 1277687 | Pseudozyma flocculosa PF-1 |
| 261164 | Psychrobacter alimentarius |
| 259536 | Psychrobacter arcticus 273-4 |
| 335284 | Psychrobacter cryohalolentis K5 |
| 1720344 | Psychrobacter sp. AntiMn-1 |
| 1028416 | Psychrobacter sp. DAB_AL43B |
| 571800 | Psychrobacter sp. G |
| 1699624 | Psychrobacter sp. P11G5 |
| 1699622 | Psychrobacter sp. P2G3 |
| 349106 | Psychrobacter sp. PRwf-1 |
| 2203895 | Psychrobacter sp. YP14 |
| 45610 | Psychrobacter urativorans |
| 313595 | Psychroflexus torquis ATCC 700755 |
| 1618207 | Psychromicrobium lacuslunae |
| 357804 | Psychromonas ingrahamii 37 |
| 314282 | Psychromonas sp. CNPT3 |
| 9402 | Pteropus alecto |
| 418459 | Puccinia graminis f. sp. tritici CRL 75-36-700-3 |
| 2094025 | Pukyongia salina |
| 2605946 | Pukyongiella litopenaei |
| 2116657 | Pulveribacter suum |
| 741275 | Punctularia strigosozonata HHB-11173 SS5 |
| 33203 | Purpureocillium lilacinum |

| NCBI Taxon Identifier | Organism Name |
|---|---|
| 1007105 | Pusillimonas sp. T7-7 |
| 861557 | Pyrenophora teres f. teres 0-1 |
| 242507 | Pyricularia oryzae 70-15 |
| 178306 | Pyrobaculum aerophilum str. IM2 |
| 340102 | Pyrobaculum arsenaticum DSM 13514 |
| 410359 | Pyrobaculum calidifontis JCM 11548 |
| 1104324 | Pyrobaculum ferrireducens |
| 384616 | Pyrobaculum islandicum DSM 4184 |
| 444157 | Pyrobaculum neutrophilum V24Sta |
| 698757 | Pyrobaculum oguniense TE7 |
| 1227555 | Pyrobaculum sp. WP30 |
| 272844 | Pyrococcus abyssi GE5 |
| 1185654 | Pyrococcus furiosus COM1 |
| 186497 | Pyrococcus furiosus DSM 3638 |
| 70601 | Pyrococcus horikoshii OT3 |
| 1609559 | Pyrococcus kukulkanii |
| 342949 | Pyrococcus sp. NA2 |
| 1183377 | Pyrococcus sp. ST04 |
| 529709 | Pyrococcus yayanosii CH1 |
| 1273541 | Pyrodictium delaneyi |
| 694429 | Pyrolobus fumarii 1A |
| 225117 | Pyrus x bretschneideri |
| 176946 | Python bivittatus |
| 192812 | Qipengyuania flava |
| 58331 | Quercus suber |
| 2703885 | Rahnella aceris |
| 745277 | Rahnella aquatilis CIP 78.65 = ATCC 33071 |
| 1151116 | Rahnella aquatilis HX2 |
| 1805933 | Rahnella sikkimica |
| 190721 | Ralstonia insidiosa |
| 105219 | Ralstonia mannitolilytica |
| 428406 | Ralstonia pickettii 12D |
| 402626 | Ralstonia pickettii 12J |
| 1366050 | Ralstonia pickettii DTP0602 |
| 1310165 | Ralstonia pseudosolanacearum |
| 1262456 | Ralstonia pseudosolanacearum FQY_4 |
| 267608 | Ralstonia pseudosolanacearum GMI1000 |
| 859656 | Ralstonia solanacearum CFBP2957 |
| 859655 | Ralstonia solanacearum CMR15 |
| 1031711 | Ralstonia solanacearum Po82 |
| 859657 | Ralstonia solanacearum PSI07 |

| NCBI Taxon Identifier | Organism Name |
|---|---|
| 365046 | Ramlibacter tataouinensis TTB310 |
| 54291 | Raoultella ornithinolytica |
| 1286170 | Raoultella ornithinolytica B6 |
| 575 | Raoultella planticola |
| 2259647 | Raoultella sp. X13 |
| 59737 | Rathayibacter iranicus |
| 33887 | Rathayibacter rathayi |
| 145458 | Rathayibacter toxicus |
| 145458 | Rathayibacter toxicus |
| 33888 | Rathayibacter tritici |
| 10116 | Rattus norvegicus |
| 1336806 | Reinekea forsetii |
| 288705 | Renibacterium salmoninarum ATCC 33209 |
| 89399 | Rhinolophus sinicus |
| 61621 | Rhinopithecus bieti |
| 61622 | Rhinopithecus roxellana |
| 1862950 | Rhizobiales bacterium NRL2 |
| 1967781 | Rhizobium esperanzae |
| 1432050 | Rhizobium etli bv. mimosae str. IE4771 |
| 1328306 | Rhizobium etli bv. mimosae str. Mim1 |
| 1432049 | Rhizobium etli bv. phaseoli str. IE4803 |
| 347834 | Rhizobium etli CFN 42 |
| 491916 | Rhizobium etli CIAT 652 |
| 348824 | Rhizobium favelukesii |
| 1041138 | Rhizobium gallicum bv. gallicum R602sp |
| 1312183 | Rhizobium jaguaris |
| 216596 | Rhizobium johnstonii 3841 |
| 1033991 | Rhizobium leguminosarum bv. trifolii CB782 |
| 395491 | Rhizobium leguminosarum bv. trifolii WSM1325 |
| 754523 | Rhizobium leguminosarum bv. trifolii WSM1689 |
| 395492 | Rhizobium leguminosarum bv. trifolii WSM2304 |
| 396 | Rhizobium phaseoli |
| 359 | Rhizobium rhizogenes |
| 311403 | Rhizobium rhizogenes K84 |
| 424182 | Rhizobium sp. IRBG74 |
| 2020311 | Rhizobium sp. Kim5 |
| 1703962 | Rhizobium sp. N1341 |
| 1703966 | Rhizobium sp. N731 |
| 1869170 | Rhizobium sp. S41 |
| 698761 | Rhizobium tropici CIAT 899 |
| 1850238 | Rhizorhabdus dicambivorans |

| NCBI Taxon Identifier | Organism Name |
|---|---|
| 1283312 | Rhizorhabdus wittichii DC-6 |
| 392499 | Rhizorhabdus wittichii RW1 |
| 666685 | Rhodanobacter denitrificans |
| 272942 | Rhodobacter capsulatus SB 1003 |
| 1850250 | Rhodobacter xanthinilyticus |
| 2026785 | Rhodobiaceae bacterium |
| 1828 | Rhodococcoides fascians |
| 1051973 | Rhodococcoides fascians D188 |
| 191292 | Rhodococcus aetherivorans |
| 1833 | Rhodococcus erythropolis |
| 1136179 | Rhodococcus erythropolis CCM2595 |
| 234621 | Rhodococcus erythropolis PR4 |
| 101510 | Rhodococcus jostii RHA1 |
| 632772 | Rhodococcus opacus B4 |
| 543736 | Rhodococcus opacus PD630 |
| 103816 | Rhodococcus pyridinivorans |
| 1435356 | Rhodococcus pyridinivorans SB3094 |
| 334542 | Rhodococcus qingshengii |
| 1829 | Rhodococcus rhodochrous |
| 1830 | Rhodococcus ruber |
| 1723645 | Rhodococcus sp. 008 |
| 1564114 | Rhodococcus sp. B7740 |
| 935199 | Rhodococcus sp. p52 |
| 1653478 | Rhodococcus sp. PBTS 1 |
| 679318 | Rhodococcus sp. WMMA185 |
| 1898103 | Rhodocyclaceae bacterium |
| 81479 | Rhodoferax antarcticus |
| 338969 | Rhodoferax ferrireducens T118 |
| 1842727 | Rhodoferax koreense |
| 1484693 | Rhodoferax saidenbachensis |
| 529884 | Rhodoluna lacicola |
| 648757 | Rhodomicrobium vannielii ATCC 17100 |
| 243090 | Rhodopirellula baltica SH 1 |
| 674703 | Rhodoplanes sp. Z2-YC6860 |
| 316055 | Rhodopseudomonas palustris BisA53 |
| 316056 | Rhodopseudomonas palustris BisB18 |
| 316057 | Rhodopseudomonas palustris BisB5 |
| 258594 | Rhodopseudomonas palustris CGA009 |
| 652103 | Rhodopseudomonas palustris DX-1 |
| 316058 | Rhodopseudomonas palustris HaA2 |
| 395960 | Rhodopseudomonas palustris TIE-1 |

| NCBI Taxon Identifier | Organism Name |
|---|---|
| 414684 | Rhodospirillum centenum SW |
| 269796 | Rhodospirillum rubrum ATCC 11170 |
| 1036743 | Rhodospirillum rubrum F11 |
| 1779382 | Rhodothermaceae bacterium RA |
| 518766 | Rhodothermus marinus DSM 4252 |
| 762570 | Rhodothermus marinus SG0.5JP17-172 |
| 308754 | Rhodovulum sp. MB263 |
| 1564506 | Rhodovulum sp. P5 |
| 35806 | Rhodovulum sulfidophilum |
| 3988 | Ricinus communis |
| 347255 | Rickettsia africae ESF-5 |
| 293614 | Rickettsia akari str. Hartford |
| 33989 | Rickettsia amblyommatis |
| 1105111 | Rickettsia amblyommatis str. GAT-30V |
| 1105110 | Rickettsia australis str. Cutlack |
| 391896 | Rickettsia bellii OSU 85-389 |
| 336407 | Rickettsia bellii RML369-C |
| 1105107 | Rickettsia canadensis str. CA410 |
| 293613 | Rickettsia canadensis str. McKiel |
| 272944 | Rickettsia conorii str. Malish 7 |
| 1032845 | Rickettsia conorii subsp. heilongjiangensis 054 |
| 315456 | Rickettsia felis URRWXCal2 |
| 652620 | Rickettsia japonica YH |
| 416276 | Rickettsia massiliae MTU5 |
| 1105112 | Rickettsia massiliae str. AZT80 |
| 109232 | Rickettsia monacensis |
| 1105114 | Rickettsia montanensis str. OSU 85-930 |
| 1105108 | Rickettsia parkeri str. Portsmouth |
| 562019 | Rickettsia peacockii str. Rustic |
| 481009 | Rickettsia philipii str. 364D |
| 1290428 | Rickettsia prowazekii str. Breinl |
| 1105096 | Rickettsia prowazekii str. BuV67-CWPP |
| 1105094 | Rickettsia prowazekii str. Chernikova |
| 1105097 | Rickettsia prowazekii str. Dachau |
| 1105098 | Rickettsia prowazekii str. GvV257 |
| 1105095 | Rickettsia prowazekii str. Katsinyian |
| 272947 | Rickettsia prowazekii str. Madrid E |
| 1290427 | Rickettsia prowazekii str. NMRC Madrid E |
| 449216 | Rickettsia prowazekii str. Rp22 |
| 1105099 | Rickettsia prowazekii str. RpGvF24 |
| 1105113 | Rickettsia rhipicephali str. 3-7-female6-CWPP |

| NCBI Taxon Identifier | Organism Name |
|---|---|
| 1105105 | Rickettsia rickettsii str. Arizona |
| 1105104 | Rickettsia rickettsii str. Brazil |
| 1105102 | Rickettsia rickettsii str. Colombia |
| 1105103 | Rickettsia rickettsii str. Hauke |
| 1105100 | Rickettsia rickettsii str. Hino |
| 1105101 | Rickettsia rickettsii str. Hlp#2 |
| 452659 | Rickettsia rickettsii str. Iowa |
| 1337396 | Rickettsia rickettsii str. Morgan |
| 1338411 | Rickettsia rickettsii str. R |
| 392021 | Rickettsia rickettsii str. 'Sheila Smith' |
| 941638 | Rickettsia slovaca 13-B |
| 1105109 | Rickettsia slovaca str. D-CWPP |
| 1182263 | Rickettsia sp. MEAM1 (Bemisia tabaci) |
| 1003202 | Rickettsia typhi str. B9991CWPP |
| 1003201 | Rickettsia typhi str. TH1527 |
| 257363 | Rickettsia typhi str. Wilmington |
| 1528098 | Rickettsiales bacterium Ac37b |
| 2486578 | Rickettsiales endosymbiont of Stachyamoeba lipophora |
| 693978 | Riemerella anatipestifer ATCC 11845 = DSM 15868 |
| 693978 | Riemerella anatipestifer ATCC 11845 = DSM 15868 |
| 1354240 | Riemerella anatipestifer CH3 |
| 1228997 | Riemerella anatipestifer RA-CH-1 |
| 1271752 | Riemerella anatipestifer RA-CH-2 |
| 992406 | Riemerella anatipestifer RA-GD |
| 41431 | Rippkaea orientalis PCC 8801 |
| 395962 | Rippkaea orientalis PCC 8802 |
| 373994 | Rivularia sp. PCC 7116 |
| 313596 | Robiginitalea biformata HTCC2501 |
| 758 | Rodentibacter pneumotropicus |
| 76731 | Roseateles depolymerans |
| 585394 | Roseburia hominis A2-183 |
| 657315 | Roseburia intestinalis M50/1 |
| 718255 | Roseburia intestinalis XB6B4 |
| 1294273 | Roseibacterium elongatum DSM 19469 |
| 187304 | Roseibium aggregatum |
| 383372 | Roseiflexus castenholzii DSM 13941 |
| 357808 | Roseiflexus sp. RS-1 |
| 441209 | Roseinatronobacter bogoriensis subsp. barguzinensis |
| 1852022 | Roseitalea porphyridii |
| 375451 | Roseobacter denitrificans OCh 114 |
| 391595 | Roseobacter litoralis Och 149 |

| NCBI Taxon Identifier | Organism Name |
|---|---|
| 257708 | Roseomonas gilardii |
| 2018065 | Roseomonas sp. FDAARGOS_362 |
| 215743 | Roseovarius mucosus |
| 2203213 | Roseovarius sp. AK1035 |
| 172042 | Rothia aeria |
| 762948 | Rothia dentocariosa ATCC 17931 |
| 680646 | Rothia mucilaginosa DY-18 |
| 756272 | Rubinisphaera brasiliensis DSM 5305 |
| 987059 | Rubrivivax benzoatilyticus JA2 = ATCC BAA-35 |
| 983917 | Rubrivivax gelatinosus IL144 |
| 42256 | Rubrobacter radiotolerans |
| 266117 | Rubrobacter xylanophilus DSM 9941 |
| 246200 | Ruegeria pomeroyi DSS-3 |
| 2293862 | Ruegeria sp. AD91A |
| 292414 | Ruegeria sp. TM1040 |
| 1379910 | Rufibacter radiotolerans |
| 1379909 | Rufibacter sp. DG15C |
| 512763 | Rufibacter tibetensis |
| 1565605 | Rugosibacter aromaticivorans |
| 394503 | Ruminiclostridium cellulolyticum H10 |
| 1572656 | Ruminococcaceae bacterium CPB6 |
| 697329 | Ruminococcus albus 7 = DSM 20455 |
| 1160721 | Ruminococcus bicirculans (ex Wegman et al. 2014) |
| 213810 | Ruminococcus champanellensis 18P13 = JCM 17042 |
| 657323 | Ruminococcus sp. SR1/5 |
| 241244 | Rummeliibacillus stabekisii |
| 2259595 | Runella rosea |
| 761193 | Runella slithyformis DSM 19594 |
| 2268026 | Runella sp. SP2 |
| 2172099 | Saccharobesus litoralis |
| 2287 | Saccharolobus solfataricus |
| 2287 | Saccharolobus solfataricus |
| 2287 | Saccharolobus solfataricus |
| 555311 | Saccharolobus solfataricus 98/2 |
| 273057 | Saccharolobus solfataricus P2 |
| 471857 | Saccharomonospora viridis DSM 43017 |
| 559292 | Saccharomyces cerevisiae S288C |
| 203122 | Saccharophagus degradans 2-40 |
| 405948 | Saccharopolyspora erythraea NRRL 2338 |
| 1179773 | Saccharothrix espanaensis DSM 44229 |
| 10224 | Saccoglossus kowalevskii |

| NCBI Taxon Identifier | Organism Name |
|---|---|
| 2009329 | Sagittula sp. P11 |
| 39432 | Saimiri boliviensis boliviensis |
| 1729720 | Salegentibacter sp. T436 |
| 2099786 | Salicibibacter kimchii |
| 1230341 | Salimicrobium jeotgali |
| 1333523 | Salinarchaeum sp. Harcht-Bsk1 |
| 309807 | Salinibacter ruber DSM 13855 |
| 761659 | Salinibacter ruber M8 |
| 2079791 | Salinibacterium hongtaonis |
| 2508880 | Salinibacterium sp. UTAS2018 |
| 407035 | Salinicoccus halodurans |
| 755307 | Salinigranum rubrum |
| 2303538 | Salinimonas sediminis |
| 2183582 | Saliniradius amylolyticus |
| 2183911 | Salinisphaera sp. LB1 |
| 1307761 | Salinispira pacifica |
| 391037 | Salinispora arenicola CNS-205 |
| 369723 | Salinispora tropica CNB-440 |
| 1307839 | Salinivirga cyanobacteriivorans |
| 1250539 | Salipiger abyssi |
| 1229727 | Salipiger profundus |
| 1792508 | Salipiger sp. CCB-MM3 |
| 632773 | Salisediminibacterium beveridgei |
| 8030 | Salmo salar |
| 1197719 | Salmonella bongori N268-08 |
| 218493 | Salmonella bongori NCTC 12419 |
| 1382510 | Salmonella bongori serovar 48:z41:-- str. RKS3044 |
| 41514 | Salmonella enterica subsp. arizonae serovar 62:z4,z23:- |
| 866913 | Salmonella enterica subsp. enterica serovar 4,[5],12:i:- str. 08-1736 |
| 1406860 | Salmonella enterica subsp. enterica serovar Agona str. 24249 |
| 454166 | Salmonella enterica subsp. enterica serovar Agona str. SL483 |
| 1173427 | Salmonella enterica subsp. enterica serovar Bareilly str. CFSAN000189 |
| 1320309 | Salmonella enterica subsp. enterica serovar Bovismorbificans str. 3114 |
| 321314 | Salmonella enterica subsp. enterica serovar Choleraesuis str. SC-B67 |
| 1271863 | Salmonella enterica subsp. enterica serovar Cubana str. CFSAN002050 |
| 439851 | Salmonella enterica subsp. enterica serovar Dublin str. CT_02021853 |
| 149539 | Salmonella enterica subsp. enterica serovar Enteritidis |
| 1412460 | Salmonella enterica subsp. enterica serovar Enteritidis str. EC20090135 |
| 1412459 | Salmonella enterica subsp. enterica serovar Enteritidis str. EC20090193 |
| 1412461 | Salmonella enterica subsp. enterica serovar Enteritidis str. EC20090332 |
| 1412462 | Salmonella enterica subsp. enterica serovar Enteritidis str. EC20090531 |

| NCBI Taxon Identifier | Organism Name |
|---|---|
| 550537 | Salmonella enterica subsp. enterica serovar Enteritidis str. P125109 |
| 550538 | Salmonella enterica subsp. enterica serovar Gallinarum str. 287/91 |
| 1225522 | Salmonella enterica subsp. enterica serovar Gallinarum/Pullorum str. CDC1983-67 |
| 1081093 | Salmonella enterica subsp. enterica serovar Gallinarum/Pullorum str. RKS5078 |
| 1124936 | Salmonella enterica subsp. enterica serovar Heidelberg str. 41578 |
| 1160717 | Salmonella enterica subsp. enterica serovar Heidelberg str. B182 |
| 1271864 | Salmonella enterica subsp. enterica serovar Heidelberg str. CFSAN002069 |
| 454169 | Salmonella enterica subsp. enterica serovar Heidelberg str. SL476 |
| 1267753 | Salmonella enterica subsp. enterica serovar Javiana str. CFSAN001992 |
| 423368 | Salmonella enterica subsp. enterica serovar Newport str. SL254 |
| 877468 | Salmonella enterica subsp. enterica serovar Newport str. USMARC-S3124.1 |
| 554290 | Salmonella enterica subsp. enterica serovar Paratyphi A str. AKU_12601 |
| 295319 | Salmonella enterica subsp. enterica serovar Paratyphi A str. ATCC 9150 |
| 1016998 | Salmonella enterica subsp. enterica serovar Paratyphi B str. SPB7 |
| 476213 | Salmonella enterica subsp. enterica serovar Paratyphi C str. RKS4594 |
| 1298917 | Salmonella enterica subsp. enterica serovar Pullorum str. S06004 |
| 439843 | Salmonella enterica subsp. enterica serovar Schwarzengrund str. CVM19633 |
| 1003191 | Salmonella enterica subsp. enterica serovar Tennessee str. TXSC_TXSC08-19 |
| 1064551 | Salmonella enterica subsp. enterica serovar Thompson str. RM6836 |
| 220341 | Salmonella enterica subsp. enterica serovar Typhi str. CT18 |
| 1132507 | Salmonella enterica subsp. enterica serovar Typhi str. P-stx-12 |
| 209261 | Salmonella enterica subsp. enterica serovar Typhi str. Ty2 |
| 527001 | Salmonella enterica subsp. enterica serovar Typhi str. Ty21a |
| 90371 | Salmonella enterica subsp. enterica serovar Typhimurium |
| 588858 | Salmonella enterica subsp. enterica serovar Typhimurium str. 14028S |
| 1008297 | Salmonella enterica subsp. enterica serovar Typhimurium str. 798 |
| 568708 | Salmonella enterica subsp. enterica serovar Typhimurium str. D23580 |
| 85569 | Salmonella enterica subsp. enterica serovar Typhimurium str. DT104 |
| 568709 | Salmonella enterica subsp. enterica serovar Typhimurium str. DT2 |
| 99287 | Salmonella enterica subsp. enterica serovar Typhimurium str. LT2 |
| 216597 | Salmonella enterica subsp. enterica serovar Typhimurium str. SL1344 |
| 909946 | Salmonella enterica subsp. enterica serovar Typhimurium str. ST4/74 |
| 718274 | Salmonella enterica subsp. enterica serovar Typhimurium str. T000240 |
| 1171376 | Salmonella enterica subsp. enterica serovar Typhimurium str. U288 |
| 990282 | Salmonella enterica subsp. enterica serovar Typhimurium str. UK-1 |
| 1271862 | Salmonella enterica subsp. enterica serovar Typhimurium var. 5- str. CFSAN001921 |
| 946362 | Salpingoeca rosetta |
| 8036 | Salvelinus alpinus |
| 927083 | Sandaracinus amylolyticus |
| 446469 | Sanguibacter keddieii DSM 10542 |
| 695850 | Saprolegnia parasitica CBS 223.65 |

| NCBI Taxon Identifier | Organism Name |
|---|---|
| 984262 | Saprospira grandis str. Lewin |
| 9305 | Sarcophilus harrisii |
| 1150468 | Scardovia inopinata JCM 12537 |
| 52773 | Schaalia meyeri |
| 322104 | Scheffersomyces stipitis CBS 6054 |
| 6185 | Schistosoma haematobium |
| 6183 | Schistosoma mansoni |
| 578458 | Schizophyllum commune H4-8 |
| 284812 | Schizosaccharomyces pombe 972h- |
| 113540 | Scleropages formosus |
| 665079 | Sclerotinia sclerotiorum 1980 UF-70 |
| 526218 | Sebaldella termitidis ATCC 33386 |
| 1199245 | secondary endosymbiont of Ctenarytaina eucalypti |
| 134287 | secondary endosymbiont of Heteropsylla cubana |
| 1835721 | secondary endosymbiont of Trabutina mannipara |
| 240427 | Secundilactobacillus paracollinoides |
| 1543721 | Sedimenticola thiotaurini |
| 1940790 | Sedimentisphaera cyanobacteriorum |
| 1941349 | Sedimentisphaera salicampi |
| 1453352 | Sediminicola sp. YIK13 |
| 573413 | Sediminispirochaeta smaragdinae DSM 11293 |
| 640132 | Segniliparus rotundus DSM 44985 |
| 88036 | Selaginella moellendorffii |
| 927704 | Selenomonas ruminantium subsp. lactilytica TAM6421 |
| 713030 | Selenomonas sp. oral taxon 136 |
| 712538 | Selenomonas sp. oral taxon 478 |
| 1884263 | Selenomonas sp. oral taxon 920 |
| 546271 | Selenomonas sputigena ATCC 35185 |
| 1936081 | Seonamhaeicola sp. S2-3 |
| 1758689 | Serinicoccus hydrothermalis |
| 9135 | Serinus canaria |
| 41447 | Seriola dumerili |
| 1458426 | Serpentinimonas maccroryi |
| 1458425 | Serpentinimonas raichei |
| 578457 | Serpula lacrymans var. lacrymans S7.9 |
| 47917 | Serratia fonticola |
| 47917 | Serratia fonticola |
| 1154756 | Serratia inhibens PRI-2C |
| 1346614 | Serratia liquefaciens ATCC 27592 |
| 1334564 | Serratia marcescens SM39 |
| 273526 | Serratia marcescens subsp. marcescens Db11 |

| NCBI Taxon Identifier | Organism Name |
|---|---|
| 435998 | Serratia marcescens WW4 |
| 682634 | Serratia plymuthica 4Rx13 |
| 768492 | Serratia plymuthica AS9 |
| 1348660 | Serratia plymuthica S13 |
| 399741 | Serratia proteamaculans 568 |
| 61652 | Serratia rubidaea |
| 768490 | Serratia sp. AS12 |
| 768493 | Serratia sp. AS13 |
| 671990 | Serratia sp. FGI94 |
| 1327989 | Serratia sp. FS14 |
| 2033438 | Serratia sp. MYb239 |
| 488142 | Serratia sp. SCBI |
| 568817 | Serratia symbiotica str. 'Cinara cedri' |
| 4182 | Sesamum indicum |
| 4555 | Setaria italica |
| 38313 | Shewanella algae |
| 326297 | Shewanella amazonensis SB2B |
| 693974 | Shewanella baltica BA175 |
| 693970 | Shewanella baltica OS117 |
| 325240 | Shewanella baltica OS155 |
| 402882 | Shewanella baltica OS185 |
| 399599 | Shewanella baltica OS195 |
| 407976 | Shewanella baltica OS223 |
| 693973 | Shewanella baltica OS678 |
| 2018305 | Shewanella bicestrii |
| 318161 | Shewanella denitrificans OS217 |
| 318167 | Shewanella frigidimarina NCIMB 400 |
| 458817 | Shewanella halifaxensis HAW-EB4 |
| 93973 | Shewanella japonica |
| 150120 | Shewanella livingstonensis |
| 323850 | Shewanella loihica PV-4 |
| 260364 | Shewanella marisflavi |
| 211586 | Shewanella oneidensis MR-1 |
| 398579 | Shewanella pealeana ATCC 700345 |
| 225849 | Shewanella piezotolerans WP3 |
| 225848 | Shewanella psychrophila |
| 399804 | Shewanella putrefaciens 200 |
| 319224 | Shewanella putrefaciens CN-32 |
| 425104 | Shewanella sediminis HAW-EB3 |
| 94122 | Shewanella sp. ANA-3 |
| 1930557 | Shewanella sp. FDAARGOS_354 |

| NCBI Taxon Identifier | Organism Name |
|---|---|
| 60480 | Shewanella sp. MR-4 |
| 60481 | Shewanella sp. MR-7 |
| 351745 | Shewanella sp. W3-18-1 |
| 2029986 | Shewanella sp. WE21 |
| 637905 | Shewanella violacea DSS12 |
| 392500 | Shewanella woodyi ATCC 51908 |
| 344609 | Shigella boydii CDC 3083-94 |
| 300268 | Shigella boydii Sb227 |
| 754093 | Shigella dysenteriae 1617 |
| 300267 | Shigella dysenteriae Sd197 |
| 623 | Shigella flexneri |
| 591020 | Shigella flexneri 2002017 |
| 1282357 | Shigella flexneri 2003036 |
| 198215 | Shigella flexneri 2a str. 2457T |
| 198214 | Shigella flexneri 2a str. 301 |
| 373384 | Shigella flexneri 5 str. 8401 |
| 1282358 | Shigella flexneri Shi06HN006 |
| 300269 | Shigella sonnei Ss046 |
| 1813821 | Shigella sp. PAMC 28760 |
| 630626 | Shimwellia blattae DSM 4481 = NBRC 105725 |
| 879274 | Shinella sp. HZN7 |
| 66692 | Shouchella clausii KSM-K16 |
| 1246626 | Shouchella lehensis G1 |
| 1454006 | Siansivirga zeaxanthinifaciens CC-SAMT-1 |
| 580332 | Sideroxydans lithotrophicus ES-1 |
| 1826607 | Silicimonas algicola |
| 1117647 | Simiduia agarivorans SA1 = DSM 21679 |
| 331113 | Simkania negevensis Z |
| 641147 | Simonsiella muelleri ATCC 29453 |
| 2109915 | Simplicispira suum |
| 886293 | Singulisphaera acidiphila DSM 18658 |
| 1608454 | Sinocyclocheilus anshuiensis |
| 75366 | Sinocyclocheilus grahami |
| 307959 | Sinocyclocheilus rhinocerous |
| 37927 | Sinomonas atrocyanea |
| 194963 | Sinorhizobium americanum |
| 1117943 | Sinorhizobium fredii HH103 |
| 394 | Sinorhizobium fredii NGR234 |
| 1185652 | Sinorhizobium fredii USDA 257 |
| 366394 | Sinorhizobium medicae WSM419 |
| 382 | Sinorhizobium meliloti |

| NCBI Taxon Identifier | Organism Name |
|---|---|
| 266834 | Sinorhizobium meliloti 1021 |
| 1286640 | Sinorhizobium meliloti 2011 |
| 693982 | Sinorhizobium meliloti AK83 |
| 698936 | Sinorhizobium meliloti BL225C |
| 1235461 | Sinorhizobium meliloti GR4 |
| 1230587 | Sinorhizobium meliloti Rm41 |
| 707241 | Sinorhizobium meliloti SM11 |
| 716928 | Sinorhizobium sojae CCBAU 05684 |
| 794846 | Sinorhizobium sp. CCBAU 05631 |
| 1842534 | Sinorhizobium sp. RAC02 |
| 471855 | Slackia heliotrinireducens DSM 20476 |
| 187101 | Sneathia vaginalis |
| 1196094 | Snodgrassella alvi wkB2 |
| 1929246 | Sodalis endosymbiont of Henestaris halophilus |
| 343509 | Sodalis glossinidius str. 'morsitans' |
| 1239307 | Sodalis praecaptivus |
| 4081 | Solanum lycopersicum |
| 28526 | Solanum pennellii |
| 4113 | Solanum tuberosum |
| 13686 | Solenopsis invicta |
| 76853 | Solibacillus silvestris |
| 1002809 | Solibacillus silvestris StLB046 |
| 2048654 | Solibacillus sp. R5-41 |
| 573370 | Solidesulfovibrio magneticus RS-1 |
| 2303331 | Solimonas sp. K1W22B-7 |
| 929556 | Solitalea canadensis DSM 3403 |
| 448385 | Sorangium cellulosum So ce56 |
| 1254432 | Sorangium cellulosum So0157-2 |
| 771870 | Sordaria macrospora k-hell |
| 4558 | Sorghum bicolor |
| 619300 | Spathaspora passalidarum NRRL Y-27907 |
| 479434 | Sphaerobacter thermophilus DSM 20745 |
| 158189 | Sphaerochaeta globosa str. Buddy |
| 158190 | Sphaerochaeta pleomorpha str. Grapes |
| 1986952 | Sphingobacteriaceae bacterium GW460-11-11-14-LB5 |
| 1010 | Sphingobacterium mizutaii |
| 743722 | Sphingobacterium sp. 21 |
| 1933220 | Sphingobacterium sp. B29 |
| 1538644 | Sphingobacterium sp. ML3W |
| 1332080 | Sphingobium baderi |
| 690566 | Sphingobium chlorophenolicum L-1 |

| NCBI Taxon Identifier | Organism Name |
|---|---|
| 120107 | Sphingobium cloacae |
| 861109 | Sphingobium indicum B90A |
| 452662 | Sphingobium indicum UT26S |
| 1855519 | Sphingobium sp. EP60837 |
| 407020 | Sphingobium sp. MI1205 |
| 1843368 | Sphingobium sp. RAC03 |
| 627192 | Sphingobium sp. SYK-6 |
| 1315974 | Sphingobium sp. TKS |
| 484429 | Sphingobium sp. YBL2 |
| 2082188 | Sphingobium sp. YG1 |
| 121428 | Sphingobium xenophagum |
| 13690 | Sphingobium yanoikuyae |
| 1609977 | Sphingomonas hengshuiensis |
| 93064 | Sphingomonas koreensis |
| 621456 | Sphingomonas melonis TY |
| 2319844 | Sphingomonas paeninsulae |
| 1560345 | Sphingomonas panacis |
| 1327635 | Sphingomonas psychrotolerans |
| 1123269 | Sphingomonas sanxanigenens DSM 19645 = NX02 |
| 2219696 | Sphingomonas sp. FARSPH |
| 1030157 | Sphingomonas sp. KC8 |
| 1390395 | Sphingomonas sp. LK11 |
| 1938607 | Sphingomonas sp. LM7 |
| 745310 | Sphingomonas sp. MM-1 |
| 1961362 | Sphingomonas sp. NIC1 |
| 1549858 | Sphingomonas taxi |
| 317655 | Sphingopyxis alaskensis RB2256 |
| 1515612 | Sphingopyxis fribergensis |
| 267128 | Sphingopyxis granuli |
| 33050 | Sphingopyxis macrogoltabida |
| 33050 | Sphingopyxis macrogoltabida |
| 292913 | Sphingopyxis sp. 113P3 |
| 1866325 | Sphingopyxis sp. MG |
| 1357916 | Sphingopyxis sp. QXT-31 |
| 1219058 | Sphingopyxis terrae subsp. terrae NBRC 15098 |
| 1913578 | Sphingorhabdus lutea |
| 1806885 | Sphingorhabdus sp. M41 |
| 2077182 | Sphingorhabdus sp. YGSMI21 |
| 335406 | Sphingosinicella microcystinivorans |
| 1892855 | Sphingosinicella sp. BN140058 |
| 3562 | Spinacia oleracea |

| NCBI Taxon Identifier | Organism Name |
|---|---|
| 1335757 | Spiribacter curvatus |
| 1260251 | Spiribacter salinus M19-40 |
| 889378 | Spirochaeta africana DSM 8902 |
| 665571 | Spirochaeta thermophila DSM 6192 |
| 869211 | Spirochaeta thermophila DSM 6578 |
| 1276258 | Spiroplasma apis B31 |
| 1114980 | Spiroplasma atrichopogonis |
| 362837 | Spiroplasma cantharicola |
| 1276227 | Spiroplasma chrysopicola DF-1 |
| 2133 | Spiroplasma citri |
| 2139 | Spiroplasma clarkii |
| 216934 | Spiroplasma corruscae |
| 1276246 | Spiroplasma culicicola AES-1 |
| 1276221 | Spiroplasma diminutum CUAS-1 |
| 315358 | Spiroplasma eriocheiris |
| 1336749 | Spiroplasma floricola 23-6 |
| 216938 | Spiroplasma helicoides |
| 273035 | Spiroplasma kunkelii CR2-3x |
| 216942 | Spiroplasma litorale |
| 838561 | Spiroplasma mirum ATCC 29335 |
| 838561 | Spiroplasma mirum ATCC 29335 |
| 1276257 | Spiroplasma sabaudiense Ar-1343 |
| 1914410 | Spiroplasma sp. NBRC 100390 |
| 1276229 | Spiroplasma syrphidicola EA-1 |
| 1276220 | Spiroplasma taiwanense CT-1 |
| 216946 | Spiroplasma turonicum |
| 504472 | Spirosoma linguale DSM 74 |
| 1178516 | Spirosoma montaniterrae |
| 2057025 | Spirosoma pollinicola |
| 1379870 | Spirosoma radiotolerans |
| 1620392 | Spongiibacter sp. IMCC21906 |
| 269673 | Sporolactobacillus terrae |
| 1476 | Sporosarcina psychrophila |
| 1930764 | Sporosarcina sp. P33 |
| 1930546 | Sporosarcina sp. P37 |
| 2283194 | Sporosarcina sp. PTS2304 |
| 1571 | Sporosarcina ureae |
| 1397361 | Sporothrix schenckii 1099-18 |
| 446470 | Stackebrandtia nassauensis DSM 44728 |
| 111780 | Stanieria cyanosphaera PCC 7437 |
| 1807358 | Stanieria sp. NIES-3757 |

| NCBI Taxon Identifier | Organism Name |
|---|---|
| 985762 | Staphylococcus agnetis |
| 985002 | Staphylococcus argenteus |
| 1280 | Staphylococcus aureus |
| 1280 | Staphylococcus aureus |
| 1280 | Staphylococcus aureus |
| 703339 | Staphylococcus aureus 04-02981 |
| 1229492 | Staphylococcus aureus 08BA02176 |
| 1321369 | Staphylococcus aureus Bmb9393 |
| 1323661 | Staphylococcus aureus CA-347 |
| 1305598 | Staphylococcus aureus M1 |
| 273036 | Staphylococcus aureus RF122 |
| 1123523 | Staphylococcus aureus subsp. aureus 11819-97 |
| 585143 | Staphylococcus aureus subsp. aureus 55/2053 |
| 1392476 | Staphylococcus aureus subsp. aureus 6850 |
| 1155084 | Staphylococcus aureus subsp. aureus 71193 |
| 1193576 | Staphylococcus aureus subsp. aureus CN1 |
| 93062 | Staphylococcus aureus subsp. aureus COL |
| 889933 | Staphylococcus aureus subsp. aureus ECT-R 2 |
| 685039 | Staphylococcus aureus subsp. aureus ED133 |
| 681288 | Staphylococcus aureus subsp. aureus ED98 |
| 1074252 | Staphylococcus aureus subsp. aureus HO 5096 0412 |
| 359787 | Staphylococcus aureus subsp. aureus JH1 |
| 359786 | Staphylococcus aureus subsp. aureus JH9 |
| 869816 | Staphylococcus aureus subsp. aureus JKD6159 |
| 985006 | Staphylococcus aureus subsp. aureus LGA251 |
| 1118959 | Staphylococcus aureus subsp. aureus M013 |
| 282458 | Staphylococcus aureus subsp. aureus MRSA252 |
| 282459 | Staphylococcus aureus subsp. aureus MSSA476 |
| 418127 | Staphylococcus aureus subsp. aureus Mu3 |
| 158878 | Staphylococcus aureus subsp. aureus Mu50 |
| 196620 | Staphylococcus aureus subsp. aureus MW2 |
| 158879 | Staphylococcus aureus subsp. aureus N315 |
| 93061 | Staphylococcus aureus subsp. aureus NCTC 8325 |
| 1368166 | Staphylococcus aureus subsp. aureus SA268 |
| 1194085 | Staphylococcus aureus subsp. aureus SA40 |
| 1201010 | Staphylococcus aureus subsp. aureus SA957 |
| 1074919 | Staphylococcus aureus subsp. aureus ST228 |
| 1074919 | Staphylococcus aureus subsp. aureus ST228 |
| 1074919 | Staphylococcus aureus subsp. aureus ST228 |
| 1074919 | Staphylococcus aureus subsp. aureus ST228 |
| 1074919 | Staphylococcus aureus subsp. aureus ST228 |

| NCBI Taxon Identifier | Organism Name |
|---|---|
| 1074919 | Staphylococcus aureus subsp. aureus ST228 |
| 1074919 | Staphylococcus aureus subsp. aureus ST228 |
| 1074919 | Staphylococcus aureus subsp. aureus ST228 |
| 523796 | Staphylococcus aureus subsp. aureus ST398 |
| 546342 | Staphylococcus aureus subsp. aureus str. JKD6008 |
| 426430 | Staphylococcus aureus subsp. aureus str. Newman |
| 1006543 | Staphylococcus aureus subsp. aureus T0131 |
| 548473 | Staphylococcus aureus subsp. aureus TCH60 |
| 663951 | Staphylococcus aureus subsp. aureus TW20 |
| 451515 | Staphylococcus aureus subsp. aureus USA300_FPR3757 |
| 451516 | Staphylococcus aureus subsp. aureus USA300_TCH1516 |
| 1028799 | Staphylococcus aureus subsp. aureus VC40 |
| 1406863 | Staphylococcus aureus subsp. aureus Z172 |
| 1458279 | Staphylococcus aureus USA300-ISMMS1 |
| 72758 | Staphylococcus capitis subsp. capitis |
| 396513 | Staphylococcus carnosus subsp. carnosus TM300 |
| 29382 | Staphylococcus cohnii |
| 70255 | Staphylococcus condimenti |
| 1282 | Staphylococcus epidermidis |
| 176280 | Staphylococcus epidermidis ATCC 12228 |
| 1449752 | Staphylococcus epidermidis PM221 |
| 176279 | Staphylococcus epidermidis RP62A |
| 246432 | Staphylococcus equorum |
| 46127 | Staphylococcus felis |
| 1283 | Staphylococcus haemolyticus |
| 279808 | Staphylococcus haemolyticus JCSC1435 |
| 1284 | Staphylococcus hyicus |
| 29384 | Staphylococcus kloosii |
| 698737 | Staphylococcus lugdunensis HKU09-01 |
| 1034809 | Staphylococcus lugdunensis N920143 |
| 155085 | Staphylococcus lutrae |
| 214473 | Staphylococcus nepalensis |
| 1276282 | Staphylococcus pasteuri SP1 |
| 170573 | Staphylococcus pettenkoferi |
| 984892 | Staphylococcus pseudintermedius ED99 |
| 937773 | Staphylococcus pseudintermedius HKU10-03 |
| 342451 | Staphylococcus saprophyticus subsp. saprophyticus ATCC 15305 = NCTC 7292 |
| 1295 | Staphylococcus schleiferi |
| 1295 | Staphylococcus schleiferi |
| 1286 | Staphylococcus simulans |
| 1194526 | Staphylococcus warneri SG1 |

| NCBI Taxon Identifier | Organism Name |
|---|---|
| 1288 | Staphylococcus xylosus |
| 1288 | Staphylococcus xylosus |
| 1288 | Staphylococcus xylosus |
| 591019 | Staphylothermus hellenicus DSM 12710 |
| 399550 | Staphylothermus marinus F1 |
| 128780 | Stenotrophomonas acidaminiphila |
| 1163399 | Stenotrophomonas maltophilia D457 |
| 868597 | Stenotrophomonas maltophilia JV3 |
| 522373 | Stenotrophomonas maltophilia K279a |
| 391008 | Stenotrophomonas maltophilia R551-3 |
| 216778 | Stenotrophomonas rhizophila |
| 1793721 | Stenotrophomonas sp. KCTC 12332 |
| 1904944 | Stenotrophomonas sp. LM091 |
| 1827305 | Stenotrophomonas sp. MYb57 |
| 2005046 | Stenotrophomonas sp. WZN-1 |
| 721885 | Stereum hirsutum FP-91666 SS1 |
| 465721 | Steroidobacter denitrificans |
| 378806 | Stigmatella aurantiaca DW4/3-1 |
| 980422 | Strawberry lethal yellows phytoplasma (CPA) str. NZSb11 |
| 1003195 | Streptantibioticus cattleyicolor NRRL 8057 = DSM 46488 |
| 1003195 | Streptantibioticus cattleyicolor NRRL 8057 = DSM 46488 |
| 519441 | Streptobacillus moniliformis DSM 12112 |
| 1311 | Streptococcus agalactiae |
| 1311 | Streptococcus agalactiae |
| 1311 | Streptococcus agalactiae |
| 1311 | Streptococcus agalactiae |
| 1309806 | Streptococcus agalactiae 09mas018883 |
| 1417990 | Streptococcus agalactiae 138P |
| 1261567 | Streptococcus agalactiae 2-22 |
| 208435 | Streptococcus agalactiae 2603V/R |
| 205921 | Streptococcus agalactiae A909 |
| 1427374 | Streptococcus agalactiae CNCTC 10/84 |
| 342616 | Streptococcus agalactiae COH1 |
| 1203670 | Streptococcus agalactiae GD201008-001 |
| 1309807 | Streptococcus agalactiae ILRI005 |
| 1318615 | Streptococcus agalactiae ILRI112 |
| 211110 | Streptococcus agalactiae NEM316 |
| 1328 | Streptococcus anginosus |
| 862970 | Streptococcus anginosus C1051 |
| 862971 | Streptococcus anginosus C238 |
| 862969 | Streptococcus constellatus subsp. pharyngis C1050 |

| NCBI Taxon Identifier | Organism Name |
|---|---|
| 696216 | Streptococcus constellatus subsp. pharyngis C232 |
| 862968 | Streptococcus constellatus subsp. pharyngis C818 |
| 1302863 | Streptococcus cristatus AS 1.3089 |
| 1247189 | Streptococcus dysgalactiae subsp. equisimilis 167 |
| 759913 | Streptococcus dysgalactiae subsp. equisimilis AC-2713 |
| 663954 | Streptococcus dysgalactiae subsp. equisimilis ATCC 12394 |
| 486410 | Streptococcus dysgalactiae subsp. equisimilis GGS_124 |
| 617121 | Streptococcus dysgalactiae subsp. equisimilis RE378 |
| 553482 | Streptococcus equi subsp. equi 4047 |
| 40041 | Streptococcus equi subsp. zooepidemicus |
| 1051072 | Streptococcus equi subsp. zooepidemicus ATCC 35246 |
| 1403449 | Streptococcus equi subsp. zooepidemicus CY |
| 552526 | Streptococcus equi subsp. zooepidemicus MGCS10565 |
| 981539 | Streptococcus gallolyticus subsp. gallolyticus ATCC 43143 |
| 990317 | Streptococcus gallolyticus subsp. gallolyticus ATCC BAA-2069 |
| 637909 | Streptococcus gallolyticus UCN34 |
| 467705 | Streptococcus gordonii str. Challis substr. CH1 |
| 1156431 | Streptococcus ilei |
| 1156431 | Streptococcus ilei |
| 1069533 | Streptococcus infantarius subsp. infantarius CJ18 |
| 1346 | Streptococcus iniae |
| 1346 | Streptococcus iniae |
| 1346 | Streptococcus iniae |
| 1318633 | Streptococcus iniae SF1 |
| 862967 | Streptococcus intermedius B196 |
| 862966 | Streptococcus intermedius C270 |
| 591365 | Streptococcus intermedius JTH08 |
| 1076934 | Streptococcus lutetiensis 033 |
| 1116231 | Streptococcus macedonicus ACA-DC 198 |
| 365659 | Streptococcus mitis B6 |
| 1198676 | Streptococcus mutans GS-5 |
| 1155071 | Streptococcus mutans LJ23 |
| 511691 | Streptococcus mutans NN2025 |
| 210007 | Streptococcus mutans UA159 |
| 1437447 | Streptococcus mutans UA159-FR |
| 927666 | Streptococcus oralis Uo5 |
| 1811193 | Streptococcus pantholopis |
| 760570 | Streptococcus parasanguinis ATCC 15912 |
| 1114965 | Streptococcus parasanguinis FW213 |
| 936154 | Streptococcus parauberis KCTC 11537 |
| 981540 | Streptococcus pasteurianus ATCC 43144 |

| NCBI Taxon Identifier | Organism Name |
|---|---|
| 189423 | Streptococcus pneumoniae 670-6B |
| 488221 | Streptococcus pneumoniae 70585 |
| 1408179 | Streptococcus pneumoniae A026 |
| 574093 | Streptococcus pneumoniae AP200 |
| 561276 | Streptococcus pneumoniae ATCC 700669 |
| 516950 | Streptococcus pneumoniae CGSP14 |
| 373153 | Streptococcus pneumoniae D39 |
| 512566 | Streptococcus pneumoniae G54 |
| 697283 | Streptococcus pneumoniae gamPNI0373 |
| 487214 | Streptococcus pneumoniae Hungary19A-6 |
| 869269 | Streptococcus pneumoniae INV104 |
| 869216 | Streptococcus pneumoniae INV200 |
| 488222 | Streptococcus pneumoniae JJA |
| 869215 | Streptococcus pneumoniae OXC141 |
| 488223 | Streptococcus pneumoniae P1031 |
| 171101 | Streptococcus pneumoniae R6 |
| 869303 | Streptococcus pneumoniae SPN034156 |
| 869304 | Streptococcus pneumoniae SPN034183 |
| 869306 | Streptococcus pneumoniae SPN994038 |
| 869307 | Streptococcus pneumoniae SPN994039 |
| 869309 | Streptococcus pneumoniae SPNA45 |
| 1130804 | Streptococcus pneumoniae ST556 |
| 487213 | Streptococcus pneumoniae Taiwan19F-14 |
| 525381 | Streptococcus pneumoniae TCH8431/19A |
| 170187 | Streptococcus pneumoniae TIGR4 |
| 1054460 | Streptococcus pseudopneumoniae IS7493 |
| 1235829 | Streptococcus pyogenes A20 |
| 487215 | Streptococcus pyogenes Alab49 |
| 1336746 | Streptococcus pyogenes HSC5 |
| 1207470 | Streptococcus pyogenes M1 476 |
| 160490 | Streptococcus pyogenes M1 GAS |
| 370552 | Streptococcus pyogenes MGAS10270 |
| 286636 | Streptococcus pyogenes MGAS10394 |
| 370554 | Streptococcus pyogenes MGAS10750 |
| 798300 | Streptococcus pyogenes MGAS15252 |
| 1010840 | Streptococcus pyogenes MGAS1882 |
| 370553 | Streptococcus pyogenes MGAS2096 |
| 198466 | Streptococcus pyogenes MGAS315 |
| 293653 | Streptococcus pyogenes MGAS5005 |
| 319701 | Streptococcus pyogenes MGAS6180 |
| 186103 | Streptococcus pyogenes MGAS8232 |

| NCBI Taxon Identifier | Organism Name |
|---|---|
| 370551 | Streptococcus pyogenes MGAS9429 |
| 471876 | Streptococcus pyogenes NZ131 |
| 193567 | Streptococcus pyogenes SSI-1 |
| 1440772 | Streptococcus pyogenes STAB901 |
| 160491 | Streptococcus pyogenes str. Manfredo |
| 1304 | Streptococcus salivarius |
| 1304 | Streptococcus salivarius |
| 1046629 | Streptococcus salivarius 57.I |
| 1048332 | Streptococcus salivarius CCHSS3 |
| 347253 | Streptococcus salivarius JIM8777 |
| 388919 | Streptococcus sanguinis SK36 |
| 1310 | Streptococcus sobrinus |
| 1759399 | Streptococcus sp. A12 |
| 1902136 | Streptococcus sp. NPS 308 |
| 1419814 | Streptococcus sp. VT 162 |
| 391295 | Streptococcus suis 05ZYH33 |
| 391296 | Streptococcus suis 98HAH33 |
| 993512 | Streptococcus suis A7 |
| 568814 | Streptococcus suis BM407 |
| 1004952 | Streptococcus suis D12 |
| 1005042 | Streptococcus suis D9 |
| 423211 | Streptococcus suis GZ1 |
| 945704 | Streptococcus suis JS14 |
| 218494 | Streptococcus suis P1/7 |
| 1184252 | Streptococcus suis S735 |
| 1246365 | Streptococcus suis SC070731 |
| 568813 | Streptococcus suis SC84 |
| 1005041 | Streptococcus suis SS12 |
| 1004951 | Streptococcus suis ST1 |
| 1007064 | Streptococcus suis ST3 |
| 1340847 | Streptococcus suis T15 |
| 1276647 | Streptococcus suis TL13 |
| 1380773 | Streptococcus suis YB51 |
| 1308 | Streptococcus thermophilus |
| 1408178 | Streptococcus thermophilus ASCC 1275 |
| 299768 | Streptococcus thermophilus CNRZ1066 |
| 1051074 | Streptococcus thermophilus JIM 8232 |
| 322159 | Streptococcus thermophilus LMD-9 |
| 264199 | Streptococcus thermophilus LMG 18311 |
| 1187956 | Streptococcus thermophilus MN-ZLW-002 |
| 767463 | Streptococcus thermophilus ND03 |

| NCBI Taxon Identifier | Organism Name |
|---|---|
| 218495 | Streptococcus uberis 0140J |
| 2498135 | Streptomonospora litoralis |
| 1886 | Streptomyces albidoflavus |
| 1940 | Streptomyces albireticuli |
| 67267 | Streptomyces alboflavus |
| 1888 | Streptomyces albus |
| 1642299 | Streptomyces alfalfae |
| 278992 | Streptomyces ambofaciens ATCC 23877 |
| 75293 | Streptomyces autolyticus |
| 227882 | Streptomyces avermitilis MA-4680 = NBRC 14893 |
| 749414 | Streptomyces bingchenggensis BCW-1 |
| 1901 | Streptomyces clavuligerus |
| 100226 | Streptomyces coelicolor A3(2) |
| 1214242 | Streptomyces collinus Tu 365 |
| 477245 | Streptomyces cyaneogriseus subsp. noncyanogenus |
| 1214101 | Streptomyces davaonensis JCM 4913 |
| 1616117 | Streptomyces formicae |
| 553510 | Streptomyces gilvosporeus |
| 1907 | Streptomyces glaucescens |
| 1172567 | Streptomyces globisporus C-1027 |
| 68214 | Streptomyces griseochromogenes |
| 455632 | Streptomyces griseus subsp. griseus NBRC 13350 |
| 1133850 | Streptomyces hygroscopicus subsp. jinggangensis 5008 |
| 1203460 | Streptomyces hygroscopicus subsp. jinggangensis TL01 |
| 39478 | Streptomyces laurentii |
| 58340 | Streptomyces lavendulae subsp. lavendulae |
| 1437453 | Streptomyces leeuwenhoekii |
| 1915 | Streptomyces lincolnensis |
| 457428 | Streptomyces lividans TK24 |
| 47763 | Streptomyces lydicus |
| 47763 | Streptomyces lydicus |
| 92644 | Streptomyces malaysiensis |
| 1303692 | Streptomyces microflavus DSM 40593 |
| 1827580 | Streptomyces nigra |
| 193462 | Streptomyces niveus |
| 1971 | Streptomyces noursei |
| 316284 | Streptomyces noursei ATCC 11455 |
| 1434306 | Streptomyces noursei ZPM |
| 146923 | Streptomyces parvulus |
| 1355015 | Streptomyces pluripotens |
| 591167 | Streptomyces pratensis ATCC 33331 |

| NCBI Taxon Identifier | Organism Name |
|---|---|
| 38300 | Streptomyces pristinaespiralis |
| 164348 | Streptomyces puniciscabiei |
| 1783515 | Streptomyces qaidamensis |
| 1343740 | Streptomyces rapamycinicus NRRL 5491 |
| 1926 | Streptomyces reticuli |
| 285473 | Streptomyces rubrolavendulae |
| 680198 | Streptomyces scabiei 87.22 |
| 1751294 | Streptomyces sp. 4F |
| 1262452 | Streptomyces sp. 769 |
| 1561022 | Streptomyces sp. CCM_MD2014 |
| 1725411 | Streptomyces sp. CdTB01 |
| 1649184 | Streptomyces sp. CFMR 7 |
| 444103 | Streptomyces sp. CNQ-509 |
| 465541 | Streptomyces sp. Mg1 |
| 1961713 | Streptomyces sp. MOE7 |
| 1265601 | Streptomyces sp. PAMC 26508 |
| 1849967 | Streptomyces sp. SAT1 |
| 862751 | Streptomyces sp. SirexAA-E |
| 953739 | Streptomyces venezuelae ATCC 10712 |
| 362257 | Streptomyces vietnamensis |
| 1935 | Streptomyces violaceoruber |
| 653045 | Streptomyces violaceusniger Tu 4113 |
| 408015 | Streptomyces xiamenensis |
| 479432 | Streptosporangium roseum DSM 43021 |
| 7668 | Strongylocentrotus purpuratus |
| 1123016 | Stutzerimonas balearica DSM 6083 |
| 316 | Stutzerimonas stutzeri |
| 316 | Stutzerimonas stutzeri |
| 379731 | Stutzerimonas stutzeri A1501 |
| 96563 | Stutzerimonas stutzeri ATCC 17588 = LMG 11199 |
| 1196835 | Stutzerimonas stutzeri CCUG 29243 |
| 1123519 | Stutzerimonas stutzeri DSM 10701 |
| 996285 | Stutzerimonas stutzeri DSM 4166 |
| 644801 | Stutzerimonas stutzeri RCH2 |
| 50429 | Stylophora pistillata |
| 796027 | Sugiyamaella lignohabitans |
| 2036206 | Suicoccus acidiformans |
| 1917485 | Sulfitobacter alexandrii |
| 421000 | Sulfitobacter donghicola |
| 1968541 | Sulfitobacter sp. D7 |
| 2070369 | Sulfitobacter sp. JL08 |

| NCBI Taxon Identifier | Organism Name |
|---|---|
| 1389005 | Sulfitobacter sp. SK012 |
| 679936 | Sulfobacillus acidophilus DSM 10332 |
| 1051632 | Sulfobacillus acidophilus TPY |
| 1670455 | Sulfodiicoccus acidiphilus |
| 330779 | Sulfolobus acidocaldarius DSM 639 |
| 1028566 | Sulfolobus acidocaldarius N8 |
| 1028567 | Sulfolobus acidocaldarius Ron12/I |
| 1435377 | Sulfolobus acidocaldarius SUSAZ |
| 930943 | Sulfolobus islandicus HVE10/4 |
| 425944 | Sulfolobus islandicus L.D.8.5 |
| 429572 | Sulfolobus islandicus L.S.2.15 |
| 1241935 | Sulfolobus islandicus LAL14/1 |
| 427317 | Sulfolobus islandicus M.14.25 |
| 427318 | Sulfolobus islandicus M.16.27 |
| 426118 | Sulfolobus islandicus M.16.4 |
| 930945 | Sulfolobus islandicus REY15A |
| 439386 | Sulfolobus islandicus Y.G.57.14 |
| 419942 | Sulfolobus islandicus Y.N.15.51 |
| 1891280 | Sulfolobus sp. A20 |
| 1620215 | Sulfuricaulis limicola |
| 1163617 | Sulfuricella denitrificans skB26 |
| 709032 | Sulfuricurvum kujiense DSM 16994 |
| 1985873 | Sulfuriferula sp. AH1 |
| 1675686 | Sulfurifustis variabilis |
| 204536 | Sulfurihydrogenibium azorense Az-Fu1 |
| 436114 | Sulfurihydrogenibium sp. YO3AOP1 |
| 563040 | Sulfurimonas autotrophica DSM 16294 |
| 326298 | Sulfurimonas denitrificans DSM 1251 |
| 273063 | Sulfurisphaera tokodaii str. 7 |
| 1223802 | Sulfuritalea hydrogenivorans sk43H |
| 760154 | Sulfurospirillum barnesii SES-3 |
| 525898 | Sulfurospirillum deleyianum DSM 6946 |
| 1854492 | Sulfurospirillum diekertiae |
| 1854492 | Sulfurospirillum diekertiae |
| 1193502 | Sulfurospirillum halorespirans DSM 13726 |
| 1150621 | Sulfurospirillum multivorans DSM 12446 |
| 206403 | Sulfurovum lithotrophicum |
| 387093 | Sulfurovum sp. NBC37-1 |
| 9823 | Sus scrofa |
| 79883 | Sutcliffiella horikoshii |
| 2494234 | Sutterella megalosphaeroides |

| NCBI Taxon Identifier | Organism Name |
|---|---|
| 292459 | Symbiobacterium thermophilum IAM 14863 |
| 269084 | Synechococcus elongatus PCC 6301 |
| 1140 | Synechococcus elongatus PCC 7942 = FACHB-805 |
| 1350461 | Synechococcus elongatus UTEX 2973 |
| 64471 | Synechococcus sp. CC9311 |
| 110662 | Synechococcus sp. CC9605 |
| 316279 | Synechococcus sp. CC9902 |
| 321332 | Synechococcus sp. JA-2-3B'a(2-13) |
| 321327 | Synechococcus sp. JA-3-3Ab |
| 1280380 | Synechococcus sp. KORDI-100 |
| 585423 | Synechococcus sp. KORDI-49 |
| 585425 | Synechococcus sp. KORDI-52 |
| 195253 | Synechococcus sp. PCC 6312 |
| 1173263 | Synechococcus sp. PCC 7502 |
| 316278 | Synechococcus sp. RCC307 |
| 32051 | Synechococcus sp. WH 7803 |
| 29410 | Synechococcus sp. WH 8103 |
| 166314 | Synechococcus sp. WH 8109 |
| 2116702 | Synechocystis sp. IPPAS B-1465 |
| 1147 | Synechocystis sp. PCC 6714 |
| 1148 | Synechocystis sp. PCC 6803 |
| 1148 | Synechocystis sp. PCC 6803 |
| 1148 | Synechocystis sp. PCC 6803 |
| 1080228 | Synechocystis sp. PCC 6803 substr. GT-I |
| 1080229 | Synechocystis sp. PCC 6803 substr. PCC-N |
| 1080230 | Synechocystis sp. PCC 6803 substr. PCC-P |
| 335543 | Syntrophobacter fumaroxidans MPOB |
| 645991 | Syntrophobotulus glycolicus DSM 8271 |
| 335541 | Syntrophomonas wolfei subsp. wolfei str. Goettingen G311 |
| 29542 | Syntrophotalea acetylenica |
| 1842532 | Syntrophotalea acetylenivorans |
| 338963 | Syntrophotalea carbinolica DSM 2380 |
| 643648 | Syntrophothermus lipocalidus DSM 12680 |
| 56780 | Syntrophus aciditrophicus SB |
| 2494374 | Tabrizicola piscis |
| 59729 | Taeniopygia guttata |
| 31033 | Takifugu rubripes |
| 2069432 | Tamlana carrageenivorans |
| 203275 | Tannerella forsythia 92A2 |
| 712710 | Tannerella serpentiformis |
| 1921510 | Tardibacter chloracetimidivorans |

| NCBI Taxon Identifier | Organism Name |
|---|---|
| 28532 | Tarenaya hassleriana |
| 299262 | Tateyamaria omphalii |
| 53336 | Tatumella citrea |
| 82987 | Tatumella ptyseos |
| 1091495 | Taylorella asinigenitalis 14/45 |
| 1008459 | Taylorella asinigenitalis MCE3 |
| 1091497 | Taylorella equigenitalis 14/56 |
| 743973 | Taylorella equigenitalis ATCC 35865 |
| 937774 | Taylorella equigenitalis MCE9 |
| 669041 | Tenacibaculum dicentrarchi |
| 584609 | Tenacibaculum jejuense |
| 1349785 | Tenacibaculum maritimum NCIMB 2154 |
| 1850252 | Tenacibaculum todarodis |
| 1911684 | Tenericutes bacterium MO-XQ |
| 1911683 | Tenericutes bacterium MZ-XQ |
| 1209989 | Tepidanaerobacter acetatoxydans Re1 |
| 1209989 | Tepidanaerobacter acetatoxydans Re1 |
| 377629 | Teredinibacter turnerae T7901 |
| 386490 | Terribacillus goriensis |
| 926566 | Terriglobus roseus DSM 18391 |
| 401053 | Terriglobus saanensis SP1PR4 |
| 1332264 | Tessaracoccus aquimaris |
| 399497 | Tessaracoccus flavescens |
| 1610493 | Tessaracoccus flavus |
| 1909732 | Tessaracoccus sp. T2.5-30 |
| 945021 | Tetragenococcus halophilus NBRC 12172 |
| 290335 | Tetragenococcus koreensis |
| 526944 | Tetragenococcus osmophilus |
| 312017 | Tetrahymena thermophila SB210 |
| 32264 | Tetranychus urticae |
| 99883 | Tetraodon nigroviridis |
| 1071381 | Tetrapisispora phaffii CBS 4417 |
| 296543 | Thalassiosira pseudonana CCMP1335 |
| 2017482 | Thalassococcus sp. S3 |
| 1298593 | Thalassolituus oleivorans MIL-1 |
| 1208320 | Thalassolituus oleivorans R6-15 |
| 1891279 | Thalassospira indica |
| 2048283 | Thalassospira marina |
| 1123366 | Thalassospira xiamenensis M-5 = DSM 17429 |
| 164330 | Thauera aminoaromatica |
| 44139 | Thauera aromatica K172 |

| NCBI Taxon Identifier | Organism Name |
|---|---|
| 96773 | Thauera chlorobenzoica |
| 1134435 | Thauera humireducens |
| 2005884 | Thauera sp. K11 |
| 353154 | Theileria annulata strain Ankara |
| 1537102 | Theileria equi strain WA |
| 869250 | Theileria orientalis strain Shintoku |
| 333668 | Theileria parva strain Muguga |
| 3641 | Theobroma cacao |
| 1089553 | Thermacetogenium phaeum DSM 12270 |
| 644966 | Thermaerobacter marianensis DSM 12885 |
| 525903 | Thermanaerovibrio acidaminovorans DSM 6589 |
| 635013 | Thermincola potens JR |
| 2026 | Thermoactinomyces vulgaris |
| 509193 | Thermoanaerobacter brockii subsp. finnii Ako-1 |
| 580331 | Thermoanaerobacter italicus Ab9 |
| 2325 | Thermoanaerobacter kivui |
| 583358 | Thermoanaerobacter mathranii subsp. mathranii str. A3 |
| 340099 | Thermoanaerobacter pseudethanolicus ATCC 33223 |
| 573062 | Thermoanaerobacter sp. X513 |
| 399726 | Thermoanaerobacter sp. X514 |
| 697303 | Thermoanaerobacter wiegelii Rt8.B1 |
| 1094508 | Thermoanaerobacterium saccharolyticum JW/SL-YS485 |
| 580327 | Thermoanaerobacterium thermosaccharolyticum DSM 571 |
| 698948 | Thermoanaerobacterium thermosaccharolyticum M0795 |
| 858215 | Thermoanaerobacterium xylanolyticum LX-11 |
| 717605 | Thermobacillus composti KWC4 |
| 525904 | Thermobaculum terrenum ATCC BAA-798 |
| 269800 | Thermobifida fusca YX |
| 469371 | Thermobispora bispora DSM 43833 |
| 759272 | Thermochaetoides thermophila DSM 1495 |
| 1121335 | Thermoclostridium stercorarium subsp. stercorarium DSM 8532 |
| 1121335 | Thermoclostridium stercorarium subsp. stercorarium DSM 8532 |
| 391623 | Thermococcus barophilus MP |
| 54077 | Thermococcus barossii |
| 1293037 | Thermococcus celer Vu 13 = JCM 8558 |
| 54262 | Thermococcus chitonophagus |
| 163003 | Thermococcus cleftensis |
| 1505907 | Thermococcus eurythermalis |
| 593117 | Thermococcus gammatolerans EJ3 |
| 71997 | Thermococcus gorgonarius |
| 1432656 | Thermococcus guaymasensis DSM 11113 |

| NCBI Taxon Identifier | Organism Name |
|---|---|
| 69014 | Thermococcus kodakarensis KOD1 |
| 523849 | Thermococcus litoralis DSM 5473 |
| 195522 | Thermococcus nautili |
| 523850 | Thermococcus onnurineus NA1 |
| 71998 | Thermococcus pacificus |
| 582419 | Thermococcus paralvinellae |
| 53952 | Thermococcus peptonophilus |
| 1712654 | Thermococcus piezophilus |
| 49899 | Thermococcus profundus |
| 187880 | Thermococcus radiotolerans |
| 604354 | Thermococcus sibiricus MM 739 |
| 72803 | Thermococcus siculi |
| 1674923 | Thermococcus sp. 2319x1 |
| 1042877 | Thermococcus sp. 4557 |
| 2008440 | Thermococcus sp. 5-4 |
| 246969 | Thermococcus sp. AM4 |
| 122420 | Thermococcus sp. P6 |
| 277988 | Thermococcus thioreducens |
| 932678 | Thermocrinis albus DSM 12173 |
| 638303 | Thermocrinis albus DSM 14484 |
| 667014 | Thermodesulfatator indicus DSM 15286 |
| 289377 | Thermodesulfobacterium commune DSM 2178 |
| 795359 | Thermodesulfobacterium geofontis OPF15 |
| 1794699 | Thermodesulfobium acidiphilum |
| 747365 | Thermodesulfobium narugense DSM 14796 |
| 289376 | Thermodesulfovibrio yellowstonii DSM 11347 |
| 1365176 | Thermofilum adornatum |
| 697581 | Thermofilum adornatum 1505 |
| 368408 | Thermofilum pendens Hrk 5 |
| 1550241 | Thermofilum uzonense |
| 1184251 | Thermogladius calderae 1633 |
| 1331910 | Thermogutta terrifontis |
| 309801 | Thermomicrobium roseum DSM 5159 |
| 471852 | Thermomonospora curvata DSM 43183 |
| 273075 | Thermoplasma acidophilum DSM 1728 |
| 273116 | Thermoplasma volcanium GSS1 |
| 1054217 | Thermoplasmatales archaeon BRNA1 |
| 768679 | Thermoproteus tenax Kra 1 |
| 999630 | Thermoproteus uzoniensis 768-20 |
| 555079 | Thermosediminibacter oceani DSM 16646 |
| 484019 | Thermosipho africanus TCF52B |

| NCBI Taxon Identifier | Organism Name |
|---|---|
| 391009 | Thermosipho melanesiensis BI429 |
| 1462747 | Thermosipho sp. 1063 |
| 1437364 | Thermosipho sp. 1070 |
| 633148 | Thermosphaera aggregans DSM 11486 |
| 1917166 | Thermostichus lividus PCC 6715 |
| 1298851 | Thermosulfidibacter takaii ABI70S6 |
| 1394889 | Thermosynechococcus sp. NK55a |
| 197221 | Thermosynechococcus vestitus BP-1 |
| 573729 | Thermothelomyces thermophilus ATCC 42464 |
| 578455 | Thermothielavioides terrestris NRRL 8126 |
| 2336 | Thermotoga maritima |
| 2336 | Thermotoga maritima |
| 243274 | Thermotoga maritima MSB8 |
| 243274 | Thermotoga maritima MSB8 |
| 243274 | Thermotoga maritima MSB8 |
| 243274 | Thermotoga maritima MSB8 |
| 309803 | Thermotoga neapolitana DSM 4359 |
| 390874 | Thermotoga petrophila RKU-1 |
| 590168 | Thermotoga petrophila RKU-10 |
| 1157948 | Thermotoga sp. 2812B |
| 1157947 | Thermotoga sp. Cell2 |
| 126740 | Thermotoga sp. RQ2 |
| 126738 | Thermotoga sp. RQ7 |
| 648996 | Thermovibrio ammonificans HB-1 |
| 580340 | Thermovirga lienii DSM 17291 |
| 498848 | Thermus aquaticus Y51MC23 |
| 56956 | Thermus brockianus |
| 751945 | Thermus oshimai JL-2 |
| 456163 | Thermus parvatiensis |
| 743525 | Thermus scotoductus SA-01 |
| 1111069 | Thermus sp. CCB_US3_UF1 |
| 262724 | Thermus thermophilus HB27 |
| 300852 | Thermus thermophilus HB8 |
| 798128 | Thermus thermophilus JL-18 |
| 762633 | Thermus thermophilus SG0.5JP17-16 |
| 1255043 | Thioalkalivibrio nitratireducens DSM 14787 |
| 713585 | Thioalkalivibrio paradoxus ARh 1 |
| 396595 | Thioalkalivibrio sp. K90mix |
| 396588 | Thioalkalivibrio sulfidiphilus HL-EbGr7 |
| 106634 | Thioalkalivibrio versutus |
| 292415 | Thiobacillus denitrificans ATCC 25259 |

| NCBI Taxon Identifier | Organism Name |
|---|---|
| 1915078 | Thioclava nitratireducens |
| 765911 | Thiocystis violascens DSM 198 |
| 765912 | Thioflavicoccus mobilis 8321 |
| 585455 | Thiohalobacter thiocyanaticus |
| 1076588 | Thiolapillus brandeum |
| 717772 | Thiomicrospira aerophila AL3 |
| 717773 | Thiomicrospira cyclica ALM1 |
| 1803865 | Thiomicrospira sp. S5 |
| 426114 | Thiomonas arsenitoxydans |
| 75379 | Thiomonas intermedia K12 |
| 40754 | Thioploca ingrica |
| 1697053 | Thiopseudomonas alkaliphila |
| 1110502 | Tistrella mobilis KA081020-065 |
| 595494 | Tolumonas auensis DSM 9187 |
| 4950 | Torulaspora delbrueckii |
| 508771 | Toxoplasma gondii ME49 |
| 717944 | Trametes versicolor FP-101664 SS1 |
| 578456 | Tremella mesenterica DSM 1558 |
| 906968 | Treponema brennaborense DSM 12168 |
| 243275 | Treponema denticola ATCC 35405 |
| 1155775 | Treponema pallidum str. Fribourg-Blanc |
| 491081 | Treponema pallidum subsp. pallidum DAL-1 |
| 455434 | Treponema pallidum subsp. pallidum SS14 |
| 666714 | Treponema pallidum subsp. pallidum str. Chicago |
| 686990 | Treponema pallidum subsp. pallidum str. Mexico A |
| 243276 | Treponema pallidum subsp. pallidum str. Nichols |
| 243276 | Treponema pallidum subsp. pallidum str. Nichols |
| 1095955 | Treponema pallidum subsp. pallidum str. Sea 81-4 |
| 491079 | Treponema pallidum subsp. pertenue str. CDC2 |
| 491080 | Treponema pallidum subsp. pertenue str. Gauthier |
| 491078 | Treponema pallidum subsp. pertenue str. SamoaD |
| 545776 | Treponema paraluiscuniculi Cuniculi A |
| 1291379 | Treponema pedis str. T A4 |
| 545694 | Treponema primitia ZAS-2 |
| 221027 | Treponema putidum |
| 1539298 | Treponema sp. OMZ 838 |
| 869209 | Treponema succinifaciens DSM 2489 |
| 7070 | Tribolium castaneum |
| 127582 | Trichechus manatus latirostris |
| 6334 | Trichinella spiralis |
| 398767 | Trichlorobacter lovleyi SZ |

| NCBI Taxon Identifier | Organism Name |
|---|---|
| 431241 | Trichoderma reesei QM6a |
| 203124 | Trichodesmium erythraeum IMS101 |
| 412133 | Trichomonas vaginalis G3 |
| 663331 | Trichophyton benhamiae CBS 112371 |
| 663202 | Trichophyton verrucosum HKI 0517 |
| 10228 | Trichoplax adhaerens |
| 7111 | Trichoplusia ni |
| 240292 | Trichormus variabilis ATCC 29413 |
| 1265309 | Tritonibacter mobilis F1926 |
| 203267 | Tropheryma whipplei str. Twist |
| 218496 | Tropheryma whipplei TW08/27 |
| 649638 | Truepera radiovictrix DSM 17093 |
| 1661 | Trueperella pyogenes |
| 1435056 | Trueperella pyogenes TP8 |
| 185431 | Trypanosoma brucei brucei TREU927 |
| 353153 | Trypanosoma cruzi strain CL Brener |
| 521096 | Tsukamurella paurometabola DSM 20162 |
| 57704 | Tsukamurella tyrosinosolvens |
| 692370 | Tsuneonella dongtanensis |
| 656061 | Tuber melanosporum Mel28 |
| 1214604 | Tumebacillus algifaecis |
| 1903704 | Tumebacillus avium |
| 246437 | Tupaia chinensis |
| 1712675 | Turicibacter sp. H121 |
| 869212 | Turneriella parva DSM 21527 |
| 336963 | Uncinocarpus reesii 1704 |
| 310581 | uncultured Sphingopyxis sp. |
| 401471 | Undibacterium parvum |
| 38504 | Ureaplasma parvum serovar 3 |
| 505682 | Ureaplasma parvum serovar 3 str. ATCC 27815 |
| 273119 | Ureaplasma parvum serovar 3 str. ATCC 700970 |
| 565575 | Ureaplasma urealyticum serovar 10 str. ATCC 33699 |
| 1850246 | Urechidicola croceus |
| 29073 | Ursus maritimus |
| 237631 | Ustilago maydis 521 |
| 633807 | Vagococcus penaei |
| 519472 | Vagococcus teuberi |
| 578460 | Vairimorpha ceranae BRL01 |
| 436907 | Vanderwaltozyma polyspora DSM 70294 |
| 1333996 | Variibacter gotjawalensis |
| 436515 | Variovorax boronicumulans |

| NCBI Taxon Identifier | Organism Name |
|---|---|
| 1246301 | Variovorax paradoxus B4 |
| 595537 | Variovorax paradoxus EPS |
| 543728 | Variovorax paradoxus S110 |
| 1795631 | Variovorax sp. PAMC 28711 |
| 2126319 | Variovorax sp. PMC12 |
| 39777 | Veillonella atypica |
| 39778 | Veillonella dispar |
| 479436 | Veillonella parvula DSM 2008 |
| 248315 | Veillonella rodentium |
| 391735 | Verminephrobacter eiseniae EF01-2 |
| 1637999 | Verrucomicrobia bacterium IMCC26134 |
| 2026799 | Verrucomicrobiota bacterium |
| 526221 | Verticillium alfalfae VaMs.102 |
| 498257 | Verticillium dahliae VdLs.17 |
| 1074311 | Vibrio alfacsensis |
| 1219076 | Vibrio alginolyticus NBRC 15630 = ATCC 17749 |
| 55601 | Vibrio anguillarum |
| 882102 | Vibrio anguillarum 775 |
| 882944 | Vibrio anguillarum M3 |
| 150340 | Vibrio antiquarius |
| 575788 | Vibrio atlanticus LGP32 |
| 553239 | Vibrio breoganii |
| 1224742 | Vibrio campbellii CAIM 519 = NBRC 15631 = ATCC 25920 |
| 1224742 | Vibrio campbellii CAIM 519 = NBRC 15631 = ATCC 25920 |
| 666 | Vibrio cholerae |
| 666 | Vibrio cholerae |
| 666 | Vibrio cholerae |
| 1134456 | Vibrio cholerae IEC224 |
| 935297 | Vibrio cholerae LMA3984-4 |
| 579112 | Vibrio cholerae M66-2 |
| 593588 | Vibrio cholerae MJ-1236 |
| 1420885 | Vibrio cholerae MS6 |
| 686 | Vibrio cholerae O1 biovar El Tor |
| 243277 | Vibrio cholerae O1 biovar El Tor str. N16961 |
| 914149 | Vibrio cholerae O1 str. 2010EL-1786 |
| 345073 | Vibrio cholerae O395 |
| 345073 | Vibrio cholerae O395 |
| 190893 | Vibrio coralliilyticus |
| 190893 | Vibrio coralliilyticus |
| 50719 | Vibrio diabolicus |
| 676 | Vibrio fluvialis |

| NCBI Taxon Identifier | Organism Name |
|---|---|
| 903510 | Vibrio furnissii NCTC 11218 |
| 687 | Vibrio gazogenes |
| 669 | Vibrio harveyi |
| 689 | Vibrio mediterranei |
| 674 | Vibrio mimicus |
| 1219067 | Vibrio natriegens NBRC 15636 = ATCC 14048 = DSM 759 |
| 28173 | Vibrio nigripulchritudo |
| 696485 | Vibrio owensii |
| 1211705 | Vibrio parahaemolyticus BB22OP |
| 1338032 | Vibrio parahaemolyticus O1:K33 str. CDC_K4557 |
| 1338034 | Vibrio parahaemolyticus O1:Kuk str. FDA_R31 |
| 223926 | Vibrio parahaemolyticus RIMD 2210633 |
| 1429044 | Vibrio parahaemolyticus UCM-V493 |
| 2025808 | Vibrio qinghaiensis |
| 190895 | Vibrio rotiferianus |
| 45658 | Vibrio scophthalmi |
| 1116375 | Vibrio sp. EJY3 |
| 1671868 | Vibrio tapetis subsp. tapetis |
| 1051646 | Vibrio tubiashii ATCC 19109 |
| 672 | Vibrio vulnificus |
| 216895 | Vibrio vulnificus CMCP6 |
| 914127 | Vibrio vulnificus MO6-24/O |
| 196600 | Vibrio vulnificus YJ016 |
| 2094242 | Victivallales bacterium CCUG 44730 |
| 3914 | Vigna angularis |
| 3916 | Vigna radiata var. radiata |
| 302167 | Virgibacillus dokdonensis |
| 1482 | Virgibacillus halodenitrificans |
| 163877 | Virgibacillus necropolis |
| 2017483 | Virgibacillus phasianinus |
| 1911587 | Virgibacillus sp. 6R |
| 403957 | Virgibacillus sp. SK37 |
| 29760 | Vitis vinifera |
| 63 | Vitreoscilla filiformis |
| 96942 | Vitreoscilla sp. C1 |
| 3068 | Volvox carteri f. nagariensis |
| 272774 | Vreelandella alkaliphila |
| 44935 | Vreelandella venusta |
| 572478 | Vulcanisaeta distributa DSM 14429 |
| 985053 | Vulcanisaeta moutnovskia 768-28 |
| 1391653 | Vulgatibacter incomptus |

| NCBI Taxon Identifier | Organism Name |
|---|---|
| 716544 | Waddlia chondrophila WSU 86-1044 |
| 1299270 | Wallemia ichthyophaga EXF-994 |
| 671144 | Wallemia mellicola CBS 633.66 |
| 865938 | Weeksella virosa DSM 16922 |
| 759620 | Weissella ceti |
| 759620 | Weissella ceti |
| 759620 | Weissella ceti |
| 137591 | Weissella cibaria |
| 1631871 | Weissella jogaejeotgali |
| 1045854 | Weissella koreensis KACC 15510 |
| 1249 | Weissella paramesenteroides |
| 1790137 | Wenyingzhuangia fucanilytica |
| 1579979 | Wenzhouxiangella marina |
| 36870 | Wigglesworthia glossinidia endosymbiont of Glossina brevipalpis |
| 1142511 | Wigglesworthia glossinidia endosymbiont of Glossina morsitans morsitans (Yale colony) |
| 214888 | Williamsoniiplasma luminosum |
| 215578 | Williamsoniiplasma somnilux |
| 1936080 | Winogradskyella sp. J14-2 |
| 754409 | Winogradskyella sp. PG-2 |
| 1548547 | Woeseia oceani |
| 246273 | Wolbachia endosymbiont of Cimex lectularius |
| 570417 | Wolbachia endosymbiont of Culex quinquefasciatus Pel |
| 163164 | Wolbachia endosymbiont of Drosophila melanogaster |
| 1236909 | Wolbachia endosymbiont of Drosophila simulans wHa |
| 1236908 | Wolbachia endosymbiont of Drosophila simulans wNo |
| 169402 | Wolbachia endosymbiont of Folsomia candida |
| 100901 | Wolbachia endosymbiont of Onchocerca ochengi |
| 292805 | Wolbachia endosymbiont strain TRS of Brugia malayi |
| 1116230 | Wolbachia pipientis wAlbB |
| 66084 | Wolbachia sp. wRi |
| 273121 | Wolinella succinogenes DSM 1740 |
| 78245 | Xanthobacter autotrophicus Py2 |
| 380358 | Xanthomonas albilineans GPE PC73 |
| 1304892 | Xanthomonas axonopodis Xac29-1 |
| 340 | Xanthomonas campestris pv. campestris |
| 314565 | Xanthomonas campestris pv. campestris str. 8004 |
| 190485 | Xanthomonas campestris pv. campestris str. ATCC 33913 |
| 990315 | Xanthomonas campestris pv. raphani 756C |
| 611301 | Xanthomonas citri pv. citri |
| 611301 | Xanthomonas citri pv. citri |
| 611301 | Xanthomonas citri pv. citri |

| NCBI Taxon Identifier | Organism Name |
|---|---|
| 611301 | Xanthomonas citri pv. citri |
| 611301 | Xanthomonas citri pv. citri |
| 611301 | Xanthomonas citri pv. citri |
| 190486 | Xanthomonas citri pv. citri str. 306 |
| 366649 | Xanthomonas citri pv. fuscans |
| 1308541 | Xanthomonas citri subsp. citri A306 |
| 1137651 | Xanthomonas citri subsp. citri Aw12879 |
| 1308548 | Xanthomonas citri subsp. citri UI6 |
| 981368 | Xanthomonas euvesicatoria pv. citrumelo F1 |
| 316273 | Xanthomonas euvesicatoria pv. vesicatoria str. 85-10 |
| 48664 | Xanthomonas fragariae |
| 56454 | Xanthomonas hortorum |
| 2754056 | Xanthomonas hortorum pv. gardneri |
| 291331 | Xanthomonas oryzae pv. oryzae KACC 10331 |
| 342109 | Xanthomonas oryzae pv. oryzae MAFF 311018 |
| 1458476 | Xanthomonas oryzae pv. oryzae PXO86 |
| 360094 | Xanthomonas oryzae pv. oryzae PXO99A |
| 129394 | Xanthomonas oryzae pv. oryzicola |
| 383407 | Xanthomonas oryzae pv. oryzicola BLS256 |
| 442694 | Xanthomonas perforans |
| 317013 | Xanthomonas phaseoli pv. phaseoli |
| 56458 | Xanthomonas sacchari |
| 487909 | Xanthomonas translucens pv. undulosa |
| 325776 | Xanthomonas vasicola pv. vasculorum |
| 925775 | Xanthomonas vesicatoria ATCC 35937 |
| 8355 | Xenopus laevis |
| 8364 | Xenopus tropicalis |
| 40576 | Xenorhabdus bovienii |
| 406818 | Xenorhabdus bovienii SS-2004 |
| 351671 | Xenorhabdus doucetiae |
| 351679 | Xenorhabdus hominickii |
| 1437823 | Xenorhabdus nematophila AN6/1 |
| 406817 | Xenorhabdus nematophila ATCC 19061 |
| 1354304 | Xenorhabdus poinarii G6 |
| 8083 | Xiphophorus maculatus |
| 264731 | Xylanibacter ruminicola 23 |
| 2509459 | Xylanimonas allomyrinae |
| 446471 | Xylanimonas cellulosilytica DSM 15894 |
| 2509457 | Xylanimonas protaetiae |
| 2371 | Xylella fastidiosa |
| 160492 | Xylella fastidiosa 9a5c |

| NCBI Taxon Identifier | Organism Name |
|---|---|
| 405440 | Xylella fastidiosa M12 |
| 405441 | Xylella fastidiosa M23 |
| 1401256 | Xylella fastidiosa MUL0034 |
| 788929 | Xylella fastidiosa subsp. fastidiosa GB514 |
| 155920 | Xylella fastidiosa subsp. sandyi Ann-1 |
| 183190 | Xylella fastidiosa Temecula1 |
| 1444770 | Xylella taiwanensis |
| 590646 | Yamadazyma tenuis ATCC 10573 |
| 284591 | Yarrowia lipolytica CLIB122 |
| 1453495 | Yersinia aldovae 670-83 |
| 263819 | Yersinia aleksiciae |
| 630 | Yersinia enterocolitica |
| 630 | Yersinia enterocolitica |
| 630 | Yersinia enterocolitica |
| 1262462 | Yersinia enterocolitica (type O:5) str. YE53/03 |
| 1443113 | Yersinia enterocolitica LC20 |
| 393305 | Yersinia enterocolitica subsp. enterocolitica 8081 |
| 994476 | Yersinia enterocolitica subsp. palearctica 105.5R(r) |
| 930944 | Yersinia enterocolitica subsp. palearctica Y11 |
| 1454377 | Yersinia frederiksenii Y225 |
| 631 | Yersinia intermedia |
| 28152 | Yersinia kristensenii |
| 419257 | Yersinia massiliensis |
| 632 | Yersinia pestis |
| 632 | Yersinia pestis |
| 632 | Yersinia pestis |
| 632 | Yersinia pestis |
| 1035377 | Yersinia pestis A1122 |
| 349746 | Yersinia pestis Angola |
| 360102 | Yersinia pestis Antiqua |
| 547048 | Yersinia pestis biovar Medievalis str. Harbin 35 |
| 229193 | Yersinia pestis biovar Microtus str. 91001 |
| 214092 | Yersinia pestis CO92 |
| 637382 | Yersinia pestis D106004 |
| 637385 | Yersinia pestis D182038 |
| 187410 | Yersinia pestis KIM10+ |
| 377628 | Yersinia pestis Nepal516 |
| 386656 | Yersinia pestis Pestoides F |
| 637386 | Yersinia pestis Z176003 |
| 633 | Yersinia pseudotuberculosis |
| 633 | Yersinia pseudotuberculosis |

| NCBI Taxon Identifier | Organism Name |
|---|---|
| 633 | Yersinia pseudotuberculosis |
| 633 | Yersinia pseudotuberculosis |
| 349747 | Yersinia pseudotuberculosis IP 31758 |
| 273123 | Yersinia pseudotuberculosis IP 32953 |
| 273123 | Yersinia pseudotuberculosis IP 32953 |
| 502801 | Yersinia pseudotuberculosis PB1/+ |
| 748672 | Yersinia pseudotuberculosis str. PA3606 |
| 502800 | Yersinia pseudotuberculosis YPIII |
| 29485 | Yersinia rohdei |
| 29486 | Yersinia ruckeri |
| 29486 | Yersinia ruckeri |
| 367190 | Yersinia similis |
| 245188 | Yoonia vestfoldensis |
| 4577 | Zea mays |
| 1470434 | Zhongshania aliphaticivorans |
| 326968 | Ziziphus jujuba |
| 347534 | Zobellella denitrificans |
| 63186 | Zobellia galactanivorans |
| 2080469 | Zoogloeaceae bacteirum Par-f-2 |
| 136037 | Zootermopsis nevadensis |
| 655815 | Zunongwangia profunda SM-A87 |
| 559307 | Zygosaccharomyces rouxii CBS 732 |
| 33074 | Zymobacter palmae |
| 555217 | Zymomonas mobilis subsp. mobilis ATCC 10988 |
| 627344 | Zymomonas mobilis subsp. mobilis ATCC 29191 |
| 622759 | Zymomonas mobilis subsp. mobilis NCIMB 11163 |
| 1194424 | Zymomonas mobilis subsp. mobilis NRRL B-12526 |
| 627343 | Zymomonas mobilis subsp. mobilis str. CP4 = NRRL B-14023 |
| 627343 | Zymomonas mobilis subsp. mobilis str. CP4 = NRRL B-14023 |
| 264203 | Zymomonas mobilis subsp. mobilis ZM4 = ATCC 31821 |
| 579138 | Zymomonas mobilis subsp. pomaceae ATCC 29192 |
| 336722 | Zymoseptoria tritici IPO323 |
